# Connectome-driven neural inventory of a complete visual system

**DOI:** 10.1101/2024.04.16.589741

**Authors:** Aljoscha Nern, Frank Loesche, Shin-ya Takemura, Laura E Burnett, Marisa Dreher, Eyal Gruntman, Judith Hoeller, Gary B Huang, Michał Januszewski, Nathan C Klapoetke, Sanna Koskela, Kit D Longden, Zhiyuan Lu, Stephan Preibisch, Wei Qiu, Edward M Rogers, Pavithraa Seenivasan, Arthur Zhao, John Bogovic, Brandon S Canino, Jody Clements, Michael Cook, Samantha Finley-May, Miriam A Flynn, Imran Hameed, Alexandra MC Fragniere, Kenneth J Hayworth, Gary Patrick Hopkins, Philip M Hubbard, William T Katz, Julie Kovalyak, Shirley A Lauchie, Meghan Leonard, Alanna Lohff, Charli A Maldonado, Caroline Mooney, Nneoma Okeoma, Donald J Olbris, Christopher Ordish, Tyler Paterson, Emily M Phillips, Tobias Pietzsch, Jennifer Rivas Salinas, Patricia K Rivlin, Philipp Schlegel, Ashley L Scott, Louis A Scuderi, Satoko Takemura, Iris Talebi, Alexander Thomson, Eric T Trautman, Lowell Umayam, Claire Walsh, John J Walsh, C Shan Xu, Emily A Yakal, Tansy Yang, Ting Zhao, Jan Funke, Reed George, Harald F Hess, Gregory SXE Jefferis, Christopher Knecht, Wyatt Korff, Stephen M Plaza, Sandro Romani, Stephan Saalfeld, Louis K Scheffer, Stuart Berg, Gerald M Rubin, Michael B Reiser

**Author notes:** Contributed equally (Frank Loesche, Shin-ya Takemura, Laura E Burnett, Marisa Dreher, Eyal Gruntman, Judith Hoeller, Gary B Huang, Michał Januszewski, Nathan C Klapoetke, Sanna Koskela, Kit D Longden, Zhiyuan Lu, Stephan Preibisch, Wei Qiu, Edward M Rogers, Pavithraa Seenivasan, Arthur Zhao).

## Abstract

Vision provides animals with detailed information about their surroundings, conveying diverse features such as color, form, and movement across the visual scene. Computing these parallel spatial features requires a large and diverse network of neurons, such that in animals as distant as flies and humans, visual regions comprise half the brain’s volume. These visual brain regions often reveal remarkable structure-function relationships, with neurons organized along spatial maps with shapes that directly relate to their roles in visual processing. To unravel the stunning diversity of a complex visual system, a careful mapping of the neural architecture matched to tools for targeted exploration of that circuitry is essential. Here, we report a new connectome of the right optic lobe from a male *Drosophila* central nervous system FIB-SEM volume and a comprehensive inventory of the fly’s visual neurons. We developed a computational framework to quantify the anatomy of visual neurons, establishing a basis for interpreting how their shapes relate to spatial vision. By integrating this analysis with connectivity information, neurotransmitter identity, and expert curation, we classified the ∼53,000 neurons into 727 types, about half of which are systematically described and named for the first time. Finally, we share an extensive collection of split-GAL4 lines matched to our neuron type catalog. Together, this comprehensive set of tools and data unlock new possibilities for systematic investigations of vision in *Drosophila*, a foundation for a deeper understanding of sensory processing.

## Introduction

Over a century of anatomical studies have carefully cataloged many cell types in the fly visual system^1–3^, and in parallel, behavioral and physiological experiments have examined the visual capabilities of flies. A wealth of high-performance visual behaviors and genetic tools for targeted access to specific cell types have made *Drosophila* uniquely suited for studying the neural circuit implementation of many visual computations^4^. Recently, the anatomical understanding of the *Drosophila* visual system has been revolutionized by connectomics, the reconstruction of neuron shapes and synaptic connections from Electron Microscopy (EM) data^5–8^. The motion vision pathway provides an impressive example. Despite decades of systematic behavioral and physiological studies, detailed EM circuit reconstructions of just a small piece of the *Drosophila* visual system transformed the field by proposing several testable hypotheses about the mechanism of directional selectivity in the T4 and T5 neurons^7,9^. Connectome analysis revealed that these cells receive asymmetric synaptic inputs from distinct neuron populations at different positions along their asymmetrically shaped dendrites. This structural information, together with a near-complete list of relevant cell types, launched a decade of experiments aiming to uncover the mechanism of directional selectivity. This prior work clearly illustrates the crucial insights into function that can come from describing the shapes of neurons and their connections. A complete visual system connectome will bring this level of analysis and incisive experimental design to all areas of visual processing.

While understanding the function of a brain region is an ongoing challenge in neuroscience, there is little mystery about the role of visual brain areas—their neurons and circuits must be involved in seeing. This core insight guides the analysis of the neurons engaged in vision: these neurons should sample and represent information from across the field of view, this information should then be transformed by a series of steps that extract increasingly selective signals, and finally, these signals should be conveyed to higher brain areas. While the mammalian retina is the iconic example of this ground plan, the same organization is found in the *Drosophila* visual system and even in the reduced visual system of the micro-wasp *Megaphragma*^10^. Studies in the mouse retina have used anatomical and functional methods to sort the retinal ganglion cells and other neurons into a comprehensive inventory of more than 40 retinal ganglion cell types^11–13^. A critical property in defining retinal cell types is their arrangement into anatomical mosaics: cells of the same type are usually organized with approximately uniform coverage and evenly spaced to sample all points in the tissue, so there are no blind spots^12–14^. We take inspiration from these efforts in fly and mouse vision that have resoundingly demonstrated the power of careful cell typing for functional analysis.

With recent progress in the acquisition and analysis of *Drosophila* EM connectomes^6,15–17^ and Light Microscopy^18,19^ (LM) data, we are struck by the unique role the fly can play in the study of vision. In *Drosophila*, these studies can be built on a foundation of comprehensive knowledge of the morphology of the relevant neurons, their neurotransmitters, and their synaptic connections, as well as access to cell-type-specific genetic tools to manipulate them^20,21^. To date, the systematic EM studies of *Drosophila* visual neurons were all performed in female brains^7,9,22–25^. Here we introduce a new EM dataset and the first detailed examination of the visual system in a male brain, thereby establishing the most comprehensive survey of any insect optic lobe. This manuscript details this new dataset along with the cell typing of all the visual neurons and a matched collection of genetic drivers. We believe these insights and resources will empower an unprecedented, holistic understanding of vision.

### Neurons of the *Drosophila* visual system

The complete Central Nervous System (CNS), including the entire brain and ventral nerve cord of a male *Drosophila* was dissected, fixed, stained, and cut into 66 20-µm slabs, which were then imaged using Focused Ion Beam milling and Scanning Electron Microscopy (FIB-SEM). Subsequently, the entire volume was aligned, synapses were detected (Extended Data Fig. 1a), and automatic segmentation was applied to extract neuron fragments. Proofreading of the visual regions on one side has been completed (Extended Data Table 1), establishing a new connectome (available via neuPrint database, see Data Availability), while proofreading of the rest of the CNS is ongoing. We provide here a complete catalog of the neurons in this visual system (Fig. 1a,b, Supplementary Table 1). The process by which individual neurons were assigned to cell types is described in Fig. 2-4, and the complete quantitative inventory is provided in Supplementary Fig. 1.

**Fig. 1.**
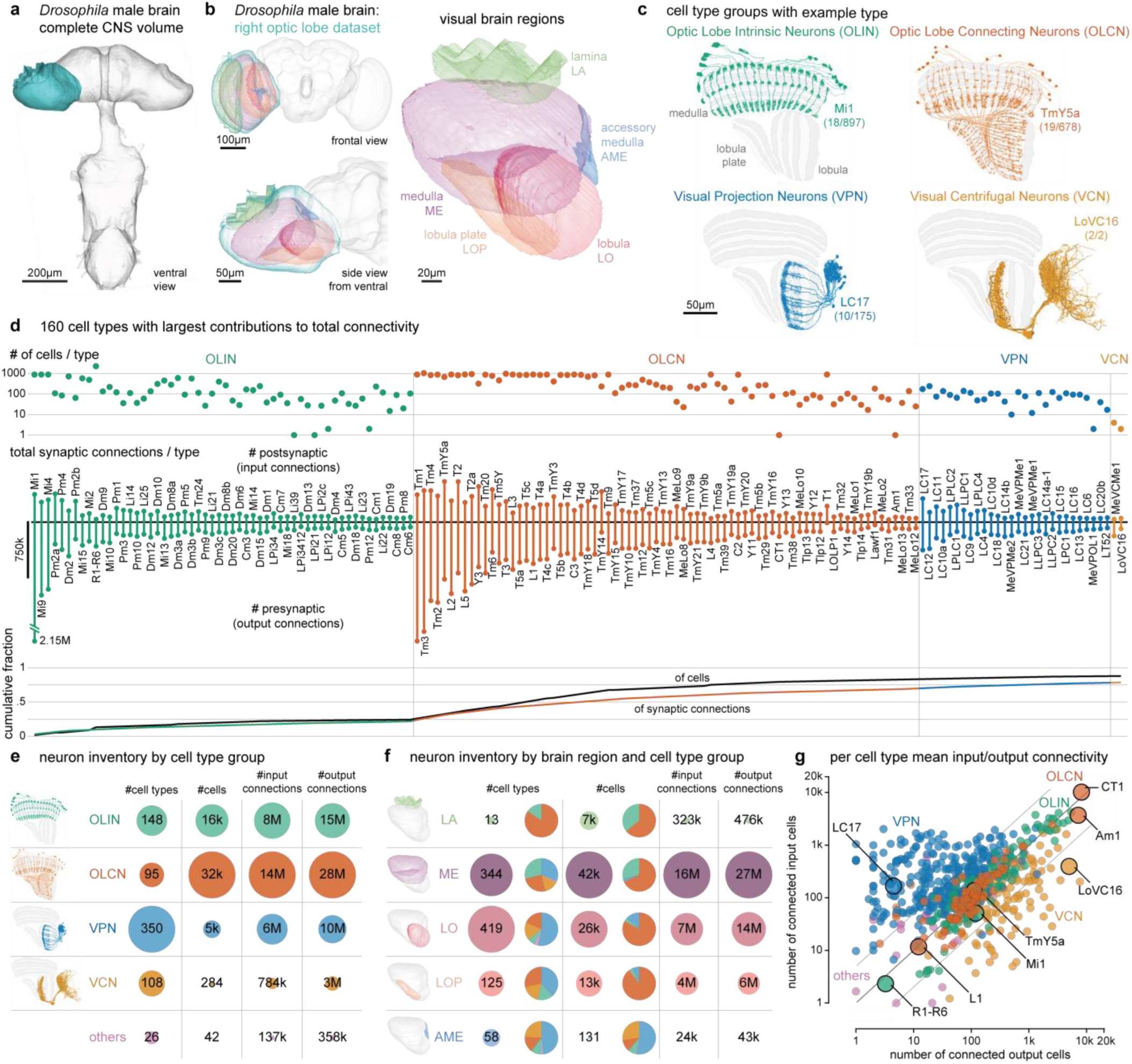
The neurons of the male *Drosophila* visual system. (a) Overview of the male central nervous system dataset, with the VNC attached. (b) This study describes the complete connectome and neuron inventory of the right optic lobe (blue). The optic lobe comprises five regions: lamina, medulla, accessory medulla, lobula, and lobula plate. (c) The 4 main groups of cell types with an example each (number cells shown / total cells of this type). (d) The number of cells and input and output synaptic connections for the 160 cell types with the largest contributions to the total connectivity in the visual system connectome are shown (all types in Supplementary Table 1). Fewer than 100 cell types account for most cells and connections. (e) Summarizing the inventory with the number of cell types, cells, and connections aggregated by cell type groups. The inventory includes a small group of “other” cell types with minimal connectivity. (f) Summary as in (e) but grouped by the five optic lobe brain regions. Counts include cell types and cells with >2% of their synapses contained within a region; many cells contribute to multiple regions. Contributions from the neurons in each cell type group are shown as pie charts. These counts summarize the connectome in the optic-lobe:v1.0 neuPrint database, and reflect the asymmetry between the completion percentage for presynapses and postsynapses (Extended Data Table 1). A few cell types are undercounted (an estimated 2,777 cells from the lamina and 459 R7/R8 photoreceptors and some Lat types in the lamina, see Methods). (g) Input/output connectivity in the optic lobe for all cell types in the inventory. The VPNs generally have more input cells than output cells, while the VCNs show the opposite connectivity pattern (excluding central brain connectivity). The example cell types from (c) and others with unusually high and low connectivity are highlighted.

**Fig. 2.**
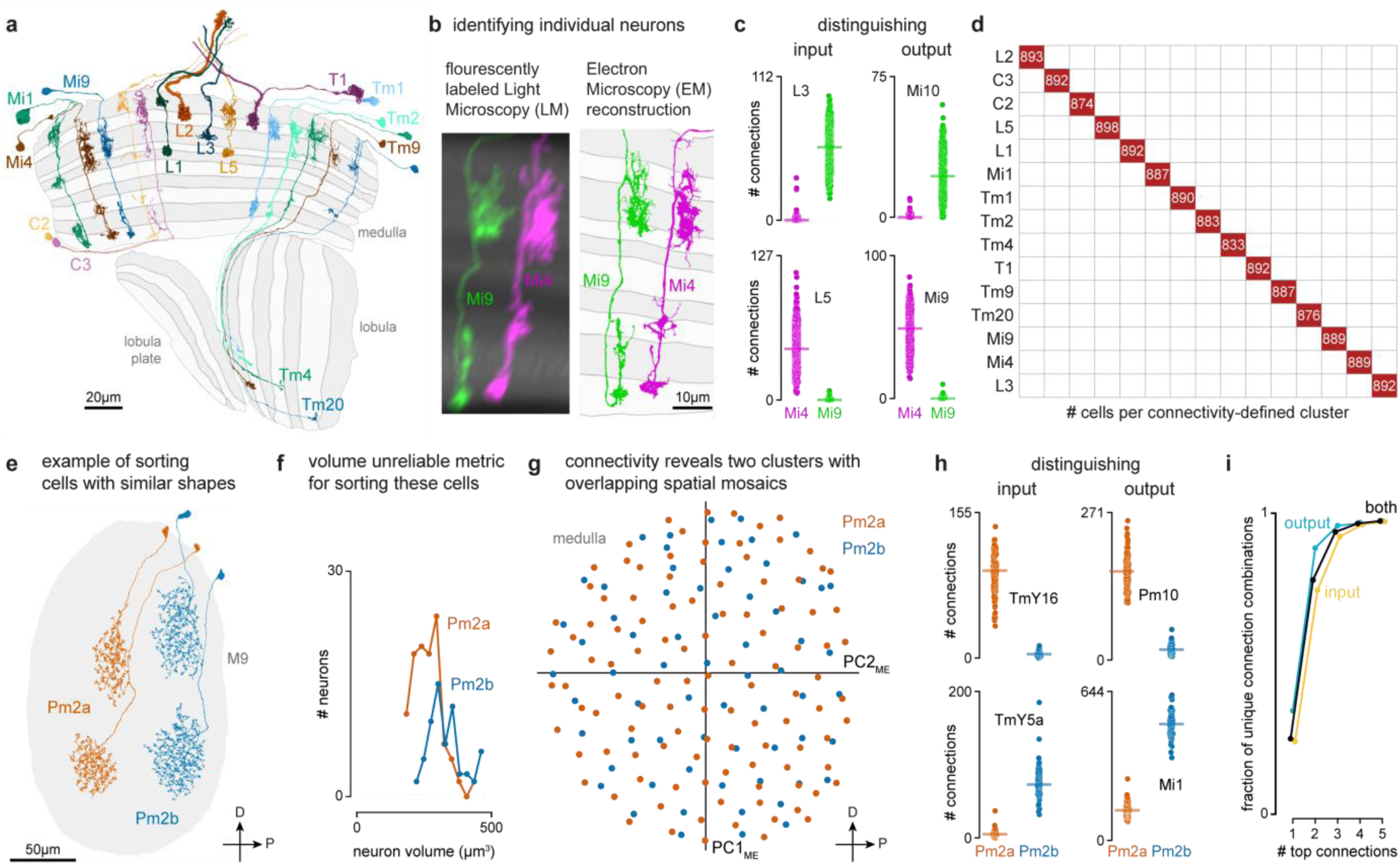
Sorting neurons into types based on morphology and connectivity. (a) Examples of 15 cell types that occur once per column in nearly every column of the visual system, shown across a slice of the optic lobe. (b) Mi9 and Mi4, shown as LM images (see methods) and EM reconstructions, are an example of neurons that appear similar but can be uniquely identified by morphology in nearly all cases. (c) Mi4 and Mi9 can be distinguished by connectivity, as shown for selected input and output cell types. (d) After cell typing all neurons, connectivity clustering sorts the cells assigned to the 15 columnar cell types in (a) into distinct groups, confirming their assignments. (e) Two examples of each of Pm2a and Pm2b neurons, medulla amacrine cells with very similar morphology. (f) The distribution of cell volume is similar for the two types. (g) Pm2a and Pm2b are sorted into two types by connectivity clustering (Extended Data Figs. 2,3), revealing 2 overlapping mosaics. The first two principal components of the centers of mass of their synapse locations are plotted. This visualization preserves the spatial relationships of the cells and aligns with major anatomical axes. (h) Selected distinguishing input and output connections for Pm2a and Pm2b cells. The combination of consistent connectivity differences with overlapping cell distributions supports the split into two types. (i) Most cell types can be distinguished by their strong connections. The proportion of unique combinations of connections across all types for the indicated number of top-ranked connections.

The fly visual system comprises several distinct anatomical regions: the lamina, medulla, accessory medulla, lobula, and lobula plate, forming the structure called the optic lobe. Homologous regions are found in visual systems across pancrustaceans^26^. These regions are referred to as neuropils, structures dense with the synaptic connections of dendrites and axons, and shown here as meshes (Fig. 1a,b) that were drawn using visible boundaries in the EM data. Similar to other invertebrate nervous systems, cell bodies are typically distributed surrounding the neuropils.

We first classified neurons based on their relationship to anatomical regions. Optic Lobe Intrinsic Neurons (OLINs) have synapses confined to a single region. Optic Lobe Connecting Neurons (OLCNs) connect two or more visual regions. A large and diverse set of neuron types connects the visual regions with the central brain. The Visual Projection Neurons (VPNs) primarily convey signals to the central brain, while Visual Centrifugal Neurons (VCNs) primarily convey central brain signals to the optic lobes. Fig. 1c shows a representative cell type from each group, visualized on a slice of the principal visual regions. Several important features of visual neurons are immediately apparent – many types form a set of repeating neurons that collectively cover large swaths of the visual brain regions with distinguishing innervation patterns that underlie their specific shapes. Nearly all connections between visual neurons are in the neuropils. However, we also found connections outside the neuropils, most frequently in the chiasmata that connect them (Supplementary Table 2, Extended Data Fig. 1d-f). For example, we find such connections in the outer chiasm involving nearly all lamina-associated neurons, extending a recent observation made for R7, R8 photoreceptors^24^.

We classified ∼53k visual system neurons into 727 cell types (Supplementary Video 1). The number of neurons that comprise a type varies widely, with 60 cell types accounting for over half of the total synaptic connections and ∼75% of the neurons. Fig. 1d highlights the 160 cell types with the largest contributions to total connectivity in the visual system connectome, including many well-studied neurons of the motion vision pathway^9,27–29^: L1, L2, L3, L5, Mi1, Mi4, Mi9, Tm1, Tm2, Tm3, Tm4, Tm9, and all 8 T4 and T5 types. It may be that the need for fast, high-resolution motion signals requires many cells and synapses. Moreover, this unbiased connectivity survey reveals many cells with substantial contributions to the network that have received minimal attention, including many Pm and TmY cell types. The inventory is summarized in Fig. 1e-f, which describes the breakdown of cell types by group and brain region. We include 26 cell types not placed in one of the four main groups, primarily central brain cell types with some limited (<2%) connectivity in visual brain regions.

The connectome contains ∼49 million connections in the visual brain regions. Fig. 1g and Extended Data Fig. 1b,c summarize the input-output connectivity of all cell types, color-coded by their group. The number of connected cells in the network spans five orders of magnitude across the visual cell types. Individual R1-6 photoreceptors and Lamina Monopolar Cells (e.g. L1) are connected to just a few other cells, while on the other extreme, several giant individual inhibitory interneurons, such as Am1 and CT1 are connected to >10,000 other cells.

Our naming scheme, explained in Methods, aims to be systematic while respecting well-established historical names. We introduced new, short names for all newly described cell types. For example, for VPNs and VCNs, the names are based on the principal neuropil, with an appended VP or VC, and a unique number. Where new evidence convinced us a previously defined cell type should be divided into several distinct types, we append a letter.

### Cell typing visual neurons

To assign cell types, we adopted a practical cell type definition that directly follows the pioneering *C. elegans* connectome^30^, the mammalian retina^12,13^, and decades of work on fly visual neurons^3,5,7,9,23,31^. We grouped neurons sharing similar morphology and connectivity patterns into types, with the aim that all neurons assigned to a type are more similar to each other than to any neurons assigned to different types. We generated type annotations in an iterative process that combined visual inspection of each cell’s morphology with computational methods for grouping cells by connectivity. Nearly all neurons in the dataset were reviewed multiple times by several team members as part of this process.

We started by naming easily recognizable, often well-known neurons, building on decades of LM studies^3,18,19^, prior EM work^7^, and our unpublished LM analyses. These neurons included the 15 columnar cell types in Fig. 2a, which are present in nearly every retinotopic medulla column (Supplementary Video 2). Each of these cell types can be identified by morphology; even cells such as Mi4 and Mi9 that appear similar are distinguishable at LM and, even more clearly, at EM resolution (Fig. 2b). Importantly, Mi4 and Mi9 neurons can be independently classified by connectivity, even when only considering a few strongly connected input and output types (Fig. 2c). Such connectivity differences can often only be assessed once connections are combined for all neurons of a cell type, because two putative type A neurons may have synaptic connections with cells of type B, but not the same individual type B cells. For this reason, we employed an iterative process in which neurons are first preliminarily typed to be used in connectivity (by type) analyses, which then confirm or refute the initial type assignment. In practice, comparisons of connectivity, including connectivity-based clustering, are already useful when only a minority of neurons is named, with increasing accuracy as more cells are typed. Indeed, with our current cell typing, the 15 columnar neurons are sorted perfectly into unique, connectivity-defined clusters (Fig. 2d).

When applying connectivity-based clustering with a preselected number of clusters to larger groups of cell types (e.g. medulla intrinsic cells, Extended Data Fig. 2), we encountered cases of (putative) cell types that clustered with different cell types or were split into multiple types. Such cases were resolved using additional criteria, such as cell morphology or, when available, genetic markers. For example, Dm6 and Dm19 cells have similar connectivity but different arbor sizes and soma locations (often indicative of their development^32^; Extended Data Fig. 2a,b) and are labeled by genetic driver lines with expression restricted to only one type. Our collection of cell-type-specific driver lines, discussed below, provided important clues about cell typing for dozens of cell types.

Another important criterion we used when deciding whether to split or merge cell types is spatial coverage. Like the cell types of the mouse retina, the populations of repeating neurons in the fly visual system also form mosaics achieving uniform coverage. To illustrate this biologically and methodologically important property, we consider two related amacrine-like cells of medulla layer M9, Pm2a and Pm2b. While Pm2a and Pm2b have similar morphologies (Fig. 2e,f), we found clear evidence for two populations that each separately cover visual space and have clear connectivity differences (Fig. 2g,h). We used similar analyses of repeating neurons to decide whether to divide groups into finer type categories (Extended Data Fig. 3).

Some challenging cases could only be resolved by using multiple types of information: connectivity, morphology, size, spatial distribution, and cell body position. For example, in a previous analysis of the R7 and R8 photoreceptors and their targets in the pathways that process color information, we found that individual neuron morphologies were idiosyncratic, such that we could not confidently sort all Tm5 neurons into types^24^. The completeness of the new connectome data overcomes these issues; we identified independent sets of connected neurons that allowed us to separately cluster Tm5a, Tm5b, and Tm29, as well as Dm8a and Dm8b, into overlapping mosaics (Extended Data Fig. 4).

Our cell clustering demonstrates how connectivity places tight constraints on cell typing. We found that >99% of cell types can be distinguished by their top five connections (Fig. 2i). Although this method does not robustly classify individual neurons, it captures essential properties of cell types and is included in our cell type summaries (Fig. 5, Supplementary Fig. 1).

### Visual system architecture from connectivity

The regions of the fly visual system, like the mammalian neocortex, have been further divided into layers wherein connections between distinct neuron populations occur. Taken together, these layers and the approximately orthogonal columns that convey retinotopy provide a representation of visual space and a biologically meaningful framework for describing visual neurons. Indeed, the distributions of arbors of many fly visual neurons have been shown to provide strong clues about their function^7,9,33,34^. Although layered structures can be seen with LM^35^, they are abstractions that emerge from the collective organization of millions of connections between cells. We, therefore, developed a computational infrastructure using principal neurons as scaffolds to delineate the visual system’s layers and columns.

We first built a coordinate system in the medulla. We assigned each of the 15 columnar neurons of Fig. 2a to hexagonal coordinates using the approach we developed in the FAFB dataset^36^. The coordinates correspond one-for-one to lenses on the compound eye (except for some lenses on the edge). We also identified coordinates along the eye’s equator, a global anatomical reference (Fig. 3a). A complete set of the columnar neurons was assigned to most medulla coordinates, while we found some incomplete sets, mostly at the margins of the medulla (Fig. 3a, Supplementary Table 3).

**Fig. 3.**
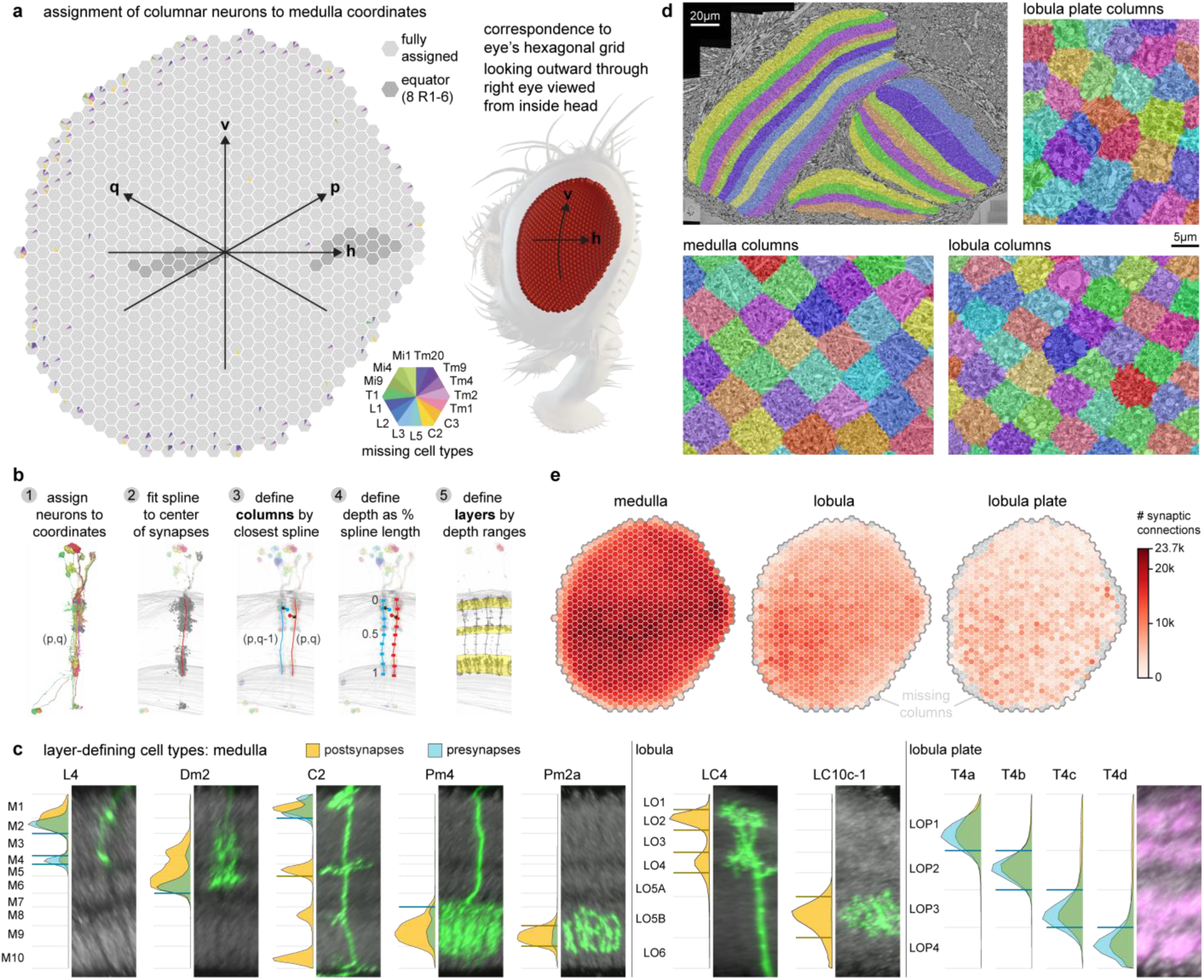
Capturing the architecture of the visual system by analyzing the connectivity of key cell types. (a) The lenses of the fly eye form a hexagonal grid and are mapped onto a hexagonal coordinate system in the medulla. The darker hexagons correspond to locations along the eye’s equator (determined by counting photoreceptors in the corresponding lamina cartridges). The 15 columnar neuron types in Fig. 2a were assigned to each coordinate. Full gray hexagons indicate a medulla location that has been assigned a set of all 15 neurons (see Methods, Supplementary Table 3). The color-coded wedges indicate the cells of a type missing at that medulla location, most of which are along the edge. (b) Process for creating columns and layers. Lobula plate columns were based on sets of T4 neurons assigned to Mi1s (Extended Data Fig. 5). (c) Layer boundaries defined based on synapse distributions of “marker" cell types, established from LM images. For each type, we show the distribution of pre/post synapses across depth together with LM images of single-neuron clones (in green; rotated and rescaled to match top and bottom of the neuropil in light grey). The lobula plate image shows the neuropil (grey, nc82-antibody) and the axon terminals of the T4 neurons (magenta). The horizontal blue/orange lines indicate the distribution cutoff around a peak that defines a boundary. The collection of these defines all layer boundaries, shown in gray. (d) The layers and columns are shown as volumes superimposed on the grayscale EM data. (e) The spatial distribution of postsynapses, by region and column (see also Extended Data Fig. 6).

We then used the synapses of all neurons assigned to each medulla coordinate to create volumetric columns for the medulla and lobula (Fig. 3b, Extended Data Fig. 5a). Lobula plate columns were built by assigning sets of T4 neurons to each Mi1 in the medulla (Extended Data Fig. 5, Supplementary Table 4), which extends the same coordinate system to all three major visual regions. Volumetric layers were constructed by bounding depth positions along columns, calibrated using benchmark cell types (visualized alongside LM examples, Fig. 3c). These volumes establish an “addressable” visual system within the accompanying neuPrint database. They can be visualized alongside the gray-scale EM data (Fig. 3d) or reconstructed neurons (Extended Data Fig. 5c,d).

These tools enable database queries that simplify otherwise complex data analyses, such as the spatial distribution of synaptic connections across the visual regions (Fig. 3e), or as a function of depth (Extended Data Fig. 6). Most significantly, the columns and layers facilitate quantification of connectivity and morphology, which form the core of our comparative dataset (Fig. 5).

### Neurotransmitters in the visual system

An essential component of a complete description of the fly visual system’s connectome is the neurotransmitters each neuron type uses to communicate across synapses. The functional sign of each synaptic connection, whether excitatory or inhibitory, depends primarily on presynaptic neurotransmitter expression, and on the receptors expressed by postsynaptic cells. Identifying neurotransmitters expressed by each neuron has been challenging and imprecise, relying on genetic techniques like reporter expression or using antibodies for labeling transmitters or their synthesizing enzymes. Moreover, we do not have robust methods for detecting neuromodulation via peptides^37^.

Recently, more reliable and scalable methods have been developed for assigning neurotransmitters to neurons. Detecting expression of neurotransmitter-synthesizing enzymes within cells using Fluorescence in situ hybridization (FISH)^38,39^ or RNAseq^40–42^ has provided transmitter identity for dozens of cell types in the fly visual system. Although EM data do not give direct information about transmitter identity, recent progress in applying machine learning to classify neurotransmitters from small image volumes around presynapses (Fig. 4a) has shown impressive accuracy^43^. We were therefore excited to apply these methods to the visual system, where the many repeating cell types and substantial prior data on neurotransmitter expression provide an unprecedented scale of training data and means to evaluate the accuracy of these predictions.

**Fig. 4.**
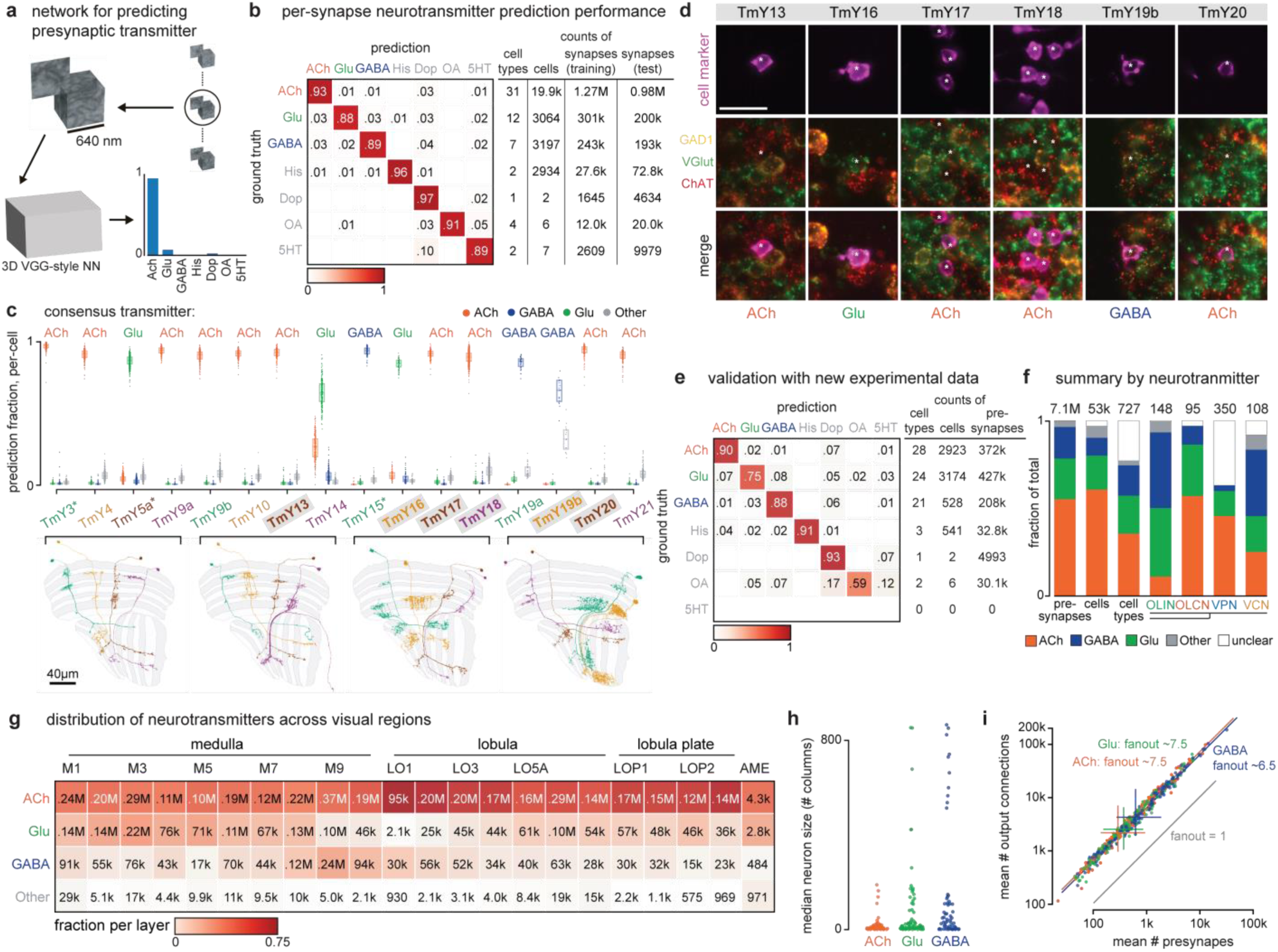
Diversity of neurotransmitter signaling in the visual system. (a) Using prior neurotransmitter expression measurements in 59 cell types as ground truth data, a VGG-style deep neural network^43^ was trained to classify presynaptic transmitter. (b) Performance of the per-(pre)synapse predictions evaluated on held-out synapses; cell types, cells, and synapses used for training and testing are tabulated. (c) Synapse predictions are aggregated for cell-type-level consensus transmitter assignments, shown for 16 TmY cell types; example morphologies below. Three cell types (asterisks) were included in training data, predictions for six types (names in gray boxes) were confirmed by new validation data. (d) Images of driver lines for those six TmY types that were assayed for neurotransmitter marker gene expression^38,39^, showing GAL4 cell marker (top), markers for ChAT, VGlut, and GAD1 (middle), and merged images (bottom), and the assigned transmitter below. (e) Performance of neurotransmitter predictions for 79 cell types evaluated with experimental data (Supplementary Table 5). (f) Neurotransmitter types counted for synapses, cells, cell types, and cell type groups. In these and following summaries, synapse-level predictions are inherited from consensus (cell-type-level) transmitters. Cells with limited predictions scored as “unclear.” (g) Spatial distribution of presynapses with neurotransmitter classifications; color scale normalized for each column. (h) Median size for OLIN and OLCN cell types. (i) Linear regression estimates of synaptic fanout, the cell-type averaged ratio of output connections to presynapses, grouped by transmitter. The fanout values are significantly different between ACh and GABA (p-value < 0.001) and Glu and GABA (p-value < 0.001), but not between Ach and Glu (p-value ∼0.64), ANCOVA.

We trained a neural network to classify synapses into one of seven transmitters (acetylcholine, glutamate, GABA, histamine, dopamine, octopamine, and serotonin) using the known transmitters of 59 cell types, contributing nearly 2 million synapses (Fig. 4b). The per-synapse accuracy of the trained network showed a modest improvement compared to predictions for the hemibrain and FAFB datasets^43^. To further validate the predictions, we collected new data on neurotransmitter expression for 66 additional cell types, primarily using EASI-FISH^38^.

As an example, we show the neurotransmitter predictions for a complete anatomical group of neurons, the TmY cells (Fig. 4c). Three of the 16 cell types (indicated by asterisks) were included in the training data, while the neurotransmitters expressed by six of these types are covered in our new, independent experimental data (labels on gray in Fig. 4c, experimental data in Fig. 4d). The fraction of the top transmitter prediction is high; 14/16 TmY cell types have >85% of synapses classified as one neurotransmitter. The consistency or accuracy of predictions is comparable between the types in the training dataset and cell types not used in training. Our experimental expression data were consistent in all six cases with the top predictions.

The prediction accuracy is further improved when the single-synapse predictions are aggregated across the cells of each type, as shown for our validation dataset of 79 cell types not included in training the neural network (Fig. 4e, Supplementary Table 5). Despite this impressive performance, there are limitations to the reliability of these predictions. Many cell types, including >100 VPN types, have limited presynapses in our data volume. Another ∼50 cell types, all with low cell counts, have predictions for octopamine, dopamine, or serotonin, which far exceeds the number of optic-lobe-associated cell types previously estimated to express these transmitters^39^. Precisely because of the scarcity of neurons confirmed to express these transmitters, they are not well represented in our training data, and accordingly, these predictions are less reliable. For cell types with low confidence predictions, unless independently confirmed by experimental data, the consensus transmitter is reported as “unclear” in the neuPrint database.

Applying the consensus neurotransmitters to the synaptic connectivity data provides new insights (Fig. 4f-i). Approximately half of the cell types express acetylcholine and are likely excitatory, and half express GABA or glutamate and are likely inhibitory (Fig. 4f). Excitatory neurons make up a majority of the region connecting OLCN and VPN types, while the OLIN and VCN types are heavily skewed towards glutamatergic, GABAergic, and aminergic cells (Fig. 4f). There are also distinct differences in neurotransmitters between the layers in each region (Fig. 4g). Furthermore, the largest neurons tend to be glutamatergic or GABAergic, and thus likely inhibitory (Fig. 4h). We also examined the synaptic “fanout” of visual system neurons, i.e., the average ratio of output connections to presynapses. These data show a remarkably consistent relationship, with a significant and potentially interesting difference across the neurotransmitters—the average fan-out of GABAergic neurons is lower than glutamatergic or cholinergic neurons, participating in ∼13% fewer connections (Fig. 4i).

Most neurons appear to release the same neurotransmitter(s) across their various terminals, and molecular profiling suggests that most neurons signal with a single dominant neurotransmitter. However, there are two clear examples of co-transmission in the visual neurons: Mi15s express the markers for both dopamine and acetylcholine, while R8 (pale and yellow) express the markers for histamine and acetylcholine^40^ (Supplementary Table 5). Several visual neurons are peptide-releasing cells, such as the l-LNVs^44^, and EM-based prediction methods have not yet been developed for peptidergic transmission.

### Quantified anatomy and connectivity

The shapes of neurons are varied and complex, but by representing visual brain regions as a coordinate system of layers and columns (Fig. 3), we could bypass much of this complexity to provide a compact, quantitative summary of innervation patterns and connections. Like a fingerprint, these summaries uniquely identify most cell types and are broadly applicable—for comparisons between cells and cell types within our dataset, with neurons imaged with LM (see Fig. 8), and for neurons in other EM volumes. These comparisons can be extended to other insect species due to the deep conservation of the optic lobe ground plan.

**Fig. 5.**
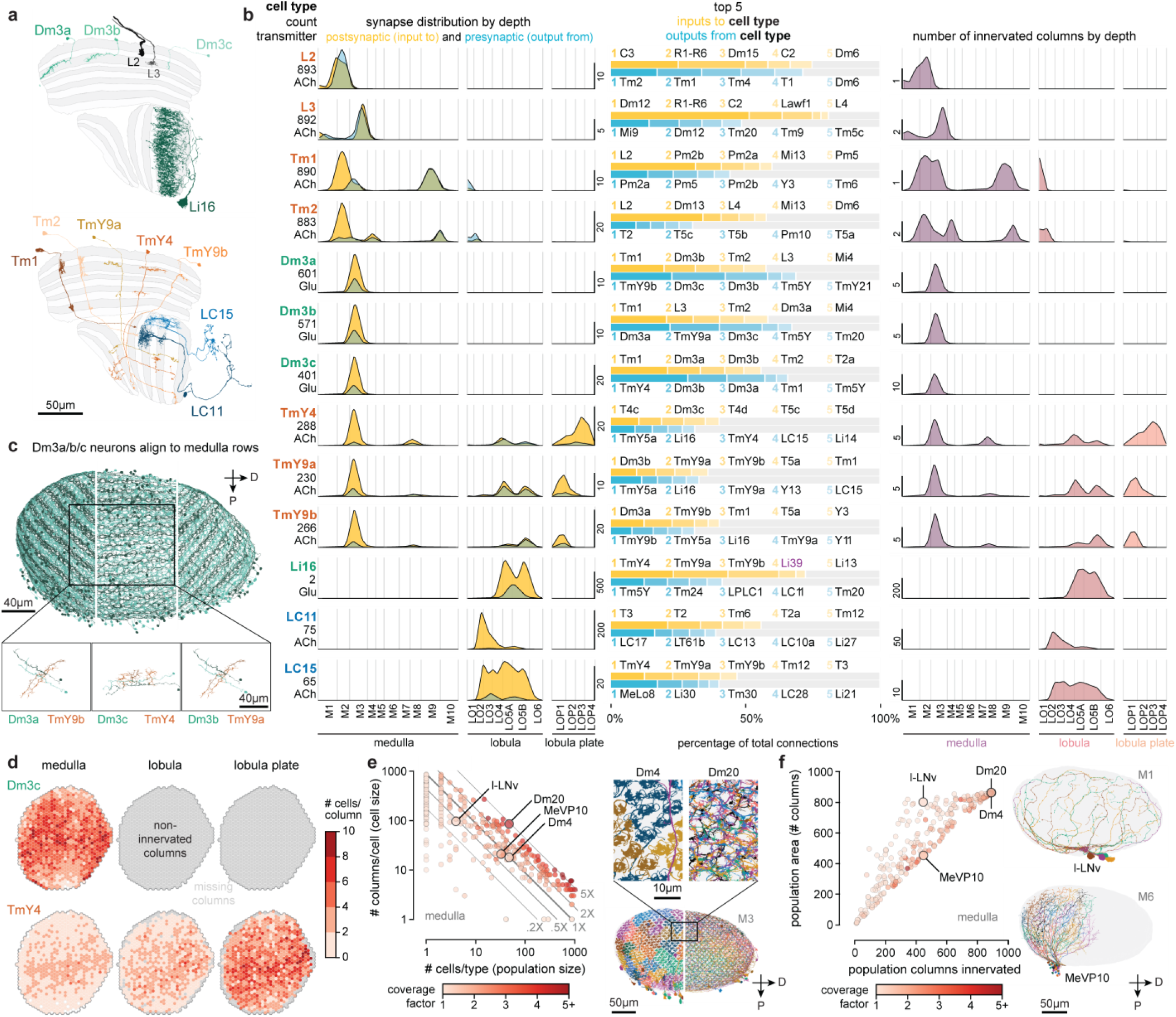
Quantitative summary of anatomy and connectivity of visual system neurons. (a) Examples of 13 cell types associated with the Dm3 “line amacrine” neurons, including their major inputs and outputs. (b) Each row presents the cell type’s quantified summary. Number of neurons and consensus neurotransmitter prediction (left-most). Distribution of presynapses and postsynapses across visual region layers (left), top five connected input and output cell types by contributed connectivity (color shade indicates rank; center), and cell size measured by depth (mean number of innervated columns, right). The visual cell types are summarized in The Cell Type Catalog (Supplementary Fig. 1). (c) The stripe-like patterns of Dm3 types each cover all M3, shown here in separate panels with neurons colored by their dominant column coordinate. The inset shows two individual Dm3s from adjacent coordinates and selected connected TmYs.(d) The number of Dm3c and TmY4 cells innervating each column as a spatial distribution by region. Similar plots for all visual cell types are in the Cell Type Explorer web resource. (e) The relationship between the number of cells (population size) and the average number of columns innervated by each cell (cell size) within the medulla (for types with >50 synapses and >5% of total connectivity therein). Color-coded by the coverage factor (per-type average number of cells per column). The 1X line indicates one-fold coverage: cell types above/below cover the whole medulla with more/less neurons per column. Selected cell types show coverage factors for different tiling arrangements: Dm4 (∼1) and Dm20 (∼5). (f) The per cell type density of medulla coverage, comparing the population’s column innervation to its convex area. Types close to the diagonal (e.g., MeVP10) densely cover the medulla, types above the diagonal (e.g., l-LNv) feature sparser coverage. Medulla layers shown face-on.

We illustrate these summaries with a fascinating group of connected neurons that includes the three Dm3 types (Fig. 5a,b). Each Dm3 type forms a striking pattern of stripes across the medulla (Fig. 5c). The stripes of each type have a distinct orientation across the medulla and each type provides substantial input to a distinct TmY type, that itself shows oriented medulla branches that are approximately orthogonal to those of the upstream Dm3 type (Fig. 5c, lower). Although prior LM studies have described Dm3 cells, also known as line amacrines, and their oriented arbors^2,3,19^, only the comprehensive EM reconstruction and annotation reveal this network in full detail. While we focus here on presenting our cell type inventory of the visual system, we note that the intriguing structure of the Dm3/TmY network has strong implications for function, as others have also recently noted^45^.

To make it easier to explore the circuits in our visual system inventory, we created compact summaries of many defining features of each cell type (example for Dm3/TmY network in Fig. 5b). The summaries present the number of individual cells of that type, the consensus neurotransmitter prediction, the mean distribution of presynapses and postsynapses (left), the top five connected input and output cell types (middle), and the mean size of the neuron as numbers of innervated columns (right). The top five input and output connections of the example neurons already capture key features of this circuit: each Dm3 type receives its main excitatory input from Tm1 neurons and provides glutamatergic (likely inhibitory) output to the other two Dm3 types, and one of the three TmYs. We were particularly intrigued to find that the three TmY types are the top inputs to LC15, a VPN with selective responses to moving long, thin bars^46^. A different VPN with a preference for small objects, LC11^47^, is instead a target of a large glutamatergic interneuron (Li16), which is another major target of the TmY cells, a potential opponent processing step that tunes the selectivity of these feature-detecting LC neurons.

The detailed morphology of the neurons was quantified as the synaptic distribution and column innervation as a function of depth within visual regions. The classic layer boundaries are shown as a reference, but the data are presented with higher depth resolution since it is clear these established neuroanatomic features do not fully capture the organization. An essential test of whether these measurements accurately profile each cell type’s distinctive morphology is to see if they can be used to sort cells. Indeed, by compiling synapse-by-depth measurements into a per-neuron feature vector, we find that most medulla neurons are sorted into their respective types (Extended Data Fig. 7), and this sorting only slightly underperforms connectivity-based clustering (cluster completeness 0.88 vs. 0.9; Extended Data Fig. 2); however, this method does not require pre-assigning cell types beyond those used to establish the columns. The cell type summaries, along with a gallery of example neurons of each of the 727 types, can be found in Supplementary Fig. 1. We developed a set of interactive webpages that summarize all visual neurons’ connectivity and serve as a browsable companion to the neuPrint web interface. The *Drosophila* Visual System Cell Type Explorer is previewed in Extended Data Fig. 8.

To survey the distribution of the neurons and connections of each cell type we analyzed their spatial coverage across visual regions (summed across layers, Fig. 5d). The example spatial coverage maps for Dm3c and TmY4 in the medulla exhibit a common trend of somewhat higher density of neurons in the area near the eye’s equator. Dm3c is a medulla intrinsic cell without processes in other regions. The coverage by TmY4 shows a noteworthy trend towards higher overlap in the lobula plate (the Cell Type Explorer web resource contains these maps for all cell types). This spatial analysis enables measurements of how neurons cover their retinotopic brain regions, directly linked to their sampling of visual space. Fig. 5e shows the relationship between the number of cells per type, the size of each type in columns, and the coverage factor (average number of cells/column) for the medulla (lobula and lobula plate plots in Extended Data Fig. 9; interactive plots on the Cell Type Explorer web resource). As expected, we find a prominent trend in which the number of cells of a type is inversely related to their size, consistent with the uniformity of coverage property of visual neurons. However, interesting differences emerge within this trend, such that cell types above the “1X” line have higher coverage factors, with more cells than are strictly necessary to cover the region. For instance, there are exactly 48 cells of each of Dm4 and Dm20, but individual Dm20 cells are much larger, featuring higher coverage. Dm4 neurons neatly tile medulla layer M3, while Dm20 cells feature substantial overlap with nearly ten cells per column. Neurons below the “1X” line have partial coverage of the medulla, but this coverage can result from two manifestations: either regional but dense coverage, illustrated by MeVP10s, or broad but patchy coverage, illustrated by l-LNvs (Fig. 5f, Supplementary Video 3). These summary data, along with the complete inventory detailed in Supplementary Fig. 1 and the Cell Type Explorer web resource, provide a comprehensive starting point for circuit discovery and experiment design.

### Specialized cell types

Our complete survey also allows us to highlight specialized sets of visual neurons defined by their anatomy and expected functional roles. The Dorsal Rim Area (DRA) is a zone of the eye whose photoreceptors are specialized for detecting polarized light^24,48^. Specialized medulla neurons integrate these R7d and R8d photoreceptors (Extended Data Fig. 10). Interestingly, we did not find DRA-specific OLCNs and, therefore, no specialized cells in the lobula.

The Accessory Medulla is a small brain region at the anterior-medial edge of the medulla, with a well-established role in clock circuitry and circadian behaviors^44,49^. Unlike the other visual brain areas, the AME does not exhibit obvious retinotopy, which supports a common view that photoentrainment of circadian rhythms should not require detailed visual-spatial information. Nevertheless, the diversity of neurons with significant connectivity in the AME (Extended Data Fig. 11) suggests an underappreciated complexity with perhaps broader roles in sensory integration and behavior.

### Communication with the central brain

The Visual Projection Neurons (VPNs) represent a remarkable compression of information. Signals from the nearly 50,000 local neurons that initially interpret the visual world are conveyed by just ∼4,500 neurons of 350 cell types to the central brain (Fig. 1e, Supplementary Table 1). Many VPN types, including the Lobula Plate Tangential, LPT^33^, the Lobula Columnar, LC^47^, and the Medulla-Tubercle, MeTu^50^, neurons have been carefully described. Nevertheless, we have identified novel cell types, and further refined prior classifications within each of these well-studied groups. Among the VPNs, the small field projections neurons of the lobula and lobula plate, the LCs, LPCs, LLPCs, and LPLCs make up ∼3000 cells, while the MeTu neurons make up over 500 more. The remaining cells are morphologically varied and target many brain regions. We present all VPNs in Fig. 6 to illustrate the diverse pathways relaying vision to central behavioral control areas. The neurons are grouped based on rough anatomical similarities, mainly the visual region in which they receive inputs. We refer readers to the summaries in Supplementary Fig. 1 for details of the connectivity and morphology within visual brain regions.

**Fig. 6:**
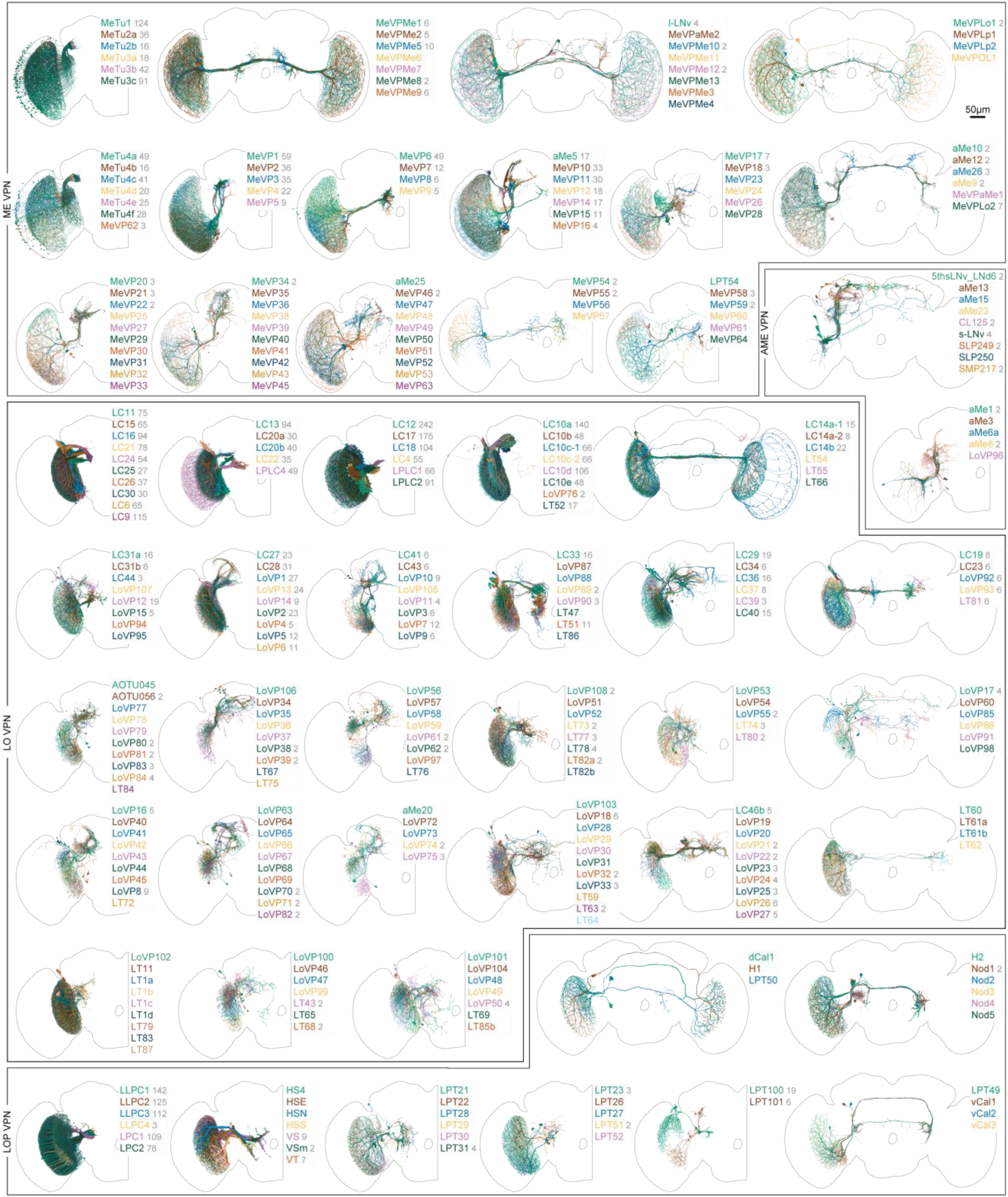
Visual Projection Neurons. All Visual Projection Neurons that connect the right optic lobe with the central brain or the contralateral optic lobe are shown. We first divided the ∼4500 VPNs into the 350 types shown here, and then placed them into 51 groups of morphologically similar types. These groups are based on the main region they receive their optic lobe inputs in, whether they project to the ipsi- or contralateral central brain or the contralateral optic lobe, and other aspects of their morphology. Unilateral cells, those whose projections do not cross the midline, are mainly shown in half-brain panels, and most cells with arbors in both brain hemispheres are shown in full-brain views. Each panel includes the names of the cell types (color-matched to the rendered neurons) and the number of individual cells of each type (in gray); types without numbers are present once per brain hemisphere. Detailed morphology within the visual brain regions can be found in the Cell Type Catalog (Supplementary Fig. 1).

The Visual Centrifugal Neurons (VCNs), 108 types across ∼270 total cells, are the final group of neurons we present as a complete set (Fig. 7). Most of these types consist of a single neuron, although there are several populations of smaller field inhibitory neurons mainly targeting the lobula plate. Among the larger VCNs are several octopaminergic cells^51^ that are likely critical for modulating visual neurons during active movement^52,53^. Many VCNs are large cells that target rather precise layers of the medulla (mostly called the MeVC neurons), the lobula (the LoVC and some LT neurons), and the lobula plate (called LPTs for historical reasons). One of these types, LoVC1 (also called IB112^6^) has recently been reported to gate the flow of specific channels of visual information during social behaviors^54^. Although most of VCNs are presumed to be inhibitory (glutamatergic or GABAergic), it is noteworthy that most of the MeVC neurons are predicted to be cholinergic. We present this full set as a single unit (Fig. 7) to serve as a reference to these understudied cells and to facilitate the systematic study of their roles in the central modulation of visual processing.

**Fig. 7:**
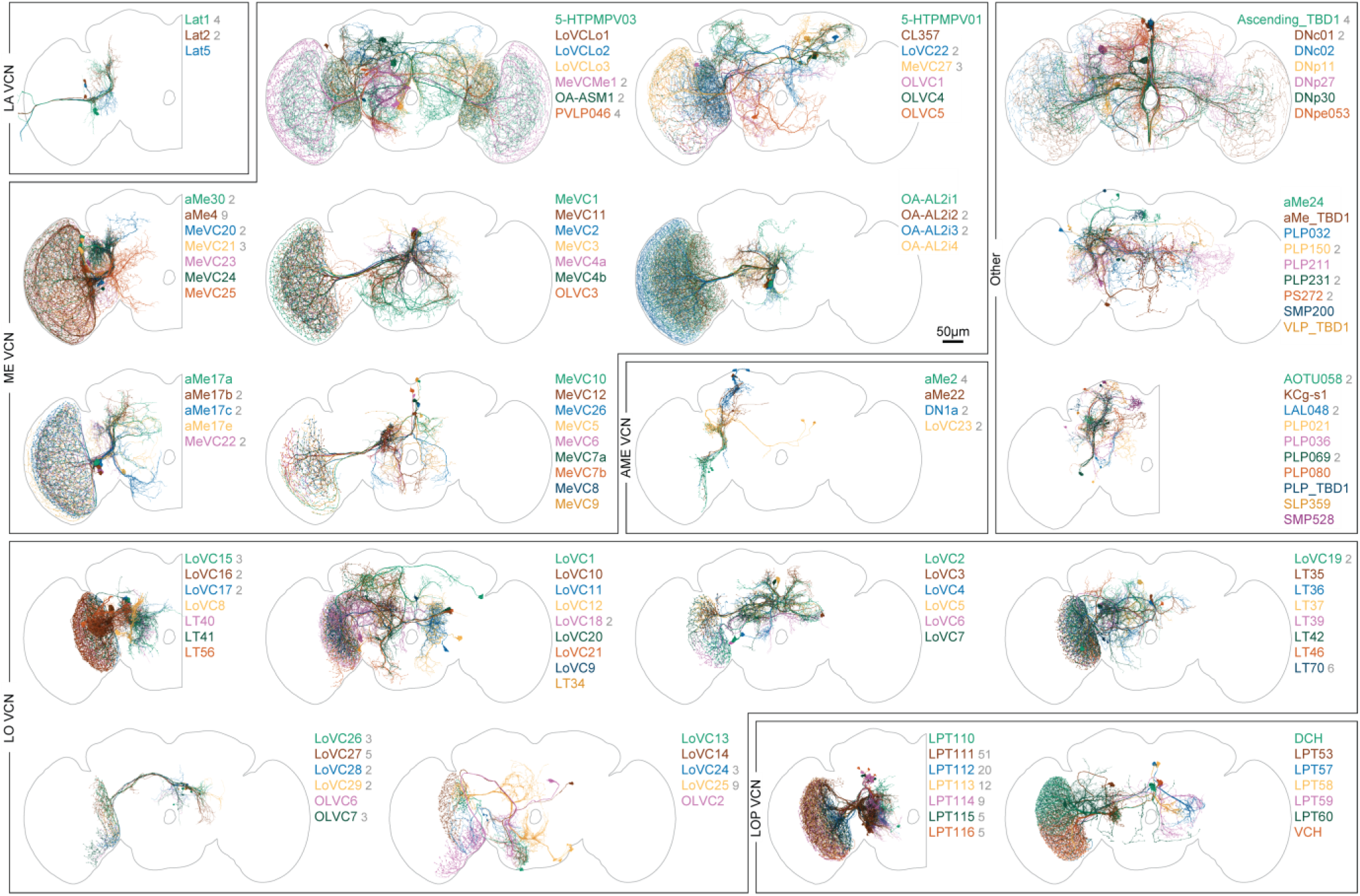
Visual Centrifugal Neurons. Visual centrifugal neurons receive major input in the central brain and project back to the optic lobes. We cataloged 108 VCN types (284 cells combined for the right optic lobe) and show them all here in groups organized by their main target regions in the optic lobes and other anatomical features (such as ipsi-, contra- or bilateral projection patterns). The figure also includes neurons (“other”) that have some optic lobe synapses in addition to central brain synapses but were not classified as VPN or VCN (see Methods). Each panel indicates the names of the cell types (color-matched to the rendered neurons) and the number of individual cells of each type (in gray); types without numbers are present once per brain hemisphere. Detailed morphology within the visual brain regions can be found in the Cell Type Catalog (Supplementary Fig. 1).

### Cell-type-specific genetic driver lines

Analysis and simulations of connectomes can suggest functions for many of its constituent cell types^55^. Testing those predictions requires genetic tools that permit the activity of specific cell types to be manipulated or marked for functional imaging or electrophysiology. For over 15 years, we have been making genetic driver lines targeted to specific cell types in the *Drosophila* visual system using the split-GAL4 intersectional method^56,57^. Using light microscopy to analyze the morphology of neurons expressed within first-generation driver lines, we developed several collections that successfully matched groups of known cell types in the lamina^21^, medulla^29^, and lobula^47^. These lines enabled targeted expression control within these cell types for functional experiments, further analysis of their anatomy^19^, and measurements of high-resolution transcriptomes^40^. Matching driver lines to well-defined cell types can be quite involved, but LM images of single neurons and populations, along with classic Golgi surveys^1,3^, provided a sufficient basis for matching most of the high-count OLCNs and OLIN types. An example of the success of our approach is that we find no new Dm types in the connectome compared to a prior LM study^19^, except for a new division of Dm3 cells and the dorsal rim types (Fig. 5c, Extended Data Fig. 10). However, the full visual system inventory (Supplementary Fig. 1) is the ideal reference for matching a much larger set of driver lines, especially the less numerous cell types. We report here a collection of 577 split-GAL4 lines (Fig. 8a; Supplementary Table 6) that have been matched to >300 EM-defined cell types across all our inventory groups.

**Fig. 8:**
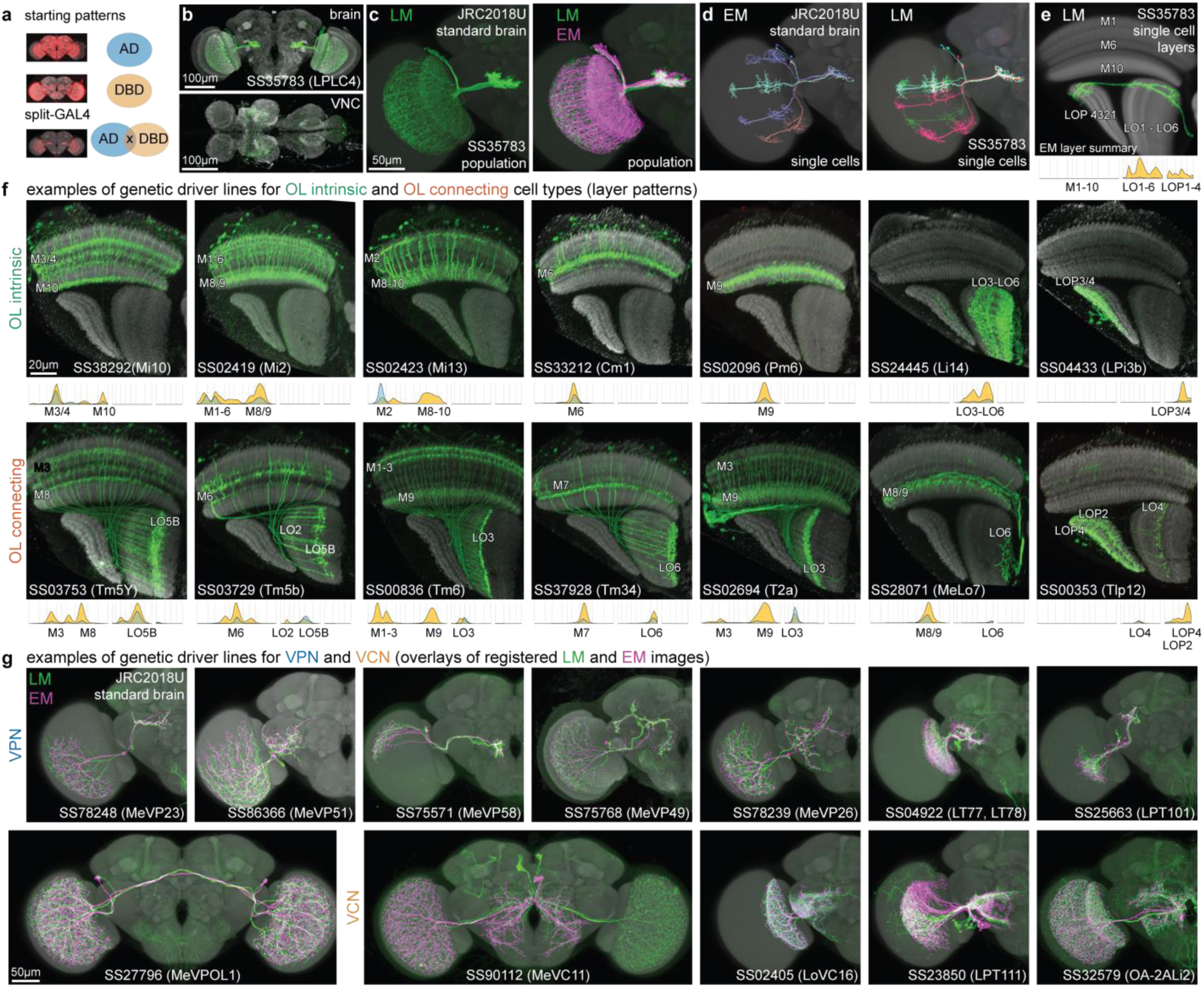
Genetic driver lines for targeting visual system neuron types, matched to the EM-defined cell types. (a) In a split-GAL4 driver line, two distinct expression patterns are genetically intersected to produce selective expression in cells of interest. We report 577 driver lines that we matched to >300 cell types. (b-e) Examples of the types of LM information used to characterize driver lines for matching to EM-defined cell types. (b) Overall expression pattern of example split-GAL4 driver line that primarily labels LPLC4 VPNs, with some “off-target” VNC expression. (c) LM image of LPLC4 neurons (driver line as in (b)) registered to and displayed on a standard brain (JRC2018U) (left) or overlayed with the skeletons of all EM-reconstructed LPLC4 neurons (registered to JRC2018U). (d) (left) EM reconstructions of individual LPLC4 cells. (right) LM images of stochastically labeled (MCFO^19^) individual LPLC4 neurons. (e) A manually segmented, registered LM image of a single MCFO-labeled LPLC4 neuron with a slice of the template brain. This image shows a different view than the images in (b)-(d), to emphasize the layer patterns in the visual system. A graphical summary of LPLC4’s innervation, as in Fig. 5b, is shown below. (f) Selected split-GAL4 lines driving expression in OLIN and OLCN types. Layer patterns of genetically labeled cell populations are shown with a neuropil marker (anti-Brp). Corresponding layers are indicated on both the LM images and the EM summary figures (see e). (g) Selected split-GAL4 labeled VPN and VCN types. Images show overlays with registered EM skeletons. Detailed layer-specific patterns can be found in the cell type catalog (Supplementary Fig. 1). Supplementary Table 6 lists split-GAL4 lines.

We found candidate matches for nearly all visual system cell types labeled by split-GAL4 lines, highlighting the completeness of our EM cell type inventory. This is all the more remarkable since most LM images are from female flies, whereas the EM reconstruction is of a male fly, suggesting that most optic lobe cell types are not sexually dimorphic. We matched 98% of the cell types in our inventory (accounting for 99.8% of cells) to cell types in the female FlyWire-FAFB dataset^15–17,25^ (Extended Data Fig. 13a,b; Supplementary Table 7), providing strong evidence for the completeness of our inventory and the absence of significant cell-type-level sexual dimorphism. The cell types that could not be matched between the datasets are candidates for sexually dimorphic neurons, with two examples shown in Extended Data Fig. 13c.

Figure 8b-e shows an example of morphological matching of a split-GAL4 driver’s expression pattern, which selectively labels a single population of visual neurons, to EM-reconstructed LPLC4s cells. We use brain registration to juxtapose EM-reconstructed neurons in the same reference as registered LM data, to compare single neuron morphology and population patterns (Fig. 8c-d). In most cases, the quantified innervation of the visual regions (Supplementary Fig. 1) provides a simple comparison that is sufficient for confident matching (Fig. 8e for LPLC4), especially for the OLIN and OLCN lines (Fig. 8f). Many VPN and VCN driver lines can be matched to EM-defined cell types using the central brain arborization patterns (visualized on a co-registered brain, Fig. 8g). We use additional information when the neurons labeled with a driver line appear to be credible matches for multiple EM-defined cell type. In these cases, we use finer differences in the layer innervations and features like the cell body distribution, regional pattern coverage, and arbor size and shape (Extended Data Fig. 12a-e). Our LM images often confirm the accuracy of the EM-reconstructed morphologies, even in a few cases where we find atypical neuron morphologies that appear to represent real but infrequent variants (Extended Data Fig. 12f). By linking genetic driver lines to EM-defined cell types, this collection establishes a powerful toolkit for probing the circuits of the visual system.

### Inter-region connectivity

Understanding the flow of visual information throughout the fly brain is a substantial undertaking, but the infrastructure we developed to catalog the visual system’s neurons provides an approach to an initial analysis. Examining major cell type groups, we asked whether specific connectivity patterns—from particular layers to others across the visual regions—are most prevalent. The first matrix in Fig. 9a shows the inter-region connectivity of the OLINs, and as expected, we find a block diagonal structure with prominent within-layer connections indicated by the high connectivity along the diagonal. The organization of the medulla’s connections supports the anatomical division into three units of more tightly interacting layers and the separation of interneurons into Dm (distal), Cm (central), and Pm (proximal). The summarized connectivity matrices (Fig. 9a) highlight the major pathways and indicate representative cell types substantially contributing to the strongest connection (Fig. 9b). The OLCNs have a much denser connectivity matrix (Fig. 9a), such that nearly every layer is connected to every other layer. However, there are more substantial connection patterns typified by prominent region-connecting cell types (highlighted in Fig. 9b). Notably, the connectivity above the diagonal is much higher than below, indicating that most connections flow in feedforward directions, e.g., from the medulla to the lobula and lobula plate. Finally, the analysis of the VPNs’ connectivity to the central brain reflects the broad classes of projection neurons (Fig. 6), with prominent connections from multiple visual regions often targeting the same central brain regions (Fig. 9b).

**Fig. 9:**
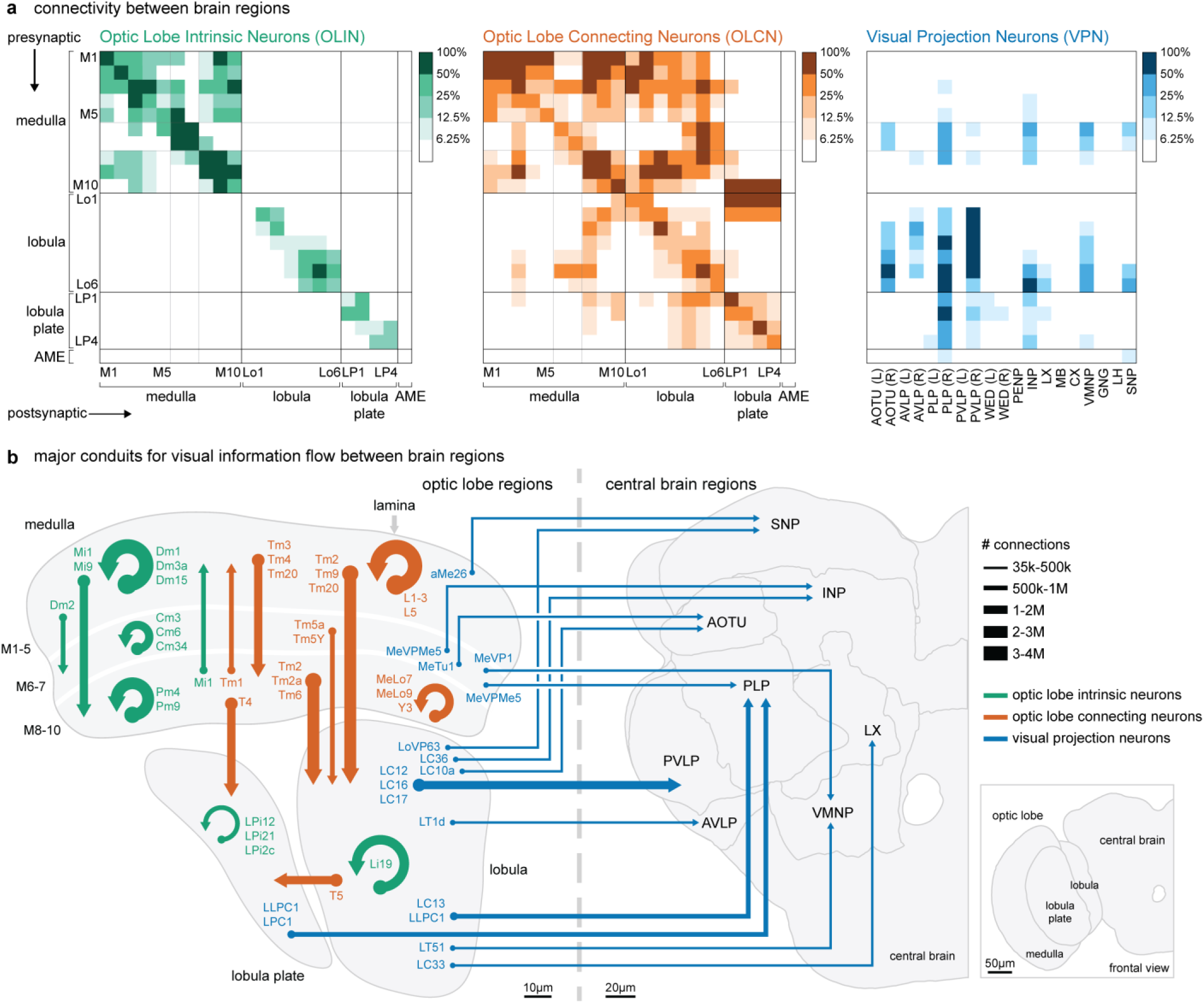
Connectivity within visual brain regions and with the central brain. (a) The connectivity between regions quantified as a weighted sum (see Methods) over all neurons from the OLIN (right), OLCN (middle), and VPN (left) groups. Optic lobe layers (Fig. 3) are treated as individual regions. The color code reflects the contributions to total connectivity: inter-region connections, the entries in each matrix, were ranked in ascending order and binned based on their contribution to the cumulative sum. (b) Schematic representation of the main conduits for visual information flow within and between brain regions based on the connectivity matrices in (a). The arrows are scaled to represent the number of connections between regions. Arrows are shown for each matrix entry in the highest 50% (darkest colors) with the addition of a few other prominent connections below this threshold. Major contributing cell types for every strong connection are indicated for each arrow. Layers of the visual regions and several central brain regions are grouped in this summary.

Interestingly, many central brain regions, such as the central complex, do not receive significant, direct visual projections. This is in part because connections outside of the top 93.75% of connections are not shown in Fig. 9. For example, there are direct VPN connections to specific Kenyon Cell types in the mushroom body that respond to visual information^58,59^. Additionally, we expect that visual signals arrive via central brain interneurons, as found for the mushroom bodies^58^, further expanding the footprint of vision in the central brain. By aggregating the visual neurons into this “projectome,” we emphasize the most prevalent connections to major brain regions. Notably, all connections are dissectible down to the individual cell types (Fig. 9b), which can be quickly found with a query of the neuPrint database. Together with the extensive collection of driver lines (Fig. 8, Supplementary Table 6), we now have a comprehensive toolkit for exploring this entire visual system and discovering how visual features are detected and where vision is integrated in the central brain.

### Concluding Remarks

Our survey has been exhaustively curated and proofread from the new connectome of one optic lobe of a male brain. In the coming months, proofreading of the rest of the central brain, and eventually the ventral nerve cord, of this connected volume will be completed. With these updates, our knowledge about the VPNs and VCNs will increase substantially as details of their central inputs and outputs are filled in. These datasets, together with an emerging connectome of a female brain^15–17^, provide a spectacular resource for understanding how vision is processed in central brain circuits and, ultimately, how vision is used to guide behavior. By mapping neural connections at unprecedented scale and resolution, connectomics overcomes the traditional trade-offs between completeness and accuracy in cataloging cell type diversity^13^. This approach has been particularly transformative in visual systems—the mammalian retina and especially in the fly—where it has facilitated detailed investigations of circuit function. In many cases, the analysis of these data directly suggests functional hypotheses. While there is some lingering skepticism about the interpretability of vast datasets or the translatability of structural maps to functional insights, these methods are already revolutionizing neuroscience on at least two levels: at their most powerful, the completeness of these surveys enables advanced analytical and modeling studies of integrative brain function, but more straightforwardly, these methods outline countless experimental roadmaps for detailed investigation of brain circuit function. The combination of connectomic surveys, detailed analysis, and genetic tools for experimental manipulation finally provide the necessary elements for bridging the gap between the stunning structural intricacy of neuronal circuits and the functional understanding the field has long sought.

## Acknowledgments

Janelia’s FlyEM Project Team (https://www.janelia.org/project-team/flyem) acquired and reconstructed the EM dataset in collaboration with the Connectomics group at Google. We thank the Janelia Research Campus for its dedicated support and FlyEM team members, in addition to the named authors who contributed, including Steering Committee members Gwyneth Card and Vivek Jayaraman. We thank Geoffrey Meissner, Janelia’s FlyLight Project Team and Project Technical Resources for help preparing and imaging fly lines; Mark Eddison, Yisheng He, and Gudrun Ihrke of Project Technical Resources for performing EASI-FISH and FISH experiments; Janelia’s Fly Core for fly care; Srini Turaga for support of N. Klapoetke; and Brett Mensch for feedback on the manuscript. We thank Cristian Goina, Hideo Otsuna, Konrad Rokicki, Robert Svirskas, and Janelia Scientific Computing Software for their work on confocal image processing, storage, and public release to Janelia websites, and Janelia’s Open Science Software Initiative for supporting the maintenance of ImgLib2 and BigDataViewer. We thank the supportive Janelia Fly community and the “FAFB optic lobe working group,” especially members of the Behnia, Chiappe, Silies, and Wernet groups for helpful discussion. This work used VVDviewer, based on software funded by the NIH: Fluorender: An Imaging Tool for Visualization and Analysis of Confocal Data as Applied to Zebrafish Research, R01-GM098151-01. This work was supported by the Howard Hughes Medical Institute and the Wellcome Trust (220343/Z/20/Z and 221300/Z/20/Z to GSXEJ with GMR, Gwyneth Card, Scott Waddell and Matthias Landgraf). This article is subject to HHMI’s Open Access to Publications policy. HHMI lab heads have previously granted a nonexclusive CC BY 4.0 license to the public and a sublicensable license to HHMI in their research articles. Pursuant to those licenses, the author-accepted manuscript of this article can be made freely available under a CC BY 4.0 license immediately upon publication.

## Author Contributions

MBR, AN, and GMR drafted the manuscript with input from SB, LEB, EG, JH, GBH, KJH, MJ, NCK, SK, FL, KDL, SP, SR, GMR, LKS, PaS, SS, and AZ. SB, JB, LEB, JC, MD, JF, EG, IH, JH, GBH, PMH, MJ, WTK, NCK, SK, FL, KDL, AN, DJO, SMP, SP, EMR, SS, PaS, ETT, LU, AZ, and TZ implemented software. KJH, ZL, and PKR prepared the sample. AN and GMR produced GAL4 driver lines. HFH, WQ, and CSX imaged the sample. CO, ToP, SP, SS, LKS, and ETT aligned the sample. SB, MJ, CO, and ShT produced the automated segmentation. GBH, CO, and PKR found the synapses. SB, JF, NCK, AN, MBR, and AZ determined neurotransmitters. JB, AN, and SS performed the LM-EM mapping. SB, JB, LEB, EG, JH, SK, FL, KDL, AN, CO, MBR, EMR, PaS, ShT, and AZ defined the neuropil compartments. SB, BSC, MC, SFM, MAF, GPH, CK, JK, SAL, ML, AL, CAM, CM, AN, NO, CO, TP, EMP, JRS, ALS, LAS, ShT, SaT, IT, AT, JJW, CW, EAY, and TY proofread the segmentation. SB, LEB, MD, EG, JH, NCK, SK, FL, KDL, AN, MBR, EMR, LKS, PaS, and AZ analyzed the connectome. SB, JB, LEB, MD, EG, JH, PMH, SK, FL, KDL, AN, MBR, EMR, SS, PaS, and AZ developed visualizations. SB, LEB, EG, JH, GSXEJ, NCK, CK, SK, FL, KDL, AN, CO, SMP, MBR, PKR, EMR, SS, PaS, ShT, ETT, LU, and AZ performed quality checks on the connectome. JH, AN, MBR, and ShT determined the cell types, and AMCF, PhS, AN, and GSXEJ matched cell types to FlyWire. SB, JF, RG, HFH, GSXEJ, CK, WK, FL, AN, SMP, SP, MBR, SR, GMR, and SS supervised internal efforts.

---

**Extended Data Figure 1.**
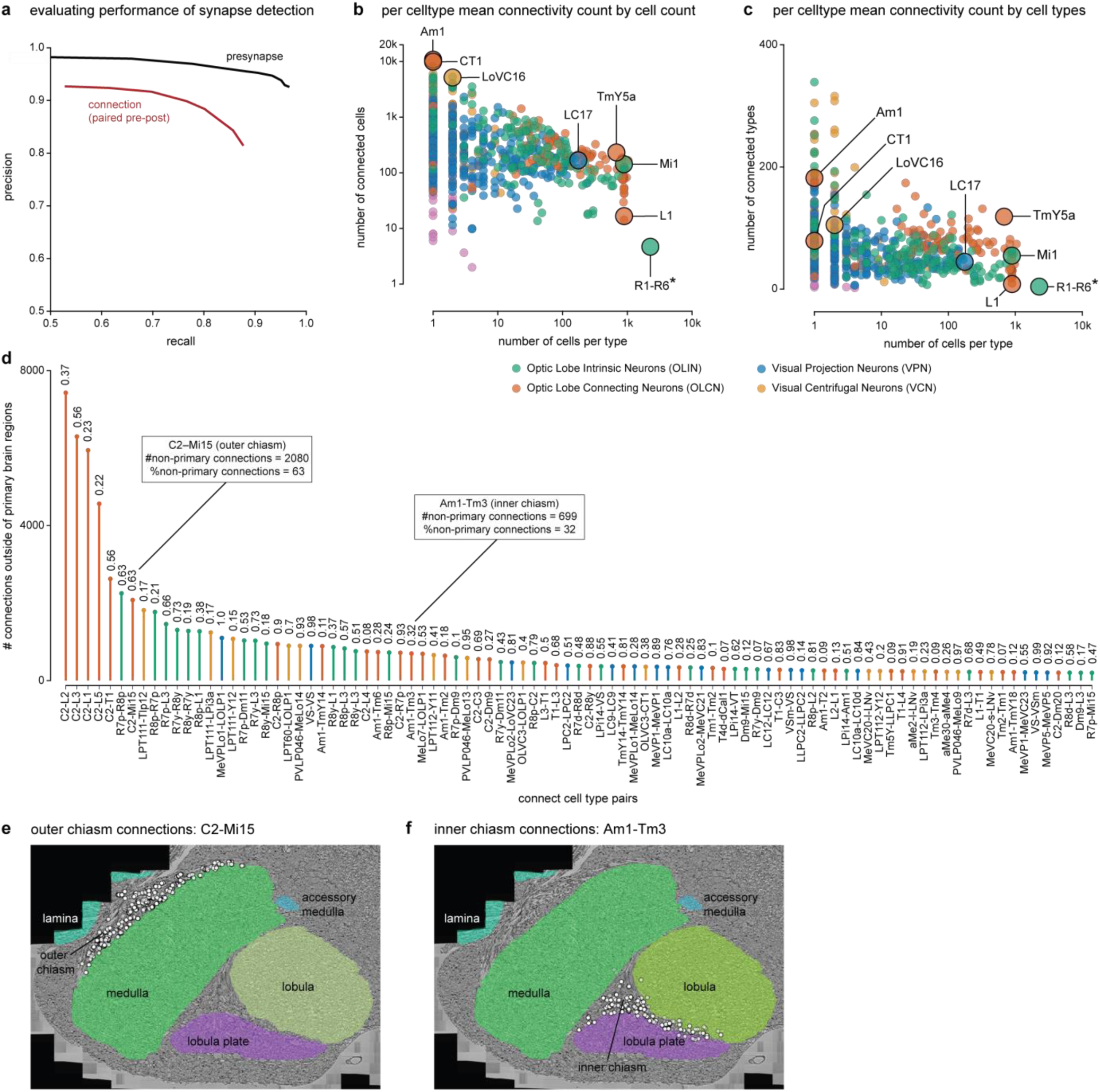
Connectivity Summary for the visual cell types (related to Fig. 1) (a) Performance of synapse detection. The precision/recall curve for presynapses (black), and connections (pre-post pairs, red). Precision is the probability that a detected feature was human-annotated, and recall is the percentage of features found by human-annotators that were also identified by the algorithm. (b) The mean number of connected cells (in the optic lobe, counting all connections > 1) as a function of the number of cells per type for all cell types in the inventory. Several examples are highlighted, as in Fig. 1g. (c) Mean number of connected cell types (in the optic lobe, counting all connections > 1) as a function of number of cells per type. The same cells are highlighted as in (b). (d) The cell type pairs with the largest contributions to connections outside of the main optic lobe regions. The plot shows the number of synaptic connections outside the primary visual regions and the fraction they contribute to all connections between the indicated cell type pairs. The connections are directional, and the pairs are ordered as A-B, where A is presynaptic to B. Many of these connections are in the outer or inner chiasm. Only pairs with >5% of their connections outside of the region boundaries are included. Additional details are in Supplementary Table 2. (e) Example of prominent connectivity in the outer chiasm: ∼60% of C2-Mi15 connections are within the outer chiasm, highlighted in (d). (f) ∼30% of Am1-Tm3 connections are within the inner chiasm, as highlighted in (d).

**Extended Data Figure 2.**
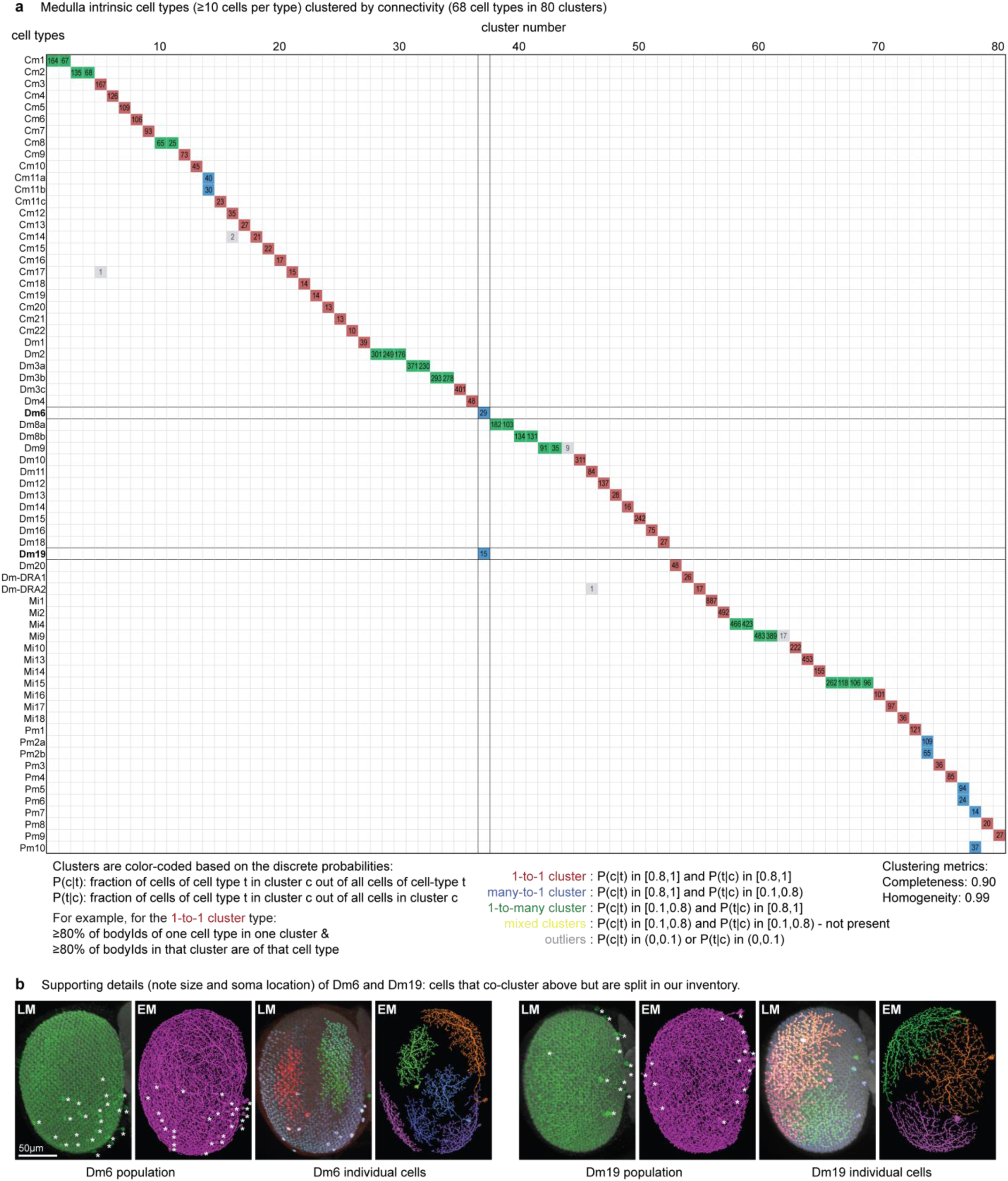
Sorting neurons with connectivity (related to Fig. 2) (a) Clustering based on connectivity for all medulla intrinsic cells with at least 10 cells per type. The 68 cell types are split into a preselected number of 80 clusters. Each cluster indicates the number of individual cells assigned to it. Cell types (Y-axis) are listed alphabetically, and clusters (X-axis) are further ordered by the number of cells per cluster. This sorting produces clusters with different cell type compositions (color-coded as indicated in the figure). We note that in all cases, 1-to-many clusters (green) for a given cell type are in a shared subtree of the dendrogram that does not contain other clusters. Compare to morphology-only clustering for the same set of neurons in Extended Data Fig. 7. (b) EM to LM comparison of cell types Dm6 and Dm19. The LM images used split-GAL4 driver lines combined with population or stochastic labeling, see methods. These cell types co-cluster in (a), but can be split by anatomy. For example, the cells have noticeably different sizes and distribution of cell bodies. Both features are visible in the EM and LM data (asterisks mark cell body locations). The two types can also be cleanly separated by further connectivity clustering (not shown), and the existence of a selective split-GAL4 driver for each indicates that they are genetically distinct.

**Extended Data Figure 3.**
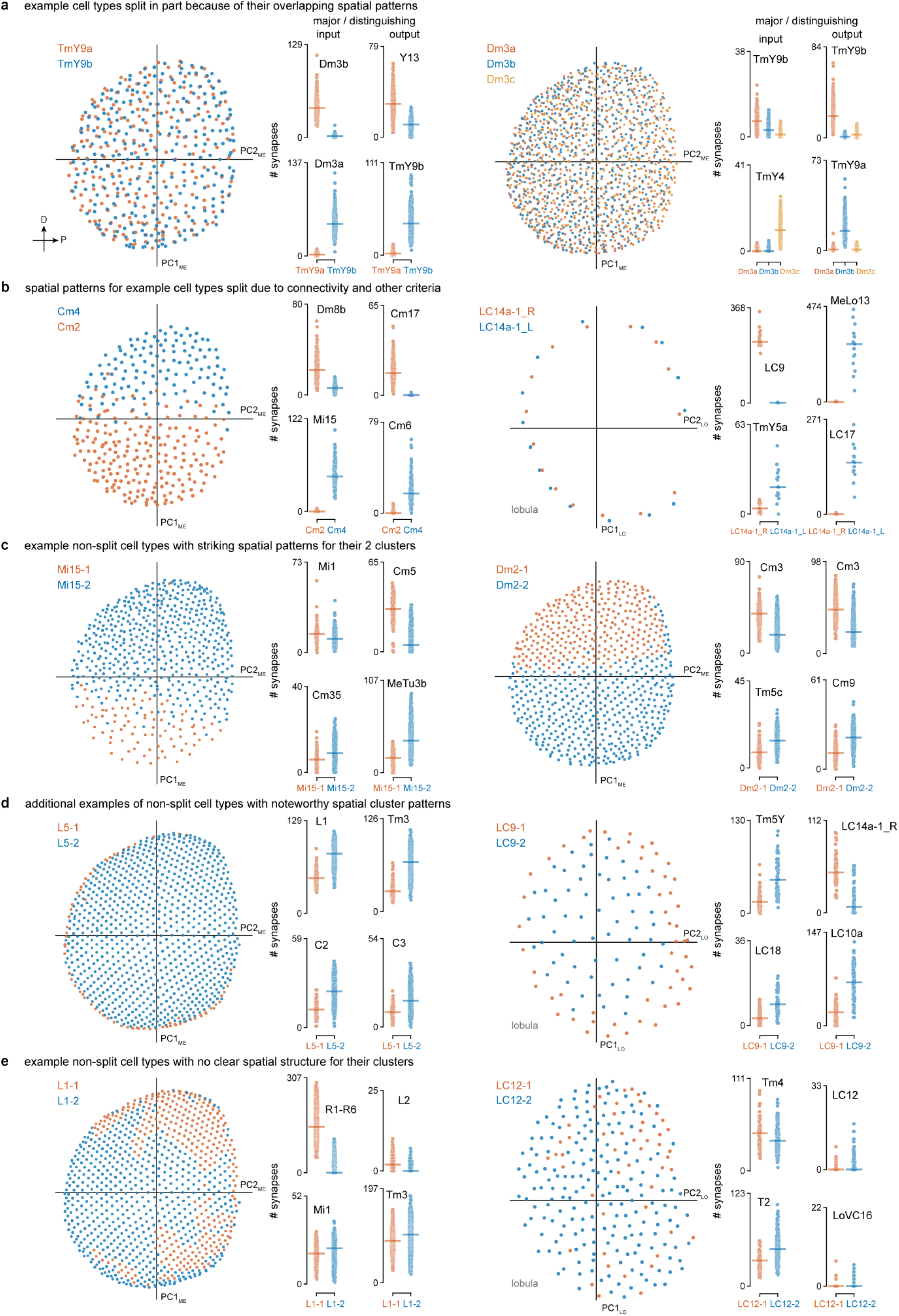
Combining connectivity clustering with spatial distributions to evaluate cell type merges/splits (related to Fig. 2) 10 groups of cell types evaluated for potential splitting into separate cell types; presentation follows the conventions of Fig. 2c,g,h, where selected distinguishing input and output connections are shown. On the right side of each panel, each point represents the sum of connections to/from the indicated cell type for a single neuron from each group. This set of examples covers the various cases we find across the dataset. Cells that are split into distinct types are indicated by their assigned names, but examples of non-split cells are indicated as −1 and −2 for the clusters of cells with the same type designation. (a) Examples of groups of similar cell types that show connectivity differences and overlapping spatial distributions (mosaics) and were split into different types. Splitting TmY9a/TmY9b and Dm3a/b/c is also supported by their different arbor orientations (Fig. 5). (b) The Cm2 and Cm4 cell types show overlapping distributions, consistent connectivity differences, and different neurotransmitter predictions. Left and right instances of cells with arbors in both optic lobes (here LC14a-1) often have distinct connectivity within the same optic lobe. Such neurons were treated as two types (e.g., LC14a-1_L and LC14a-1_R) in some analyses. (c) Examples of cell types with subclusters with distinct spatial distributions. Mi15 and Dm2 subclusters occupy different domains along the DV axis; note that the overall density of Mi15 cells also differs along this axis. Such divisions suggest regional differences within these cell populations, but the absence of strong, consistent connectivity differences prevented division into distinct types. (d) Other examples of cell types that were not split owing to the lack of strong connectivity differences between the subclusters that show striking spatial distribution patterns: L5 (cells at margin separate) and LC9 (center vs. perimeter). (e) Additional examples of cell types that were not split and show no obvious structure in the spatial distributions of subclusters: L1 and LC12.

**Extended Data Figure 4.**
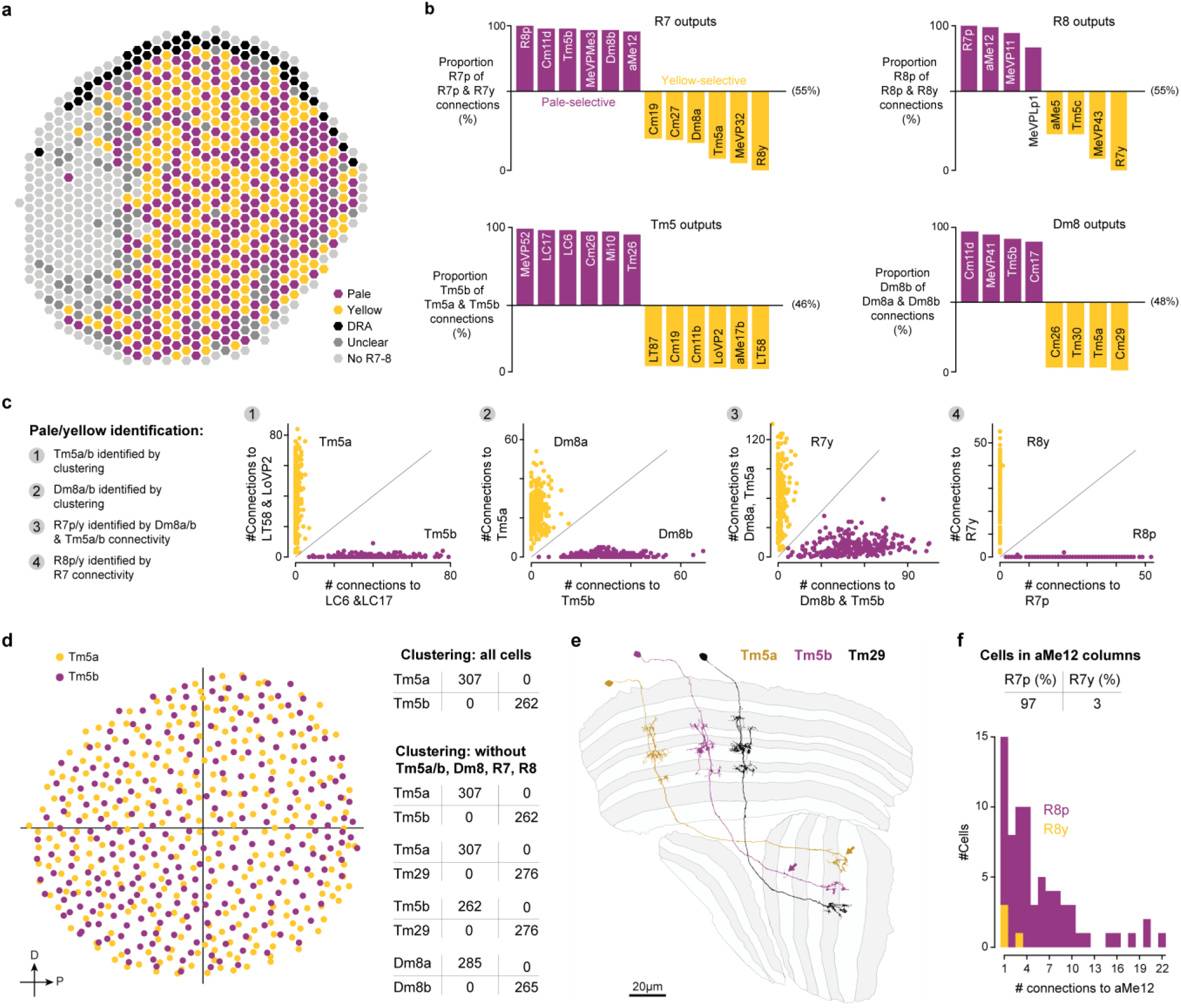
Functionally specialized inner photoreceptor types supply the visual system with color and polarized light information (related to Fig. 2) (a) Summary eye map of pale and yellow columns that supply color information, and dorsal rim area (DRA) columns that supply polarized light information. A subset of the photoreceptors in one part of the medulla was very heavily stained, resulting in highly fragmented terminals that could not be combined into identifiable neurons. Eye map uses the coordinate system based on medulla columns introduced in Fig. 3a. (b) Plots of pale and yellow pathway connections. For each synaptic connection pair, bars indicate the fraction of pale-pathway connections, e.g. (# R7p to Tm5b connections) / (# R7 to Tm5b connections). The baseline is the fraction of connections expected from the fraction of pale-pathway cells of that type, e.g. # R7p cells / # R7 cells. (c) Overview of the pale/yellow identification process. Step 1: Clustering connectivity with lobula cell types unambiguously categorizes Tm5 cells as Tm5a and Tm5b; summed connectivity with a subset (LC6 & LC17, LT58 & LoVP2) is shown for individual Tm5 cells. Step 2: Clustering of connectivity with Tm5a, Tm5b and other cell types unambiguously categorizes Dm8 cells as Dm8a and Dm8b; connectivity with Tm5a and Tm5b shown. Step3: R7 cells are classified as pale (R7p) or yellow (R7y) using their connectivity with Dm8a/b and Tm5a/b. Step 4: R8 cells are unambiguously classified as pale (R8p) or yellow (R8y) using their connectivity with R7p/y. (d) Left: Spatial clustering for Tm5a/b, following conventions of Fig. 2 and Extended Data Fig. 3. Right: Results of classifying Tm5a/b cells using connectivity with all cell types (top), and Tm5a/b, Dm8a/b, and Tm5a/b and morphologically similar Tm29 when Tm5a/b, Dm8, R7, and R8 are excluded (below). (e) Examples of Tm5a, Tm5b and Tm29 morphology. Many Tm5a cells have hook-shaped terminals in the lobula (yellow arrow) and slightly narrower medulla processes than Tm5b or Tm29 cell types; many Tm5b cells have a small process in the lobula layer 2 (purple arrow) missing in TM5a and Tm29. (f) Pale/yellow classification was corroborated using colocalization (top) and connectivity (bottom) of inner photoreceptors with aMe12, a cell type previously identified as pale-specific^24^. Top: Fraction of R7p and R7y in 97 manually identified columns innervated by aMe12. Bottom: Distribution of the number of identified synaptic connections with aMe12 of R8p/y cells.

**Extended Data Figure 5.**
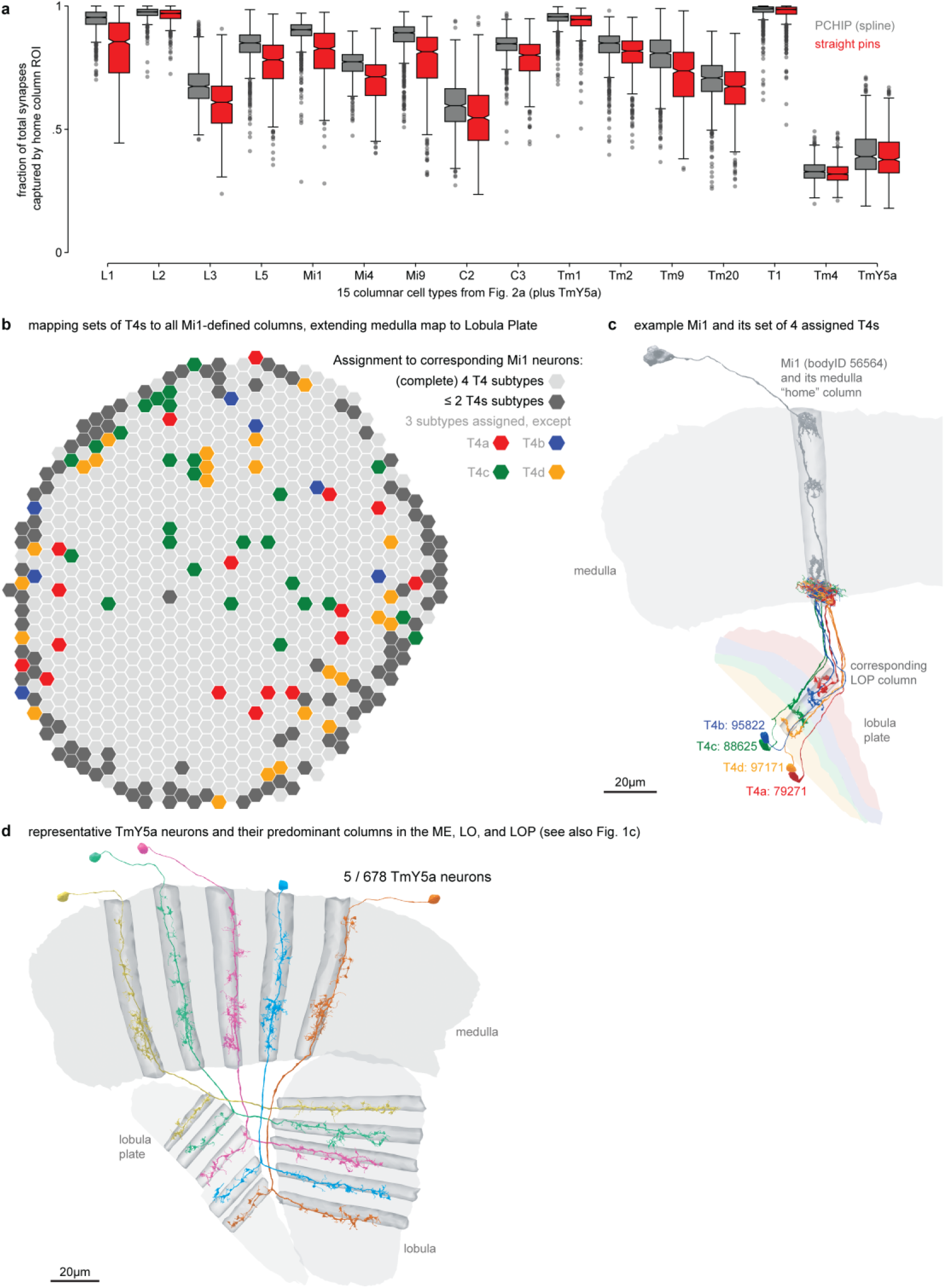
Supporting data for the creation of columns and their extension to the lobula plate (related to Fig. 3) (a) Comparison of straight lines to spline-based method for defining medulla column centerlines. Plots shows the fraction of synapses in the home column (the column with the largest synapse count) of each neuron of a cell type. For each cell type used to construct the medulla columns (first 14), the fraction is higher for splines than for straight lines. For the other 2 cell types, the fractions are comparable. Boxes show first, second, and third quartiles; whiskers are drawn at 1.5 times the interquartile range below and above the first and third quartiles, respectively. (b) A lobula plate coordinate system was derived from the medulla coordinates by mapping neurons of the four T4 types to individual Mi1s. In most hexagons (712, in light grey), all 4 types could be matched to a single Mi1, but in 75, only 3 types could be matched (the missing neuron is indicated in color). In the 105 dark hexagons near the perimeter, only 2 or fewer T4s could be matched to an Mi1. Matching criteria include connectivity and distance (see Methods). (c) An example Mi1 and the set of 4 assigned T4 neurons. (d) TmY5a neurons are the most numerous TmY cells and are used to visualize the column assignments across neuropils. This example shows a few TmY5a cells with the corresponding home columns in the medulla, along with the lobula and lobula plate columns with the same hexagonal coordinate.

**Extended Data Figure 6.**
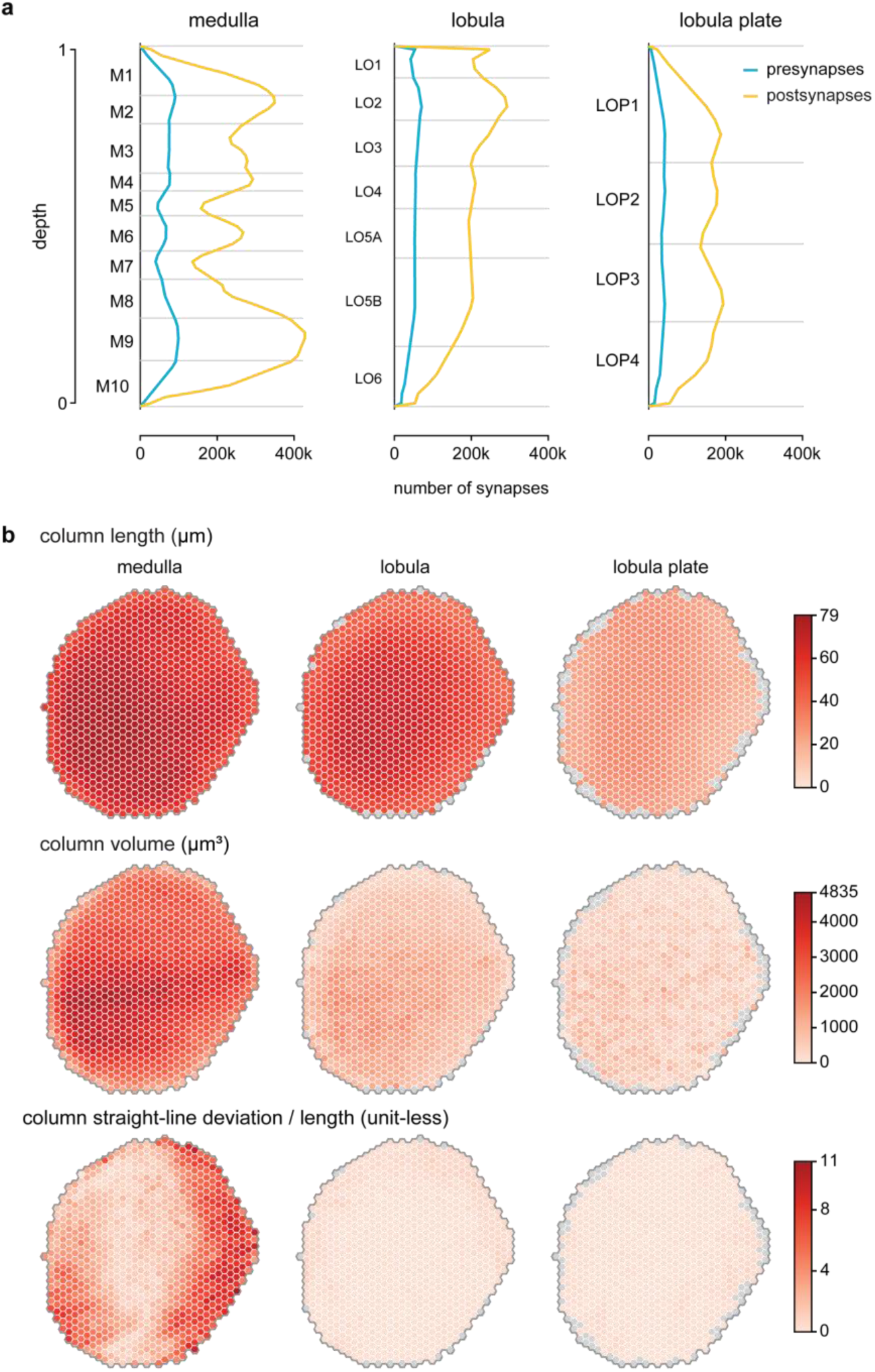
Summary of synapse distributions by columns and layers (related to Fig. 3) (a) The spatial distribution of presynapses (blue) and postsynapses (yellow) as a function of depth in the 3 main visual brain regions. The ratio of pre-to-post is approximately 1:6 and this is conserved across most of the layers of the visual system. (b) Quantification of the dimensions of the columns in the main visual brain regions. As expected, columns in the medulla are longest, the lobula columns somewhat shorter, and the lobula plate columns much shorter. Column volume is shown in the middle row. There is a noteworthy increase in volume near the equator of the eye and towards the front (left) that is especially prominent in the medulla. The bottom row shows the straightness of columns normalized by column length. Medulla columns, especially near the anterior and posterior margin are found to have the greatest curvature. While this reflects the nature of that neuropil, it is also in part due to the richer set of neurons used to define medulla columns. Gray columns around the periphery indicate columns that are not present in that brain region.

**Extended Data Figure 7.**
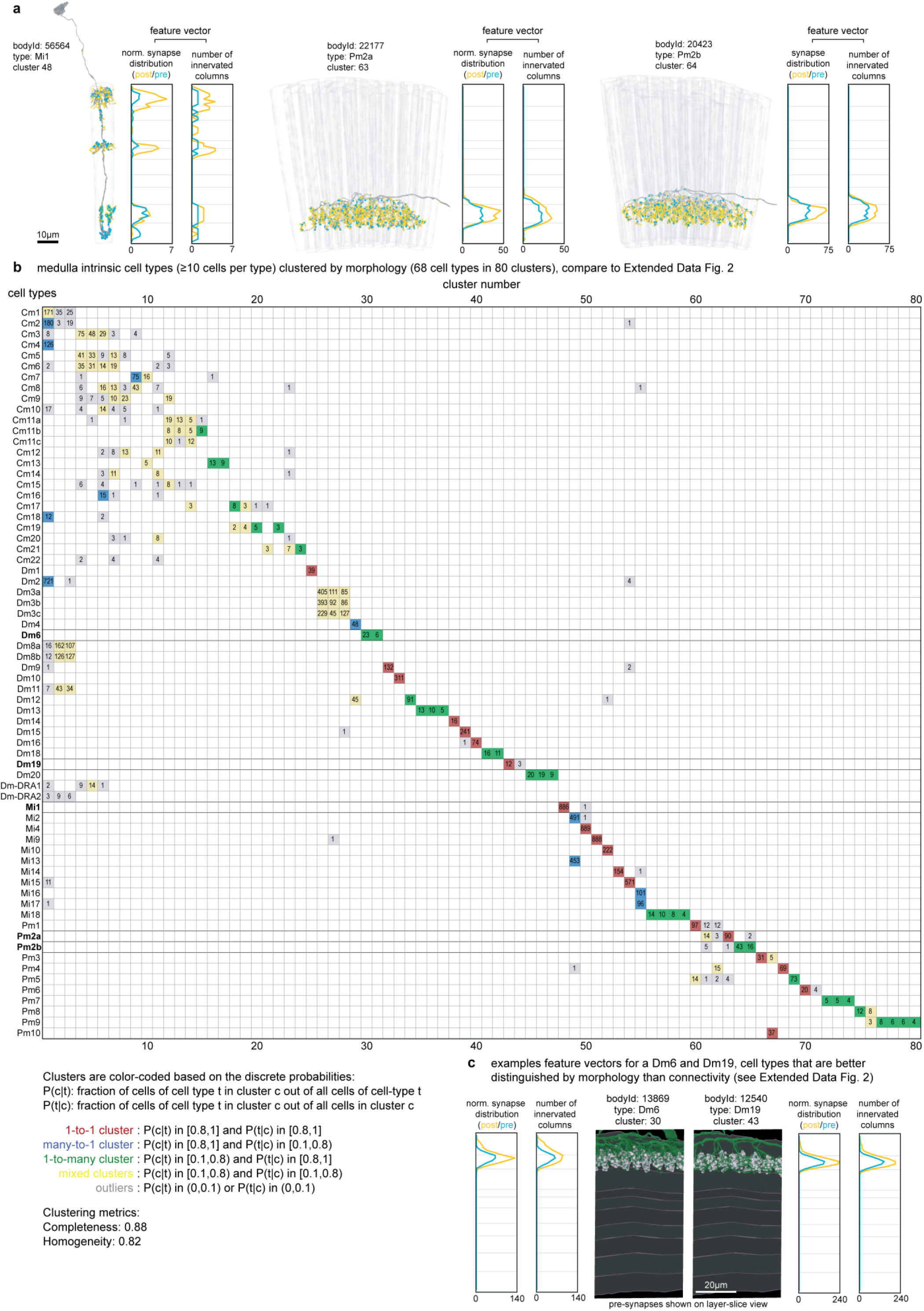
Clustering neurons using quantified anatomy (related to Fig. 5) (a) Examples of 3 neurons, each of a different type, to illustrate the aspects of quantified morphology used to construct a feature vector for each cell. Each neuron is shown together with its primary columns and the layer distribution of its presynapses and postsynapses. The feature vector comprises scaled synapse distributions and the number of innervated columns per depth, separately for the presynapses and postsynapses (see Methods). The pre/post synapse distribution is normalized such that the sum equals the sum of pre/post number of innervated columns for that cell. It is noteworthy that Pm2a/b neurons can be more easily distinguished by the total number of innervated columns than by their volume (see Fig. 2f). (b) Confusion matrix of clustering the same 68 medulla intrinsic cell types as in Extended Data Fig. 2 where clustering was based on connectivity, but here based on the quantified morphological feature vectors. The data are sorted and color-coded as in Extended Data Fig. 2 to facilitate comparison. (c) As an example of morphologically similar neurons from different cell types, we highlight the feature vectors for one Dm6 and one Dm19, representing these cell types that are more separable by their quantified morphology (see (b)) than by their connectivity (Extended Data Fig. 2). The feature vectors show a noticeable difference in the innervation of layer M1, which we confirmed by visualizing the presynapses of both cells in a layer-slice view.

**Extended Data Figure 8.**
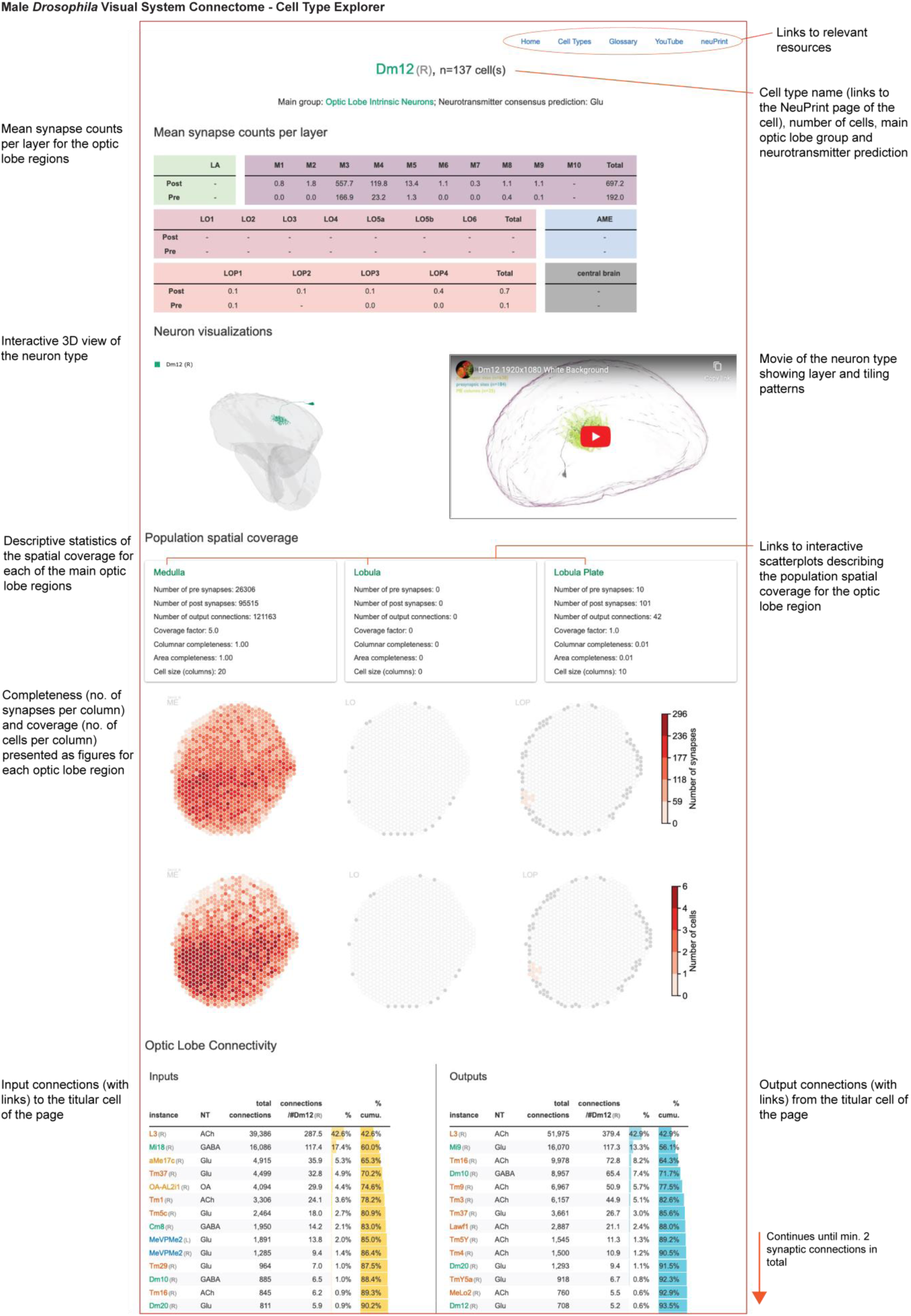
Interactive visual system Cell Type Explorer web resource (related to Fig. 5) The *Drosophila* Visual System Cell Type Explorer is a set of interactive webpages designed to offer more details about the retinotopy and connectivity for each cell type in the optic lobe. Each page features the name of the cell type at the top, which links to the cell type’s neuPrint page, and includes information on its assigned neuron group and predicted neurotransmitter. Below this header are tables displaying mean presynapse and postsynapse counts across every optic lobe region, with the main regions divided into layers. These tables include mean (by cell) synapse counts within the central brain as well. The next section presents a 3D interactive view of a representative cell alongside a linked video that illustrates the cell type’s layer and tiling patterns, as well as the entire cell population. The “Spatial coverage” section further describes the distribution of synapses and cell counts per column across the main optic lobe regions (introduced in Fig. 5d), including data on cell size and total synapse counts. The bottom of the page is devoted to the optic lobe connectivity table, arranged by the magnitude of total synaptic connections. This table color-codes each cell type according to its main group, listing on the left ("Inputs") those cell types that provide input to the profiled cell type, and on the right ("Outputs"), those cell types that receive input from it. The webpages are interconnected, allowing users to navigate between cell types via the Optic Lobe Connectivity table. Navigation is further facilitated by links at the page’s top to the Home page, where users can search for cell types by name, an Index page listing all optic lobe cell types, and a Glossary page that clarifies the terminology used throughout the site. We also provide interactive versions of the scatter plots in Fig. 5e,f, and Extended Data Fig. 9, so users can discover cell types by their coverage properties.

**Extended Data Figure 9.**
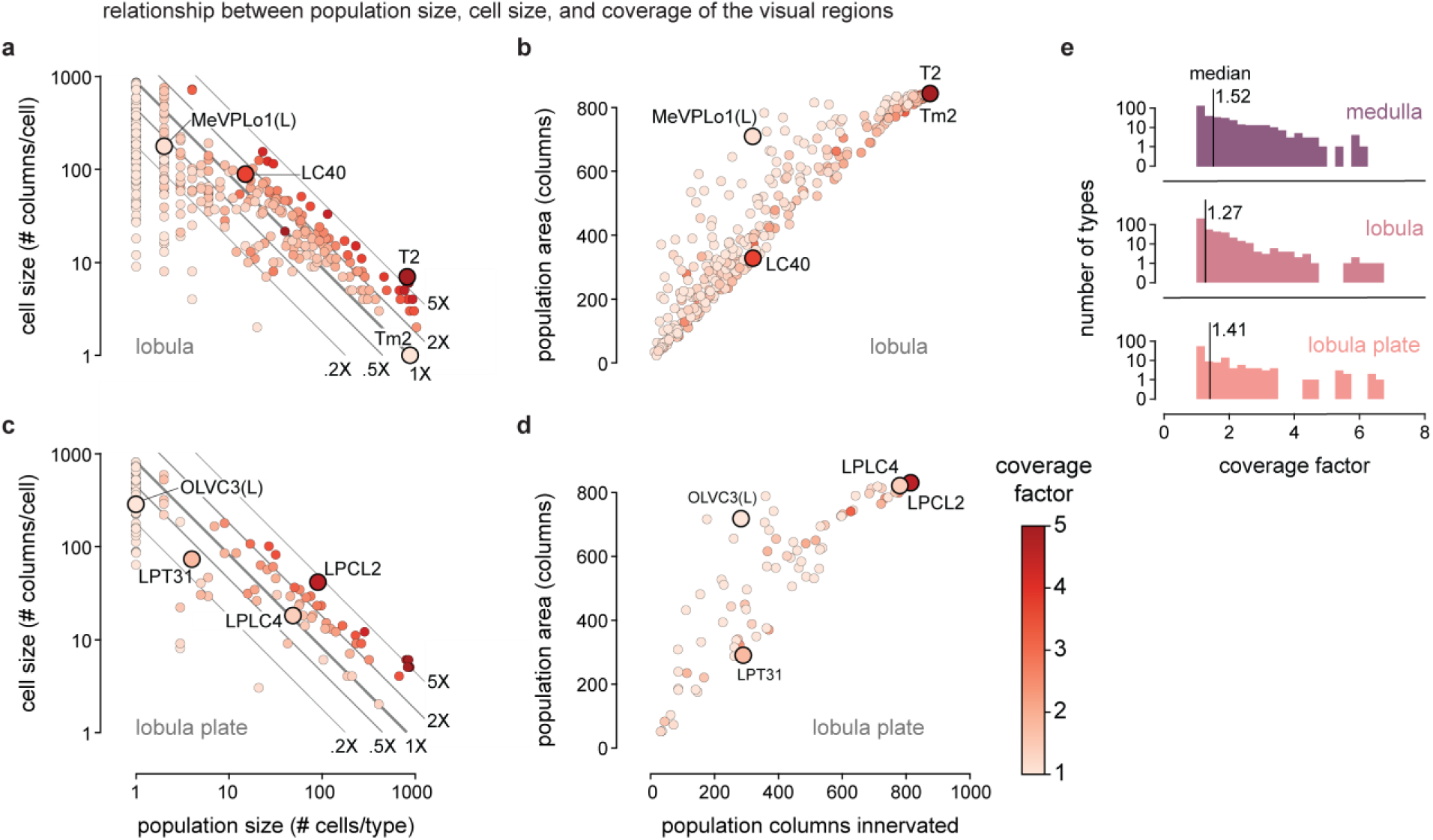
Coverage and density analysis for the cell types in the lobula and lobula plate (related to Fig. 5) (a) The relationship between population size (total number of cells) and cell size (the average number of columns innervated by a single cell) of cell types within the lobula, color-coded by coverage factor (the average number of cells that innervate a single column). The diagonal lines are guides to the ratio of global coverage: 1X for a given cell size, this is the “optimal” population size to tile the columns of the region, 2x and 5x more neurons and 0.5x and 0.2x fewer neurons than would be needed to fully cover the region at this cell size. Selected types to highlight how the coverage factor reflects the tiling properties of the neurons are T2 (6.72), Tm2 (1.03). (b) Summary plots of the density with which cell types innervate the lobula, summarized by plotting the number of columns innervated against the convex area (in units of columns) covered by the total population. Only types with at least 5% of their total synapses and at least 50 synapses in total within the chosen brain region were included in the plots. Selected types to highlight how the ratio between the columns innervated and area covered captures the density of innervation: MeVPLo1 (left instance, columns: 319, area: 710), LC40 (columns: 320, area: 329). (c) Same as (a) but for types assigned to the lobula plate. Selected types to highlight how the coverage factor reflects the tiling properties of the neurons are LPLC2 (4.63), LPLC4 (1.41). (d) As for (b) but for types assigned to the lobula plate. Selected types to highlight how the ratio between the columns innervated and area covered captures the density of innervation OLVC3 (left instance, columns: 283, area: 718), LPT31 (columns: 290, area: 290). (e) Distribution of coverage factor values in the three main optic lobe regions (top to bottom: medulla, lobula, lobula plate). Vertical black lines indicate the median coverage factor per region.

**Extended Data Figure 10.**
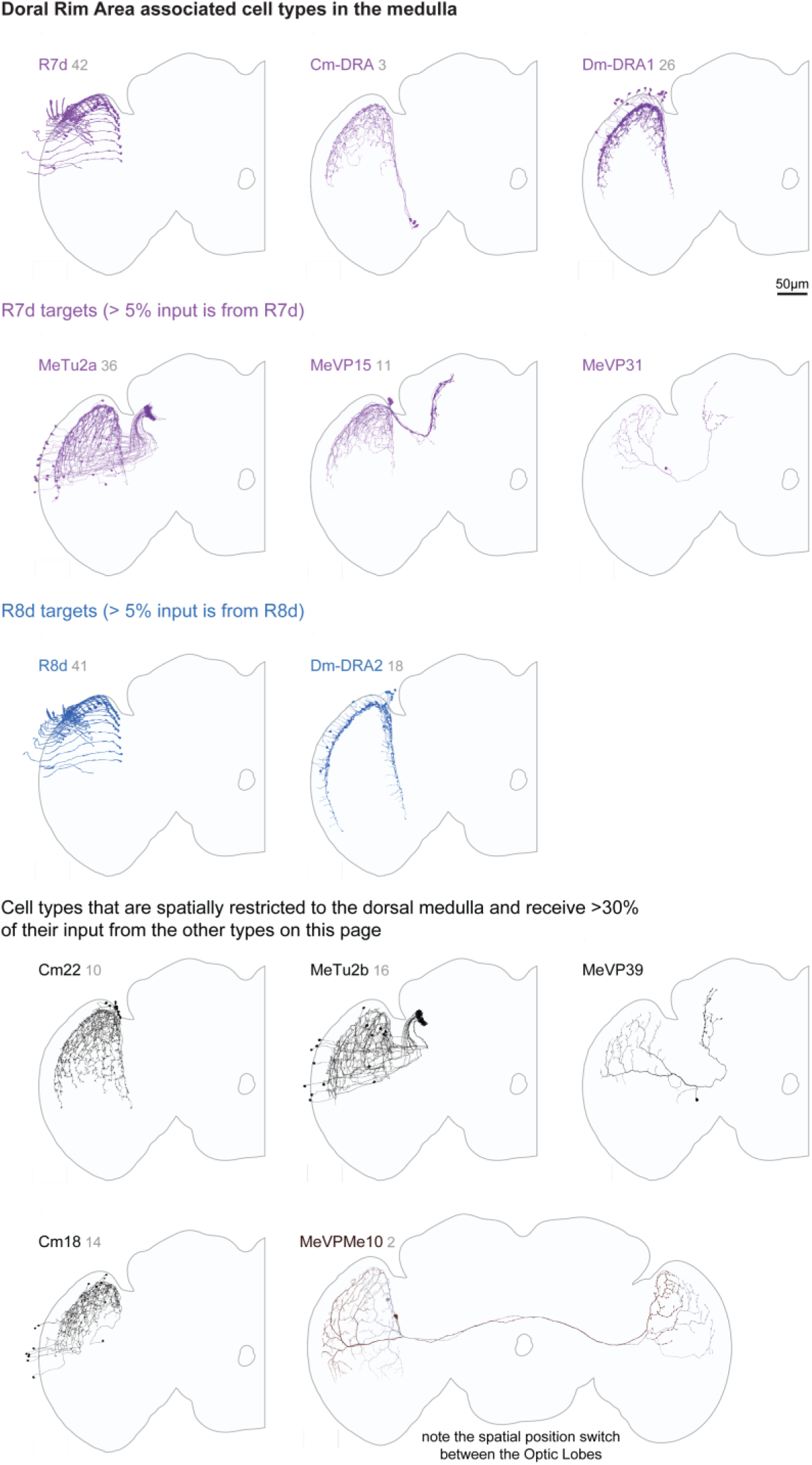
Neurons associated with the Dorsal Rim Area (related to Fig. 6) The Dorsal Rim Area (DRA) of the eye is a specialized zone of photoreceptors engaged in detecting polarized light. In the dorsal medulla, there are specialized cell types not found elsewhere in the medulla that process the output of the DRA photoreceptors. The primary cell types of the medulla area that correspond to the DRA of the eye are shown and organized into groups. The first two groups show R7d (magenta) and R8d (blue) photoreceptors and their main targets, which for R7d include both VPNs and medulla intrinsic cells. The lower panels show additional cell types that are identified as other components of the medulla DRA network by their regional arborizations in the dorsal medulla and their connections with cells in the top panels and with each other. One of these cell types is a VPN that connects the DRA regions of the two optic lobes. Each panel shows all members of each cell type (for the right OL). The R7d and R8d photoreceptors are undercounted (see Extended Data Fig. 3 and Methods).

**Extended Data Figure 11.**
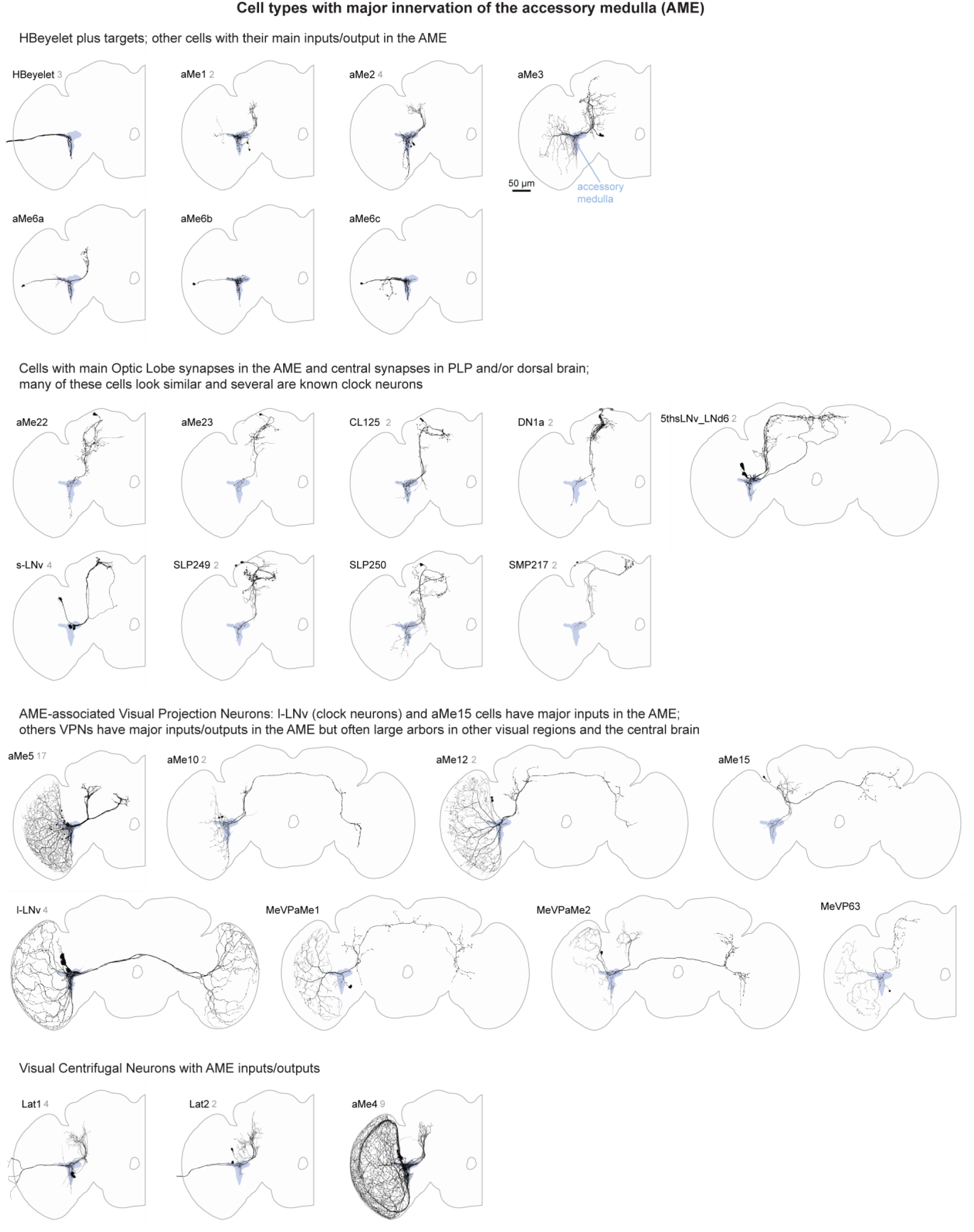
Selected neurons associated with the Accessory Medulla (related to Fig. 6) The Accessory Medulla (AME) is a small brain region located at the anterior-medial edge of the medulla that is mainly known for its role in circadian regulation. It contains processes of several clock neuron types and a diverse group of VPN and VCN cells. This page does not include all AME-associated neurons but shows examples of cell types and cell type groups with processes in the aMe. For each cell type shown, all identified individual cells (for the right OL) are included.

**Extended Data Figure 12.**
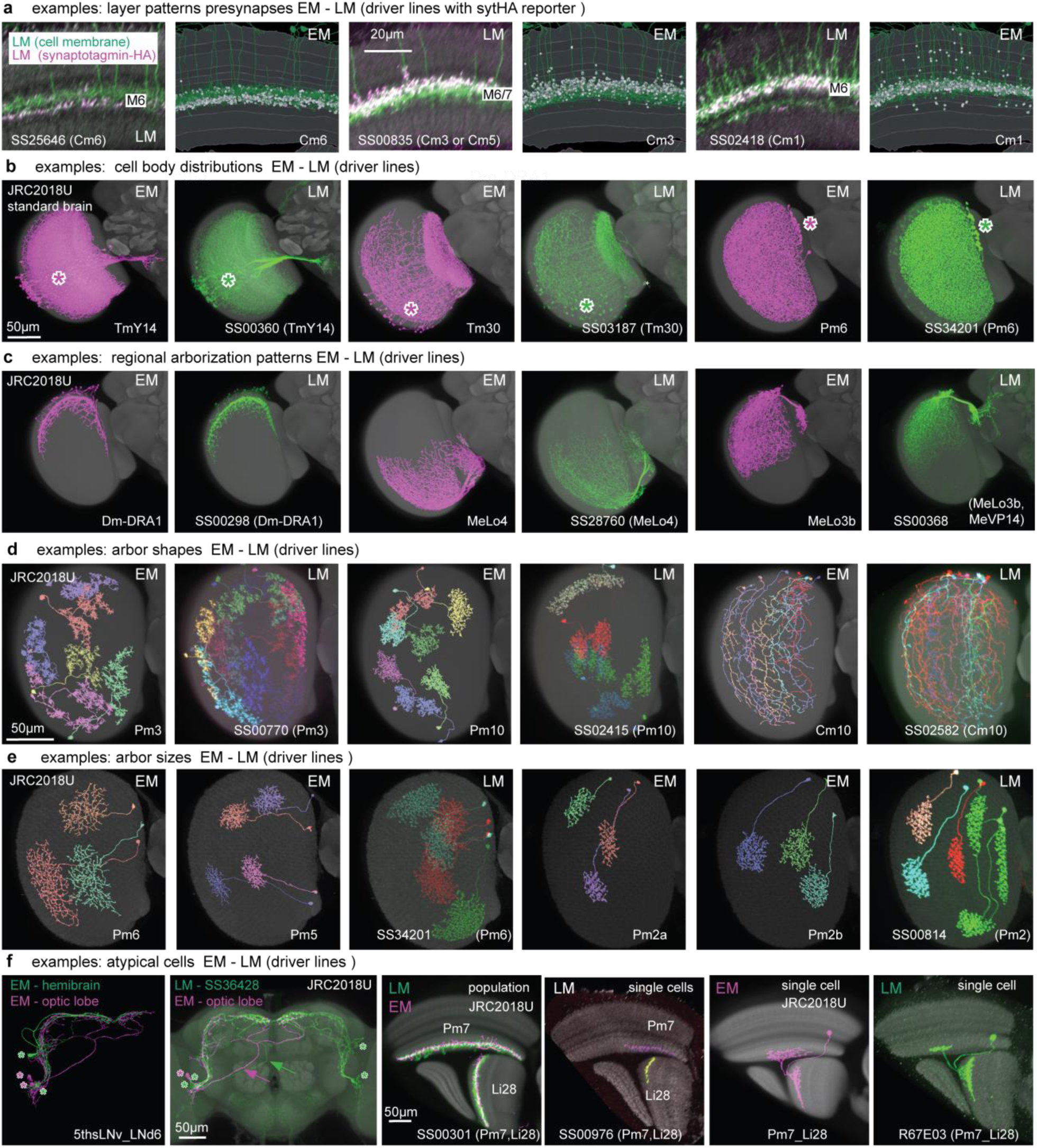
Supporting examples of anatomical features used for EM-LM matching (related to Fig. 8) Images in (b-e) show registered LM or EM images displayed on the JRC2018U template brain. LM images in (a-c) are based on full expression patterns; those in (d,e) show stochastic labeling. (a) Distribution of presynaptic sites in different medulla layers for three Cm cell types. LM images show projections through reoriented substacks (selected to show medulla layers) of the expression of a synaptic marker (syt-HA; magenta) and a membrane marker (green) driven by the indicated split-GAL4 lines. EM-based images show presynaptic sites (magenta) and EM meshes (green) of cells of the indicated types in a slice of the medulla selected to show layer patterns. (b) Cell body locations. While the soma location of individual cells is variable, general areas with cell bodies (indicated by asterisks) are similar within a cell type. Examples: TmY14 (cell bodies in a wedge-shaped subregion of the medulla cell body rind (MECBR), Tm30 (cell bodies in the ventral MECBR), and Pm6 (cell bodies in a cluster at the dorsal-medial edge of the medulla). (c) Regional arborization patterns. Examples: Dm-DRA1 (dorsal rim), MeLo4 (ventral medulla and lobula) and MeLo3b (dorsal medulla and subregion of dorsal lobula). The SS00368 driver also labels MeVP14 cells which overlap with MeLo3b in the medulla but extend into the central brain. (d) Arbor shape. Pm10 terminals have a more compact shape than those of Pm3 cells. Cm10 cells spread across nearly the full length of the medulla along the DV axis but are much narrower along the AP axis. (e) Arbor size indicates that SS34201 is expressed in Pm6 cells. SS00814 cells appears to be a better match to Pm2a than to Pm2b cells but the driver might also express in a combination of these cell types. (f) Examples of atypical cells observed in the EM reconstructions and LM examples with similar morphology, indicating the EM morphology is unlikely to be a reconstruction error. (Left two panels) Two clock neurons (the 5th s-LNv and LNd6) that typically have different cell body locations (asterisks) and similar projection patterns, show similar cell body locations and, in one case, an unusual axonal path in the optic lobe dataset (cells in magenta, asterisks mark cell body locations). For comparison, hemibrain reconstructions of the same cell types are shown in green. Most available LM images show the typical morphology, but we found one brain in which the expression pattern of a split-GAL4 line in one hemisphere matches the cell shape and cell body distribution seen in the optic lobe dataset. (Right four panels) An unusual cell (annotated as Pm7_Li28 in the EM dataset) has LM counterparts. From left to right: Optic lobe layer pattern of a driver line labeling both Pm7 and Li28 overlaid with registered EM reconstructions of these cells. Stochastic LM labeling of a Pm7 and an Li28 cell. The combined Pm7_Li28 cell in the EM (displayed on the standard brain). An LM example of a similar cell is displayed in a similar view.

**Extended Data Figure 13.**
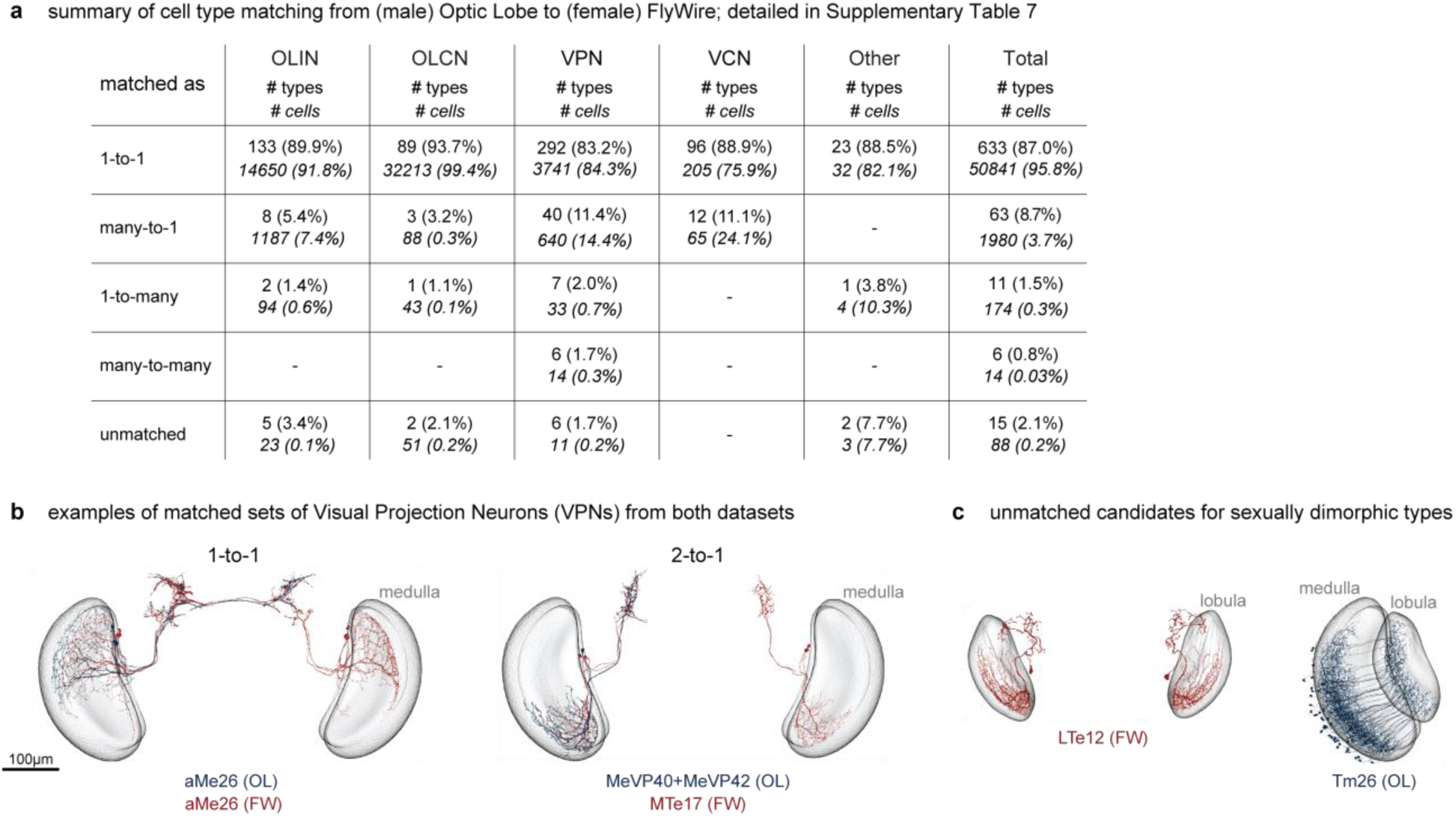
Matching cell types between the male optic lobe and FlyWire datasets (related to Supplementary Table 7) (a) Summary table of cell type matching from the (male) Optic Lobe (OL) to the (female) FlyWire (FW) dataset^15–17,25^. The full set of matches across cell types is detailed in Supplementary Table 7. This table shows the number of matched cell types and cells for each dataset, categorized by optic lobe cell type groups (Fig. 1c) and the level of matching: 1-to- 1, many-to-1, 1-to-many, many-to-many, and unmatched. The counts here are referenced to the male optic lobe dataset. (b) Examples of matched sets of neurons from both datasets. The left panel shows an example of a "1-to-1" match, with the neuron type aMe26 from both datasets. For most cell types, the FlyWire annotations of Schlegel et al.^16^ identify neurons on both sides of the brain. The right panel shows an example of a "2-to-1" match, with the cell types MeVP40 and MeVP42 from the OL dataset matched to the type MTe17 in FW. (c) Examples of candidate dimorphic neurons, each identified in one dataset but not the other. The left panel shows the neuron type LTe12 from the FW dataset, while the right panel shows the cell type Tm26 from the OL dataset (only the right optic lobe is shown). Tm26 appears to be the cell type previously reported as male-specific^60^, and an additional unmatched OL cell type (LoVP92, not shown, see Supplementary Table 7) resembles a different previously described male-specific cell type^61^. Images in b and c are shown at the same scale and from the same perspective.

**Extended Data Table 1:** The counts of presynapses and the completeness of connections for the optic lobe brain regions and the layers of the medulla, lobula, and lobula plate. “Pre” completeness is the percentage of presynapses contained in “traced” neurons. “post” completeness is the percentage of postsynapses assigned to traced neurons. Connection completeness refers to the percentage of synapses for which both the presynape and the postsynapse are in traced neurons. Neurons designated as "status = Traced" in neuPrint have passed multiple rounds of quality checks, are believed to contain no false merges, and have all of their main branches within the volume reconstructed (whenever possible, corroborated by comparison to other neurons of the same type). The lamina posed substantial reconstruction challenges (see Methods).

| Neuropil | presynapses<br>($\times 10^3$) | completeness (percentage) | | |
| --- | --- | --- | --- | --- |
|  |  | pre | post | connection |
| medulla (total) | 4144 | 96 | 56 | 55 |
| layer M1 | 488 | 93 | 60 | 57 |
| layer M2 | 407 | 97 | 64 | 62 |
| layer M3 | 643 | 94 | 50 | 47 |
| layer M4 | 254 | 97 | 54 | 53 |
| layer M5 | 164 | 92 | 52 | 48 |
| layer M6 | 447 | 96 | 52 | 50 |
| layer M7 | 213 | 98 | 47 | 46 |
| layer M8 | 434 | 98 | 56 | 55 |
| layer M9 | 766 | 97 | 62 | 61 |
| layer M10 | 328 | 96 | 62 | 61 |
| lobula (total) | 1943 | 98 | 51 | 50 |
| layer LO1 | 117 | 98 | 68 | 67 |
| layer LO2 | 316 | 98 | 59 | 58 |
| layer LO3 | 276 | 98 | 55 | 54 |
| layer LO4 | 283 | 97 | 50 | 49 |
| layer LO5A | 244 | 98 | 49 | 48 |
| layer LO5B | 482 | 98 | 47 | 46 |
| layer LO6 | 224 | 98 | 43 | 42 |
| lobula plate (total) | 869 | 98 | 61 | 60 |
| layer LOP1 | 265 | 98 | 59 | 58 |
| layer LOP2 | 232 | 98 | 61 | 59 |
| layer LOP3 | 192 | 98 | 63 | 62 |
| layer LOP4 | 180 | 98 | 61 | 60 |
| accessory medulla | 7 | 97 | 55 | 54 |
| lamina | 223 | 16 | 31 | 5.1 |

## Supplementary Information

**Supplementary Fig. 1 (related to Fig. 5):**

**Summary of the anatomy and connectivity of all visual system neurons**

**Supplementary Video 1**: A selected neuron for each cell type in the visual system catalog. Cell names are shown, and the cells are color-coded to match the main groups of Fig. 1c.

**Supplementary Video 2**: The regions of the *Drosophila* visual system and the neurons of the 15 columnar cell types used to define columns (Fig. 2a).

**Supplementary Video 3**: Neuron populations from cell types selected to illustrate different patterns of sampling visual space with medulla coverage patterns. The four cells highlighted in Fig. 5e,f are featured.

**Supplementary Table 1**: Cell type names, counts, and neurotransmitter predictions for the inventory of visual neurons. The table lists all of the right-side instances (majority of cells, instance labels end with “_R”) followed by the left-side instances (instance label end with “_L”). Related to Fig. 1, Supplementary Fig. 1.

**Supplementary Table 2**: Fraction of synaptic connections outside the primary visual system regions. Related to Extended Data Fig. 1.

**Supplementary Table 3**: The bodyIDs and assigned coordinates for the 15 columnar call types of Fig. 2a. Related to Fig. 3.

**Supplementary Table 4**: T4 type bodyIDs matched to Mi1 bodyIDs for creating columns in LOP. Related to Extended Data Fig. 5.

**Supplementary Table 5**: Neurotransmitter ground truth (for training) and experimental validation data. Related to Fig. 4.

**Supplementary Table 6**: List of split-GAL4 lines and the cell types labeled by these lines. Related to Fig. 8.

**Supplementary Table 7**: Preliminary matches for the (male) optic lobe cell types to the annotated cell types in the (female) FlyWire dataset. Cell type names, counts and neurotransmitter predictions from both datasets are listed, together with URL links to visualizations of the neurons of the matched cell types. The neurotransmitter predictions that cannot be compared are greyed out, while the mismatched predictions are colored in red. Further details in Methods. Related to Extended Data Fig. 13.

## Methods

### EM sample preparation

The sample preparation followed our recently described methods^1,2^ and are reproduced here with minimal modifications from the recent manuscript on the “MANC” Ventral Nerve Cord (VNC) Electron Microscopy (EM) volume^3^ to preserve the consistency and accuracy of the reported methods:

Five-day-old males from a cross between wildtype Canton S strain G1 x w^11^^18^ were raised on a 12-hour day/night cycle and dissected 1.5 hours after lights-on. The main difference from previous work occurred during this dissection step. For this new sample we dissected the entire central nervous system (CNS) as a unit, including the brain, ventral nerve cord, and the neck. The main difficulty, requiring many attempts and extreme care, was dissection without damaging the relatively fragile neck connective. An undamaged neck connective is necessary to reconstruct an entire CNS. The optic lobe reported here is a subset of this complete CNS reconstruction. Samples were fixed in aldehyde fixative, enclosed in bovine serum albumin (BSA) and post-fixed in osmium tetroxide. Samples were then treated with potassium ferricyanide and incubated with aqueous uranyl acetate followed by lead aspartate staining. A Progressive Lowering Temperature (PLT) dehydration procedure was applied to the samples. After PLT and low-temperature incubation with an ethanol-based UA stain, the samples were infiltrated and embedded in Epon (Poly/Bed 812; Luft formulations). Collectively, these methods optimize morphological preservation, allow full-brain preparation without distortion, and provide increased staining intensity, enabling faster FIB-SEM imaging. Each completed sample was examined by x-ray CT imaging (Zeiss Versa 520) to check for dissection damage and other potential flaws and to assess the quality of the staining.

### EM sample preparation: hot knife cutting

The *Drosophila* CNS is roughly 700 µm wide and more than 1000 µm long, making it too large to image by FIB-SEM without milling artifacts. We therefore subdivided the brain right to left (vertical cuts in the orientation shown in Fig. 1a), and the VNC lengthwise (horizontal cuts in the orientation of Fig. 1a), both into 20 µm thick slabs using an approach developed during the hemibrain project^1^. This allowed imaging of individual slabs in parallel across multiple FIB-SEM machines. During embedding, the CNS was oriented such that the long axis of the VNC was perpendicular to the block face, with its caudal tip closest to the block face. The block was then trimmed into a fin shape^4^ (600 µm wide and several millimeters long in the cutting direction with a sloped trailing edge) to a depth encompassing all of the VNC and half of the neck, leaving the brain in the untrimmed part of the block. This VNC and half-neck were then sectioned into a total of 31 slabs, each 20 µm thick, using our previously described “hot-knife” ultrathick sectioning procedure^4^ (oil-lubricated diamond knife (Diatome 25° angle Cryo-Dry), 90°C knife set point temperature, mm/s). The remaining part of the block, containing the brain and upper half of the neck, was then reoriented and re-embedded in Epon so that the long axis of the brain was perpendicular to the new block face. This block was trimmed into a fin shape as before, and hot-knife sectioned (cutting speed of 0.025 mm/s) into a total of 35 slabs, each 20 µm thick. Each slab was imaged by light microscopy for quality control, then flat-embedded against a sturdy backing film, glued onto a metal FIB-SEM mounting tab, and laser-trimmed using previously described methods^4^. Each mounted slab was then x-ray CT imaged (0.67 µm voxel size) to check preparation quality, to provide a reference guide for FIB-SEM milling, and to establish a z-axis scale factor for subsequent volume alignment. All 66 slabs were FIB-SEM imaged separately, and the resulting volume datasets were stitched together computationally (as discussed in the section **EM volume alignment**) and used in the ongoing reconstruction of the entire CNS. The right optic lobe was contained in 13 of these slabs.

### EM volume imaging

The imaging methods in this section are reproduced with minimal modifications from the recent manuscript describing the MANC EM volume^3^ to ensure the consistency and accuracy of the reported methods:

The 35 brain and 31 VNC slabs were imaged using seven customized FIB-SEM systems in parallel over a period of almost a year. Unlike the FIB-SEM machines used for the hemibrain project^1^, this new platform replaced the FEI Magnum FIB column with the Zeiss Capella FIB column to improve the precision and stability of FIB milling control^5^. FIB milling was carried out by a 15-nA 30 kV Ga ion beam with a nominal 8-nm step size. SEM images were acquired at 8-nm XY pixel size at 3 MHz using a 3-nA beam with 1.2 kV landing energy. Specimens were grounded at 0V to enable the collection of both secondary and backscattered electrons.

### EM volume alignment

Alignment of the EM volumes was performed with an updated version of pipeline used for the hemibrain^1^, that was also used for the MANC VNC^3^ volume. The “render” web services were used for serial section alignment of the 35 brain slabs, followed by automatic adjustment for milling thickness variation^6^. The surfaces of the slabs were automatically identified using a combination of a hand-engineered^7^ and machine-learning-based cost estimation^3^ prior to graph-cut computation^3,7^, followed by manual refinements using a custom BigDataViewer^8^-based tool to interactively correct remaining issues. As with the MANC and hemibrain, the series of flattened slabs was then stitched using a custom, distributed method for large-scale deformable registration to account for deformations introduced during hot-knife sectioning. A custom BigDataViewer-based tool was developed to interactively help the automatic alignment in challenging regions. The code used for the “render” web services can be found at https://github.com/saalfeldlab/render and the code used for surface finding and hot-knife image stitching is available at https://github.com/saalfeldlab/hot-knife.

### EM volume segmentation

Segmentation was carried out as described in the MANC VNC volume^3^, but with some differences described below. Flood-filling network (FFN) inference was only applied in areas likely to contain segmentable tissue. To detect these, we used the following heuristic: the CLAHE-normalized images were downsampled to 32×32×32 nm^3^ voxel size and then filtered in-plane (XY) with a high-pass filter *I_hp_* = *I* + *min*3*x*3(255 − *I*), where *I* is the pixel intensity and *min*3*x*3 is the minimum function, applied convolutionally with a kernel size of 3 x 3 pixels. We then computed 2D (in-plane) connected components of the filtered images thresholded at a value of 220, and used the largest connected component within every section to exclude the corresponding parts of the EM images from segmentation.

We trained a semantic segmentation model for the volume by manually annotating selected segments from the FFN segmentation into 8 disjoint classes: “do-not-merge”, “glia”, “trachea”, “soma”, “non-glia-nuclei”, “neuropil”, “muscle”, “glia-nuclei”, and using these to train a convolutional neural network model to classify every voxel of the dataset at 16×16×16 nm^3^ resolution into one of these classes. For every segment, we then computed the corresponding class using a majority voting scheme and only included “soma”, “non-glia-nuclei” and “neuropil” segments in the agglomeration graph. The neural network used for this process had the same architecture as the FFN (“residual convstack”), but with all convolutional layers applied in the “VALID” mode.

Instead of manually annotating all nuclei in the dataset, we relied on the predictions of the semantic segmentation model. We computed 3D connected components of voxels classified as glial and non-glial nuclei, and postprocessed them twice according to the following procedure: apply morphological erosion with a radius of 5 voxels, recompute 3D connected components, and then remove components smaller than 10,000 voxels. We took the centroids of the remaining objects to be nuclei centers and disallowed distinct nuclei to be merged in the agglomeration graph. We applied a heuristic procedure for merging nuclei segments and surrounding soma segments. We applied morphological dilation with a radius of 5 voxels to every nucleus generated in the previous step. We next computed the number of overlapping voxels between these expanded nuclei and FFN segments of at least 10,000 voxels at 16×16×16 nm^3^ with a majority class of “soma” or “nonglia-nuclei”. All segments matched with a specific nucleus were then merged.

### EM volume synapse identification

We performed synapse prediction using the same methods as described in the MANC VNC reconstruction^3^. We obtained ground truth synapse data from the larger CNS sample and used it to train the networks for pre-synaptic T-bar detection and for post-synaptic partner detection The network weights for T-bar detection were first initialized using the previously-trained detector^3^, then fine-tuned using the available CNS training data. To quantify performance of the synapse identification within the optic lobe, 14 cubes of 300×300×300 voxels were randomly selected within the optic lobe volume, and synapses within each subvolume were densely annotated. In aggregate, the subvolumes contained 287 annotated T-bars and 2425 annotated post-synaptic partners. The overall precision-recall plots for T-bars alone, and for the synapse as a whole (both components) are shown in Extended Data Fig. 1a. This performance is substantially better than the accuracy achieved in the hemibrain^1^ and similar to the performance on the more recent MANC reconstruction^3^.

### EM volume Proofreading

Proofreading followed similar methods to those documented for the hemibrain^1^ and MANC^3^ connectomes and used a software toolchain for connectome reconstruction including NeuTu^9^, Neu3^10^, and DVID^11^. One difference for this volume is that we automatically generated point annotations for cell bodies (as discussed in **EM volume segmentation**), then added point annotations to neck fibers and nerve bundles, at early stages of the proofreading process. These annotations were fed back into the agglomeration stage, where they were used to forbid merges between the corresponding segments. This considerably reduced the number of falsely merged bodies, which are difficult and time-consuming to separate. An important difference in the optic-lobe connectome was the role that cell typing played in quality control and setting some proofreading priorities (see **Overview of cell typing)**. As we were typing cells from an early stage during the proofreading, we frequently compared the morphology and connectivity of cells assigned the same types, serving as a robust check for incomplete shapes or reconstruction errors. In the optic lobe dataset, the vast majority of cells and synapses belong to repeating cell types, so this parallel cell typing and proofreading effort was especially helpful in arriving at a high-quality connectome.

An important metric for evaluating the “completeness” of a connectome, or a region of a connectome, is the percentage of all synapses where both the pre- and post-synaptic partners belong to identified neurons. We provide this completion percentage for the optic lobe regions in Extended Data Table 1, and these summary metrics show this new connectome to be based on one of the highest-quality reconstructions to date. The completion rate is high, and relatively uniform across regions. Across the whole dataset, the connection completeness is 52.7% (54.2% when the partial lamina, discussed below, is excluded). This is considerably higher than the hemibrain, where the corresponding metric is 37.5%^1^.

The lamina completion metrics (Extended Data Table 1) are considerably worse than the other optic lobe neuropils. The lamina is the most peripheral neuropil of the optic lobe, where the axons of outer photoreceptors contact their targets. In this volume, the lamina was particularly difficult to reconstruct for two reasons: 1) it is incomplete with a noteworthy stair-step profile due to the trimmed edges of the 20µm slabs, and 2) the severed axons of photoreceptors are generally quite heavily stained and therefore were much harder to segment than other neurons in the volume. While it was not our original intention to analyze the lamina, the more complete segments of it are quite useful, especially for estimating the location of the eye’s equator^12^, as shown in the eye map of Fig. 3a. For this reason, and despite the sample’s limitations, we invested considerable effort in the lamina’s reconstruction, but the completeness metrics reflect the lower quality and completeness of the data for this neuropil. The heavily stained inner photoreceptors (R7/R8) that target the medulla also present very considerable challenges for segmentation and proofreading. Nevertheless, we proofread most of these cells, except for those in one patch of the medulla where these terminals were too fragmented to assemble into neurons. The distribution of these photoreceptors is detailed in Extended Data Fig. 4a. We estimate the counts of the missing cells in the section **Estimating the count of photoreceptors and lamina intrinsic neurons**.

### LM-EM volume correspondence

To compare neurons imaged by Light Microscopy (LM) and those reconstructed from EM data, we transformed the 3D coordinates of the whole central brain EM volume, including the optic lobes, into reference coordinates for a standard reference brain used for for LM data. The spatial transformation was obtained by registering the synapse cloud of the male CNS EM volume onto the JRC2018M *Drosophila* male template^13^, following an approach similar to that described for the hemibrain volume^1^. To produce an image volume, the synapse predictions were rendered at a resolution close to the JRC2018M template (512nm isotropic). Next, we performed a two-step automatic image registration procedure using Elastix^14^, with the JRC2018M template as a fixed image and the rendered synapse cloud as the moving image. The first step estimates an affine transformation, which is used to initialize the second step, during which a nonlinear (B-spline) transformation is estimated. We then manually fine-tuned this transformation using BigWarp^15^ to place 91 landmarks across the datasets to define a thin-plate spline transformation to correct visible misalignments. This defined an initial transformation that we used to warp neuron skeletons traced from the hemibrain dataset^1^ into the coordinate system of this new (male CNS) EM volume. We identified putative neuron correspondences between hemibrain neurons and preliminary neuron traces from this volume and used these to identify remaining misalignments. We corrected these in BigWarp by placing 82 additional landmarks, bringing the final total of landmarks to 173. The composition of the transformations from these steps (affine, b-spline, thin-plate-spline) is the spatial transformation from the JRC2018M template to the EM space. We also estimated the inverse of the total transformation using code found in https://github.com/saalfeldlab/template-building. As there are transformations already defined between JRC2018M, JRC2018F, and JRC2018U, establishing the correspondence for one of the three automatically does it for the other two. Fig. 8c,d,g and Extended Data Fig. 12b-f show many applications of the image registration to compare neurons from the optic lobe dataset to similar cells in LM and EM (hemibrain) images, using the JRC2018U template.

### Defining anatomical regions in the EM volume

Prior to segmentation, the neuropils were defined using synapse point clouds. The neuropil boundaries were initialized based on the JRC2018M *Drosophila* male template^13^ after registration with our EM volume, as described in section **LM-EM volume correspondence.** Then the boundaries of the right optic lobe regions were refined with hand-drawn corrections based on the grayscale data. While some boundaries are quite clear, others, such as those surrounding the inner chiasm of the optic lobe, are not sharp. The Regions of Interest (ROIs) for each neuropil were drawn with the goal of enclosing the maximum number of synapses within the corresponding neuropil (without overlapping with adjacent neuropils).We named the regions following the standard nomenclature for the optic lobe^16^ and central brain^17^. For the optic lobe we then used an iterative process to columns and layers, first reported in classic neuroanatomical studies^16,18,19^. We match all of the classically defined layers except for the 5^th^ layer of the lobula, which we divide into LO5A and LO5B following our previous definition from LM data^20^. This process is described in the section **Defining column centerlines and ROIs** and the section **Layer ROIs.** The resulting ROIs are available in neuPrint and are systematically named. Column ROIs are named by concatenating the neuropil, brain hemisphere, and hexagonal coordinates (e.g., LOP R col 15 21), and layer ROIs are named by their neuropil, brain hemisphere, and layer number (e.g., ME R layer 07).

### Connectome data access overview

The primary data source for our analysis is the optic-lobe:v1.0 dataset in neuPrint^21^, a Neo4J^22^ graph database. We access three levels of nodes in the database: (i) “Segment” (ii) “SynapseSet”, which are collections of synaptic sites representing T-bars and postsynaptic densities (PSDs) and (iii) “Synapse” which refers to the individual synaptic site. These nodes have a hierarchical relationship wherein the “Segment” nodes contain “SynapseSet” nodes, and the “SynapseSet” nodes contain “Synapse” nodes. Both the “Segment” and “SynapseSet” nodes connect to other “Segment” and “SynapseSet” nodes, respectively.

These connections are calculated based on the relationship between individual “Synapse” nodes. All “Synapse” nodes derive from an “Element” type and inherit their properties, such as the 3D (x,y,z) coordinate in the EM volume. The overwhelming majority of “Segment” nodes correspond to small, unproofread, and unnamed fragments with few synapses and are not relevant to most analyses. To enable faster queries, the subset of “Segment” nodes that are relevant for typical analyses are additionally labeled as “Neuron” nodes. To qualify as a Neuron, a Segment must have at least 100 synaptic connections, a defined “type,” “instance,” or known soma location. In most data analyses presented here, we primarily work with named Neurons, i.e. “Neuron” nodes with defined “type” and “instance” properties. We ignore non-Neuron Segments and further exclude Neurons whose types are suffixed with “_unclear”.

For full clarity, in the graph database syntax^21^, the nodes are “:Neuron” “:Segment” and have a “:Contains” relationship to “:SynapseSet”s which have a “:Contains” relationship with “:Synapse”s. “:Neuron”s and “:SynapseSet”s have “:ConnectsTo” relationships to other “:Neuron”s and “:SynapseSet”s respectively. These relationships are calculated based on the “:SynapsesTo” relationship between individual “:Synapse” nodes.

In neuPrint data loaders, regions of interest (ROIs) are defined as connected volumes, i.e. spatially contiguous collections of voxels. Consequently, each element with a spatial location can be assigned to any number of ROIs, such as ROIs we defined for layers, columns, and neuropils. This is a simple assignment for point objects like postsynapses, presynapses, and column pins (defined below). Since synaptic connections are typically based on a single presynapse and several postsynaptic densities, neuPrint uses the convention of assigning the connection to the ROIs of the postsynaptic site.

While our analysis focuses on the visual system, several cell types have processes in the central brain. Although proofreading and naming efforts are ongoing and, therefore, not as complete as for the visual system, the reconstructed arbors are included in the dataset, as skeletons (and meshes) but without the synaptic connections. In the database, the aggregated summaries of synaptic sites and their innervated brain regions representing the current state of the reconstruction effort are included as properties of neurons. These aggregations rely on the confidence threshold of 0.5 set for the optic-lobe:v1.0 dataset. As a result, the summaries are consistent with the late 2023 snapshot of the reconstruction (optic-lobe:v1.0), but the synapse properties of “:Neuron”s differ from the number of “:Synapse”s they contain (as central brain connections are incomplete, only half the connection is present).

Within the python environment, neuprint-python^23^ provides access to the neuPrint servers and translates function calls to Neo4J Cypher queries (https://connectome-neuprint.github.io/neuprint-python/docs/). We rely on the carefully selected, reasonable default values of the neuPrint database (the most relevant is the synapse confidence threshold of 0.5). To simplify analysis, we aggregate data as close to the source as possible either in pandas DataFrames or using neuprint-python’s “fetch_custom” function with optimized Neo4j Cypher queries. The code we share in the repository (see **Code availability**) uses a combination of these methods for data access. For estimating the spatial location of an element, such as a synapse, in relation to columns and layers, we added the data type “:ColumnPin” to neuPrint, which represents points positioned along the center lines of each column (of the medulla, lobula, and lobula plate) and with properties representing the identity of and depth within a column.

In addition to the neuPrint Neo4J database, we provide neuron morphologies and boundaries of ROIs in the form of meshes through Google Storage buckets. Meshes are generated from ROIs with marching cubes, followed by Laplacian smoothing and decimation by quadrics (https://github.com/sp4cerat/Fast-Quadric-Mesh-Simplification). Meshes for brain regions, columns, and layers are available in three different spatial resolutions between 256nm and 1024nm (per side of voxels), while meshes of most individual neurons were generated from 16nm voxels.

Data specific to our analysis and retrieved by other methods are stored inside the code repository in the “/params” folder, such as:

- The “Primary_cell_type_table” groups the named neuron types into the 4 main categories OLIN, OLCN, VPN, VCN, and “other” (see section **Cell type groups**).
- The identification of example cells (and their bodyIDs) for each neuron type, which we call “star neurons” inside the “all_stars” spreadsheet (see section **Gallery plots**).
- We provide some heuristically defined “Rendering_parameters” in a second spreadsheet (see section Gallery plots).
- Additional files contain parameters used in the generation of pins and layers: “pin_creation_parametera” (see **Defining column centerlines and ROIs** and **Layer ROIs**).

The data inside the neuPrint database, precomputed meshes stored at the Google Cloud, and the data inside the params folder is sufficient for replicating our data analysis and figures, for example using the source code we provide in our repository (see **Code availability)**.

We provide our code in “notebooks” (literate programming^24^) to explain analytical steps or command-line scripts for long-running processes. Both rely on functions for prototypical implementations or object-oriented components shared between several of our applications.

### Visualization of reconstructed EM neurons

The neurons in our dataset are all in the *right* optic lobe, that is, on the *fly’s right side* of the brain. However, we represent this optic lobe on the left side of our images (see Fig. 1a), as this is the view most familiar to anatomists (e.g. looking down at a sample in a microscope) and is directly comparable to LM data. To provide further intuition for these perspective changes, we show the coordinate system of the medulla’s correspondence to retina (and lenses) of the right eye, when viewed from inside the brain and looking outwards (Fig. 3a).

Throughout the manuscript and associated materials, EM-reconstructed neurons are rendered using several methods introduced here, and detailed in later sections. Neurons are generally shown as an orthographic projection of a 3D view; most are programmatically rendered in Blender^25^ and described in **Pipeline for rendering neurons**.

Many visual system neurons are shown on an optic lobe “slice” view, an anatomical section containing neurons of interest and relevant brain regions. Parts of the neurons may lie outside the sliced volume, so we use a “pruning” step to show layer innervations (required due to curvature of the optic lobe regions).

- The **Pipeline for rendering neurons** section describes the method for producing the images in: Fig. 1c,e; Fig. 2a,b,e; Fig. 4c; Fig. 5a,e,f; Fig. 9b; Extended Data Figs. 4e, 7c, 12a, and the Cell Type Catalog (Supplementary Fig. 1).
- For “full brain” views of rendered neurons we used methods described in **Pipeline for rendering neurons** for producing the images in: Fig. 6; Fig. 7; Fig. 9b; Extended Data Figs. 10, 11.
- We used some manually produced, custom Blender visualizations for the images in: Fig. 1b,f; Fig. 5c; Extended Data Figs. 5c,d, 7a.
- The web-based viewer, neuroglancer^26^, was used to produce the images in: Fig. 1a; Fig. 3b,d; Extended Data Fig. 1e,f; Extended Data Fig. 13b,c.
- The Light Microscopy (LM) images and the comparisons with EM-reconstructed neurons registered to a standard reference brain are produced by methods described in **Split-GAL4 lines expressing in visual system neurons** and shown in: Fig. 2b; Fig. 3c; Fig. 4d; Fig. 8; Extended Data Figs. 2b, 12.

### Pipeline for rendering neurons

While neurons have complex, extended 3D shapes, it is remarkable that 2D visual representations of neurons have typically served as extremely rich descriptions of cell types across generations of neuroscientists, from hand drawings of Golgi stained neurons^16,18^ to computer-generated ray traces of reconstructed EM volumes^27^. Following open-science principles, we developed a rendering pipeline based on the state-of-the-art open-source software Blender^25^ that allows us to produce static printable images similar to best practices developed over more than a century of communicating microscopic structures, as well as movies for accessible science communication. The Cell Type Explorer web resource contains interactive figures^28^ affording the individual exploration of the neuron’s morphology.

To render detailed, high-resolution printable images and movies, we developed a novel software pipeline and integrated neuVid^29^, a pipeline for making videos, into our workflow. In both cases, we begin with a bodyID for individual neurons (for which we most commonly use “star neurons” selected from a curated list of representative neurons for each cell type, listed in the “all_stars” spreadsheet in the code repository “/params” directory). We then query neuPrint for additional information about this neuron, such as the layer and column innervation, cell type names, synapses, and neighboring cells (of the same type). Based on five templates, we combine this information with material properties and virtual camera location (“Rendering parameters” spreadsheet in the code repository “/params” directory) into JSON description files. In a next step, we use either our own image pipeline or neuVid for movies to generate the output. Based on the image and movie descriptions, we download required meshes for neurons and brain regions from a google cloud storage bucket (see **Data Availability**) and cache them temporarily inside the “/cache” folder in the local file system.

The rendering is done using Blender for both the movies and the images. For the movies we rely on the neuVid package^29^ to move the camera and assemble individual frames of high-resolution images. We used this to make Supplementary Videos 2 and 3, and neuron-specific videos that are linked from the Cell Type Explorer web resource.

The rendered, static images of neurons are taken from two main directions, similar to the frontal and side view of the optic lobe in Fig. 1b. For the full brain view (e.g. Fig. 6, 7), we placed the camera anterior to the animal with a posterior viewing direction of the orthographic projection. For the optic lobe “slice” view, we position the orthographic camera laterally and within a few degrees of the dorsoventral axis (similar to the side view of Fig. 1b). We developed the rendering pipeline to allow free positioning of the camera, which we take advantage of in a few of our figures. In our standard renderings with the full brain view, we produced whole brain images for neurons with midline-crossing projections and produced images of slightly more than half a brain for all other neurons. To render representative examples of all optic lobe neurons in an optic lobe “slice” view, we established three camera positions: an equatorial view used for most neurons, as well as ventral and dorsal views that are better suited for visualization of a subset of neurons and types. In Supplementary Fig. 1, the slice position of each panel is indicated with a letter: E, D, V.

The full brain view benefits from outlines of the different brain regions, specifically the two optic lobes and the central brain. The 3D meshes are projected onto a plane orthogonal to the viewing direction of the camera and spatially towards the posterior end. The neurons are then plotted in their original coordinates and rendered in Blender.

The optic lobe slice views (e.g. Fig. 1c, 2a, and all the panels of the gallery in the Cell Type Catalog, Supplementary Fig. 1) required quite a bit of fine-tuning to establish representative visualizations of individual neurons that capture their layer-specific arborization patterns, while visualizing the layers, and also capturing the axons, cell body fibers, soma locations, and, where relevant, central brain arborizations. We obtained a visual representation of the layers of the visual regions by intersecting the medulla, lobula, lobula plate, and accessory medulla meshes with a rectangular cuboid of around 2.5nm thickness near the location of the neuron. For the movies, we use a similar approach with 100nm thick cuboids. The intersections between layer meshes and cuboids are then represented as a wireframe around the edges of the newly created mesh. We use the wireframe to emphasize the boundary of the slice with a darker color. The neurons themselves are sliced with a thicker rectangular cuboid. Depending on the cell and innervation pattern, we use between 750nm and 3µm thick cuboids to constrain the visible parts of a neuron. For most neurons, we use slices only inside the layered brain regions (medulla, lobula, and lobula plate) while we show all of the reconstructed arbors outside these three regions. This allows us to capture the layer-specific arborization patterns (due to the curvature of the optic lobe regions, projections through thick slices obscure most of these patterns). For about 100 neurons, we extend the slice outside the layered brain regions, and for 7 cell types (Cm-DRA, Cm27, Li37, LPi14, LPi4a, LPT30, and Mi20), we manually pruned arbors to achieve the visualization goals described in the section on the **Gallery Plots**.

To show innervation patterns in specific medulla layers, such as in Fig. 2e and Fig. 5e, f, our optic lobe “slice” view was modified so that the camera was turned approximately 90 degrees for a face-on view of the medulla layers. In these images, only the relevant medulla layer was rendered.

For renderings that include synapses (Extended Data Fig. 7c and 12a), we only show synapses to/from the indicated neurons, and only the synapses inside of the bounding rectangular cuboid are rendered. The synaptic sites are represented by a sphere and assigned a visually pleasing diameter.

The generated raster graphics are stored on the local file system. In the final step of assembling individual panels, or the galleries of images, such as in Fig. 6 and 7 and Supplementary Fig. 1, we use the JSON description to add text and combine several images into a single page using the pyMuPDF^30^ library.

We defined workflows for long-running and multi-step computations with snakemake^31^. While we prototyped and could produce most images on a personal computer running Blender, with >700 cell types and several views of each type to render, we found snakemake to be very helpful in distributing the workload across the nodes of a compute cluster.

### Cell type groups

To facilitate further analyses and the data presentation, we assigned each cell type to one of five main groups: OLIN, OLCIN, VPN, VCN and other. Assignments to groups were done at the cell type level, not the cell level. In general, OLINs (optic lobe intrinsic neurons) and OLCNs (optic lobe connecting neurons) are defined as cell types that have essentially all their synapses (>98% of both input and output connections) within the optic lobe, with OLINs restricted to a single region of the right optic lobe: lamina, LA(R); medulla, ME(R); lobula, LO(R); lobula plate, LOP(R); accessory medulla, AME(R) (the abbreviated label of each is the name of the corresponding ROIs in neuPrint). OLCNs are the cell types that connect two or more of these regions. VPNs (Visual Projection Neurons) receive substantial inputs in the optic lobe and project to the central brain, VCNs (Visual Centrifugal Neurons) also connect the optic lobe and central brain but with the opposite polarity. “Other” was used primarily for cell types with central brain and optic lobe synapses with a comparatively small proportion of their connections in the right optic lobe (and therefore did not fit well into the VPN or VCN groups).

Photoreceptors (R1-R6 and R7/R8) were included with OLIN types, as this matches the distribution of their synapses. However, they could alternatively be considered OLCN cells (given their projection patterns).

Ascending and Descending neurons were placed in the “other” group. We note that some of these have substantial optic lobe synapses and, therefore, could alternatively be classified as VPN (e.g., DNp30) or VCN (e.g., Ascending_TBD1) types.

While most cell types can be unambiguously placed into one of these groups based on synapse distributions and overall morphology, there are more marginal cases that required further criteria, in particular for deciding between VPN, VCN and “other” groups for some cell types. For most cells, the assignment was based on relative numbers of synaptic connections. For a VCN (VPN) we required >10% of its total output (input) connections to be located within the optic lobe. The AME was treated in the same way as other OL regions for this purpose (placing, for example, the s-LNv clock neurons in the VPN group). Additional criteria used when classifying cells as VPNs or VCNs included the proportion of downstream synaptic connections relative to total synaptic connections in the optic lobe and central brain, the grouping of related cell types, and details of arbor structure (for example, for cells that primarily receive input in one part of the optic lobe and have major output connections in both another optic lobe region and the central brain). When grouping the cell types, we aimed to keep the number of cell types in the “other” category low; we, therefore, chose a relatively low minimum proportion of optic lobe synapses (10%, see above) for VPNs and VCNs and (tentatively) classified a few ambiguous types as VPN or VCN based on the balance of the available evidence.

### Estimating the count of photoreceptors and lamina intrinsic neurons (related to Fig. 1)

To provide a nearly complete count of cells in the optic lobe (Fig. 1 plus the estimate listed in the figure legend), we must account for two technical limitations of the dataset. First, because the lamina is incompletely contained in our volume, the number of reconstructed Lai and R1-6 cells is an underestimate of the total count of these cells. Second, due to poor segmentation of some photoreceptor cells, not all R7 and R8 photoreceptors in the medulla could be reliably identified, resulting in another undercount (Extended Data Fig. 4a). An approximate number of R1-6 was obtained as the total number of non-edge columns times six (again rounded to the nearest ten). Finally, we counted labeled Lai cells (labeled using split-GAL4 driver SS00808) in three optic lobes and used the rounded average (210) as our estimate for Lai.

### Overview of cell typing (related to Figs. 1,2)

We used both morphology and synaptic connectivity for grouping cells into types. Morphology-based typing was generally done by direct visual inspection of reconstructed EM bodies; the combination of features such as cell body positions and the number, size, and layer position of their arbors gives cells of many optic lobe cell types a distinct appearance, that with experience, is often directly recognizable. Since the initial typing was carried out in parallel to proofreading, our review of cell morphologies also served to identify apparent reconstruction errors and to direct targeted proofreading efforts. Morphology-based cell typing was facilitated by the considerable amount of available prior information on optic lobe cell types^16,20,32–35^ both from published descriptions and unpublished light microscopy analyses (see **Split-GAL4 lines expressing in visual system neurons**). Cell type annotations were performed in an iterative fashion. As the proportion of typed (or preliminarily typed) neurons increased, we increasingly relied on synaptic connectivity (see **connectivity-based cell clustering**) to confirm or refine annotations, identify outliers, and place cells into preliminary groups for further review and annotation based on morphology.

Over several cycles of this procedure, and because of the high-level of completeness in our dataset, we assigned types to (nearly) all reconstructed EM bodies above a minimum size (∼100 combined in- and output synapses). All EM bodies (except many in the lamina), that our team of expert proofreaders believe are individual neurons, have been annotated with a type and instance. To do so, we used all available evidence to label some atypical cells (e.g. near the margins) and gave about 350 remaining bodies names including “_unclear”. These unclear types provide some information (e.g. “R7_unclear”) but indicate the currently unresolved typing status (about ∼300 of these “unclear” bodies are R7 or R8 photoreceptor neurons that we considered too incomplete for more precise typing). Most of the remaining unnamed bodies in the dataset above the 100-connection threshold are smaller fragments that are primarily severed parts of annotated neurons and bodies in the lamina for which annotation and proofreading are less complete than in the other parts of the optic lobe. To support proofreading efforts and connectivity-based clustering, we also assigned candidate types to some smaller fragments; most of these were eventually merged with other EM bodies, but some such “fragment” annotations are still available in the “instance” field of neuPrint. Additional aspects of cell typing are addressed in the sections on **Cell type nomenclature**, **cell type groups**, and **connectivity-based cell clustering**.

### Cell type nomenclature (related to **Figs 1,2**)

There can be no perfect nomenclature for this diverse set of neurons that respects the historical usage while systematically describing the details of each cell type. In addition to following the historical precedent, our aim was to use short names which are typically easier to remember and use, and in most cases give some indication that (and often how) a cell type is associated with the visual system. Most of the cell type names used in the visual system dataset, both the existing names and new names introduced here, consist of a short “base name” for example, “Tm” “Dm” or “LoVP” that provides a broad classification of a neuron, followed by a number that distinguishes the individual types within this group.

Many of the individual cell type names for OLINs and OLCNs are based on the systematic names provided in the Golgi studies by Fischbach and Dittrich^16^, which provided many of the base names we extended for naming new cell types. In addition to Fischbach and Dittrich, sources of names (and descriptions) for previously described neurons include: several OLINs and OLCNs^33,36–40^, Dm and some Pm cells^32^, Tm5a/b/c^41^, DRA neurons^42,43^, LCs^20,34,35^, LPC/LLPCs^44^, MeTus^45^, LPTs^46^, OA neurons^47^, and many VCN/VPNs and some Lis^1^. For new names we introduced some additional cell type base names, mainly as part of a new systematic naming scheme for VPNs and VCNs.

Most new names for VPNs or VCNs follow a format that starts with “Lo” “Me” or “Lp” indicating the main region a cell type has input synapses (for VPNs) or output synapses (for VCNs) in, followed by “VP” or “VC” and a number. Some VCNs have prominent outputs in 2 or more regions, in which case we use “OL” instead. For bilateral cells, an additional indicator for the regions in the second (left) optic lobe is included after “VP” or “VC” (e.g. MeVPMe1). In addition to these systematic names, a few new names are based on existing naming schemes, extending the base names that had been used for groups of similar cell types (e.g LC10e, LLPC4).

We kept many hemibrain names for VPN and VCN neurons^1^, but in cases where prior names omit any indication of prominent visual system connections, we introduced new synonyms in place of names based on central brain regions (e.g. LoVC16 for “PVLP132”). For the (few) cell types with central brain and optic lobe synapses that were not classified as VPN or VCN but as “other” (described in **Cell type groups**), we used their hemibrain names. In 5 cases, we applied a placeholder name including “_TBD” since we expect that future central brain data will suggest a more appropriate designation.

We also introduced two new cell type base names, “Cm” and “MeLo.” The Cm cells are “central medulla” OLINs, named to complement the existing “Dm” (distal medulla) and “Pm” (proximal medulla) base names. This base name is applied to medulla intrinsic neurons with their main arbors in layers M6 and M7. The “MeLo” base name is a simple contraction of the first two letters of medulla and lobula, and is applied to OLCN cell types that connect the medulla and lobula but do not project through the inner chiasm (in contrast to “Tm” cells). “Cm” and “MeLo” replace the “Mti” and “ML” labels that we had used for a small number of cell types from these groups in prior work^42^.

Some established names for visual system cell types include letters to indicate further divisions beyond a shared base name and number, for example “LC10” types are split into LC10a, LC10b, etc. In most cases, we have continued to use these existing names (and introduced a few names of this type, for example, to subdivide an existing type). The use of letters and numbers, at least in the visual system, has not been standardized, i.e. LC10a and LC10b are not necessarily more closely related than say, LC6 and LC16. We also do not make a formal distinction (as done in the hemibrain dataset^1^ for central brain neurons) between types identified through morphology vs. connectivity. Many cell types that are most readily distinguished by connectivity still show some other anatomical differences, at least at the population level. In two cases, we name subdivisions of cell types for which the existing names already include a letter (LC10c-1 / LC10c-2 and LC14a-1 / LC14a-2) but do not advocate for more widespread use of this practice.

For cell types with morphological variability in features that are suggested by base names, the same name was applied to all cells of the type. For example, several Tlp13 cells have branches into the medulla that make them resemble “Y” neurons, but as this is not a feature of the entire type, we keep the “Tlp” base name.

There are many gaps in the numbers attached to base names. For example, our list of TmY neurons starts with TmY3 and TmY4, skipping “TmY1” and “TmY2.” We do this to minimize confusion with prior usage. Some cell types illustrated and named by Fischbach and Dittrich (for a long time, the main, if not only, source of cell type information for neurons in the *Drosophila* optic lobe) appear to represent variants of other types. For example, Fischbach’s “Tm6” and “Tm14” both match our Tm6; “Mi8” is likely an atypical Mi1, and “Tm15” and “Tm20” both resemble our Tm20. Second, to avoid incorrect matches with earlier studies (some cell types are difficult to match to the limited information available from single-view Golgi drawings), several recent efforts by us and others have started with non-overlapping sets of numbers when naming additional types (a practice continued here for many cell types, for example, the Tm cells).

### Summarized inventory of visual neurons and connectivity

Figure 1 shows a summarized overview of the visual neurons contained in the optic-lobe:v1.0 dataset. Supplemental Table 1 lists the 727 unique cell types and 770 unique cell instances.

For Fig.1d, we consider all the cell types that belong to the 4 main neuron groups defined in **cell type groups**—OLINs, OLCNs, VPNs, and VCNs—and report the number of input and output connections for each cell type. There are different ways to count connections in the dataset, so it is important to clarify the conventions we use. A “Synaptic Connection” (or simply connection) is a paired presynapse on one body and a matched postsynapse on another. Autapses (self-connections) have been expunged from the dataset, and most of our analysis is restricted to named neurons. Since connections are directed and between two bodies, each side of the connection can be counted as an “input connection” or an “output connection.” The counts in Fig. 1 report these input and output connections for the 160 cell types listed in Fig. 1d (pooled across all neurons of each type) or for all cells in each group (Fig. 1e) or by region and group (Fig. 1f). Fig. 1 is the only place we report these rawer counts for connections. In the remaining analyses and materials, we report connections limited to only those between identified (named) cells, so the counts found on, e.g., the Cell Type Explorer web resource are expectedly lower (average connection completion percentage for the dataset is ∼53%). “Presynapses” is used for counts of T-bars (presynaptic active zones). “Postsynapses” is used to refer to counts of PSDs opposing presynapses. Because synaptic connections are asymmetric, with one presynapse typically opposed to multiple postsynapses, the count of postsynapses in a neuron is identical to the number of input connections, but the count of presynapses is not the same as the number of output connections. This difference is related to the neurons’ “fan-out,” which is examined in Fig. 4i. There is also an asymmetry in completion percentage between presynapses and postsynapses (Extended Data Table 1), which explains why the output connection numbers are higher than input connections when querying the database for neuron connections (as in Fig 1). Nearly all (∼97%) outputs (via presynapses) have been assigned to a neuron, while ∼55% of post-synapses are assigned to named neurons.

In Fig.1f, a neuron was assigned to one of the five optic lobe regions if >2% of the neuron’s summed presynapse and postsynapses are contained within the corresponding region ROI. Similarly, a neuron type was assigned to one of the optic lobe regions if >2% of all the summed presynapse and postsynapses of all the neurons belonging to that cell type are within the corresponding region ROI. All five lamina tangential (Lat) types are confirmed to have arbors in the lamina (LM data, not shown), but because the distal lamina is not in the EM volume, we capture very few of their synapses and therefore some Lat types are not counted in Fig. 1f. For Fig.1g we considered all the connections (>1) between the 727 cell types and report the mean number of connected input cells and output cells for each cell type.

### Connectivity-based cell clustering (related to Fig. 2, Extended Data Fig. 2)

To group cells into types by connectivity, we used aggregate connectivity to named cell types as the basis for clustering. For each reconstructed EM body to be clustered, we calculated the sum of its connections, further split into inputs and outputs, to all cells of each of the named cell types in the dataset (not including connections with cells with “_unclear” type annotations). Connections with cell types with synapses in both optic lobes were further split by instance (right and left side) since the connectivity in the right optic lobe of the left and right instances of cells of the same type can differ substantially. Small EM bodies annotated as “fragments” of a cell type (e.g. the instance “MeLo9_fragment_R” in neuPrint) were included with the corresponding cell types when calculating the summed connection weights. Excluding “unclear” cells and adding “fragments” was mainly important during the active proofreading stage of the reconstruction, when many EM bodies still had obvious reconstruction errors or were clearly incomplete. Finally, since the dataset reported here does not include detailed connectivity in the central brain, we only used connections within the right optic lobe, which for this purpose was the sum of connections in the LA(R), ME(R), LO(R), LOP(R) and AME(R) ROIs.

The resulting connectivity table was used as input for hierarchical clustering. Clustering was performed in python using the scipy.hierarchy^48^ and fastcluster^49^ libraries. We used cosine distance as metric and Ward’s linkage method. This choice was primarily based on the observed excellent agreement between our expert-curator-led morphology-based cell typing (not to be confused with the independent clustering based on quantified morphology described in **Morphology clustering**) and connectivity clustering for well-characterized cell types (see Fig. 2 and Extended Data Fig. 2). The key aspect of using cosine distance appears to be the normalization that is part of calculating this metric; L2 normalization of each row of the connectivity table followed by clustering with Ward linkage and Euclidean distances produced very similar results. Clustering output was divided into a preselected number of clusters (using scipy.hierarchy.fcluster with the “maxclust” criterion). For morphologically distinct cell types, connectivity clustering primarily served as a method for preselecting cells for subsequent morphology-based typing. For other cell types, connectivity was the primary determinant of cell groupings and reviewing cell morphologies was mainly used to identify outliers such as cells with apparent reconstruction errors. We note, however, that for several cell types initially separated by connectivity clustering, we were able to identify at least some morphological differences between the two cell populations (e.g. differences in arbor spread or cell body locations).

To illustrate clustering of larger groups of cells, we used 15 columnar cell types (Fig. 2d) and OLIN neurons of the medulla with ≥ 10 cells per type as examples (Extended Data Fig. 2a). For the first group, we set the number of clusters to the number of types (in this case independently defined by morphology) and observed a one-to-one correspondence between cell type annotations and cluster assignments. For the second example, we selected a number of clusters slightly exceeding the number of types (80 for the 68 medulla OLIN with ≥ 10 cells/type). In this case, while most clusters only contain cells of a single type, some cell types are not separated (but can be by reclustering) and some other types (combined as one type based on shared features, spatial coverage, or genetic information from driver lines) are split into multiple clusters. This example illustrates why additional criteria (as mentioned below) are required to decide which clusters should be annotated as distinct types.

Connectivity differences between two cell populations do not automatically imply that these represent different types. For this reason, we considered many factors, including cell morphology, consistency and magnitude of the connectivity differences and, when available, genetic information in the form of split-GAL4 driver line patterns to help decide when to split cells into separate types. In addition, we relied on the spatial distribution of repeating neurons in the optic lobe—many cells form mosaics covering a region (Fig. 2g, Extended Data Fig. 3). For this analysis, we used connectivity clustering to split the cells of existing types or combinations of similar types into two (or more) groups and plotted the positions of individual cells in each group (calculated as the center of mass of the positions of their synapses). Positions were transformed into a 2D representation by plotting the first two principal components (obtained for a set of preselected synapses for each brain region to standardize the view across cell types; denoted as e.g. PC1_ME_ and PC2_LO_, for the first and second PCs, for viewing the synapses of candidate cell types in the medulla and lobula, respectively). In general, we considered the combination of a split into two regular, overlapping patterns (mosaics), especially when combined with consistent connectivity differences, as a basis for dividing a cell population into two types. Extended Data Fig. 3 shows examples of the outcomes from applying this method, ranging from clear cases for splitting cells with overlapping mosaics, to regional patterns or patterns with no clear spatial structure along with only modest connectivity differences, which were not further subdivided.

We note that some of our cell types consist of single cells that we named individually (often applying established names) because the available evidence suggests they are uniquely identifiable. However, some cells with unique names belong to small groups of types that together cover part or all of the columns and which could alternatively be combined using similar coverage criteria (examples are the HS cells, typed separately as HSE, HSN and HSS, or the CH cells, typed separately as DCH and VCH).

### Assigning neurons to medulla hexagonal coordinates (related to Fig. 3)

We assigned neurons to a total of 892 hexagonal coordinates in the medulla, in a multistep process that iteratively refined the assignments. First, we manually assigned medulla coordinates to individual cells of 11 types using presynapses and postsynapses of the following connections: L1-Mi1, L2-Tm1, L3-Mi9, L4-Tm2, L5-Mi4, and L3-Tm20. Using the spatial distribution of the medulla presynapses of these 11 cell types, we calculated a straight-line central axis (a “proto-column”) for every coordinate using the first principal component.

Second, we used these linear axes to allocate hexagonal coordinates for 13 cell types using the following paired connections in the medulla: L1 (L1-Mi1, L1-L5, L1-C3), Mi1 (L1-Mi1, L5-Mi1 Mi1-C2), C3 (L1-C3, L5-C3, Mi1-C3), T1 (C3-T1), L2 (C3-L2, L2-T1), Tm2 (L2-Tm2), Tm1 (L2-Tm1, C3-Tm1), L5 (L1-L5, L2-L5, L5-Mi1), Mi4 (L5-Mi4, Mi1-Mi4), Mi9 (Mi4-Mi9, Tm2-Mi9, C3-Mi9), Tm20 (Mi4-Tm20, Tm1-Tm20, L2-Tm20), C2 (L1-C2, Mi1-C2, L5-C2), L3 (L3-Mi9, L3-Tm20, L3-Mi1). For every paired connection, synapses were assigned to the nearest coordinate axis, and individual cells were labeled with the mode of the synapse assignments. For individual cells where paired connections labeled different columns, we manually inspected the cell’s location in relation to the cells of the same type in neighboring coordinates to assign the coordinate location. We also used manual inspection in relation to its neighbors to verify coordinates where cell types were missing or duplicated, and to verify the assignment of every cell.

Third, we used the presynapses from these 13 cell types to calculate prototypes for the curved central axis for every column, which we refer to as columnar “pins” or column centerlines (described below). These prototype column pins were then used to automatically assign columns to the following 15 cell types: L1, L2, L3, L5, C2, C3, Mi1, Mi4, Mi9, T1, Tm1, Tm2, Tm4, Tm9, and Tm20. As before, we visually inspected the location of every cell in relation to the cells assigned to neighboring coordinates to verify coordinates where cell types were missing or duplicated and to verify the assigned coordinate of every cell. The final coordinate assignments are summarized in Fig. 3a and Supplementary Table 3. We do not use L4 cells, which have multi-columnar processes, or Tm3 cells, which are not reliably assignable to single the common coordinates of the other cell types. By using a large set of neuron types, the identification of columnar coordinates was robust to variability within cell types and reconstruction errors.

### Assigning T4s to Mi1s to extend the coordinate system to the lobula plate (related to Fig. 3, Extended Data Fig. 5)

To create columns in the lobula plate and a mapping between medulla and lobula plate columns, we used the neurons of each T4 type: T4a, T4b, T4c, and T4d. We used T4s for 3 reasons: (1) The axons of each type innervate one of the four layers of the lobula plate, (2) their dendrites receive strong Mi1 input, and (3) there are roughly as many T4 neurons of each type as there are medulla columns (T4a: 849, T4b: 846, T4c: 883, T4d: 860 and Mi1: 887). The major difficulties in assigning T4 neurons to medulla columns are: (i) T4 dendrites are not strictly columnar but innervate ∼6 medulla columns, and (ii) there are many cases where a unique assignment of individual Mi1 to individual T4 of each type is not possible. But once all T4’s dendrites are assigned to medulla coordinate, we can use the same coordinate assignment for its axons in the lobula plate, linking medulla columns to lobula plate columns.

To achieve a nearly unique assignment between T4 neurons of each type and Mi1 neurons, we set up a global optimization problem (called “maximum weight matching” in graph theory). First, for each T4 type, which we denote as T4x, we create a connectivity matrix 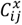 containing the number of connections between Mi1 neuron *i* and T4x neuron *j*. Then, to account for the bias in the number of synapses in different parts of the medulla, we normalize each connection by the total number of connections of each a Mi1 neuron to all T4x neurons, i.e., we define a normalized connectivity matrix 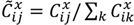. Finally, we obtained putative Mi1-T4x assignments by using 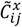 as the cost matrix in the Python linear_sum_assignment function from the scipy.optimize library^48^ (setting the parameter “maximize” to “True”).

Putative Mi1 to T4x assignments were rejected if the distance between the neurons was too large. We measured the distance between an Mi1 neuron *i* and T4x neuron *j* as 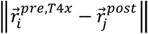 where ‖. ‖ denotes the Euclidean norm, 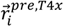 is the mean position of presynapses from Mi1 neuron *i* to any T4x neuron, and 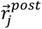 is the mean position of postsynapses of T4x neuron *j*. The distribution of distances between putative Mi1-T4x pairs showed a gap around 8μm (which is approximately the distance between columns in medulla layer M9), which we chose as the distance threshold above which we rejected the putative Mi1-T4x assignment. The results of this assignment procedure are detailed in Supplementary Table 4 and summarized in Extended Data Fig. 5.

To create columns in the lobula plate, we defined valid assignments as groups of at least 3 T4s of different types assigned to the same Mi1. The mean position of presynapses on the axons of these T4s provided us with at least 3 points with which to create the lobula plate columns. This procedure created 712 Mi1-T4x valid groups with all four types and an additional 75 groups with three types.

### Defining column centerlines and ROIs (related to Fig. 3)

Column ROIs subdivide a neuropil ROI and are based on the center lines of columns (“pins”). The basic idea of a column pin is that it is a smooth, slightly bent line that goes through the center of synapses of certain neurons assigned to the same hexagonal coordinate (as described in previous sections). More formally, a pin is a list of 3D points (“pin points,” called ColumnPin in neuPrint) that start at the top of a neuropil ROI and end at the bottom.

Starting from a neuron to hexagonal coordinate assignment in the medulla (15 cell types, as described in section **Assigning neurons to medulla hexagonal coordinates**), we construct pin *α* corresponding to hexagonal coordinates *α* following 8 steps:

1. For all neurons assigned to the hexagonal coordinate *α*, get the 3D positions *r*_*i*_ of their (pre and post) synapses in the medulla, where *i* runs from 1 to the number of synapses.
2. Find the axes of biggest variation of *r*_*i*_ using principal component analysis (PCA). Fix the sign of the axis of biggest variation (PC 1) such that PC 1 points from the top to the bottom of the medulla ROI.
3. Compute the Euclidean distances *l*_*i*_ of *r*_*i*_ from their mean ∑_*i*_ *r*_*i*_ in the PC 2-PC 3 plane, and drop the *r*_*i*_ for which *l*_*i*_ < *f*_*lat*_*σ*_*lat*_, where 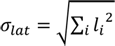 is the root mean squared distance of all *r*_*i*_. The factor *f*_*lat*_*=2* was chosen manually such that the *r*_*i*_ with *l*_*i*_ ≥ *f*_*lat*_*σ*_*lat*_ looked like they were part of a different column.
4. For the remaining *r*_*i*_, define their normalized projection onto PC 1 as *t*_*i*_ such that *t*_*i*_ = −1 corresponds to the top and *t*_*i*_ = 1 to the bottom of the medulla ROI. To clarify, this PC 1 was computed in step 2, using all *r*_*i*_.
5. For those *r*_*i*_ with *t*_*i*_ < 0.1, we compute a new PC 1 and find its intersection with the top of the medulla ROI–we denote this 3D point as *r*_*top*_. If there are less than *N*_*PC*_ points with *t*_*i*_ < 0.1 then we take the points with the *N*_*PC*_ smallest *t*_*i*_’s instead to compute the new PC 1. We chose the integer *N*_*PC*_ = 800 manually to ensure a robust estimation of *r*_*top*_. An analogous computation, using the points with *t*_*i*_ > 0.2 or the points with the *N*_*PC*_ largest *t*_*i*_’s and the intersection of their newly computed PC 1 with the bottom of the medulla ROI, defines *r*_*bottom*_.
6. Define a list *L*′_*α*_ for which the first element is *r*_*top*_, the last element is *r*_*bottom*_ and intermediate elements are determined by rank ordering the *r*_*i*_ by *t*_*i*_. Define another list *L*_*α*_ for which the first element is *r*_*top*_, the last element is *r*_*bottom*_ and the intermediate elements are the averages of the elements in *L*′_*α*_ with their *N*_*avg*_ neighbors. The integer *N*_*avg*_ = 260 was determined manually such that the elements in *L*_*α*_ formed approximately a smooth line. This line was often helical and typically did not go through the center of the *r*_*i*_.
7. To create a line that goes through the center of the *r*_*i*_, find the shortest sublist *O*_*α*_ of *L*_*α*_ for which the first element is *r*_*top*_, the last element is *r*_*bottom*_ and the 3D Euclidean distance between neighboring points is at most *d*_*max*_. We defined *d*_*max*_ as the 3D Euclidean distance between *r*_*top*_ and *r*_*bottom*_ divided by *N*_*tang*_. We chose the integer *N*_*tang*_ = 7 manually such that *O*_*α*_ corresponds to a line that goes approximately through the center of the *r*_*i*_. Notice that the minimal length of *O*_*α*_ is *N*_*tang*_ + 1.
8. For each of the 3 spatial coordinates of points in *O*_*α*_, use a spline interpolation (we used a Piecewise Cubic Hermite Interpolating Polynomial, PCHIP) to uniformly sample *O*_*α*_ with *N*_*samp*_ points. This list of length *N*_*samp*_ defines pin *α*. We determined *N*_*samp*_ = 121 manually by checking if the synapse distributions along a pin looked approximately smooth and did not change much with small changes in *N*_*samp*_.

To create pins in the lobula and lobula plate, we first need a neuron to hexagonal coordinate assignment. In the lobula, this assignment was simply inherited from the medulla by choosing neurons that innervate both neuropils (namely, Tm1, Tm2, Tm4, Tm9, Tm20). For the lobula plate, we used T4 neurons because they innervate both the medulla and the lobula plate. We assigned T4 neurons to hexagonal coordinates by assigning T4 neurons of each subtype to Mi1 neurons (from section **Assigning T4s to Mi1s to extend the coordinate system to the lobula plate**) and then using the Mi1’s hexagonal coordinate assignment.

The steps to construct pins in the lobula and lobula plate were similar to the 8 steps to construct pins in the medulla but there were also important differences:

A) Instead of two separate PCA’s for the top and bottom of each pin (step 5 above), we performed one PCA using all the *r*_*i*_ from step 2. With the exception of the top of the lobula, we defined *r*_*top*_ and *r*_*bottom*_ as the intersection of this newly computed PC 1 with the top and bottom of the neuropil ROI, respectively. For the top of the lobula, *r*_*top*_ was computed differently because the top of the lobula ROI contains ridges from the chiasm, which we did not want to replicate in our pins. (However, they are necessarily replicated in our column ROIs as they subdivide a neuropil ROI). Instead, we defined *r*_*top*_ as the extrapolation of the newly computed PC 1 to the median position of the points with the 37 smallest *t*_*i*_’s (from step 4). Defined this way, the *r*_*top*_ of different lobula columns also lie on a slightly curved surface rather than on ridges.
B) In the lobula plate, PCA (in steps 2 and A) was not performed on the collection of all synapses of neurons but instead on the collection of spatially averaged synapses of neurons. To be clear, in the lobula plate, the number of points on which PCA was performed equaled the number of neurons (3 or 4). The reason is that the lobula plate ROI is not as thick at the medulla and lobula, such that the spatial variation of all synapses along the column (from top to bottom) can be comparable to that in the lateral directions. Using synapses would result in a PC 1 tilted away from the columnar direction (between the top and bottom of the neuropil).
C) We chose different empirical parameters: in the lobula, we used *f*_*lat*_ = 1.5, *N*_*avg*_ = 37, *N*_*tang*_ = 2, *N*_*samp*_ = 76, and in the lobula plate, *f*_*lat*_ = 1, *N*_*avg*_ = 56, *N*_*tang*_ = 2, *N*_*samp*_ = 51.
D) In the lobula and lobula plate, we imposed 3 criteria to make a pin (which would have been automatically satisfied in the medulla):

i. The number of *r*_*i*_ after step 3 is at least *N*_*avg*_.
ii. The 3D Euclidean distance between *r*_*top*_ and the spatial average of the *r*_*i*_ with the 5% largest *t*_*i*_’s is at most *d*_*max*_ (as defined in step 7).
iii. The 3D Euclidean distance between *r*_*bottom*_ and the spatial average of the *r*_*i*_ with the 5% smallest *t*_*i*_’s is at most *d*_*max*_. Criterion (i) is required to perform step 7. Criteria (ii-iii) were imposed because otherwise, the synapse positions *r*_*i*_ (after step 3) could be far from both the top and bottom neuropil ROI, e.g. for the lobula if the cell type Tm20 was missing. In this case, the pin would often be tilted. We hypothesize that a more complete neuron-to-hexagonal coordinate assignment in the lobula and lobula plate could obviate the need for these criteria.

Based on these parameter *N*_*samp*_, we produce medulla pins with 121 points along their centerline, lobula plate pins with 76 points and lobula plate pins with 51 points. This discretization is used for measuring the position of a synapse along each pin, which we refer to as its *depth*. But because of the criteria in D, only 870 pins were created in the lobula and 783 pins in the lobula plate. These pins were less regular than those in the medulla. Therefore, for the lobula and lobula plate, we iteratively refined the pins by regularizing existing pins using neighbors and filling in missing pins. The iterative algorithm replaces pins *α*_*old*_ with pins *α*_*new*_ as follows:

- Regularization: For every created pin *α*_*old*_, find the created pins *β*_*old*_ with hexagonal coordinates *β*_*old*_ that differ from *α*_*old*_ by at most 2. For every *β*_*old*_ and for every 3D point in the list of the pin *β*_*old*_, subtract the first 3D point in the list of the pin *β*_*old*_. The median of these lists across *β*_*old*_ plus the first 3D point in the list of the pin *α*_*old*_ defines the new pin *α*_*new*_.
- Filling-in: For every missing pin *α*_*old*_, find (i) the 4 created pins *β*_*old*_ with hexagonal coordinates *β*_*old*_ that differ from *α*_*old*_ by ±1 along either of the two hexagonal coordinates (but not both), or, if (i) does not exist, (ii) the 2 created pins *β*_*old*_ where *β*_*old*_ differs from *α*_*old*_ by ±1 along one hexagonal coordinate. If neither (i) nor (ii) exists, then there is no filling-in. Otherwise, the new pin *α*_*new*_ is defined as the median of the pins *β*_*old*_ across *β*_*old*_.

The iteration is stopped if no more pins are filled in. This took 2 iterations in the lobula, resulting in 875 pins, and 3 iterations in the lobula plate, giving 817 pins.

Given all the pins of a neuropil ROI, any 3D point within the neuropil ROI can be assigned a pin by minimal Euclidean distance, i.e., it is assigned to pin *α* if the minimal Euclidean distance to pin *α* is smallest among all pins. The minimal Euclidean distance of a point to a pin is defined as the minimum Euclidean distance of the point to any pin point of that pin. We used this to show that the curved centerlines are a more accurate model of the shape of the neurons in the medulla, and therefore of the columns, as compared to a straight line (quantified in Extended Data Fig. 5a, where presynapses and postsynapses are assigned to columns through this method). The pinpoints are stored as ColumnPins in neuPrint, which can be used for efficient assignments of synapses to columns and corresponding depth, for details see **Connectome data access overview**.

The column ROIs shown in Fig. 3d were created by uniformly sampling a box surrounding the corresponding neuropil ROI at a resolution of 512nm. Then, all sampled 3D points within the neuropil ROI were assigned a pin as described. All 3D points assigned to the same pin constitute a column ROI. The column ROIs were upsampled if an assignment at a finer resolution needed to be made. Assignments at different resolutions produce slightly different results. These column ROIs can facilitate quantification as in Fig. 3e and Extended Data Fig. 6, and also visualization (e.g. Extended Data Fig. 5c,d).

### Layer ROIs (related to Fig. 3)

Since a pin is an ordered list of 3D points going from the top to the bottom of a neuropil ROI in a fixed number of steps, we can view the order of the list as an indicator of layer depth. Specifically, for any 3D point in a neuropil ROI, we find the order of the closest 3D pin point (in terms of Euclidean distance); we define “depth” as the order normalized to a number between 0 and 1 such that 0 correspond to the first points in pins and 1 to the last points. By assigning a depth to a synapse, we compute 1D synapse distributions. For the distributions in Fig. 3c and Extended Data Fig. 6a, the synapse distributions were subsampled by a factor of 2. For the 1D synapse distribution of a cell type, we take all the synapses of all its neurons in each of the 3 neuropils where pins are defined.

Layer ROIs subdivide a neuropil ROI and are based on depth thresholds that define layer boundaries. Each depth threshold was chosen as the cutoff of a peak in the presynapse or postsynapse distribution of one cell type (except for the lobula plate, see later in this paragraph). The choice of cell type, presynapses or postsynapses, which peak, and if we chose the upper or lower cutoff of a peak, was inspired by LM images selected to match prior notions of these layers. In Fig. 3c we show the correspondence between EM and LM and the selected criteria for defining layers. In the lobula plate, each depth threshold was defined as the mean of two T4 type depth thresholds. We used the mean of the thresholds of two types since there are gaps in depth between the different T4 types (and so defining the top of e.g. layer 3 separately from the bottom of layer 2 is not useful), as can be clearly seen in the LM image (Fig. 3c, left).

The peak cutoffs on 1D synapse distributions were determined by the following steps, which depend on one parameter *frac_peaks*:

1. Find the threshold *syn_thre* on the 1D synapse distribution such that the fraction of the distribution above *syn_thre* is *frac_peaks*.
2. The lowest depth value at which the 1D synapse distribution surpasses *syn_thre* is defined as the lower cutoff of the first peak. The next depth value that is at least 3 depth points away from the lower cutoff, at which the 1D synapse distribution falls below *syn_thre* is defined as the upper cutoff of the first peak. Lower and upper cutoffs for other peaks, at larger depths, are defined similarly. Each cutoff must be at least 3 depth points away from other cutoffs.

We chose *frac_peaks* for each neuropil roughly based on the peakedness of the 1D synapse distributions:

*frac_peaks=0.85* in the medulla, *frac_peaks=0.8* in the lobula, and *frac_peaks=0.75* in the lobula plate.

To make layer ROIs, we could apply the same depth threshold to all pins, however, if done directly, we found that layers exhibit “blockiness” owing to the discretization of the pins. We therefore applied a smoothing procedure. For each layer, we defined smooth layer meshes as alpha shapes, which depend on two parameters α and *frac_ext*. They were constructed as follows:

1. Find the pin points at the lower and upper layer boundaries.
2. Find the upper and lower edge pin points from the upper and lower pin points, respectively.
3. Find the upper and lower edge mesh points by taking the neuropil ROI mesh points and taking the points that are closest to the upper and lower edge pin points, respectively.
4. Displace the upper and lower edge mesh points further away from the corresponding upper and lower edge pin points by adding *frac_ext* times their difference.
5. Use the upper and lower pin points and the displaced upper and lower edge mesh points to define an alpha shape with parameter α.

For the medulla we used α=0.0006 and *frac_ext=1,* and for the lobula and lobula plate, α=0.0004 and *frac_ext=0.5*. Note that smaller α corresponds to more smoothing and smaller *frac_ext* to smaller layers in the lateral dimensions. The resulting smooth layer meshes could be slightly overlapping and/or missing small volumes of the neuropil ROI. These smooth layer meshes were only used to construct layer ROIs.

Similar to the column ROIs, the layer ROIs shown in Fig. 3d were created by uniformly sampling a box surrounding the corresponding neuropil ROI at a resolution of 512nm. Then, all sampled, 3D points within the neuropil ROI were assigned a layer with a 2-step procedure. First each 3D point in a neuropil ROI is assigned to a putative layer by depth, resulting in “blocky” layers. Second, the assignment is finalized by looping through the smooth layer meshes (from top to bottom) and assigning a layer if the point was contained within that smooth layer mesh. The putative assignment was necessary to account for the small volumes in the neuropil ROI that were not covered by any of the smooth meshes. Volumes covered by two smooth layer meshes are handled by assigning it to the later layer. All 3D points assigned to the same layer constitute a layer ROI. The layer ROIs were then implemented in neuPrint, which is how we use them to acquire synapses within each “layer” in Fig. 4g, Fig. 9a, Fig. 5b, and the Cell Type Explorer web resource.

### Training and evaluation data for transmitter predictions (related to Fig. 4)

Neurotransmitter assignments used as “ground truth” for training and evaluation of transmitter predictions (see below) came from either the literature or new experiments and are summarized in Supplementary Table 5. We primarily used data from the literature^36,40,46,47,50–55^ for training data and performed new experiments to generate an expanded set of evaluation data. Both published and new experiments use the expression of molecular markers, detected at the mRNA or, in a few cases, protein level, as indicators of transmitter. New data were generated using Fluorescence in situ hybridizations (FISH). We used two adaptations of this method, referred to as FISH^56,57^ and EASI-FISH^58^. Members of the Janelia Project Technical Resources and FLyLight Project teams performed the EASI-FISH and FISH experiments, respectively, following published protocols. We used Fiji to assess the results of these experiments qualitatively and to select sections for display. We routinely adjusted the brightness and contrast of the image stacks for both evaluation and display. Cell types of interest were identified by using split-GAL4 lines to specifically label these cells. In some cases, multiple cell types were examined using the same driver line; this was possible when soma locations of these cell types were clearly different or if the cells of a group of neurons with overlapping cell body locations showed signals with the same FISH probes. In most FISH and EASI-FISH experiments, we only probed for markers for cholinergic, glutamatergic, and GABAergic transmission (ChaT/VAChT, VGlut, and GAD1 probes) and only followed this with probes for OA, 5-HT, and Dop markers in a few cases when we suspected an aminergic phenotype, or if the results with the first probe set showed no clear signals in cells of interest. We acknowledge that this approach may miss some cases of co-transmission but made many experiments far more efficient. It is also partly justified by published distributions of FISH markers for aminergic transmission^57^ which show only a few cells with clear labeling in or near the optic lobes or, in the case of dopamine, include many small cell bodies in the medulla cell body rind but suggest (through co-labeling with a split-GAL4 driver) that these belong primarily, if not exclusively, to Mi15 neurons^57^. Supplementary Table 5 lists neurotransmitter data for 88 cell types, not part of the training dataset. The 79 evaluated as “validation with new experimental data” in Fig. 4e, are based on this set, excluding the uncertain types ([Cm11], [Pm2]), those with co-transmission or “unclear” (Mi15, CL357, l-LNv, MeVC27, OLVC4, T1), and assigning Histamine to HBeyelet, R8p, R8y and counting the latter two as one type.

### Neurotransmitter prediction (related to Fig. 4)

We predicted neurotransmitter identity for 7,014,581 presynapses in the optic-lobe dataset, following an existing method^59^. We first trained an image classifier network on EM volumes (640 x 640 x 640 nm^3^) to predict each presynaptic site as one of seven possible neurotransmitter types (and their common abbreviations: acetylcholine/ACh, glutamate/Glu, GABA, histamine/His, dopamine/Dop, octopamine/OA, and serotonin/5HT). The ground truth neurotransmitter types are described in the section **Training and evaluation data for transmitter predictions** and were based on prior experimental data, detailed in Supplementary Table 5. Entire neurons (for ACh, Glu, GABA, and His) or individual synapses (due to the low number of neurons for Dop, OA, 5HT) were partitioned into disjoint training, validation, and testing sets of 70%, 10%, and 20% respectively, optimizing for similar class frequency in each partition. Models included an additional class to indicate non-synaptic or unrecognized structures, trained by sampling random locations in the bounding box of all synaptic locations.

In neuPrint, the computed probability of each neurotransmitter type is attached to the presynapses, and can be queried using the properties: *ntAcetylcholineProb*, *ntGlutamateProb*, *ntGabaProb*, *ntHistamineProb*, *ntDopamineProb*, *ntOctopamineProb*, and *ntSerotoninProb*. The values range from 0 (impossible) to 1.0 (certain), sum to 1, and should be interpreted as relative probabilities. The accuracy of the synapse-level neurotransmitter predictions is assessed in Fig. 4b. While the training data were restricted to high confidence (*≥*0.9) synapses, for evaluating the performance of the trained network we produced a confusion matrix (Fig. 4b) showing classification results on the testing dataset and an additional 1,014,151 synapses with a detection confidence < 0.9.

In addition to these synapse level predictions, we provide aggregated neurotransmitter predictions (with quality controls applied, as explained below) at the neuron and cell type level, which are stored in neuPrint as properties *predictedNt* and *celltypePredictedNt*, respectively. First, we computed the neuron-level neurotransmitter confidence score as previously described^59^ and assigned the most frequent presynaptic neurotransmitter as the neuron’s prediction if that cell has *≥*50 presynapses and confidence *≥* 0.5. Similarly, we computed the cell type-level confidence score and assigned the most frequent presynaptic neurotransmitter as each cell type’s prediction if the type has *≥*100 presynapses (summing over all neurons of the type) and confidence *≥* 0.5. Individual neurons and cell types that did not meet these criteria were labeled with “unclear” neurotransmitters. Finally, we provide a consensus neurotransmitter type (in neuPrint, property *consensusNt*). Cell types with predictions for Dop, OA, 5HT, that were not supported by the high-confidence experimental data listed in Supplementary Table 5 had their consensus type set to “unclear” as these neurotransmitters were underrepresented in the training data and independent measurements of the abundance of these transmitters did not match their prevalence in the predictions. The consensus neurotransmitters are reported in the Cell Type Explorer web resource, the Cell Type Catalog (Supplemental Fig. 1), and the basis for the summary data in Fig. 4.

### Number of innervated columns of a cell type (related to Fig. 5, Supplementary Fig. 1)

The column pins we have built for the medulla, lobula, and lobula plate provide a framework for quantifying the size of visual system cells, by measuring the number of columns a neuron has synapses in as a function of depth. We do this two different ways, a direct method, which is used in Fig. 5b and in the Cell Type Catalog (Supplementary Fig. 1), takes advantage of the pin points stored in neuPrint to assign synapses to depths along each pin, and this method is described in **Summary of connectivity and size by depth**. We also wanted to summarize the size of neurons in each neuropil with a single number, but simply summing the column assignments of all synapses across depth led to overestimates of cell size that did not match our expectations. We therefore implemented a trimming method, explained below, that was used as the basis of the size and coverage analysis in Fig. 5d-f, Extended Data Fig. 9, and the Cell Type Explorer web resource.

The trimming procedure was implemented to solve a problem we frequently encountered: columnar neurons like L1 or Mi1 in the medulla have a clear “home” column (the column with the largest synapse count summed over depth), but a few synapses are often sprinkled away from their home column. As a result, the direct column count, when summed over depth, would typically be larger than what we intuitively expect (e.g. columnar neurons should be of size ∼1 in units of columns). To correct for this overestimation, we trim these excessive synapses using the following steps:

1. For each neuron, we rank the columns by how many synapses they contain (summed over depth), with the main column having rank 1 (columnar cells have a clear home column, but many cell types do not). The cumulative fraction of the number of synapses as a function of rank has a convex shape that eventually flattens out around 1, and the curve is similar for most neurons of the same cell type.
2. We then take the median of the cumulative fraction curve across neurons and use a knee/elbow finder to identify the rank (*rank\**) at which the curve defined by the median cumulative fraction versus rank (with the point (0,0) added) had the highest curvature. We used the knee/elbow finder algorithm “kneedle”^60^, for which we specified that the curve is concave and increasing, and we used the default parameter S=1.
3. If the median cumulative fraction at *rank\** is less than 0.775, we take *rank\** to equal the rank where the median cumulative fraction is at least 0.995 (or, if this value is not reached, its maximum).
4. Then, for every neuron separately, we discard all synapses in columns with synapse count less than that in the column of rank *rank\**. If all columns have unique synapse counts, this is the same as discarding all synapses in columns of rank larger than *rank\**.

After the trimming step, we proceed to count columns and then average these distributions across neurons of the same cell type.

### Spatial coverage of visual regions: cell size, coverage factor, completeness (related to Fig. 5)

Spatial coverage metrics were calculated per neuron type for each of the major optic lobe regions (i.e. the medulla, lobula, and lobula plate). Metrics were calculated separately for neurons from the same neuron type depending on the hemispheric location of their somas by querying the neuPrint database for their instance (e.g. “aMe12_L” or “aMe12_R”). Individual synapses were assigned to columnar hexagonal coordinates (see **Defining column centerlines and ROIs)** by using their assignment in the neuPrint database. When we summarize the size of the cell in column units or conversely the number of cells per column we use the trimming procedure above. In all other cases, we use the direct column occupancy data.

The spatial patterns are represented on the hexagonal “eye map” introduced in Fig. 3, as heatmaps of the number of cells and number of synapses per column. The raw, untrimmed column occupancy data were used to summarize the number of synapses per column whereas the trimmed data was used when plotting the number of cells per column (Fig. 5d and the Cell Type Explorer web resource). The number of synapses per column reflects the sum of both “pre” and “post” synapses in that column. Whereas, when calculating the number of synapses across all neuron types within a given optic lobe region (Fig. 3e) only “post” synapses were included to avoid double counting “pre” and “post” synapses at the same connection. The maximum color scale value was set as the 98th percentile value of the highest value across the medulla, lobula and lobula plate for both plots (Cell Type Explorer web resource).

We quantify cell size (Fig. 5e, Extended Data Fig. 9, Cell Type Explorer web resource) as the median number of columns innervated by neurons of the type within a designated optic lobe region, after synapses had been trimmed, per neuron, based on the trimming procedure described in the section above.

To examine the amount of spatial overlap between individual neurons of the same type we calculated the coverage factor (Fig. 5e-f, Extended Data Fig. 9, Cell Type Explorer web resource) for each optic lobe region. This metric is calculated as the mean number of neurons that contribute synapses to a single column, across all occupied columns of the specified optic lobe region, after trimming using the method described above.

To assess the proportion of total columns innervated by a neuron type, we calculated the “columnar completeness” per region for each type to describe the proportion of total columns within each optic lobe region that are occupied by synapses from all neurons of the type (for this calculation, we use raw column occupancy data without trimming). This metric is provided as a count of innervated columns in Fig. 5f and Extended Data Fig. 9, and as the proportion of total columns (0-1) in the Cell Type Explorer web resource.

The extent to which neuron types densely or sparsely innervated a particular optic lobe region was investigated by comparing the total number of columns innervated by neurons of that type, related to the “columnar completeness” described above, and the columnar area covered by all neurons of the type. For most cell types the “columnar area” was found by fitting a convex hull around the hexagonal coordinates based on their assignment in neuPrint, and calculating its area. For the cell types where the convex hull was a poor approximation of the area covered, we instead counted the innervated columns; this included aMe2, aMe10, Ascending_TBD_1, Cm-DRA, Dm-DRA1, Dm-DRA2, MeLo11, MeTu4f, MeVP15, MeVP20, MeVPMe8, Mi16, R7d, R8d, TmY19b, LC14b in Medulla, aMeVP_TBD_1, LC14a-1, LC14a-1, LC14a-2, LC14a-2, LC14b, LC31a, Li37, LoVC17, LoVP12, LoVP26, LoVP49, LoVP92, LPT31, LT80, LT81, TmY19b in Lobula, and LPT31, LC14b and LPT100 in the Lobula Plate. The columnar area metric (Fig. 5f, Extended Data Fig. 9, Cell Type Explorer web resource) enabled us to quantitatively distinguish cell types that densely cover fractional parts of a region from types that sparsely cover the entire region.

### Summary of connectivity and size by depth (related to Fig. 5, Supplementary Fig. 1)

To produce the synapse distribution by depth, featured prominently in the Cell Type Catalog (Supplementary Fig. 1), we first identify all synapses of each target cell from the designated cell type. Using a k-dimensional tree, we find the nearest neighboring column pin by Euclidean distance for each presynapse and postsynapse. The depth property of the column pin associates each synapse with one of the 121 medulla, 76 lobula, or 51 lobula plate depth bins (see **Defining column centerlines and ROIs).** Across all neurons of a cell type, we calculate the mean count separately for presynapses and postsynapses, per depth. We smooth the distribution per brain region with a Savitzky-Golay filter (window size 5, 1st-order polynomials) before plotting on the left-side panel of the summary data figures. Separately we provide the percentage of synapses located within the AME over the cell type’s total number of synapses, again separated by presynapses and postsynapses.

To quantify each cell type’s size as a function of depth, we find the depth for all synapses. Since columns are implemented as ROIs in neuPrint (see **Connectome data access overview**), each synapse comes with an associated column. For each cell type, we calculate the mean count of columns with synapses (combining pre and post) per depth and smooth the distribution per brain region with a Savitzky-Golay filter (window size 5, 1st-order polynomials) before plotting on the right-most panel of the summary data.

The center panel of the cell type summary contains a connectivity summary. For all neurons of a target cell instance (effectively type, but separately accounting for right and left instances), we find all synapses and their connecting partner cells. We order them by the fraction of input and output connections and show the top five after removing unnamed segments.

### Morphology clustering (related to Fig. 5 and Extended Data Fig. 7)

To test if the 68 medulla OLIN cell types (11,102 neurons), mentioned in the section **Connectivity-based cell clustering** (Extended Data Fig. 2), could be split into cell types by their morphology alone, we assigned each neuron a feature vector of length 244. The features capture the pre/post synapse and pre/post size distributions across depths of the medulla, similar to the first and third columns in Fig. 5b but for individual neurons instead of cell types (and for the medulla only). Additionally, the number of innervated columns is split into presynapse and postsynapse innervations and is calculated directly from the pins (introduced in the section **Defining column centerlines and ROIs** rather than the column ROIs stored in neuPrint). We also subsampled the depth by a factor of 2 (as described in **Layer ROIs**). We used a confidence of 0.9 for the included synapses, based on the expectation that higher confidence synapses would produce more robust feature vectors, but did not systematically explore this.

Concretely, the first 61 features are the number of presynapses across depths of the medulla, normalized to the number of innervated columns of presynapses (with no trimming). This normalization means that we multiply the number of presynapses across depths of medulla with one and the same factor *N*_*pre*,*size*_/*N*_*pre*,*syn*_ where *N*_*pre*,*syn*_ equals the sum of the number of synapses across depths in medulla, and *N*_*pre*,*syn*_ equals the sum of number of innervated columns across depths in medulla. While this normalization is not biologically motivated, it follows the common strategy of bringing different features to the same scale in order to compare them. The second 61 features are similar to the first 61 but for post instead of presynapses. The third 61 features are the untrimmed number of columns innervated by presynapses (without further normalization). The final 61 features are the number of columns innervated by postsynapses. Example feature vectors for three neurons are shown in Extended Data Fig. 7a.

For the clustering, we used the same standard library described in the **Connectivity-based cell clustering** section but used Euclidean distance as the metric for comparing feature vectors. We again make a flat cut in the hierarchical clustering diagram to obtain 80 clusters. The corresponding confusion matrix and the completeness and homogeneity scores^61^ are shown in Extended Data Fig. 7b.

### Confusion matrices of cell type clustering

To facilitate comparison of the connectivity and morphology clustering, we present the results as identically formatted “confusion” matrices in Extended Data Fig. 2a and Extended Data Fig. 7b. The numbers in the confusion matrices are the nonzero numbers of neurons for a given cell type in each cluster.

The rows (cell type names) are ordered lexicographically. The columns (cluster identities) are ordered by the number of neurons in a cell type as follows. Starting with the first cell type, we pick the first cluster as the one with the largest number of neurons of that cell type; the second cluster is the cluster with the second largest number; we continue picking clusters until the number of neurons in each remaining cluster is 5% or less for that cell type. Then we continue picking from the remaining clusters using the second cell type with an analogous procedure as for the first cell type. The colors in the confusion matrices correspond to five non-overlapping categories:

1. red: 1-to-1 clusters: At least 80% of neurons of one cell type are in one cluster and at least 80% of neurons in that cluster are of that cell type.
2. blue: many-to-1 clusters: At least 80% of neurons of one cell type are in one cluster but less than 80% of neurons in that cluster are of that cell type.
3. green: 1-to-many clusters: Less than 80% but at least 10% of neurons of one cell type are in a cluster but at least 80% of neurons in those clusters are of that cell type.
4. yellow: mixed clusters: Less than 80% but at least 10% of neurons of one cell type are in a cluster and less than 80% but at least 10% of neurons in those clusters are of that cell type.
5. gray: outliers: everything else that is nonzero.

These categories are also described as probabilities in Extended Data Fig. 2a and Extended Data Fig. 7b.

### Cell Type Explorer web resource analysis (related to Extended Data Fig. 8)

We provide a set of interactive, interlinked webpages to facilitate quick browsing of connections between all the cell types in our inventory. Each cell type is detailed on a single webpage, except for the cases of bilateral neurons with both left and right instances in the right optic lobe. These cell types are described on two pages, one per instance. On each cell type’s webpage, the mean presynapses and postsynapses for many ROIs are provided. The synapses in the medulla, lobula, and lobula plate layers were determined by querying synapses within the specified layer ROIs (described in section **Layer ROIs**) for each neuron in the optic lobe. The lamina and accessory medulla do not have layers, and so the synapses were queried for the whole ROI. In the central brain, queries were conducted on all primary ROIs but excluding the left and right optic lobes. These synapses are then presented as the mean values in across neurons of a given type in each of these ROIs.

The coverage factor, columnar and area completeness, and cell size (in columns) were quantified for each of the medulla, lobula, and lobula plate, as described in section **Spatial coverage of visual regions: cell size, coverage factor, completeness.**

The connectivity table displayed on the webpages was generated by fetching all connecting neurons either upstream (inputs, T-bars) or downstream (outputs, postsynaptic densities) from the cell type named on the webpage. It lists the total number of connections between the input/output cell type and the titular cell type (“total connections”), the total connections divided by the number of the titular cell type’s cell count (“connections/[cell type]”), and provides both the percentage and cumulative percentage of the total connection count. The table displays all connections between neurons with a mean count > 1.0. Most cells have a long tail of weak connections that are not displayed on the webpages. All connections, including the omitted weaker connections, can be found in neuPrint.

The Cell Type Explorer is available at https://reiserlab.github.io/male-drosophila-visual-system-connectome/ and for download as a set of static html pages at DOI:10.5281/zenodo.10891950.

### Gallery plots (related to Supplementary Fig. 1, and renderings of selected neurons throughout)

For the gallery plots in Supplementary Fig 1, we used the optic lobe “slice views” detailed in the section **Pipeline for rendering neurons.** We established three camera orientations in Blender^25^ that produced three slices oriented equatorially, dorsally, and ventrally, labeled in panels as E, D, and V, respectively. Over multiple rounds of manual curation, we selected representative neurons that captured many features of each cell type (when there are more than 1 to choose from). We choose cells with a primary orientation close to one of the three slices. An additional consideration was to select cells that, when projected along these camera views, had cell bodies that could be visualized outside the brain regions, wherever possible (cell bodies are generally outside of the brain regions neuropils; this is simply an artifact of projecting 3D bodies onto 2D images). The width of the slice used for the projection of the neuron varied with the morphology of the individual cell. For cells that required a thicker slice, the layer pattern of the projection was affected as the layers are curved--an effect most prominent for neurons rendered on the D and V slices. Therefore, while every effort was taken to show neurons with characteristic layer innervation patterns, the accompanying summary of synapses by depth should be used as the characteristic layer innervation pattern of the cell type. For a small number of cells with morphology not captured by the three standard slices, we used a combination of custom slices to generate projections on the dorsal, equatorial, or ventral slice views that faithfully conveyed the morphology of the cells. For example, we used this strategy to capture the vertical cell body process for the Cell Type Catalog of the Mi20 cell type. We refer to the set of selected neurons for each cell type as the “star neurons,” and the parameters used for rendering the projections of the neurons, including the slice used and slice thickness (width), are listed in the “params/all_stars.xlsx” file in the code repository (also see **Connectome data access overview**).

### Split-GAL4 lines expressing in visual system neurons

Supplementary Table 6 summarizes our collection of split-GAL4 lines for visual system neurons. The majority of driver lines are newly reported here, while we also include lines from our previous publications and another (Dionne et al, in prep.) that is in preparation. Further details on the construction and annotation of these lines are described below. Supplementary Table 6 includes annotations of the main optic lobe cell type(s) that each lines drive expression in. Images used for characterizing the lines are available at https://splitgal4.janelia.org/cgi-bin/splitgal4.cgi. Split-GAL4 lines that are currently maintained as fly stocks (the majority) can be requested via the same website and many are also available from the Bloomington stock center. Most of the remaining lines could be remade using publicly available AD and DBD hemidrivers.

Newly reported split-GAL4 driver lines were generated as previously described^20,40,62^. Briefly, we selected candidate hemidriver (AD and DBD) combinations based on images of GAL4 line expression patterns (overall expression patterns^63,64^ or MCFO-labeled individual cells^32,65^ and tested whether these combinations could drive the expression of a UAS reporter specifically in cells of interest. Promising AD/DBD combinations were combined into stable fly stocks and the expression patterns retested. For detailed characterization, we examined images of overall expression patterns in the brain and VNC at low resolution (20x objective) and, for most lines, additional high resolution (63x objective) images of cell populations, MCFO-labeled individual cells or both. Most split-Gal4 screening, sample preparation and imaging were performed by the FlyLight Project Team, using established methods^20,32,62,66^ and publicly available protocols (https://www.janelia.org/project-team/flylight/protocols). The FlyLight split-GAL4 project has been summarized in a recent publication^67^.

Images shown in Fig. 8 and Extended Data Fig. 12 were displayed using VVD viewer^68^. Details of the underlying genotypes (e.g. reporters used) and images (e.g. objective used) for each panel are included in Supplementary Table 6 (column “Figure details”) with the following abbreviations: MCFO-1 and MCFO-7^32^, 20xChr (for 20XUAS-CsChrimson-mVenus in *attP18*^69^ and syt-HA (for pJFRC51-3XUAS-IVS-Syt::smHA in *su(Hw)attP1*) and myr-smFLAG (for pJFRC225-5XUAS-IVS-myr::smFLAG in *VK00005*)^32,62^. The latter two transgenes were used a combined stock but most images only show the myr-smFLAG reporter (as indicated in the table). For overlays of EM skeletons and LM images, both were registered to a template brain (see section **LM-EM volume correspondence**; JRC2018U)^70^ and overlayed in VVD viewer. In most cases, images show projections through the full brain in an anterior view or through a subvolume selected and oriented to display the layer pattern within the optic lobe. Some LM images are of left optic lobes, these were mirrored to match the location of the EM skeletons in the right optic lobe.

Assignment of LM images to EM cell types was done by visual comparison of LM and EM morphologies, at multiple levels of detail. The range of anatomical features used for LM/EM matching is illustrated in Fig. 8 and Extended Data Fig. 12. In Supplementary Table 6, matched cell types are listed with the names used in Cell Type Catalog (Supplementary Fig. 1) and throughout this manuscript. We cannot provide confident 1- for-1 matches between optic lobe cells expressed in every driver line and EM-defined cell types. Some LM/EM matches are to groups of cell types rather than to single types, some represent more tentative, candidate matches. This is usually the case for cell types that are primarily distinguished by EM-connectivity and/or by relying on features that are difficult to assess with the available LM images. Such cases are identified in Supplementary Table 6 as follows: Cell type names in square brackets denote expression in one or more of a group of cell types, and type names in parentheses indicate lower confidence matches within a group of annotations. For example, “LC31a, (LC31b)” means that we are confident a driver is expressed in LC31a and probably also LC31b. “[Pm2a,(Pm2b)]” describes a driver with expression in Pm2a or Pm2b, or both, but for which we consider expression in Pm2a more likely than expression in Pm2b. In a few cases, primarily for brevity, we list groups of cell types under a single name, e.g. “[Tm]” for expression in as yet unidentified Tm cells or “T4” for expression in T4a, T4b, T4c and T4d.

The completeness of the EM cell type inventory greatly facilitated the LM/EM matching efforts. A potential limitation to this completeness are sexually dimorphic neurons, but we find very little evidence for them. Most images used for annotation of the split-Gal4 collection are of female flies whereas this EM reconstruction is of a male optic lobe. Yet nearly every visual system cell types in the LM images of most lines could be readily matched to the EM-defined cell types, with one exception mentioned in the main text.

The cell type annotations in Supplementary Table 6 are intended to facilitate finding driver lines that may be useful for experiments on specific optic lobe neurons of interest. Because driver lines with expression in some “off-target” cell types can still be valuable for many experiments, we included some lines with additional expression in the collection. We note that our annotations generally do not include these off-target cell types in the central brain, VNC and, in some cases, other (minor) visual system expression patterns. The images used for annotating the lines are available online (see above and **Data Availability**) and could be consulted if details of any additional expression may prove critical for a planned experiment.

### Matching cell types between the male optic lobe and FlyWire datasets (related to Extended Data Fig. 13 and Supplementary Table 7)

To facilitate comparison between the male optic-lobe (OL) and female FlyWire-FAFB datasets^71–74^, we matched cell types across datasets. This matching was performed after the completion of cell typing in the OL dataset. The methods used for matching are summarized below. A full listing of the cell type matching is provided as Supplementary Table 7, and summarized in Extended Data Fig. 13a. We note that this is a nearly complete, yet preliminary, matching of the cell types. We expect that upcoming data on central brain and left optic lobe connectivity in the male brain, along with advances in neuron annotations in FlyWire, will contribute to further validation and refinement of the cell type matches between the datasets.

Supplementary Table 7 contains the cell type name given in the optic lobe dataset, and the name(s) for the corresponding cell types in FlyWire. FlyWire cell types are based on Schlegel *et al*.^72^ and Matsliah *et al.*^73^ and in some cases two different type name annotations, one from each reference, are provided. We note that additional annotations for some FlyWire cells beyond those included in these cell type references (for example, the subdivisions of MeTu neurons in Garner *et al.*^45^) are not included in our comparison.

For most cell types, we matched cells between the datasets as one-to-one matches, however, ∼13% of types are matched as many-to-one, one-to-many, or many-to-many. “1-to-many” matches are cases in which a single optic lobe cell type is split into multiple types in FlyWire; “many-to-1” indicates a correspondence between multiple optic lobe types and a single Flywire type. Example of a “1-to-1” match and a “2-to-1” match are shown in Extended Data Fig. 13b. For OLIN and OLCN cell types for which there are both Schlegel *et al*.^72^ and Matsliah *et al.*^73^ types, the classification of the matches refers to the match between the OL and Matsliah *et al.*^73^ types. For 6 VPN cell types, the matches are classified as “many-to-many”, these cell types could be matched as groups in each dataset but we were not confident about the matches between individual types within these groups. We also identified several cell types, classified as “unmatched”, that appear to be present in one dataset but absent from the other. Some of these types are candidates for male- or female-specific neurons, but others likely represent biological variability between the two flies or technical variability due to incomplete reconstruction of some cells (examples in Extended Data Fig. 13c).

Supplementary Table 7 also includes cell type counts for each dataset (with counts for the two sets of FlyWire type annotations listed separately). For most cell types, the FlyWire dataset with the Schlegel *et al*.^72^ annotations cover the cells on both the right and left brain hemispheres, and in these cases, we provide the counts from both sides. The table also provides neurotransmitter predictions from both datasets. For the OL cell types, the listed predictions are the consensus predictions (from neuPrint; provided in Supplementary Table 1 and described in section **Neurotransmitter prediction**). For FlyWire, the table lists the most frequently predicted neurotransmitter^59^ across the neurons of a given type (union of the Schlegel *et al*.^72^ and Matsliah *et al.*^73^ annotations). FlyWire types where the top two predicted neurotransmitters are tied are labeled as “unclear”. Since some hemibrain types were renamed in our study (to provide more systematic names to optic lobe neurons), we include hemibrain matches in Supplementary Table 7 based on the hemibrain-FlyWire matches reported in Schlegel *et al*.^72^.

To match cell types, we first used NBLAST to identify morphologically similar neurons between the FlyWire and the OL datasets following the procedure described in Schlegel *et al*.^72^ for hemibrain-FlyWire matching. For FlyWire, we used previously prepared skeletons^71^ for all neurons; for OL, we downloaded skeletons from neuPrint (release optic-lobe:v1.0). Skeletons were pruned to remove twigs smaller than 5 microns and then transformed from OL into FlyWire (FAFB14.1) space using a combination of non-rigid transforms. Once in FlyWire space, they were resampled to 0.5 nodes per micron of cable to approximately match the resolution of the FlyWire L2 skeletons and then turned into dotprops. The NBLAST was then run both in forward (FlyWire→OL) and reverse (OL→FlyWire) direction and the minimum between both was used.

From the NBLAST scores we extracted, for each FlyWire neuron, a list of potential OL matching neurons based on the 5 top hits. A sample of individual FlyWire neurons for each type was co-visualized with their potential OL hits in neuroglancer and the corresponding cell type match was recorded. Although the registration of the optic lobes of the two datasets is imperfect in places, this approach was sufficient to identify candidate matches at the cell type level. All neurons for matched cell types were then co-visualized to confirm the match, verifying that the total number of neurons was also consistent between datasets. While this approach was sufficient to match most cell types across datasets, in a few cases, we also used the output of connectivity co-clustering (with coconatfly or cocoa, see below) to identify and confirm matches. The main use cases were morphologically similar OLIN and OLCN types, mostly among those that had only been typed unilaterally by Matsliah *et al.*^73^ and were not present in the Schlegel *et al*.^72^ typing.

Neuroglancer scenes were prepared with co-aligned OL and FlyWire meshes. The static FlyWire meshes from materialization 783^71^ were transformed to OL space (using the transformation to the male brain described in section **LM-EM volume correspondence** for the final step). Cell type annotations for OL neurons were sourced from neuPrint (release optic-lobe:v1.0). FlyWire cell type annotations were sourced from https://github.com/flyconnectome/flywire_annotations^72^. For intrinsic and connecting neurons (OLIN/OLCN), we also matched to the cell types described in Matsliah *et al.*^73^ as downloaded on April 24, 2024 from https://codex.flywire.ai/api/download?data_product=visual_neuron_types&data_version=783. For convenience, a neuroglancer annotation layer was prepared to display Schlegel *et al*.^72^ and Matsliah *et al.*^73^ cell types. At the time of writing, Matsliah *et al.*^73^ described a larger number of OLIN/OLCN cell types than Schlegel *et al*.^72^ but only provided them for one optic lobe in FlyWire. These scenes are provided with a unique URL, per OL cell type, in Supplementary Table 7. For the OLIN and OLCN cell types with both Schlegel *et al*.^72^ and Matsliah *et al.*^73^ annotations, the linked neuroglancer scenes include both FlyWire types; subsets of the displayed cells can be selected through regular expression searches of the scenes.

Skeleton and mesh processing, spatial transforms, and NBLAST were performed using the navis^28^ and fafbseg-py python packages (see https://github.com/navis-org/navis). Connectivity co-clustering was performed using coconatfly (https://natverse.org/coconatfly) and cocoa (https://github.com/flyconnectome/cocoa). Code and neuroglancer links are available at https://github.com/flyconnectome/ol_annotations. Connectivity data for FlyWire neurons can be obtained from the FlyWire CAVE annotation backend, for example, through https://fafbseg-py.readthedocs.io/en/latest.

The initial set of matches obtained as described above was independently reviewed by a separate team member. Similar to the initial matching, this review primarily relied on comparisons of morphology via visual inspection of the neuroglancer scenes. For a minority of types, mainly in the OLIN and OLCN groups, that are difficult to distinguish by morphology alone, we also reviewed connectivity patterns. This was done either by comparison of aggregate cell type to cell type connectivity or connectivity clustering or co-clustering (using adaptations of the approach described in the methods section **Connectivity-based cell clustering**). Clustering was consulted in some cases as it provides a quick overview of cell-level matches and includes untyped cells. Overall, this review only resulted in very minor adjustments (e.g. identifying untyped candidate matches for some initially unmatched types, indicated in the “notes” column of Supplementary Table 7) of the initial matches.

The comparison of predicted neurotransmitters for the matched cell types provides yet another independent evaluation of both the putative matches and the neurotransmitter predictions (in two independent EM volumes, with independent training data, but using the same method of Eckstein *et al.*^59^). In general, the concordance is very high. In the OL dataset we used a conservative policy for reporting consensus transmitter predictions, therefore 168 of the 727 cell types have “unclear” predictions (130 of which are VPNs). Of the remaining OL cell types, there are 9 with “unclear” predictions in FlyWire, and 9 with no cross-dataset matches, so no neurotransmitter predictions that could be compared. Once these cell types are excluded (without double counting), there are 543 cell types whose predictions can be evaluated, of which 502 agree (92.5%). Of the 41 mismatched predictions, 8 are OL predictions for histamine, which was not included in the FlyWire training or predictions. Of the remaining 33 mismatched predictions, 26 are OL predictions for glutamate and FW predictions for GABA (interestingly, there are no mismatched predictions for the opposite assignments).

### Brain region connectivity (related to Fig. 9)

This analysis aims to quantify the amount of connectivity among brain regions, with a weighted sum of their connected neurons. The brain regions considered in Fig. 9 are the visual system layer ROIs (and the AME) and prominent grouped central brain regions.

If one neuron receives input connections from brain region A and sends output connections to brain region C, then how much does it contribute to the A→C connectivity? We first count the output connections in C, and then compute the fraction of input connections that are in A. We consider the product of the count and the fraction as the contribution of this neuron to the A→C connectivity. For example, a neuron X has 10 upstream connections in region A and 20 in region B, and has 30 downstream connections in region C and 15 in region D. Then X’s contributions to inter-region connectivity are 10 for A→C, 20 for B→C, 5 for A→D, and 10 for B→D.

We use the visual system neuron groups (defined in **Cell type groups**), and analyzed the inter-region connectivity of all OLINs, OLCNs, and VPNs separately. Each is represented by one connectivity matrix. For the OLIN and OLCN analysis, we examine the connectivity between layer regions: medulla layers {1,2,3,4,5,6,7,8,9,10}, lobula layers {1,2,3,4,5A,5B,6}, and Lobula Plate layers {1,2,3,4}. For the VPNs, we analyzed connectivity between these 21 optic lobe regions and central brain regions. Several brain regions have been grouped into super regions to simplify this summary^17^. The regions are abbreviated in Fig. 9a, and many are illustrated in Fig. 9b (projection view): Anterior Optic Tubercle (AOTU), Anterior Ventrolateral Protocerebrum (AVLP), Posterior Lateral Protocerebrum (PLP), Posterior Ventrolateral Protocerebrum (PVLP), Wedge (WED), Periesophageal Neuropils (PENP), Inferior Neuropils (INP), Lateral Complex (LX), Mushroom Body (MB), Central Complex (CX), Ventromedial Neuropils (VMNP), Gnathal Ganglia (GNG), Lateral Horn (LH), and Superior Neuropils (SNP).

## Data availability

The connectome data is directly accessible via the neuPrint database server: https://neuprint.janelia.org/?dataset=optic-lobe:v1.0

The Cell Type Explorer web resource is available at: https://reiserlab.github.io/male-drosophila-visual-system-connectome/index.html and can also be downloaded as a zip file from DOI:10.5281/zenodo.10891950

Meshes of neurons and boundaries of ROIs are provided as Google Cloud Storage buckets at gs://flyem-optic-lobe/v1.0/segmentation and gs://flyem-optic-lobe/rois/, respectively.

The LM-EM transformation vectors from our EM sample to the JRC2018M template brain are at: https://figshare.com/s/e17528e5e2c44ba78b5d, also stored at: <u>gs://flyem-optic-lobe/transforms/MaleCNS_JRC2018M.h5</u>. In that file, the "dfield" transformation vectors map points from EM space to LM template space, and "invdfield" vectors map points in the opposite direction.

Images of split-GAL4 driver lines are available at https://splitgal4.janelia.org/cgi-bin/splitgal4.cgi.

## Fly stock availability

Fly stock availability is detailed in Supplementary Table 6. Most split-GAL4 driver lines are available at https://splitgal4.janelia.org/cgi-bin/splitgal4.cgi. In addition, many lines have been deposited in the Bloomington *Drosophila* stock center. For additional lines that are currently not maintained as combined stocks, most component AD and DBD hemidrivers are available from the Bloomington stock center.

## Code availability

The python source code to replicate our analysis and visualizations follows established best practices for computational analysis^75^. In the version-controlled code repository (https://github.com/reiserlab/male-drosophila-visual-system-connectome-code), we document dependencies such as Pandas^76^, NumPy^77^, SciPy^48^, Jupyter^78^, Plotly^79^, snakemake^31^, and Trimesh^80^.

In the code repository’s “docs” folder, we share a range of technology-specific “getting started” guides. This includes a guide for setting up the required python-3.10 (or newer) and our software on Linux, Windows, or Mac computers, running simple analysis, and using the workflow management system snakemake to replicate our figures.

## Notes

https://neuprint.janelia.org/?dataset=optic-lobe:v1.0

https://reiserlab.github.io/male-drosophila-visual-system-connectome/index.html

https://github.com/reiserlab/male-drosophila-visual-system-connectome-code

https://splitgal4.janelia.org/cgi-bin/splitgal4.cgi

### Competing Interest Statement

The authors have declared no competing interest.

### Summary of Updates

The major addition is Extended Data Figure 13 and Supplementary Table 7, covering the matching of cell types to from the male Optic Lobe to FlyWire-FAFB dataset. Two authors and several references were added. Minor additional corrections to the manuscript

