## Supplementary Fig 1 for "Connectome-driven neural inventory of a complete visual system"

**Supplementary Fig. 1: Summary of the anatomy and connectivity of visual system neurons**

The 68-page summary of all the neurons follows the conventions of Fig. 5 (see Methods section **Summary of connectivity and size by depth**). The quantified morphology and distribution of pre- and post-synapses (mean of all cells of the type), together with the top five connected cells, are found on odd pages. For details of cell type names, see the methods section **Cell type nomenclature**. Some bilateral neurons are found in the right optic lobe, and to distinguish between the right and left hemisphere versions of the cell type, we treat them separately and append an (R) or (L) to the cell type's name. In the top 5 connectivity data in the center panel, left-hemisphere cell types are indicated with a magenta label. The synapse distributions are plotted as counts of synapses in each bin along columns of the brain region, with the scale indicated on the right-hand side. Medulla columns: 121 bins (0.54µm mean length), lobula columns: 76 bins (0.75 µm mean length), lobula plate columns: 51 bins (0.5µm mean length). The length of columns varies by spatial position (see Extended Data Fig. 6b). Annotated example of per-cell type summary:

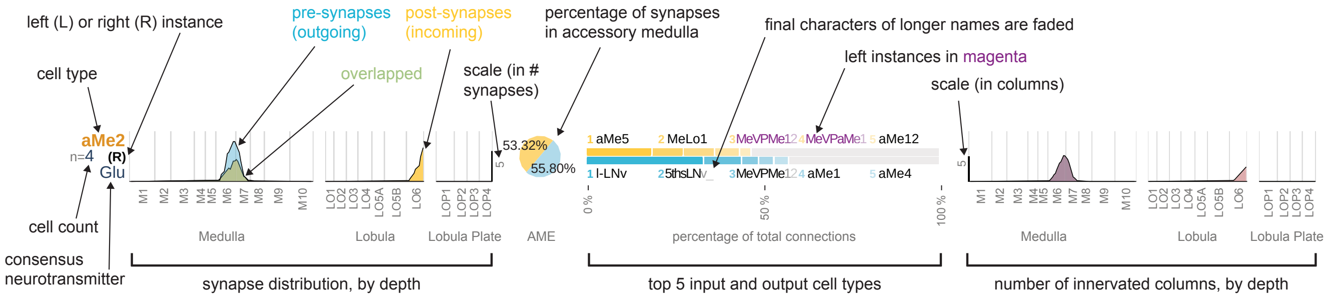


A gallery of rendered example neurons (see Methods section **Gallery of representative neurons**) is paired with the summary data and shown on even pages. A representative neuron of each type has been selected and is shown in a sliced view to reveal the innervation patterns of the visual regions (scale bar = 50µm); the central brain arbors of most VPN and VCN neurons are not shown in their entirety. Each slice is taken from one of three locations, indicated by D (dorsal), E (equatorial), or V (ventral), with most neurons shown in the E slice, except for cells best represented in more dorsal or ventral locations. The layers are sheared relative to the slicing planes in the D and V locations, so the layer patterns should be viewed as suggestive, but the more accurate description is found in the corresponding synapse and size (by depth) data plots.

The PDF document contains bookmarks with cell type names (linked to the summary data). The document is most conveniently viewed in "two-page view" and tested to work well with Adobe Acrobat.

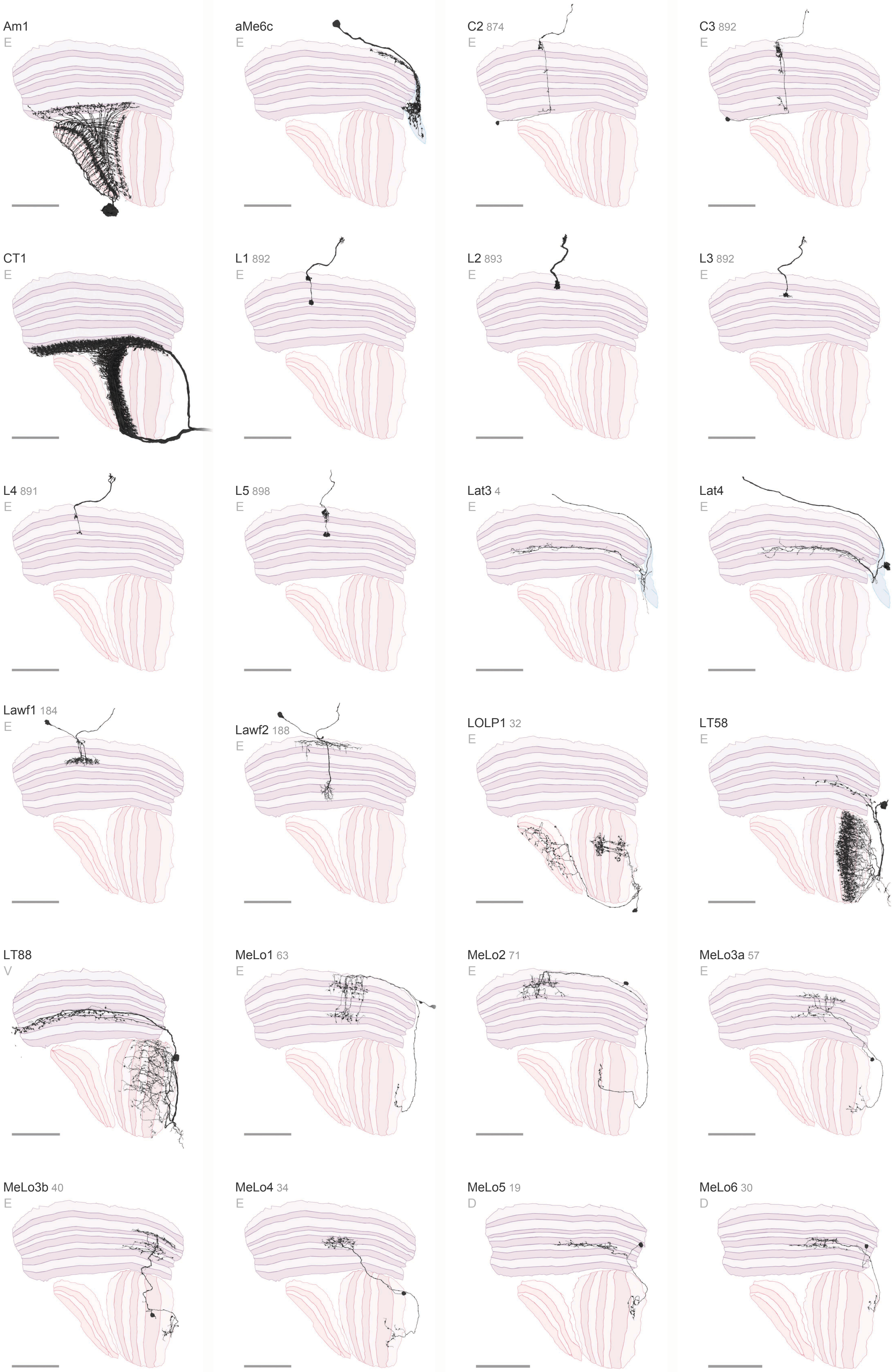

Am1
E
aMe6c
E
C2 874
E
C3 892
E
CT1
E
L1 892
E
L2 893
E
L3 892
E
L4 891
E
L5 898
E
Lat3 4
E
Lat4
E
Lawf1 184
E
Lawf2 188
E
LOLP1 32
E
LT58
E
LT88
V
MeLo1 63
E
MeLo2 71
E
MeLo3a 57
E
MeLo3b 40
E
MeLo4 34
E
MeLo5 19
D
MeLo6 30
D

### Optic Lobe Connecting Neurons 1 / 4

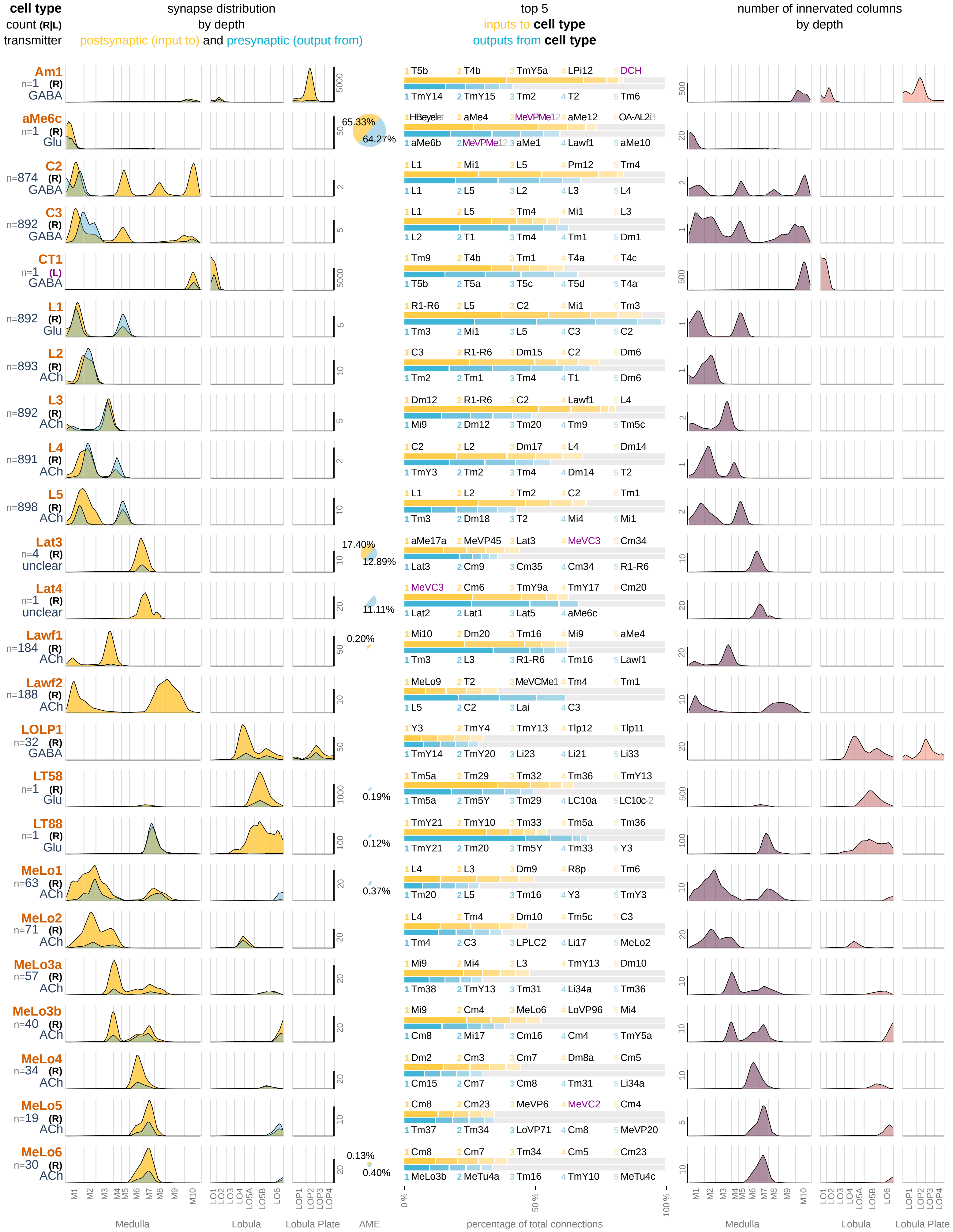

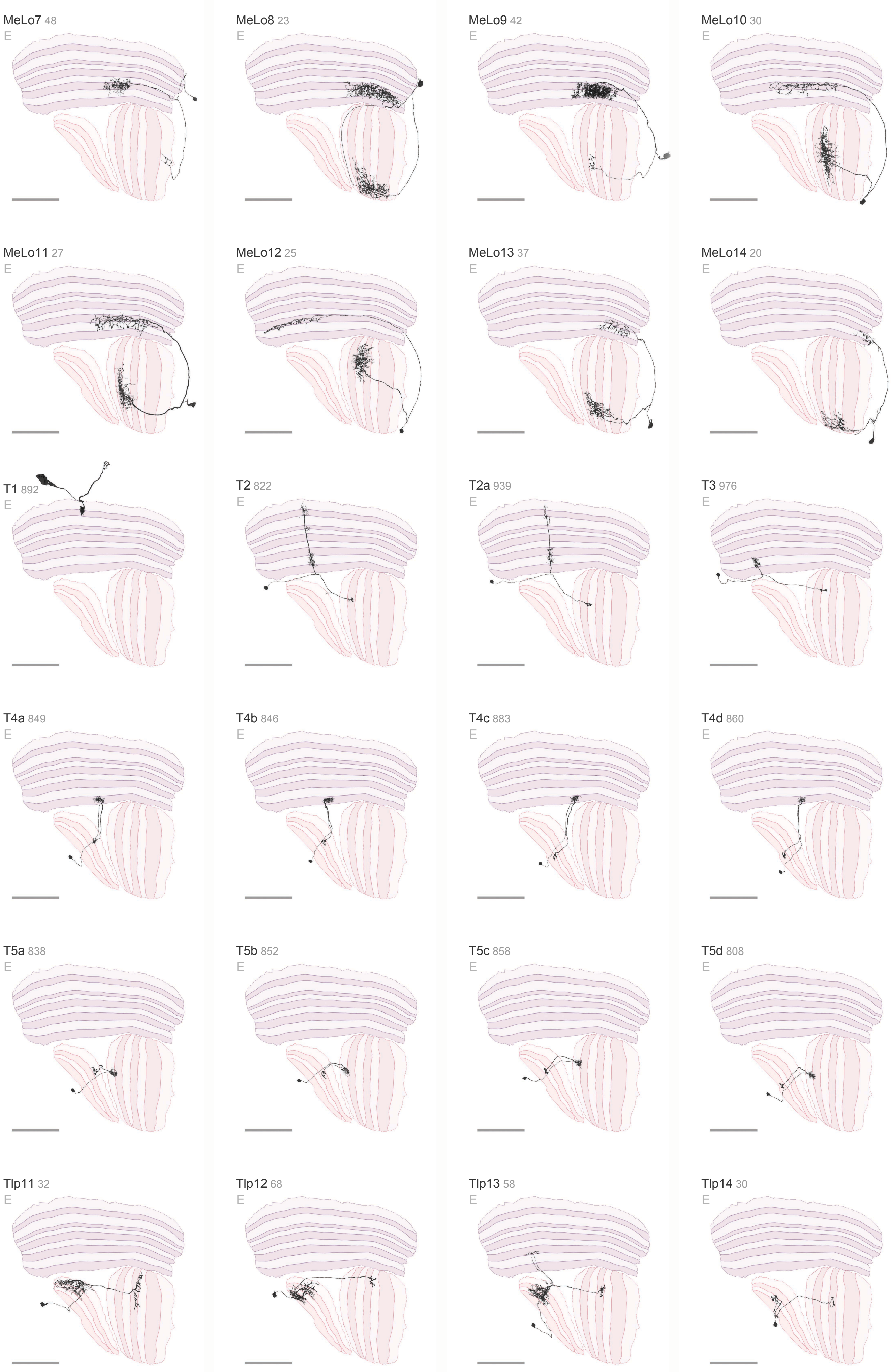

MeLo7 48
E
MeLo8 23
E
MeLo9 42
E
MeLo10 30
E
MeLo11 27
E
MeLo12 25
E
MeLo13 37
E
MeLo14 20
E
T1 892
E
T2 822
E
T2a 939
E
T3 976
E
T4a 849
E
T4b 846
E
T4c 883
E
T4d 860
E
T5a 838
E
T5b 852
E
T5c 858
E
T5d 808
E
Tlp11 32
E
Tlp12 68
E
Tlp13 58
E
Tlp14 30
E

### Optic Lobe Connecting Neurons 2 / 4

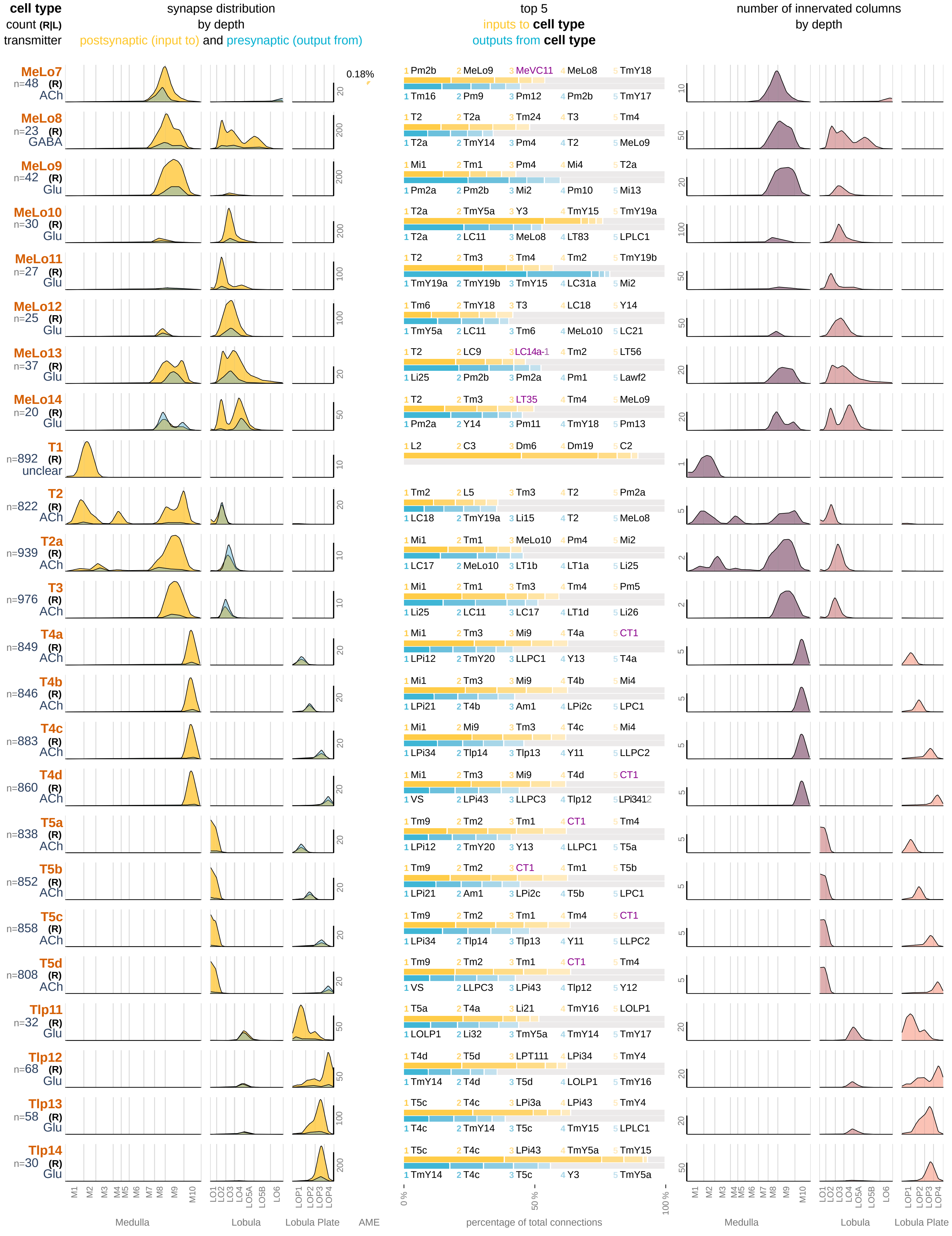

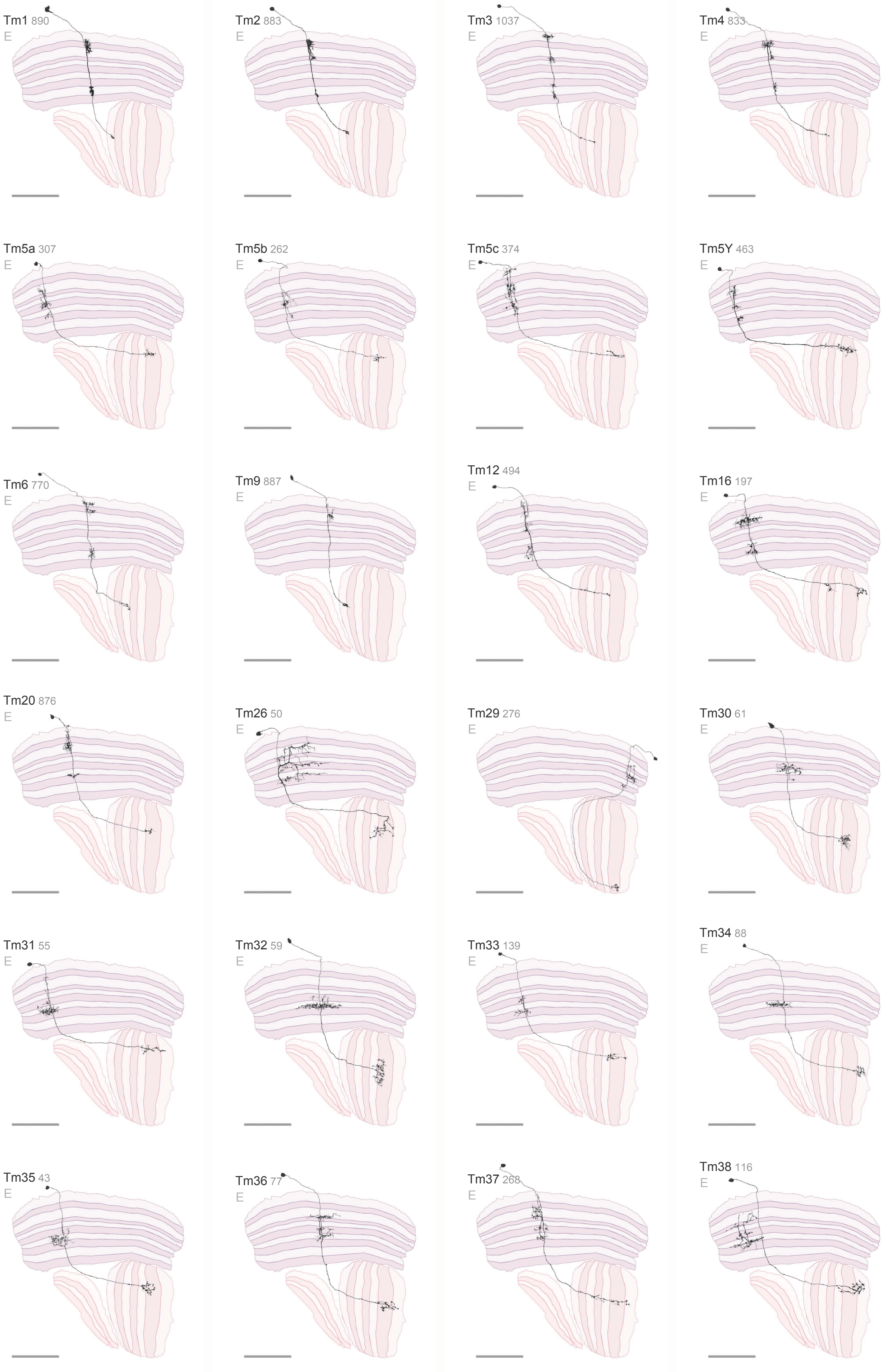

Tm1 890
E
Tm2 883
E
Tm3 1037
E
Tm4 833
E
Tm5a 307
E
Tm5b 262
E
Tm5c 374
E
Tm5Y 463
E
Tm6 770
E
Tm9 887
E
Tm12 494
E
Tm16 197
E
Tm20 876
E
Tm26 50
E
Tm29 276
E
Tm30 61
E
Tm31 55
E
Tm32 59
E
Tm33 139
E
Tm34 88
E
Tm35 43
E
Tm36 77
E
Tm37 268
E
Tm38 116
E

### Optic Lobe Connecting Neurons 3 / 4

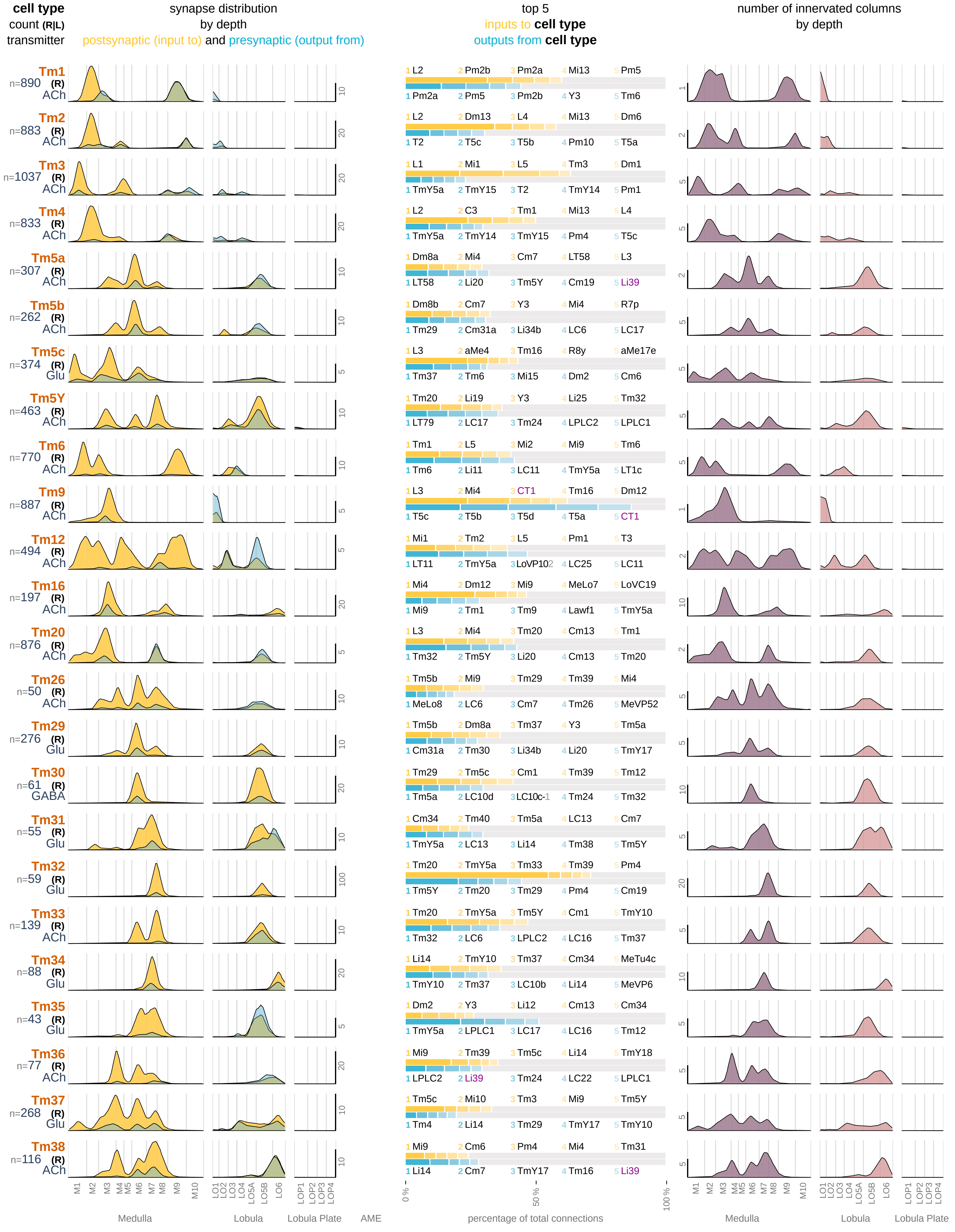

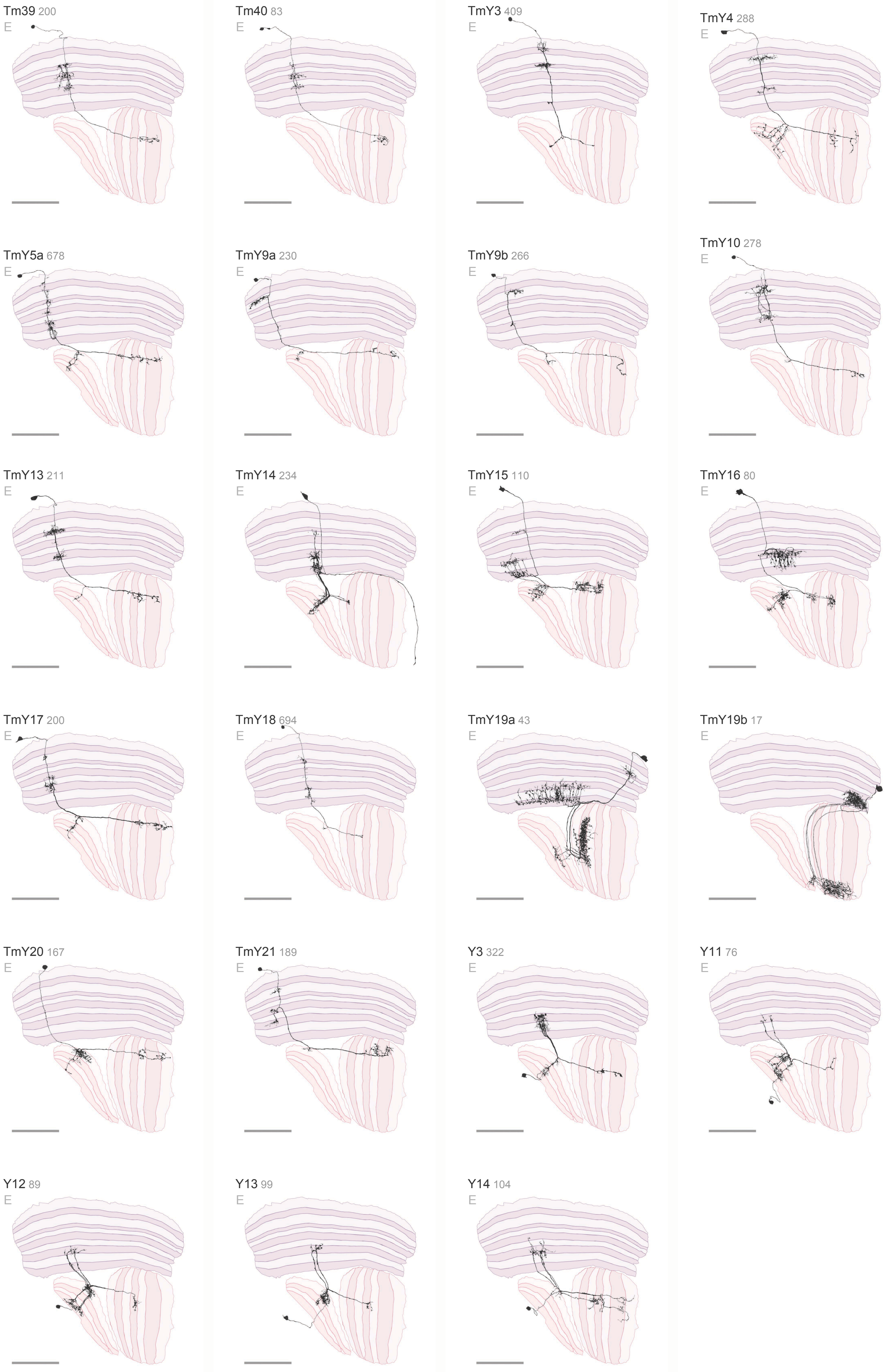

Tm39 200
E
Tm40 83
E
TmY3 409
E
TmY4 288
E
TmY5a 678
E
TmY9a 230
E
TmY9b 266
E
TmY10 278
E
TmY13 211
E
TmY14 234
E
TmY15 110
E
TmY16 80
E
TmY17 200
E
TmY18 694
E
TmY19a 43
E
TmY19b 17
E
TmY20 167
E
TmY21 189
E
Y3 322
E
Y11 76
E
Y12 89
E
Y13 99
E
Y14 104
E

### Optic Lobe Connecting Neurons 4 / 4

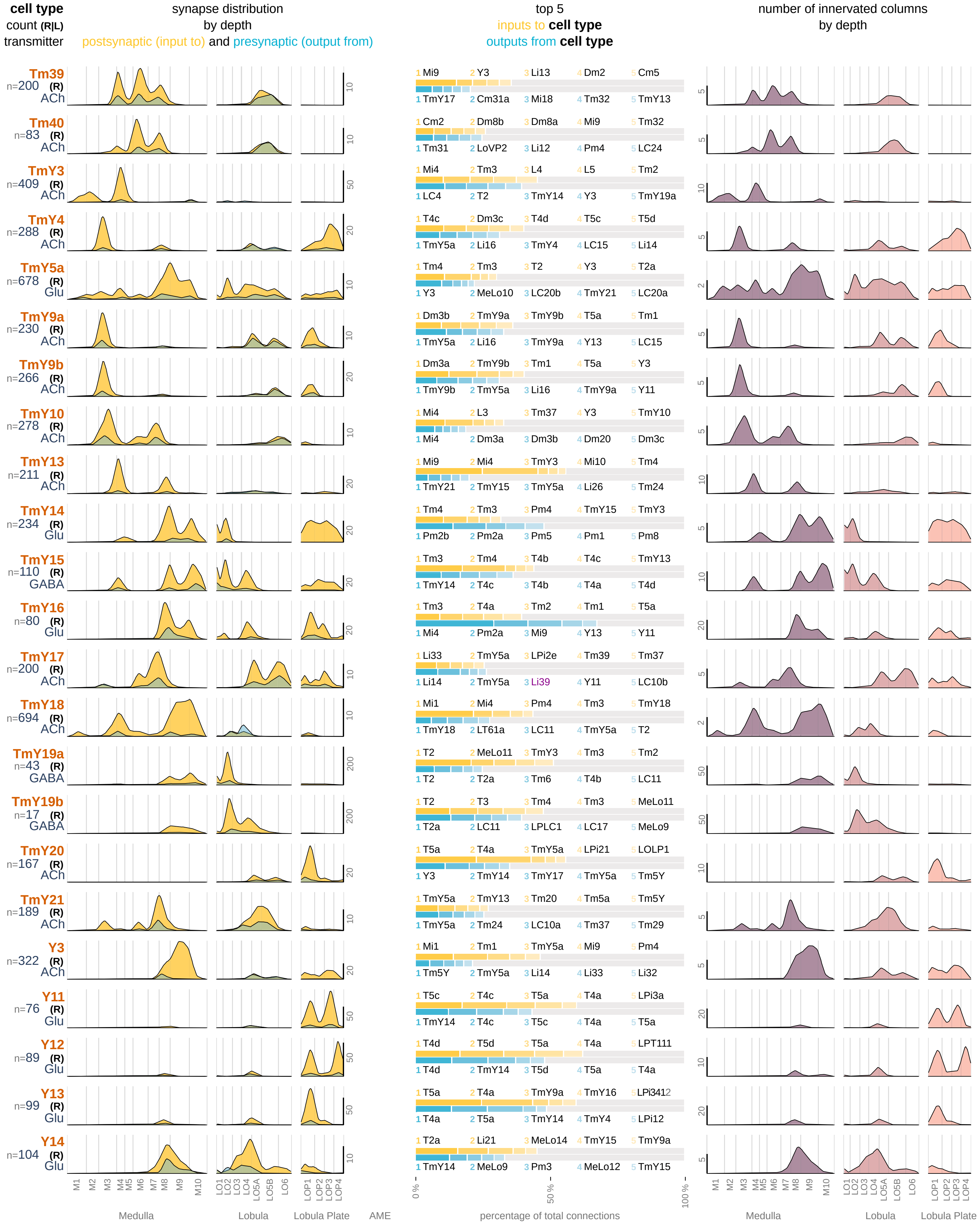

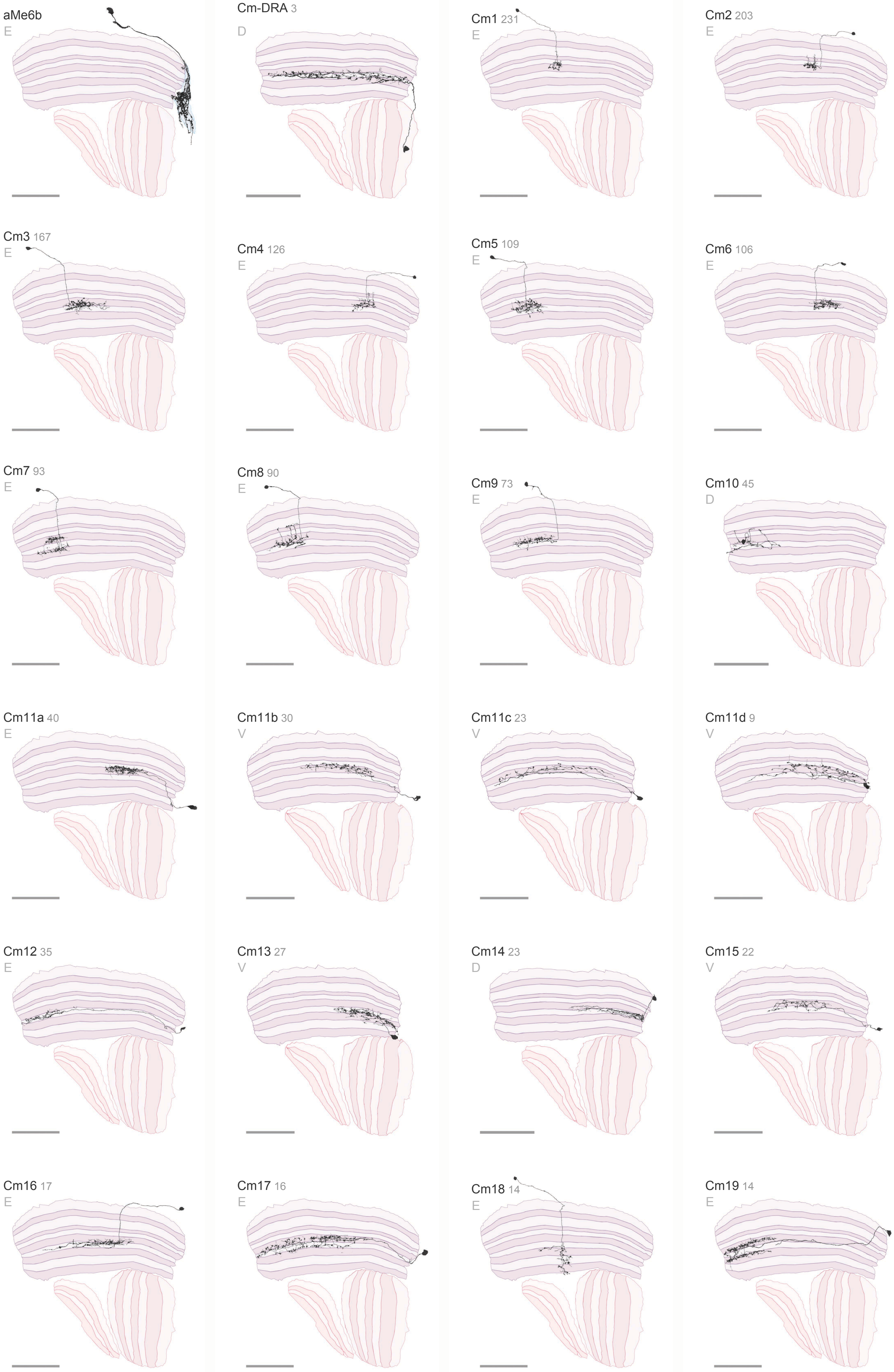
aMe6b
E
Cm-DRA 3
D
Cm1 231
E
Cm2 203
E
Cm3 167
E
Cm4 126
E
Cm5 109
E
Cm6 106
E
Cm7 93
E
Cm8 90
E
Cm9 73
E
Cm10 45
D
Cm11a 40
E
Cm11b 30
V
Cm11c 23
V
Cm11d 9
V
Cm12 35
E
Cm13 27
V
Cm14 23
D
Cm15 22
V
Cm16 17
E
Cm17 16
E
Cm18 14
E
Cm19 14
E

### Optic Lobe Intrinsic Neurons 1 / 7

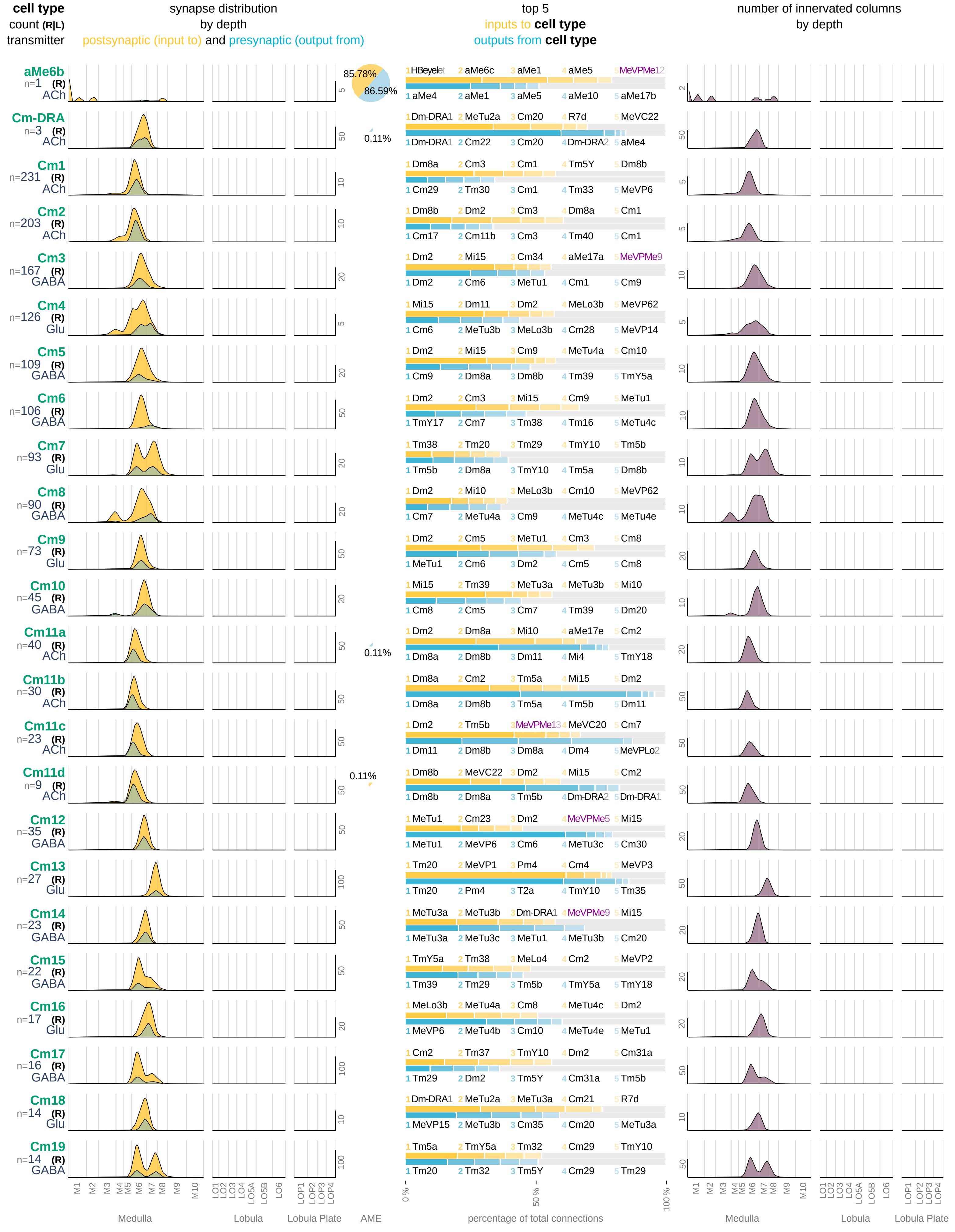

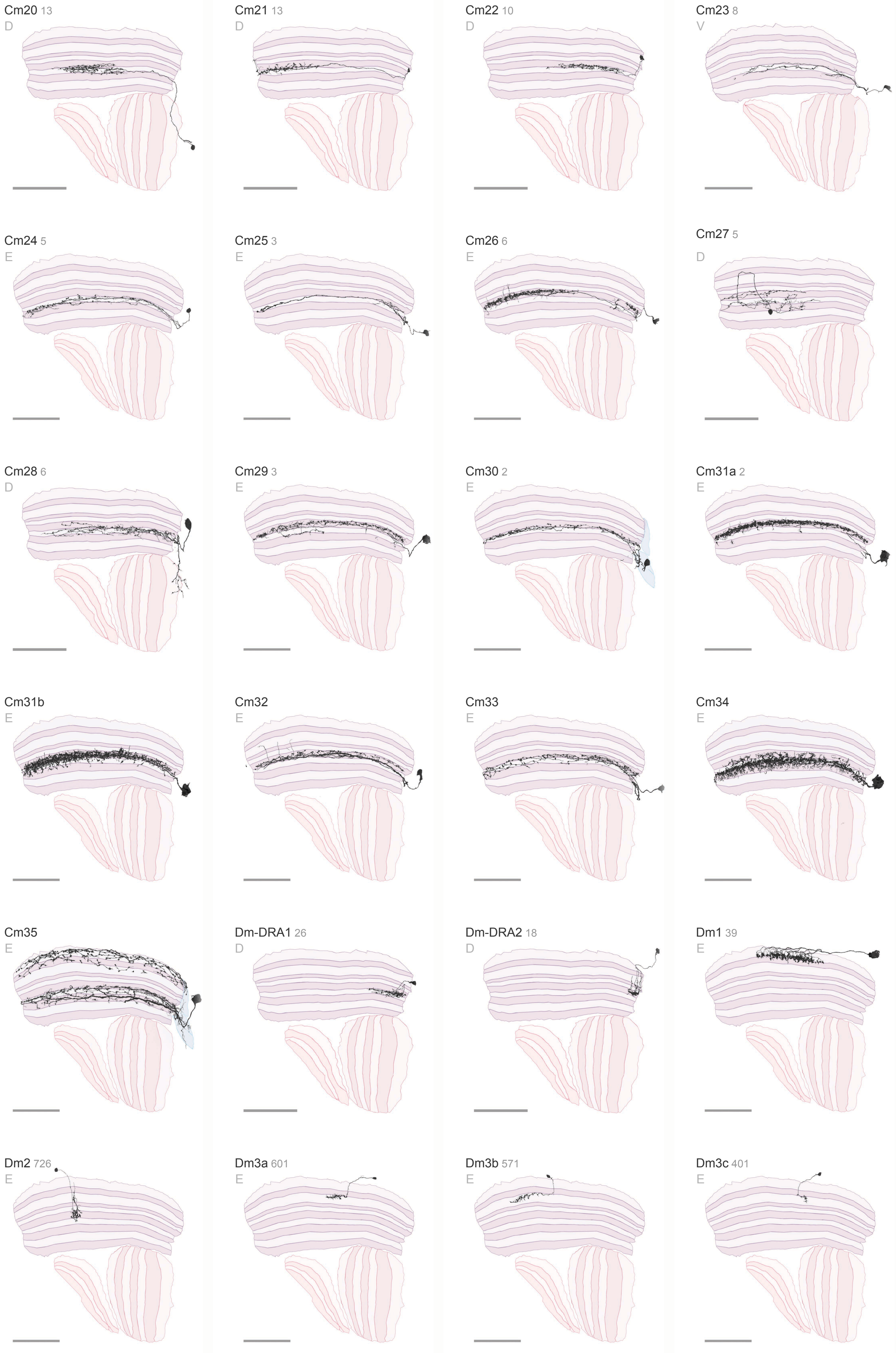

Cm20 13
D
Cm21 13
D
Cm22 10
D
Cm23 8
V
Cm24 5
E
Cm25 3
E
Cm26 6
E
Cm27 5
D
Cm28 6
D
Cm29 3
E
Cm30 2
E
Cm31a 2
E
Cm31b
E
Cm32
E
Cm33
E
Cm34
E
Cm35
E
Dm-DRA1 26
D
Dm-DRA2 18
D
Dm1 39
E
Dm2 726
E
Dm3a 601
E
Dm3b 571
E
Dm3c 401
E

#### Optic Lobe Intrinsic Neurons 2 / 7

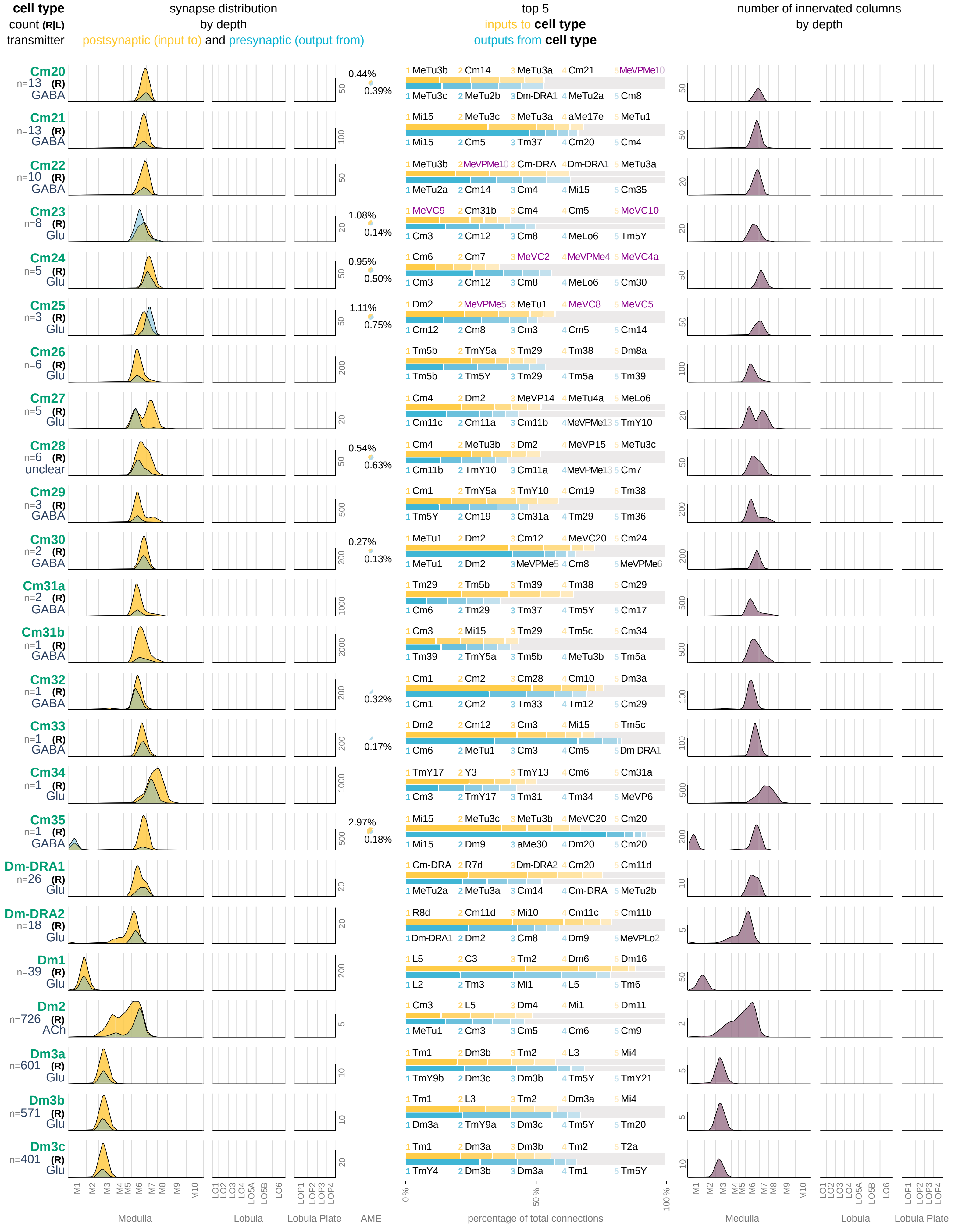

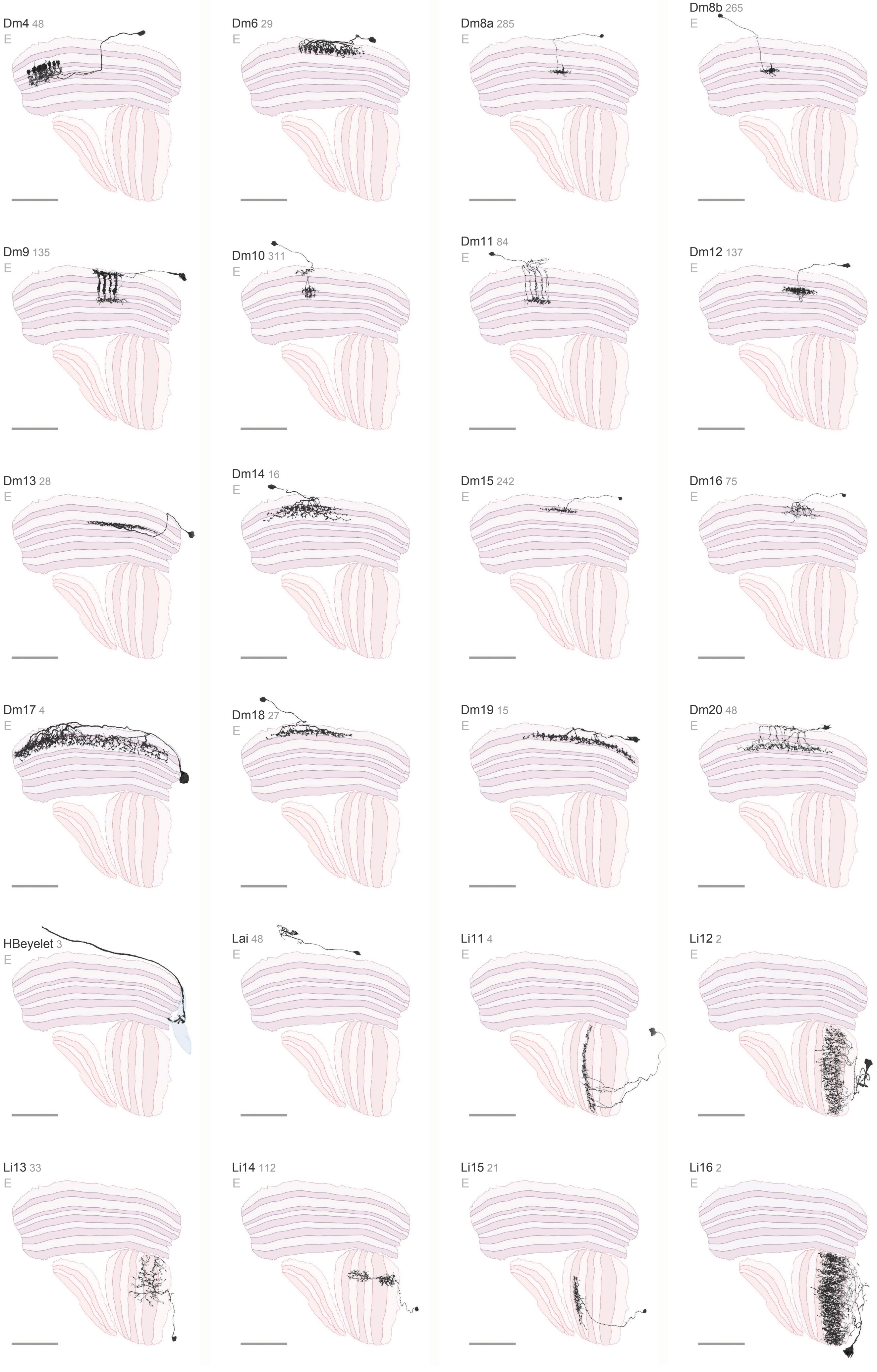

Dm4 48
E
Dm6 29
E
Dm8a 285
E
Dm8b 265
E
Dm9 135
E
Dm10 311
E
Dm11 84
E
Dm12 137
E
Dm13 28
E
Dm14 16
E
Dm15 242
E
Dm16 75
E
Dm17 4
E
Dm18 27
E
Dm19 15
E
Dm20 48
E
HBeyelet 3
E
Lai 48
E
Li11 4
E
Li12 2
E
Li13 33
E
Li14 112
E
Li15 21
E
Li16 2
E

#### Optic Lobe Intrinsic Neurons 3 / 7

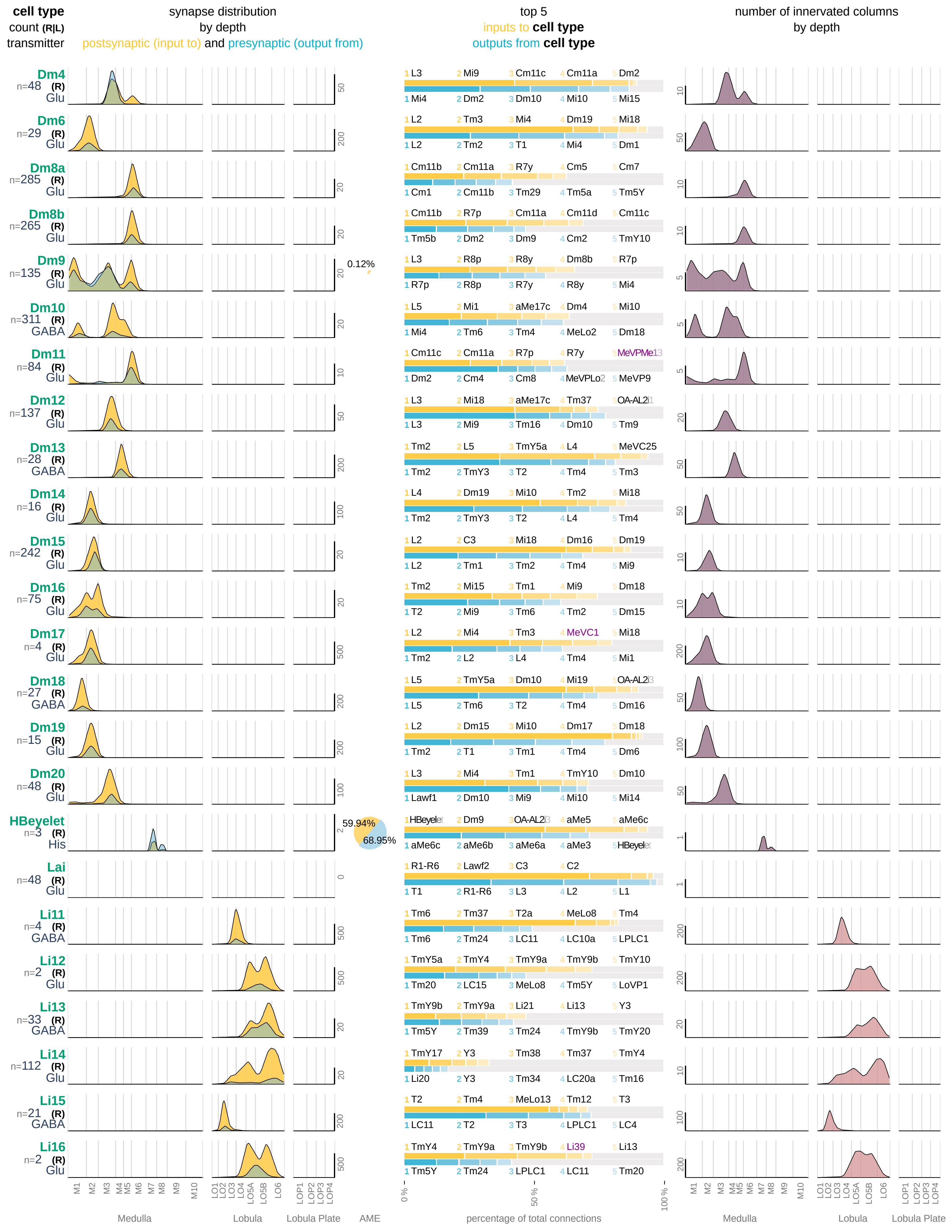

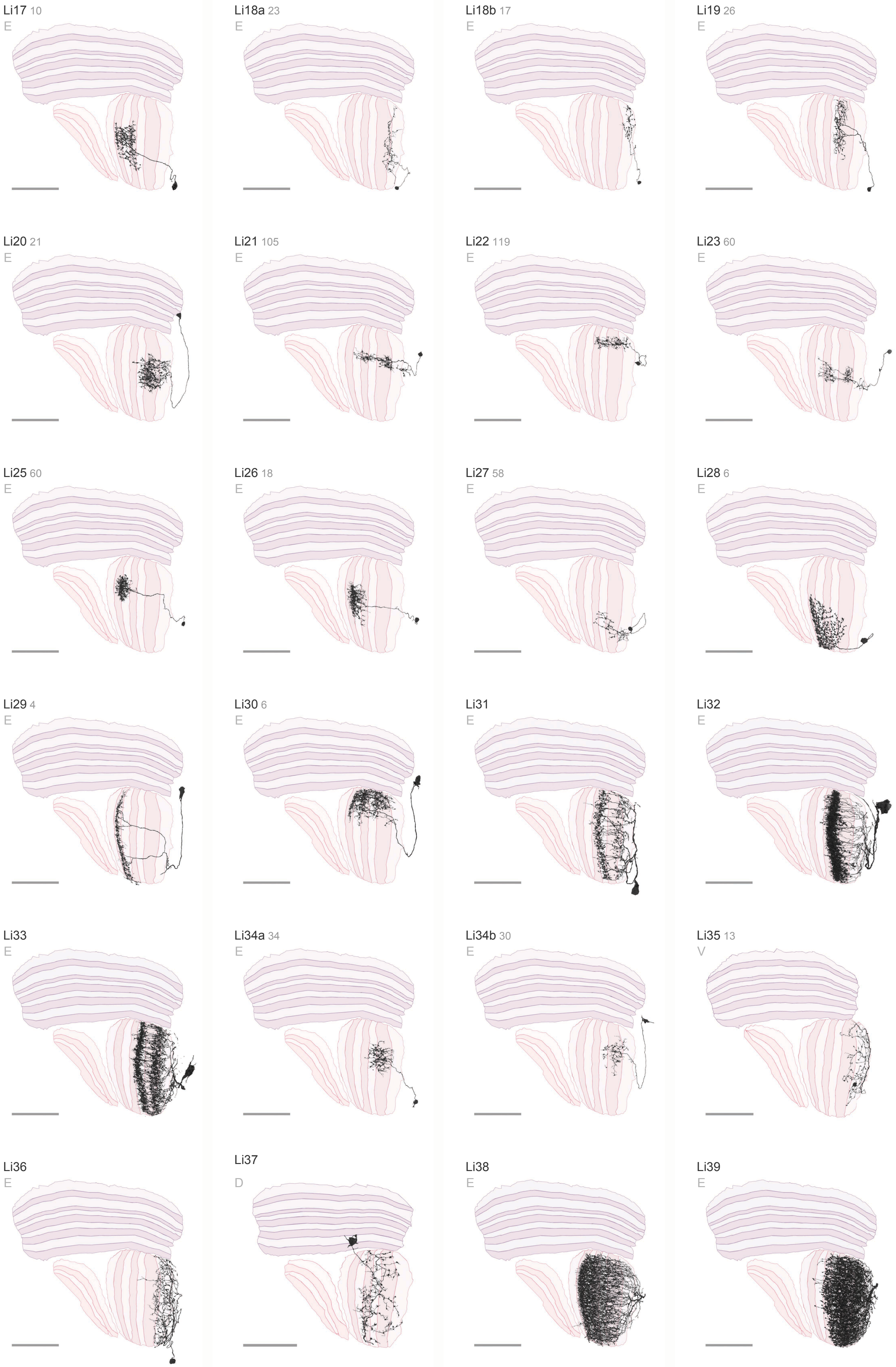

Li17 10
E
Li18a 23
E
Li18b 17
E
Li19 26
E
Li20 21
E
Li21 105
E
Li22 119
E
Li23 60
E
Li25 60
E
Li26 18
E
Li27 58
E
Li28 6
E
Li29 4
E
Li30 6
E
Li31
E
Li32
E
Li33
E
Li34a 34
E
Li34b 30
E
Li35 13
V
Li36
E
Li37
D
Li38
E
Li39
E

### **Optic Lobe Intrinsic Neurons** 4 / 7

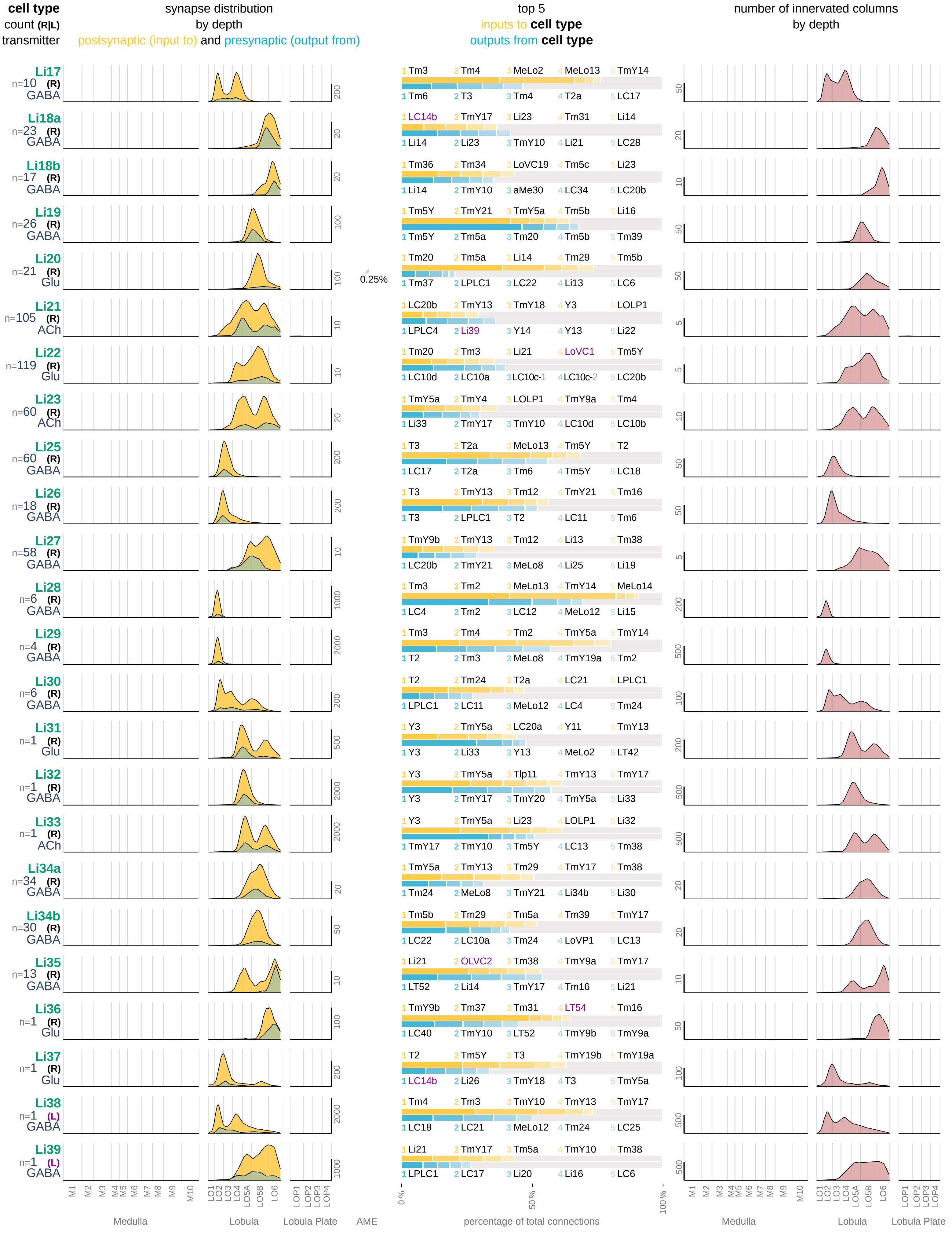

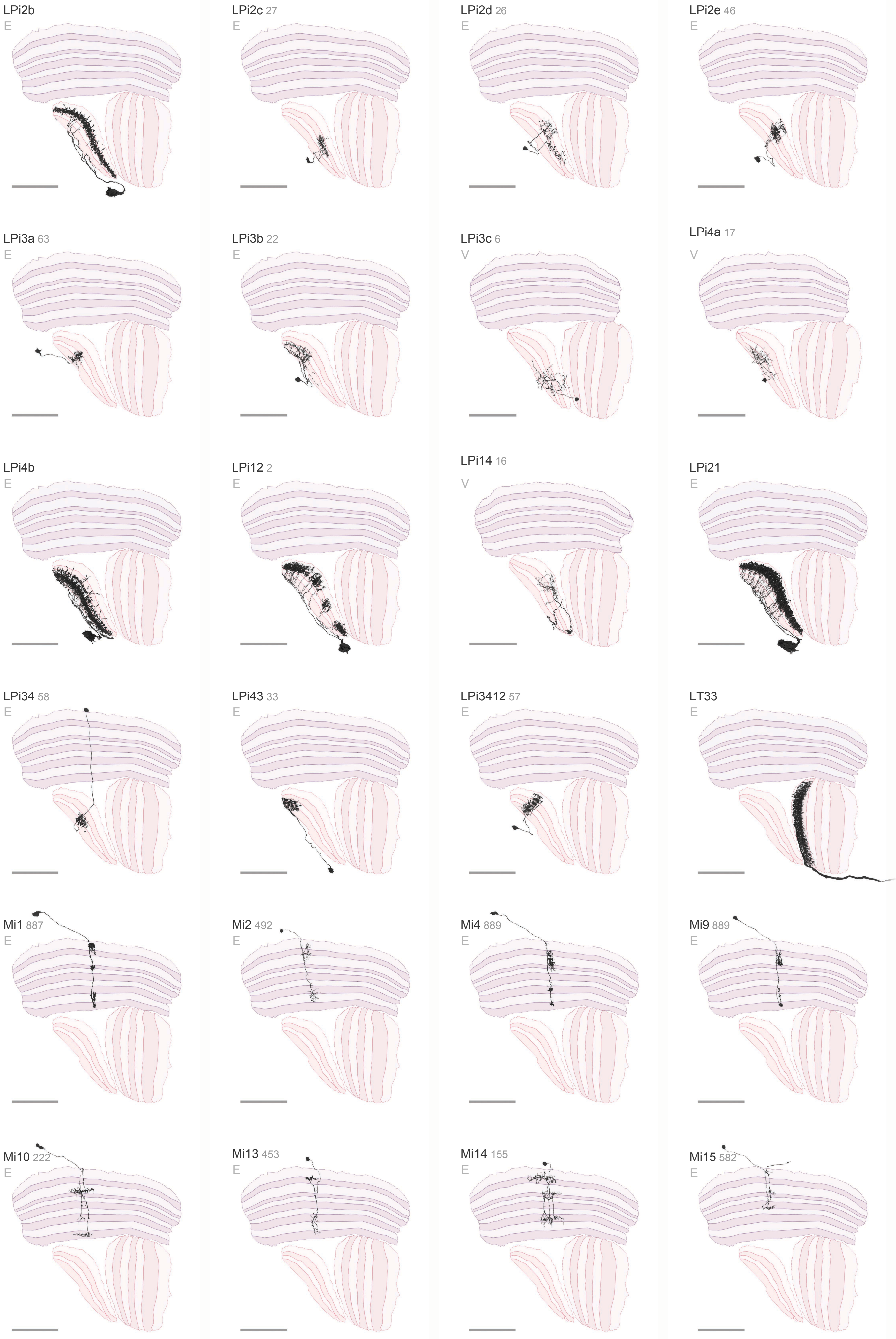
LPi2b
E
LPi2c 27
E
LPi2d 26
E
LPi2e 46
E
LPi3a 63
E
LPi3b 22
E
LPi3c 6
V
LPi4a 17
V
LPi4b
E
LPi12 2
E
LPi14 16
V
LPi21
E
LPi34 58
E
LPi43 33
E
LPi3412 57
E
LT33
E
Mi1 887
E
Mi2 492
E
Mi4 889
E
Mi9 889
E
Mi10 222
E
Mi13 453
E
Mi14 155
E
Mi15 582
E

### Optic Lobe Intrinsic Neurons 5 / 7

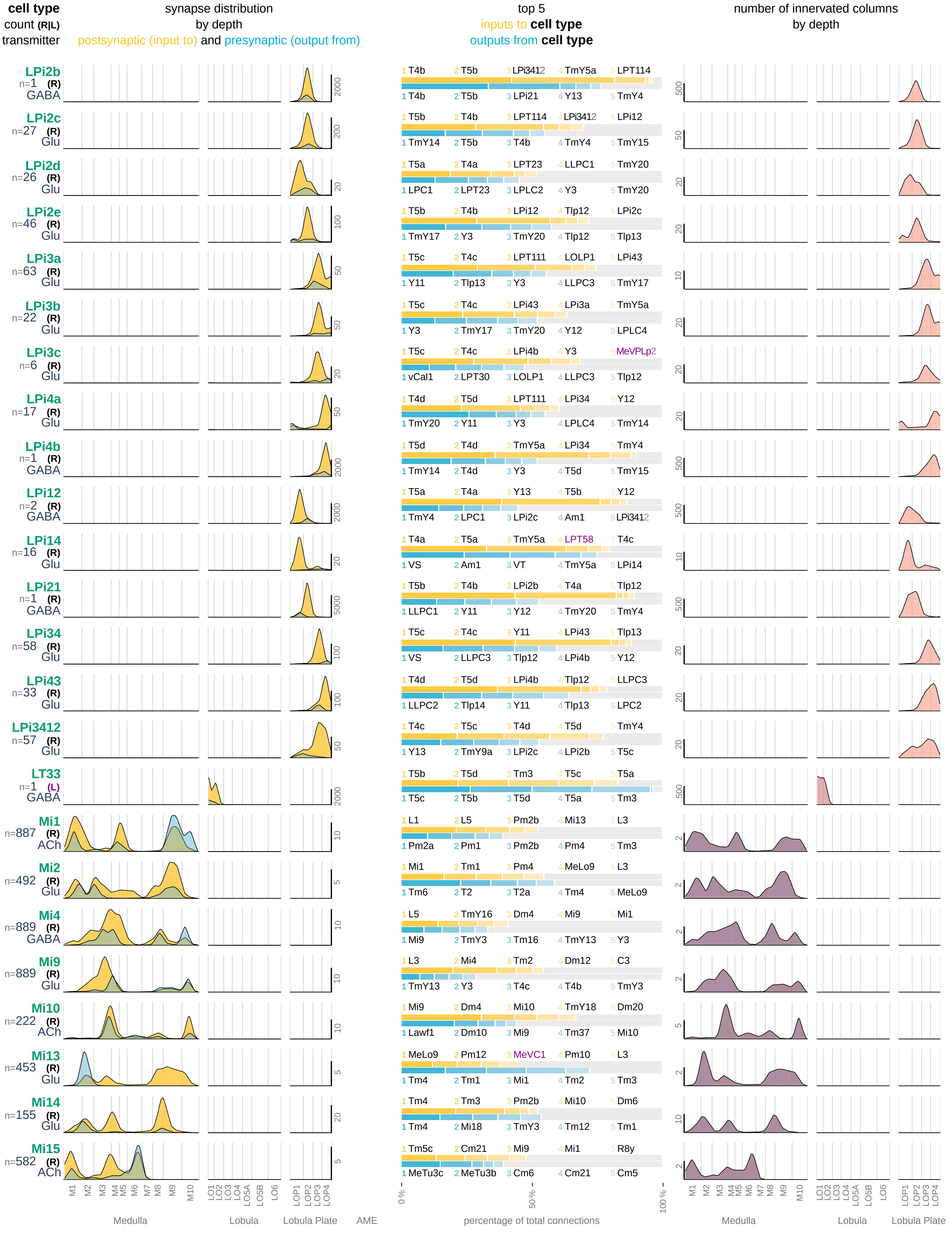

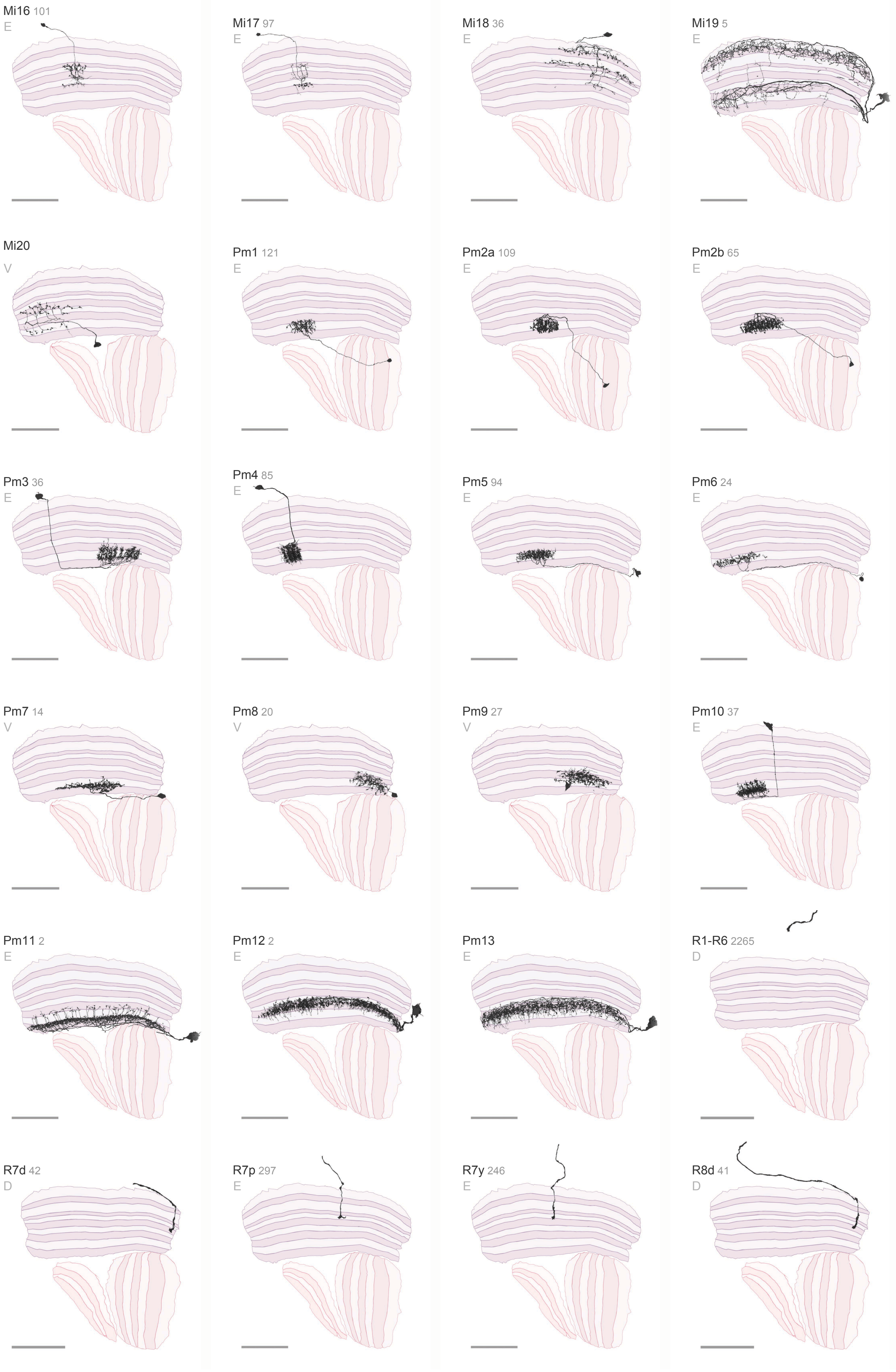

Mi16 101
E
Mi17 97
E
Mi18 36
E
Mi19 5
E
Mi20
V
Pm1 121
E
Pm2a 109
E
Pm2b 65
E
Pm3 36
E
Pm4 85
E
Pm5 94
E
Pm6 24
E
Pm7 14
V
Pm8 20
V
Pm9 27
V
Pm10 37
E
Pm11 2
E
Pm12 2
E
Pm13
E
R1-R6 2265
D
R7d 42
D
R7p 297
E
R7y 246
E
R8d 41
D

### Optic Lobe Intrinsic Neurons 6 / 7

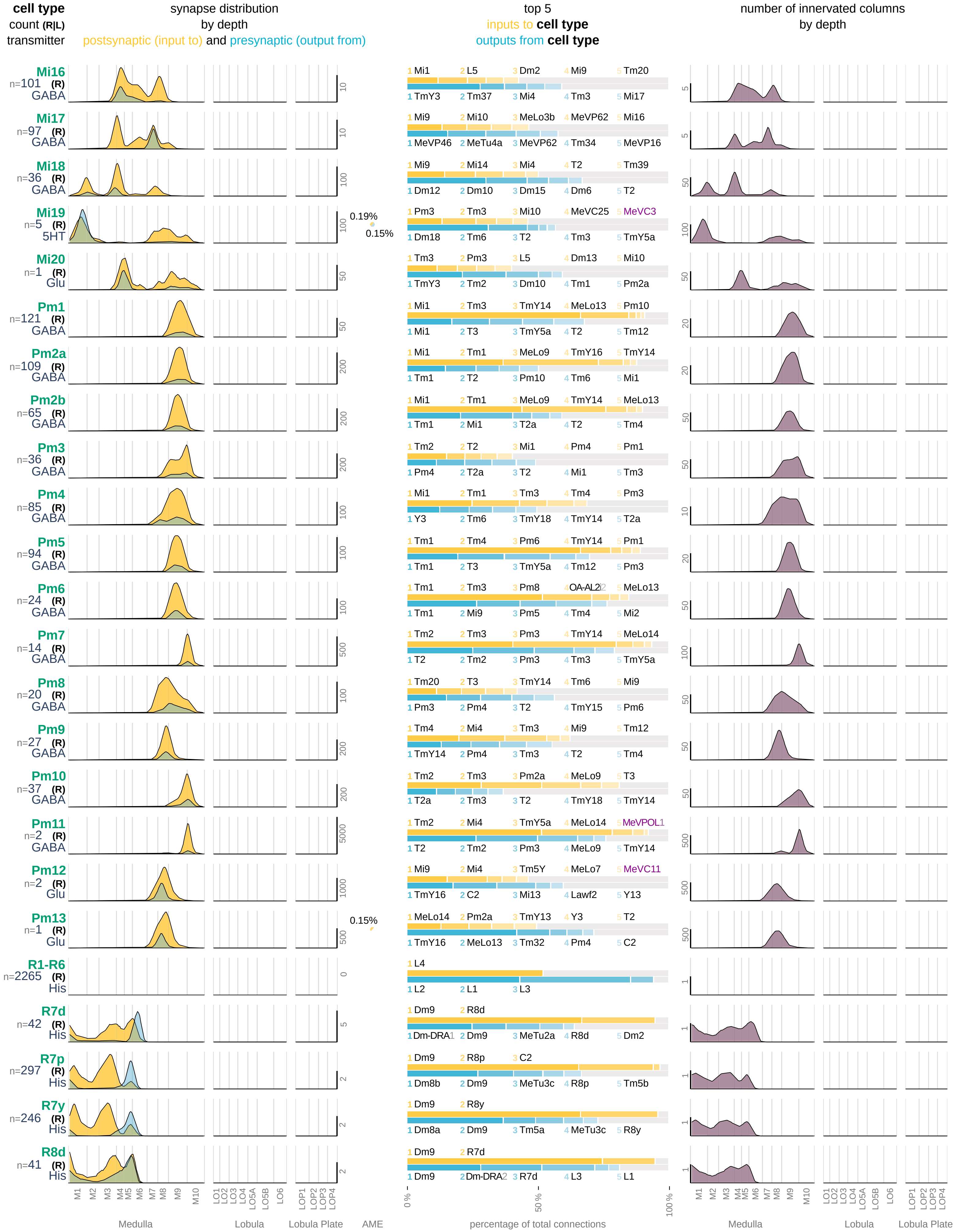

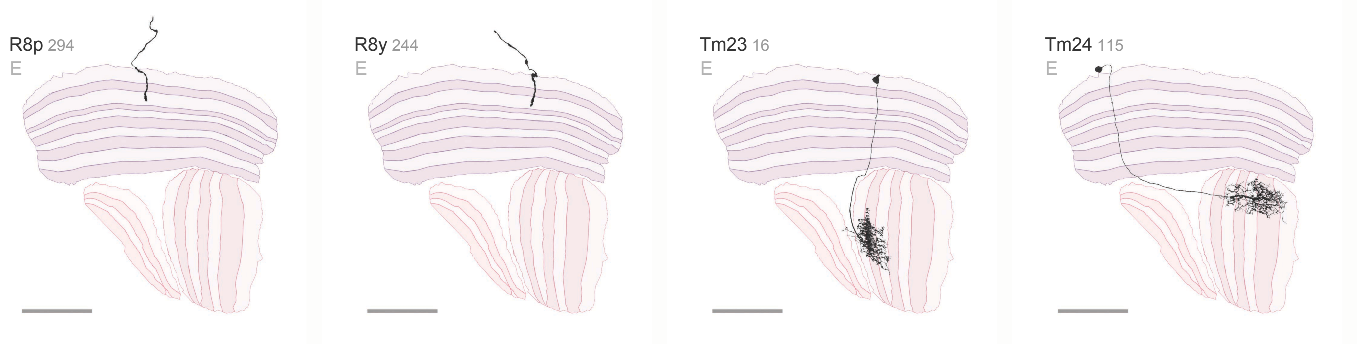
R8p 294
E
R8y 244
E
Tm23 16
E
Tm24 115
E

### **Optic Lobe Intrinsic Neurons** 7 / 7

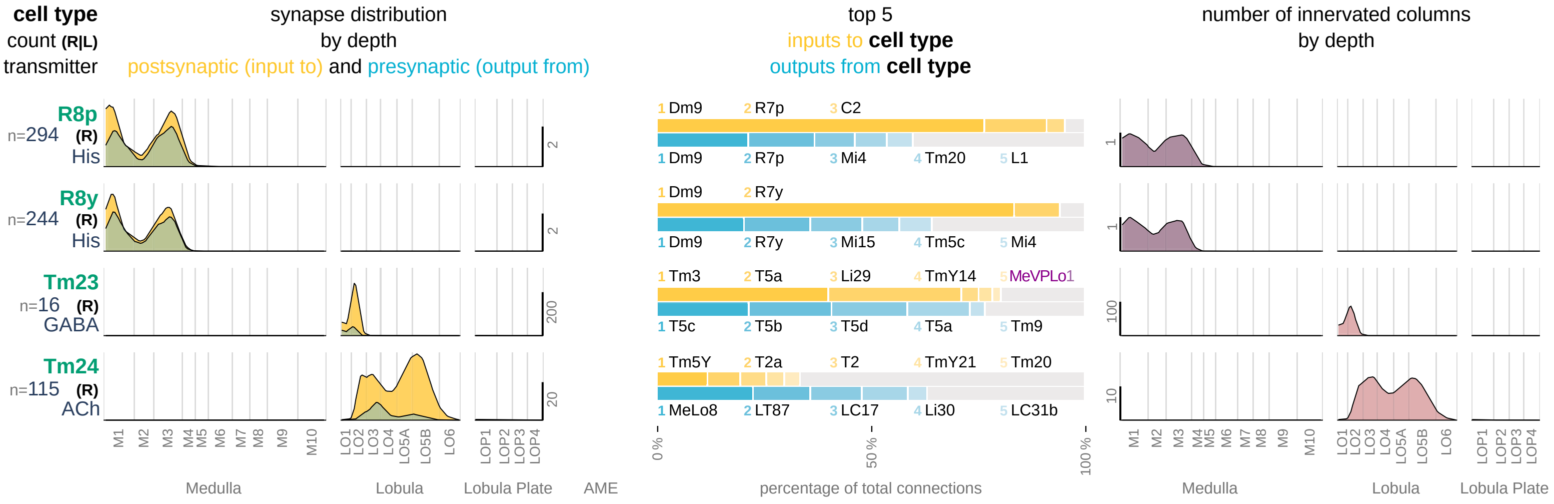

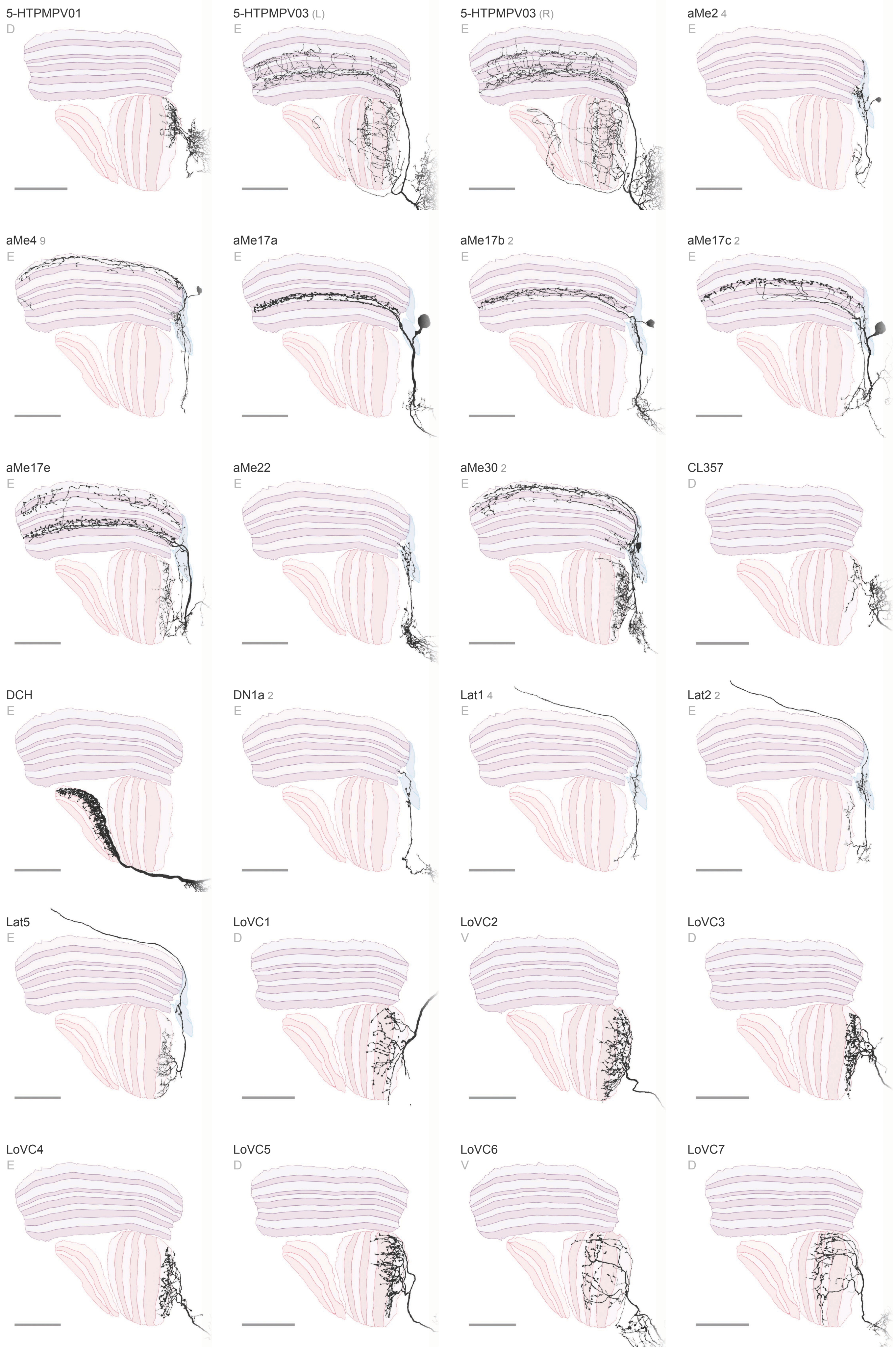

5-HTPMPV01
D
5-HTPMPV03 (L)
E
5-HTPMPV03 (R)
E
aMe2 4
E
aMe4 9
E
aMe17a
E
aMe17b 2
E
aMe17c 2
E
aMe17e
E
aMe22
E
aMe30 2
E
CL357
D
DCH
E
DN1a 2
E
Lat1 4
E
Lat2 2
E
Lat5
E
LoVC1
D
LoVC2
V
LoVC3
D
LoVC4
E
LoVC5
D
LoVC6
V
LoVC7
D

### Visual Centrifugal Neurons 1 / 5

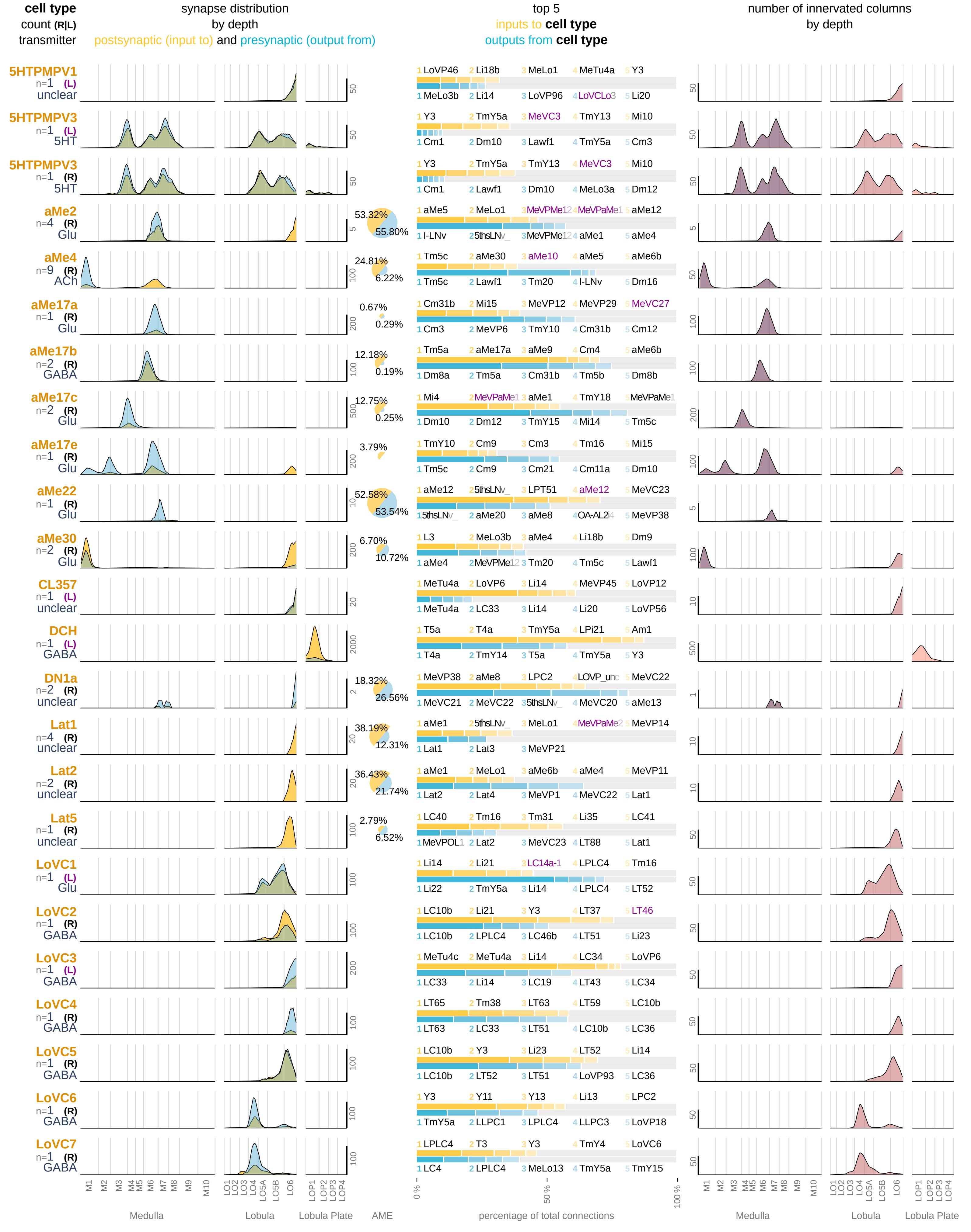

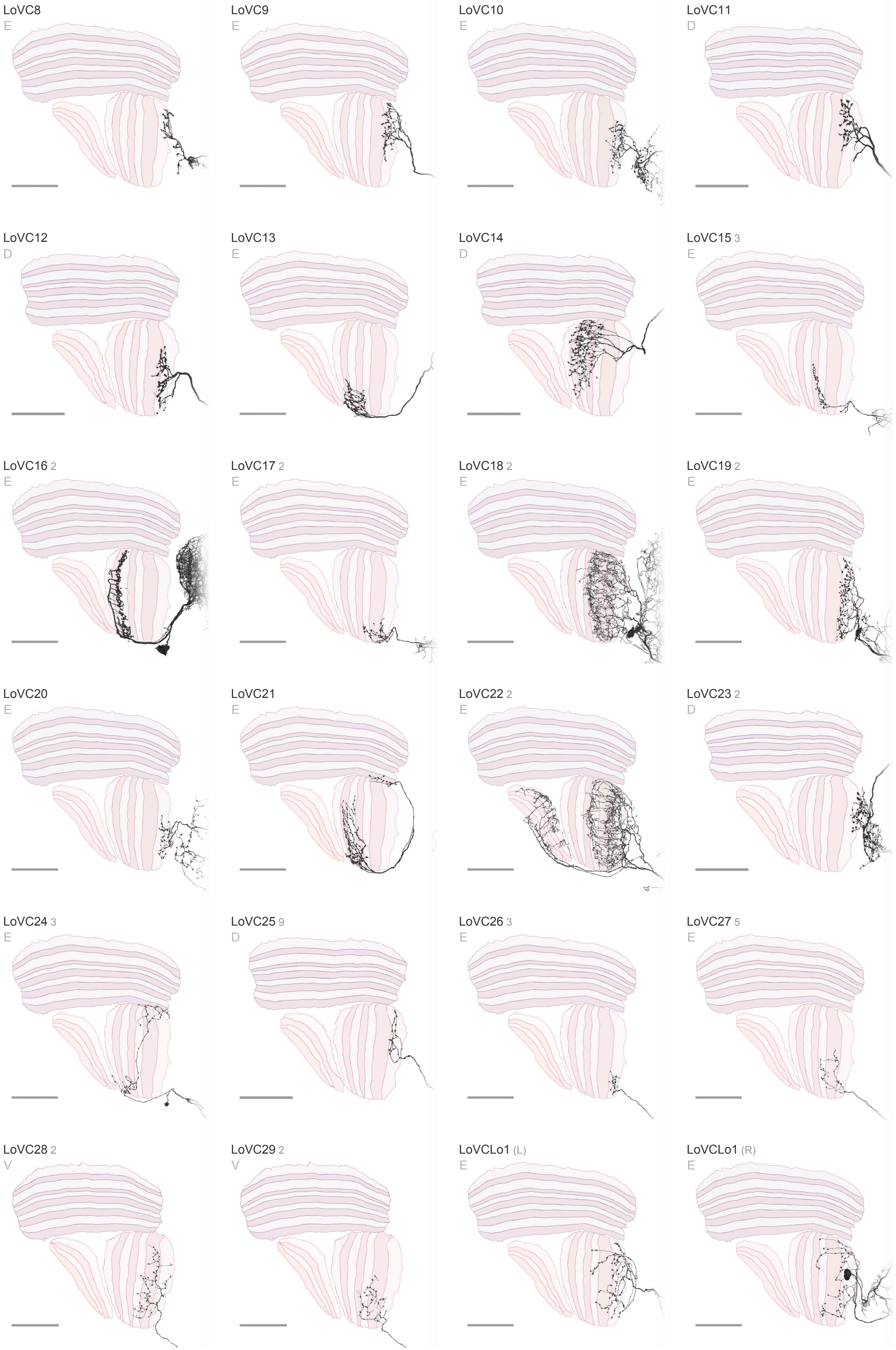

LoVC8
E
LoVC9
E
LoVC10
E
LoVC11
D
LoVC12
D
LoVC13
E
LoVC14
D
LoVC15 3
E
LoVC16 2
E
LoVC17 2
E
LoVC18 2
E
LoVC19 2
E
LoVC20
E
LoVC21
E
LoVC22 2
E
LoVC23 2
D
LoVC24 3
E
LoVC25 9
D
LoVC26 3
E
LoVC27 5
E
LoVC28 2
V
LoVC29 2
V
LoVCLo1 (L)
E
LoVCLo1 (R)
E

#### Visual Centrifugal Neurons 2 / 5

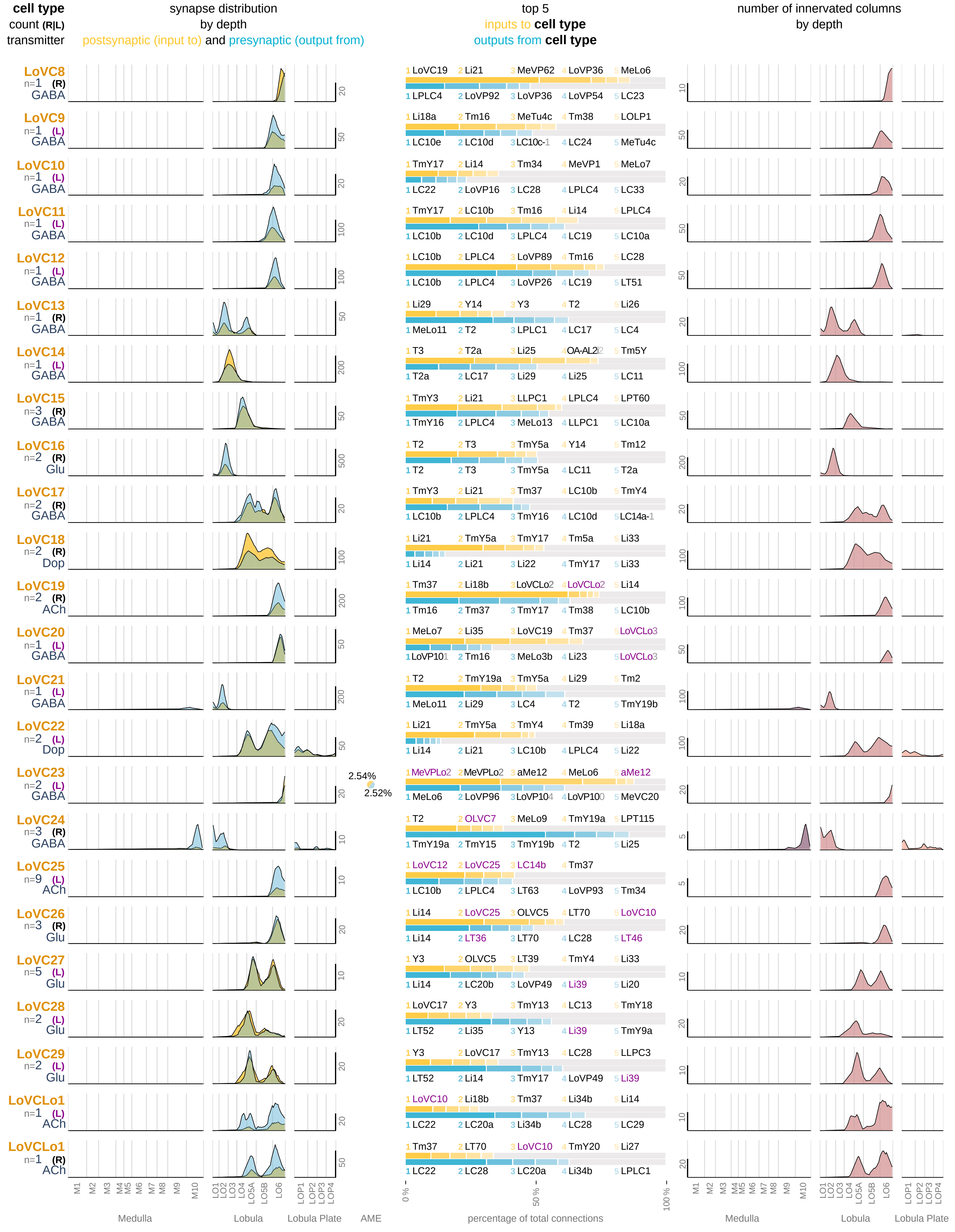

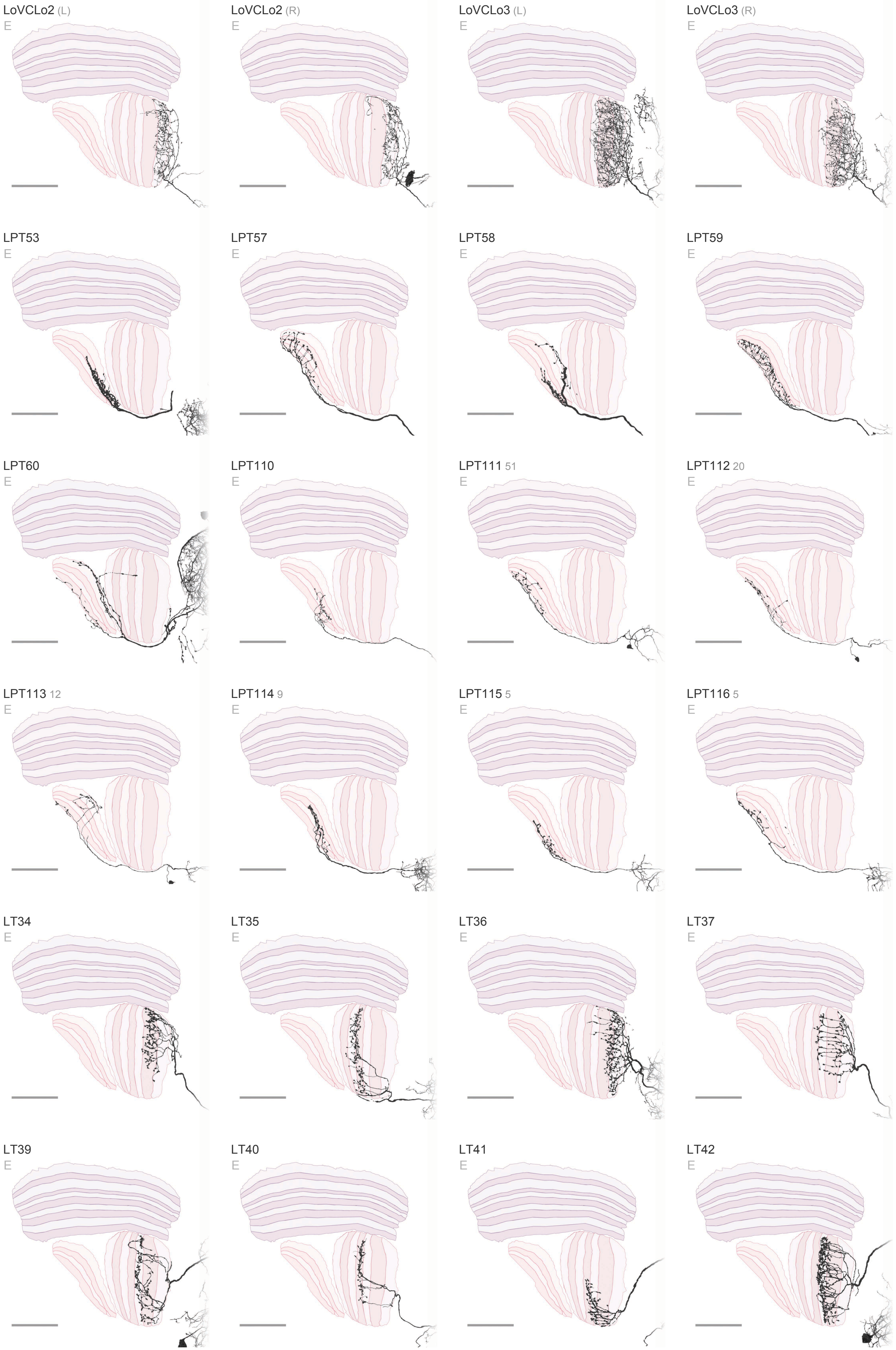
LoVCLo2 (L)
E
LoVCLo2 (R)
E
LoVCLo3 (L)
E
LoVCLo3 (R)
E
LPT53
E
LPT57
E
LPT58
E
LPT59
E
LPT60
E
LPT110
E
LPT111 51
E
LPT112 20
E
LPT113 12
E
LPT114 9
E
LPT115 5
E
LPT116 5
E
LT34
E
LT35
E
LT36
E
LT37
E
LT39
E
LT40
E
LT41
E
LT42
E

### Visual Centrifugal Neurons 3 / 5

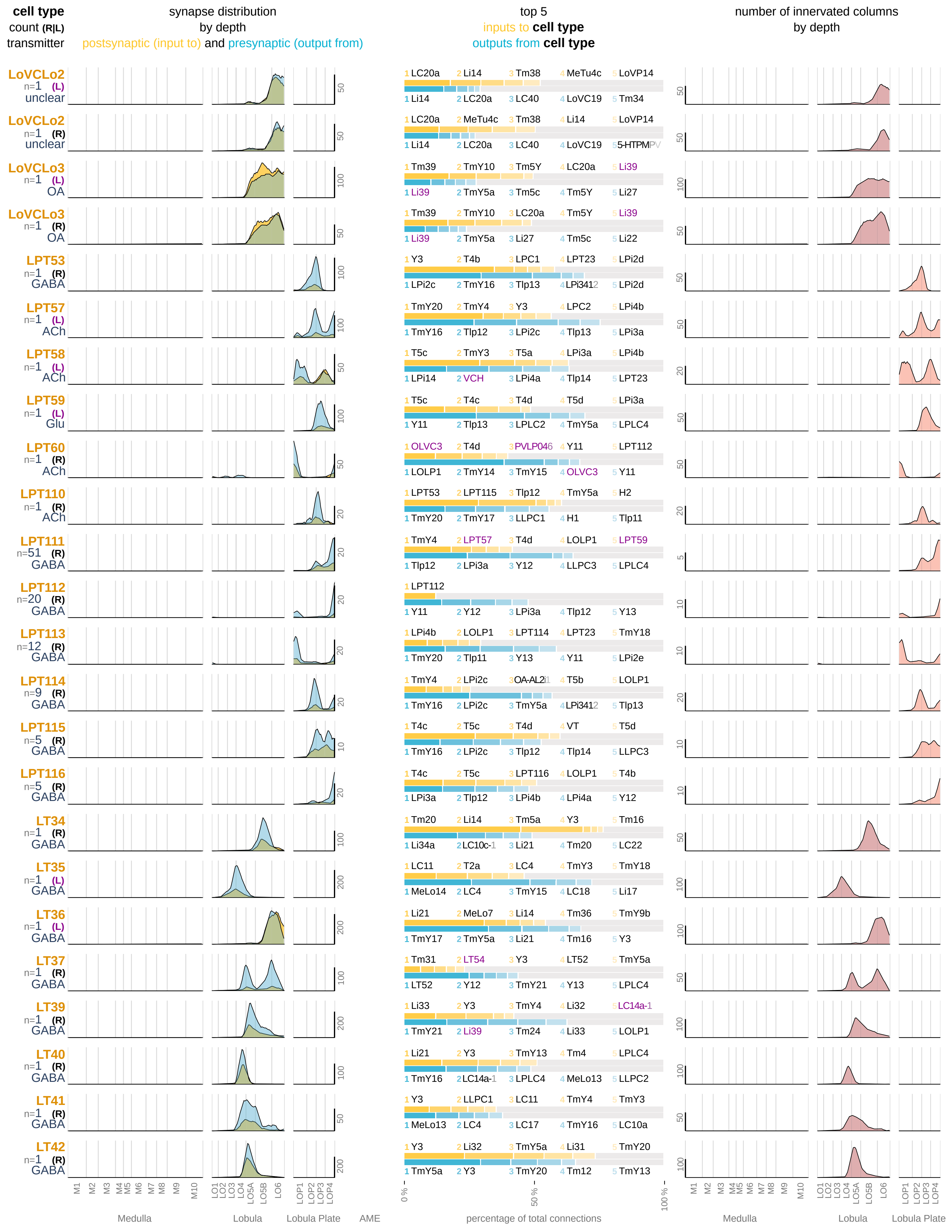

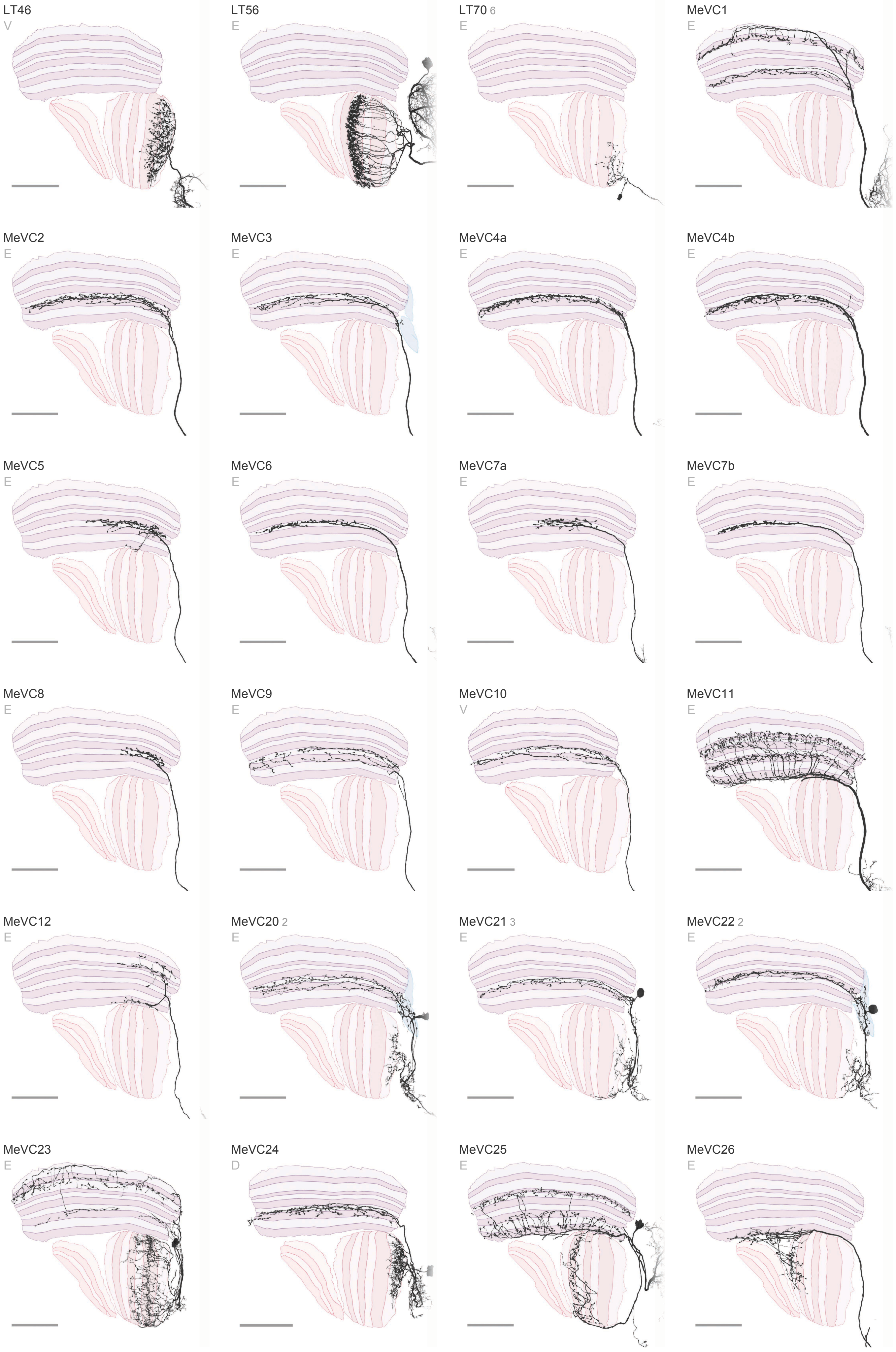

LT46
V
LT56
E
LT70 6
E
MeVC1
E
MeVC2
E
MeVC3
E
MeVC4a
E
MeVC4b
E
MeVC5
E
MeVC6
E
MeVC7a
E
MeVC7b
E
MeVC8
E
MeVC9
E
MeVC10
V
MeVC11
E
MeVC12
E
MeVC20 2
E
MeVC21 3
E
MeVC22 2
E
MeVC23
E
MeVC24
D
MeVC25
E
MeVC26
E

### Visual Centrifugal Neurons 4 / 5

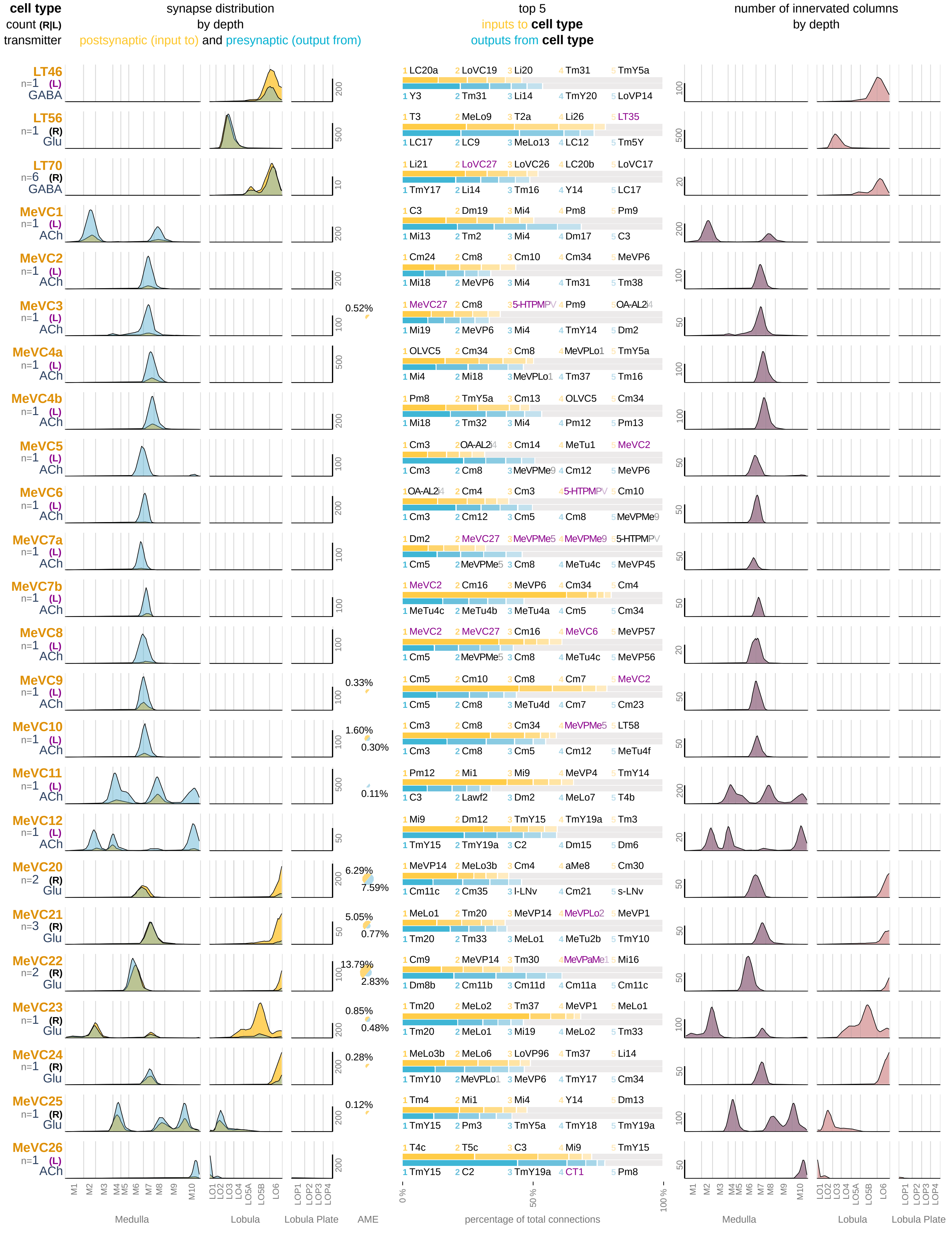

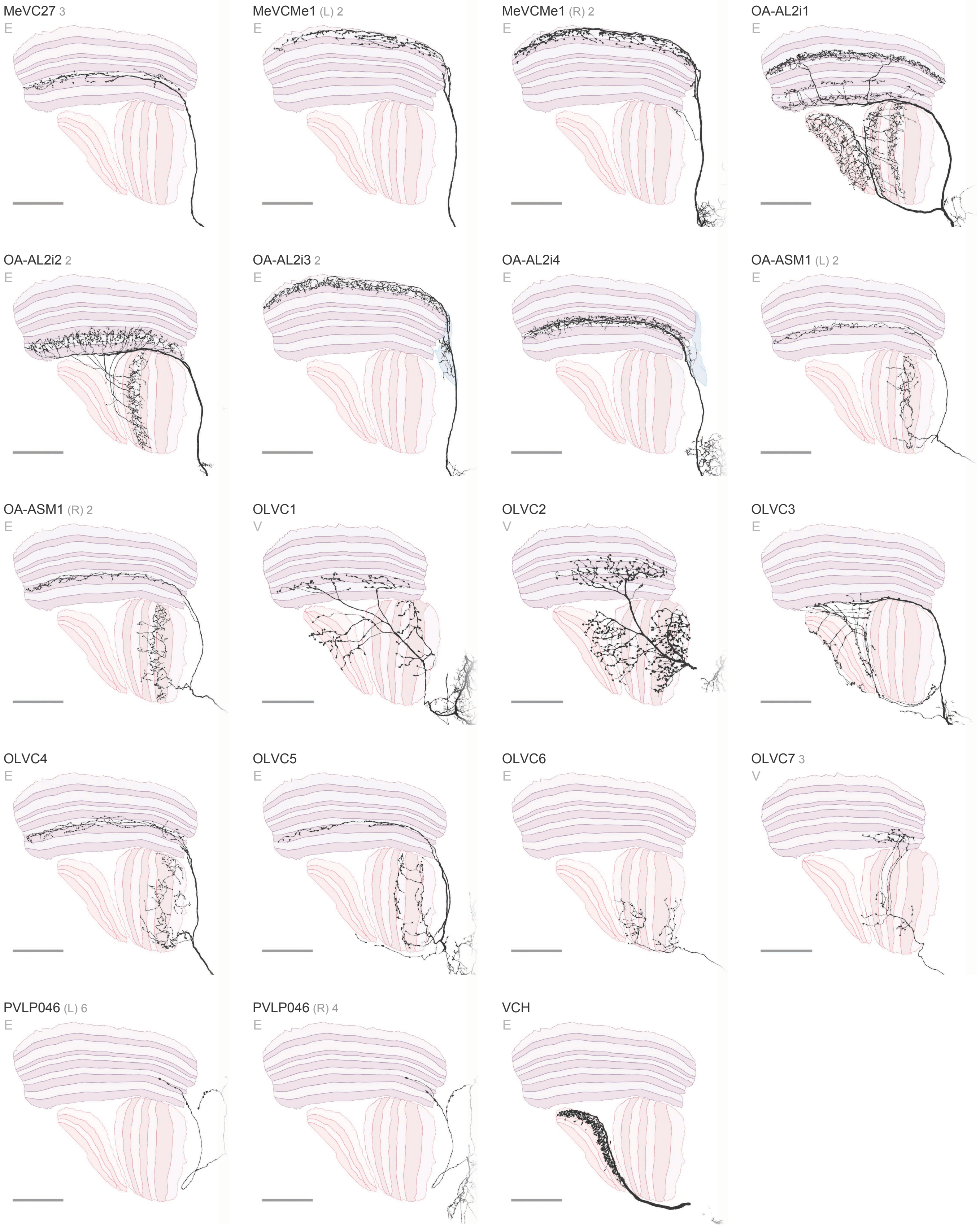

MeVC27 3
E
MeVCMe1 (L) 2
E
MeVCMe1 (R) 2
E
OA-AL2i1
E
OA-AL2i2 2
E
OA-AL2i3 2
E
OA-AL2i4
E
OA-ASM1 (L) 2
E
OA-ASM1 (R) 2
E
OLVC1
V
OLVC2
V
OLVC3
E
OLVC4
E
OLVC5
E
OLVC6
E
OLVC7 3
V
PVLP046 (L) 6
E
PVLP046 (R) 4
E
VCH
E

### Visual Centrifugal Neurons 5 / 5

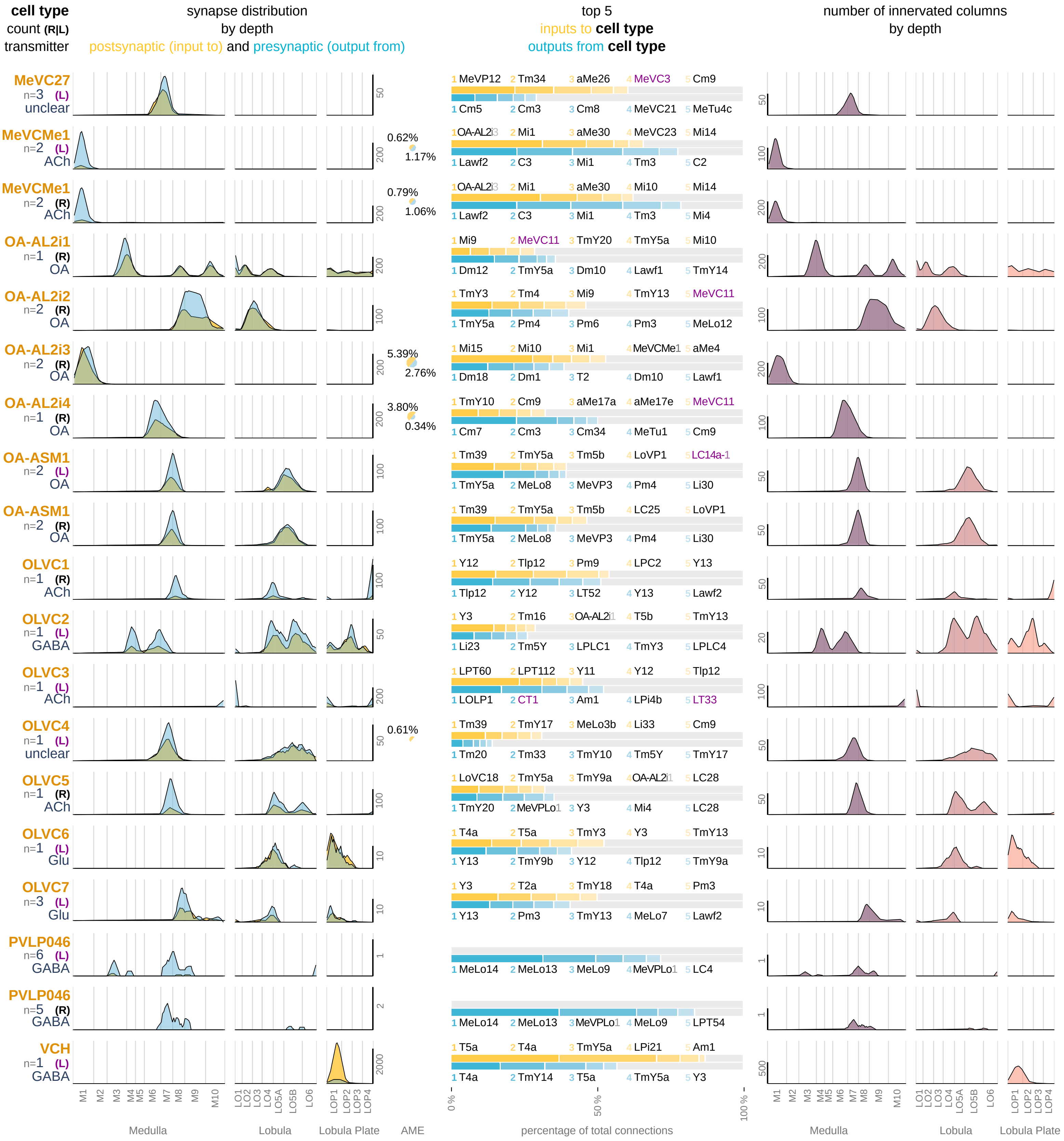

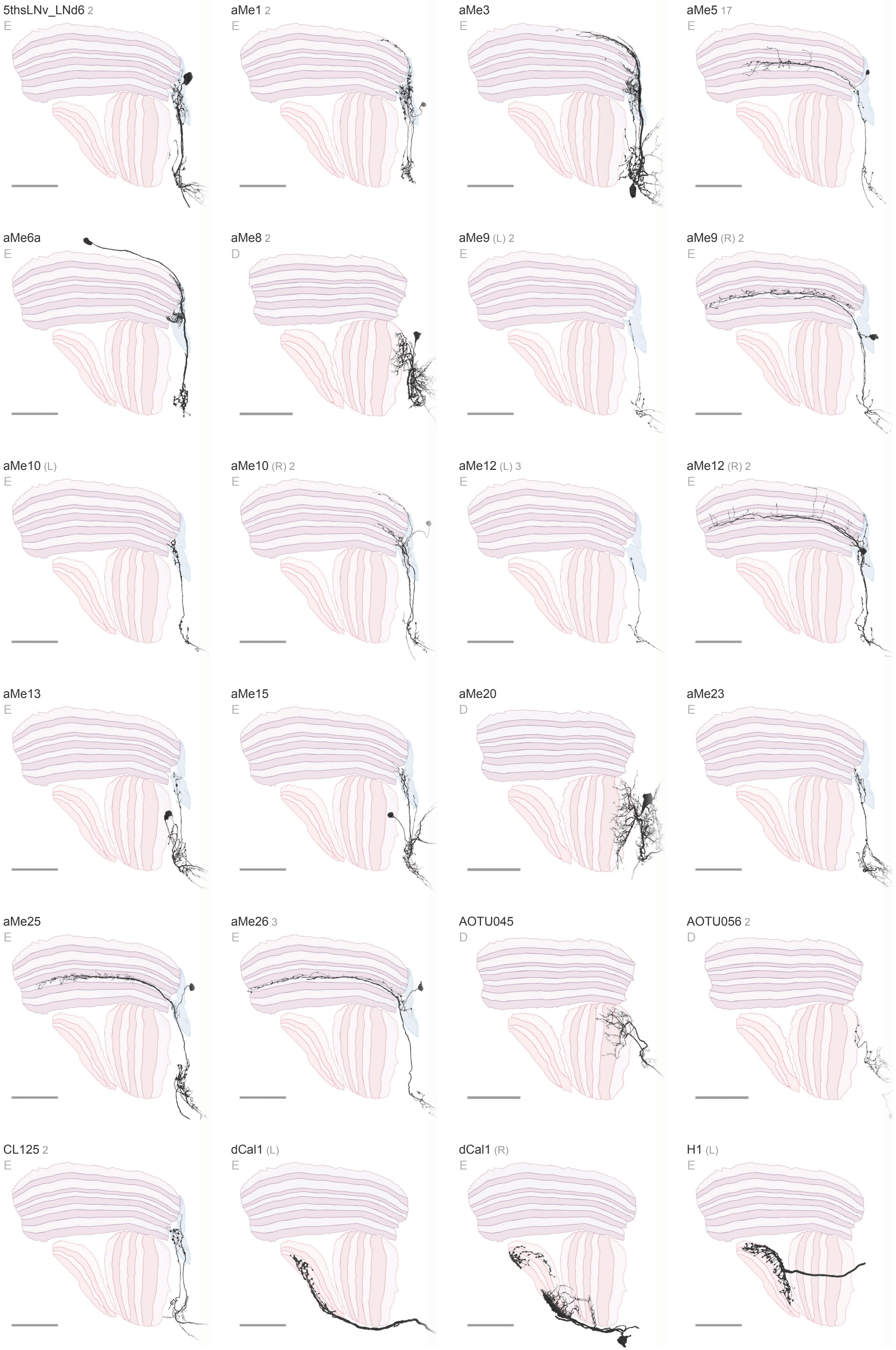
5thsLNv_LNd6 2
E
aMe1 2
E
aMe3
E
aMe5 17
E
aMe6a
E
aMe8 2
D
aMe9 (L) 2
E
aMe9 (R) 2
E
aMe10 (L)
E
aMe10 (R) 2
E
aMe12 (L) 3
E
aMe12 (R) 2
E
aMe13
E
aMe15
E
aMe20
D
aMe23
E
aMe25
E
aMe26 3
E
AOTU045
D
AOTU056 2
D
CL125 2
E
dCal1 (L)
E
dCal1 (R)
E
H1 (L)
E

#### Visual Projection Neurons 1 / 16

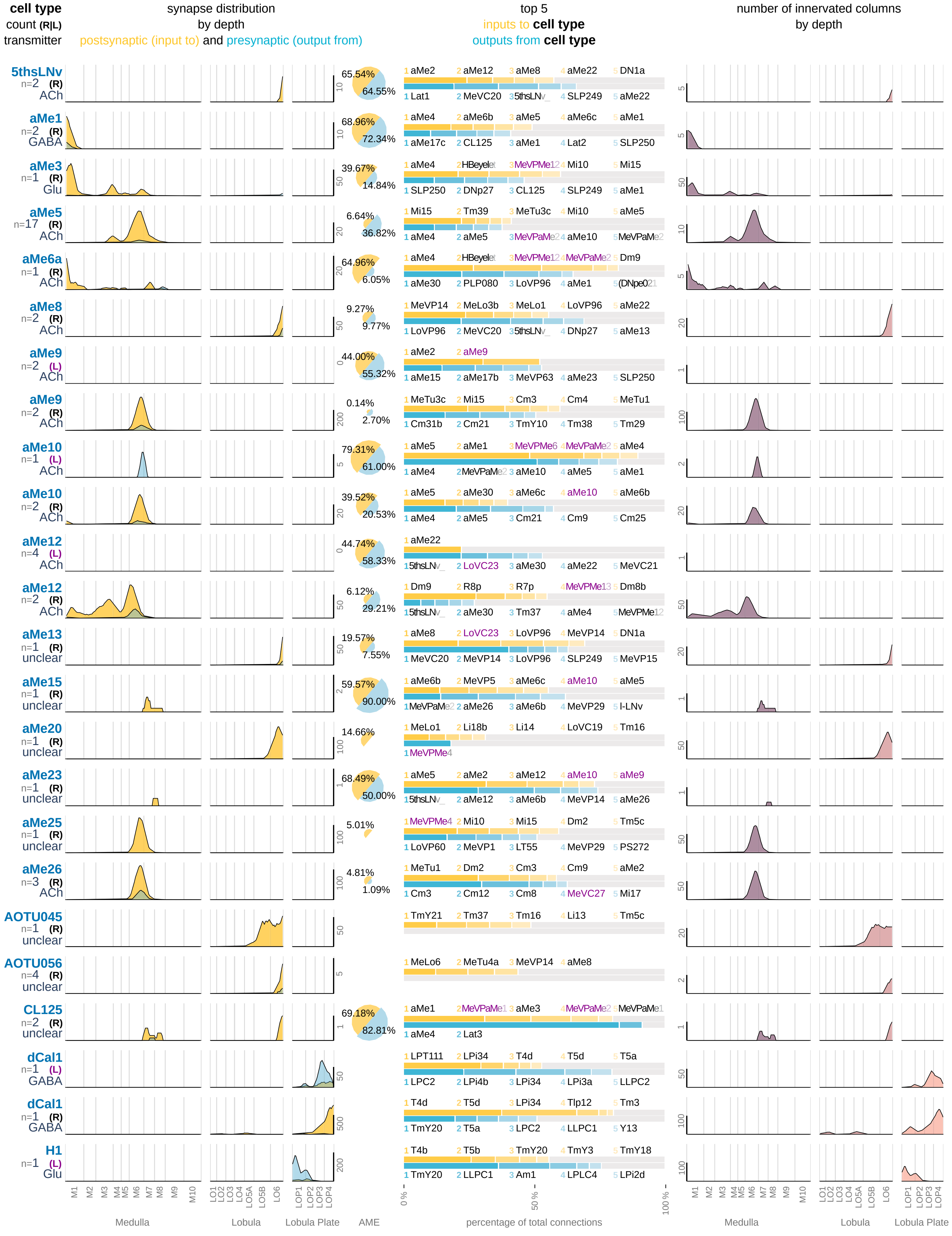

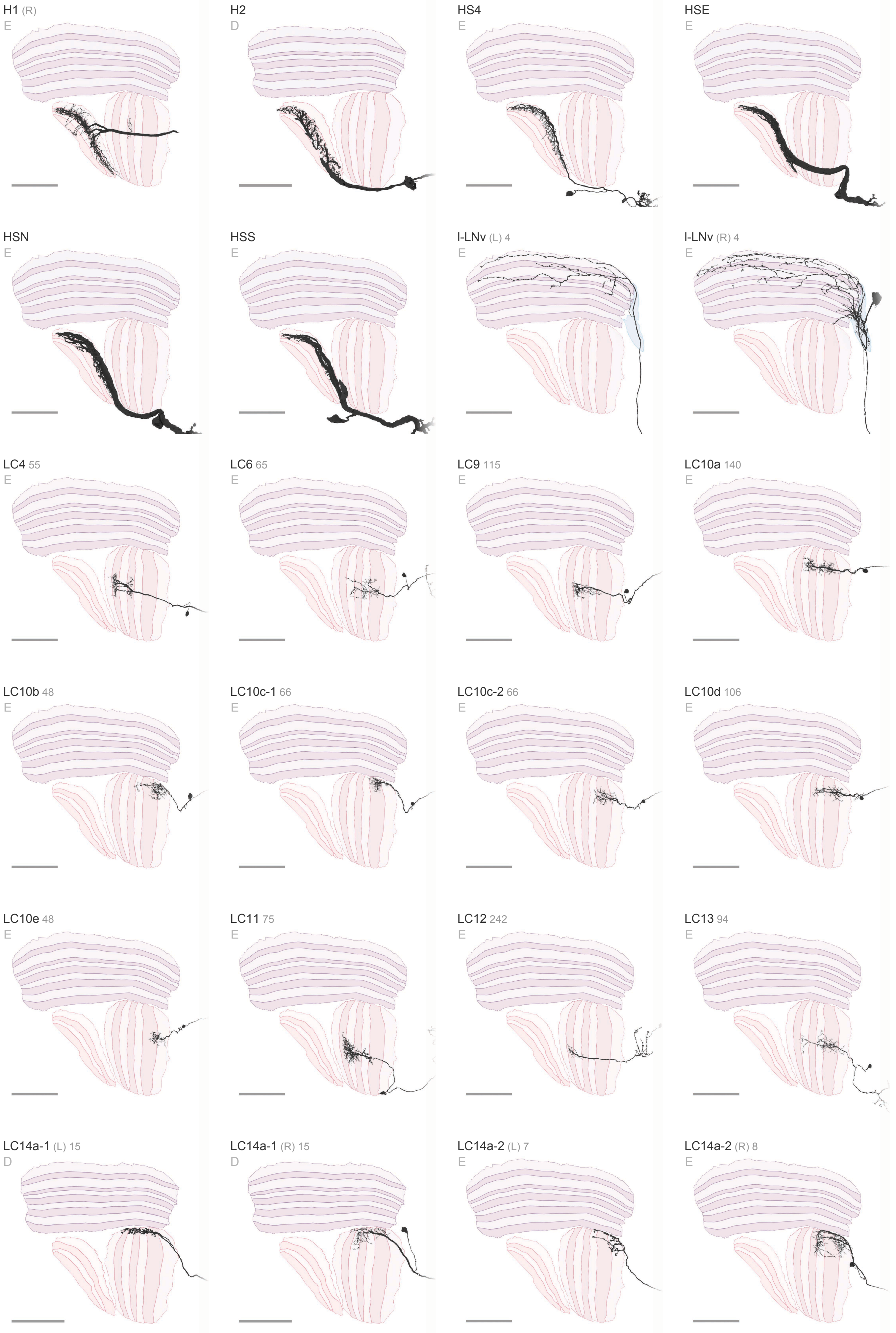
H1 (R)
E
H2
D
HS4
E
HSE
E
HSN
E
HSS
E
l-LNv (L) 4
E
l-LNv (R) 4
E
LC4 55
E
LC6 65
E
LC9 115
E
LC10a 140
E
LC10b 48
E
LC10c-1 66
E
LC10c-2 66
E
LC10d 106
E
LC10e 48
E
LC11 75
E
LC12 242
E
LC13 94
E
LC14a-1 (L) 15
D
LC14a-1 (R) 15
D
LC14a-2 (L) 7
E
LC14a-2 (R) 8
E

**Visual Projection Neurons** 2 / 16

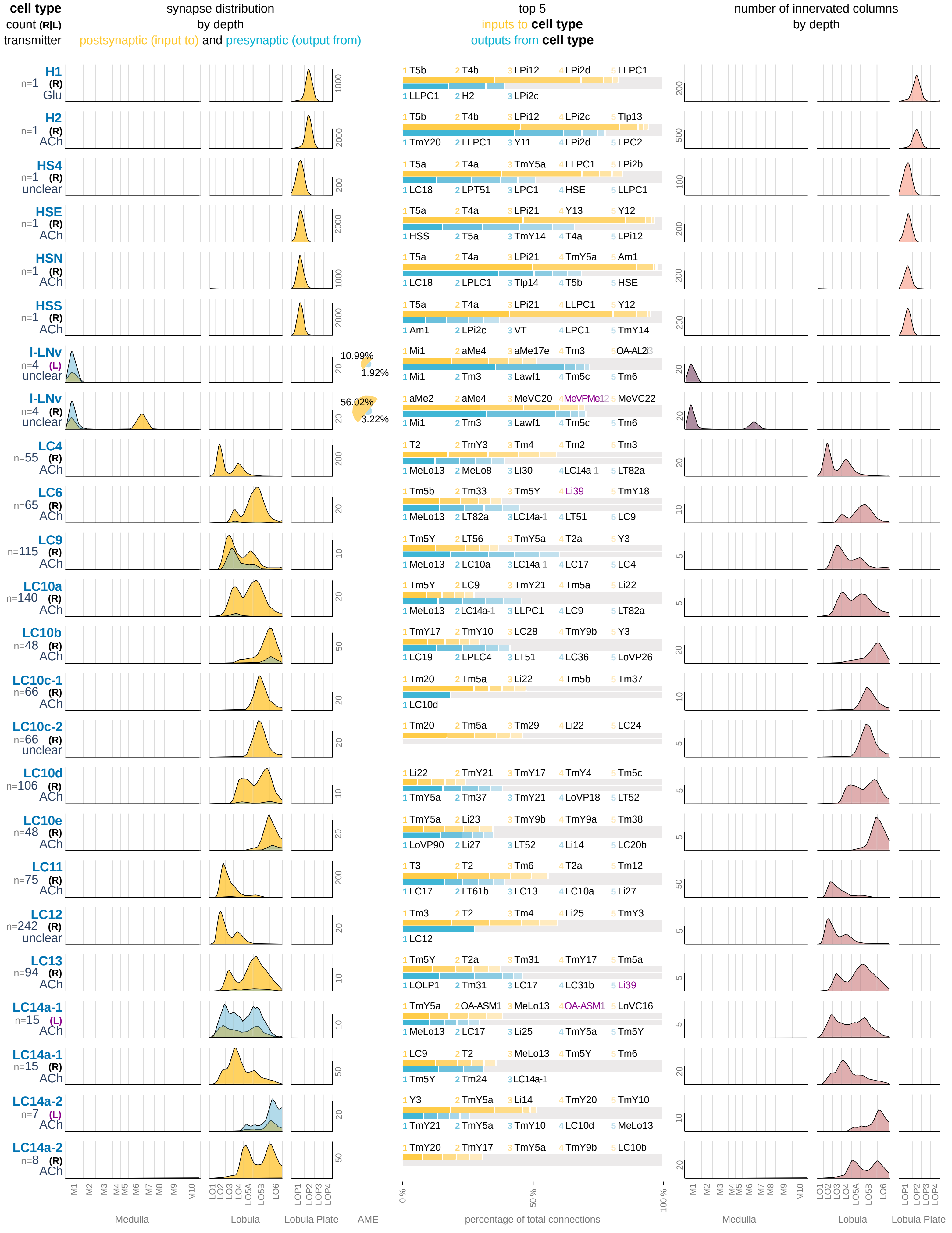

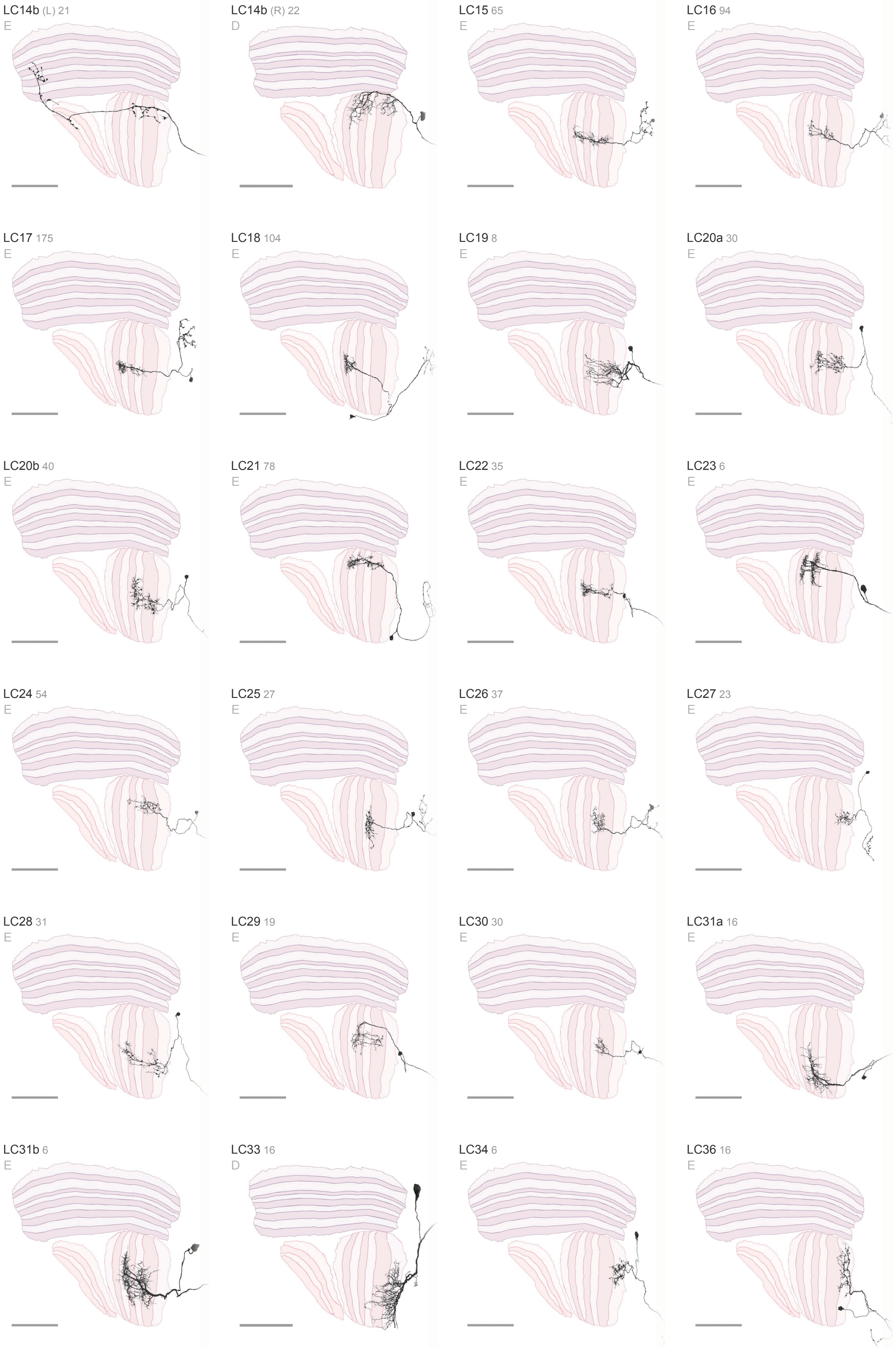
LC14b (L) 21
E
LC14b (R) 22
D
LC15 65
E
LC16 94
E
LC17 175
E
LC18 104
E
LC19 8
E
LC20a 30
E
LC20b 40
E
LC21 78
E
LC22 35
E
LC23 6
E
LC24 54
E
LC25 27
E
LC26 37
E
LC27 23
E
LC28 31
E
LC29 19
E
LC30 30
E
LC31a 16
E
LC31b 6
E
LC33 16
D
LC34 6
E
LC36 16
E

### Visual Projection Neurons 3 / 16

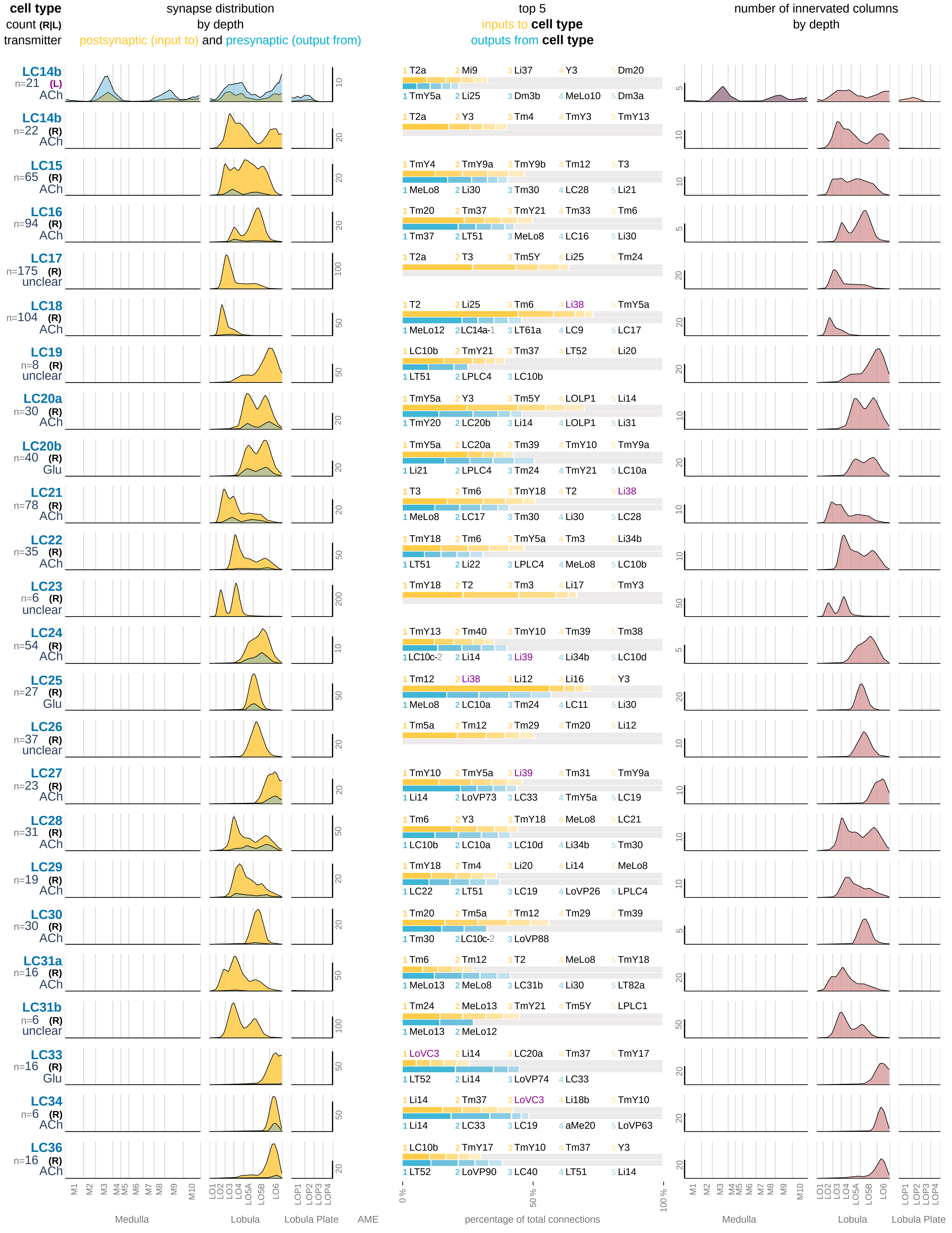

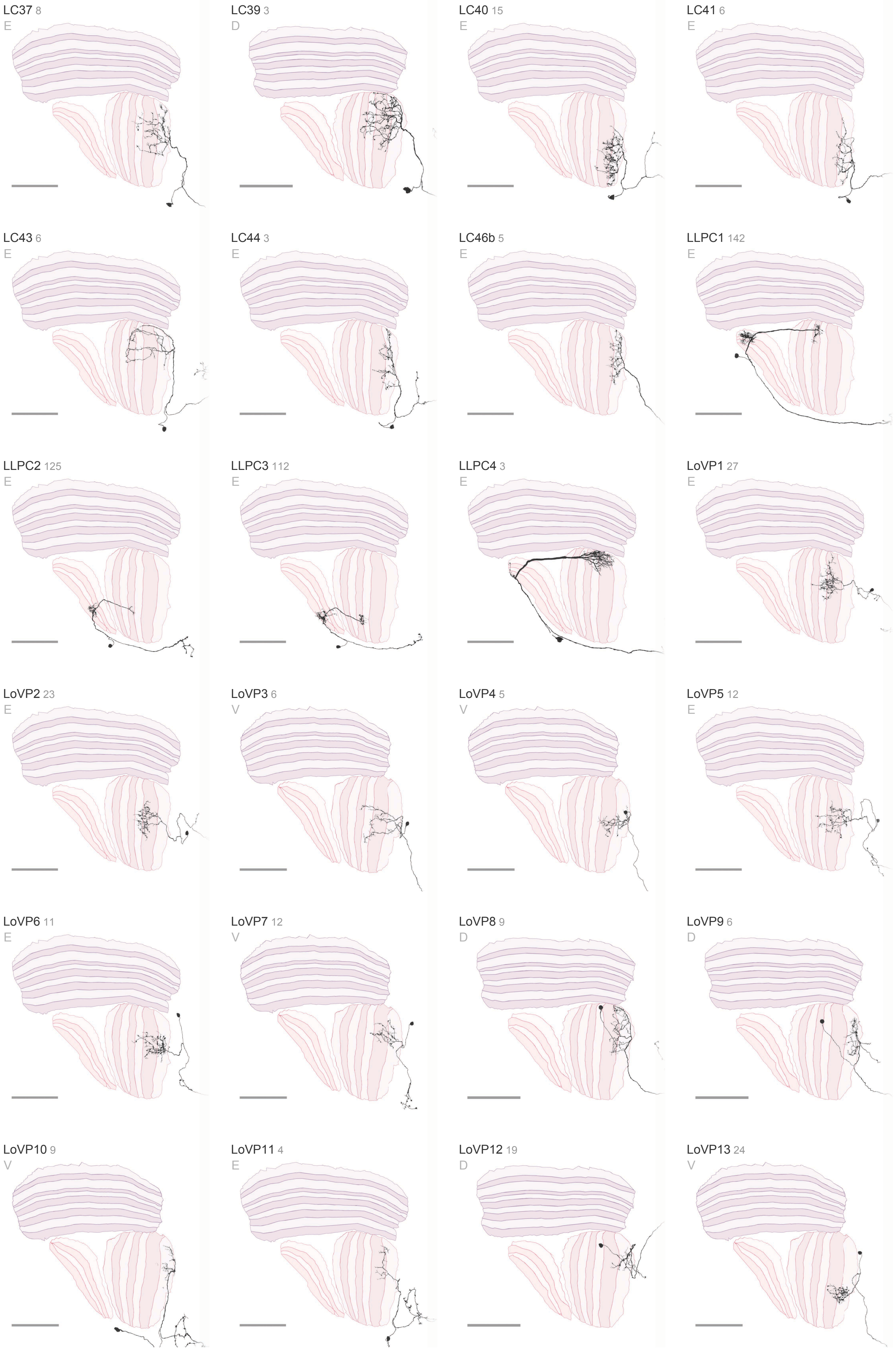
LC37 8
E
LC39 3
D
LC40 15
E
LC41 6
E
LC43 6
E
LC44 3
E
LC46b 5
E
LLPC1 142
E
LLPC2 125
E
LLPC3 112
E
LLPC4 3
E
LoVP1 27
E
LoVP2 23
E
LoVP3 6
V
LoVP4 5
V
LoVP5 12
E
LoVP6 11
E
LoVP7 12
V
LoVP8 9
D
LoVP9 6
D
LoVP10 9
V
LoVP11 4
E
LoVP12 19
D
LoVP13 24
V

### Visual Projection Neurons 4 / 16

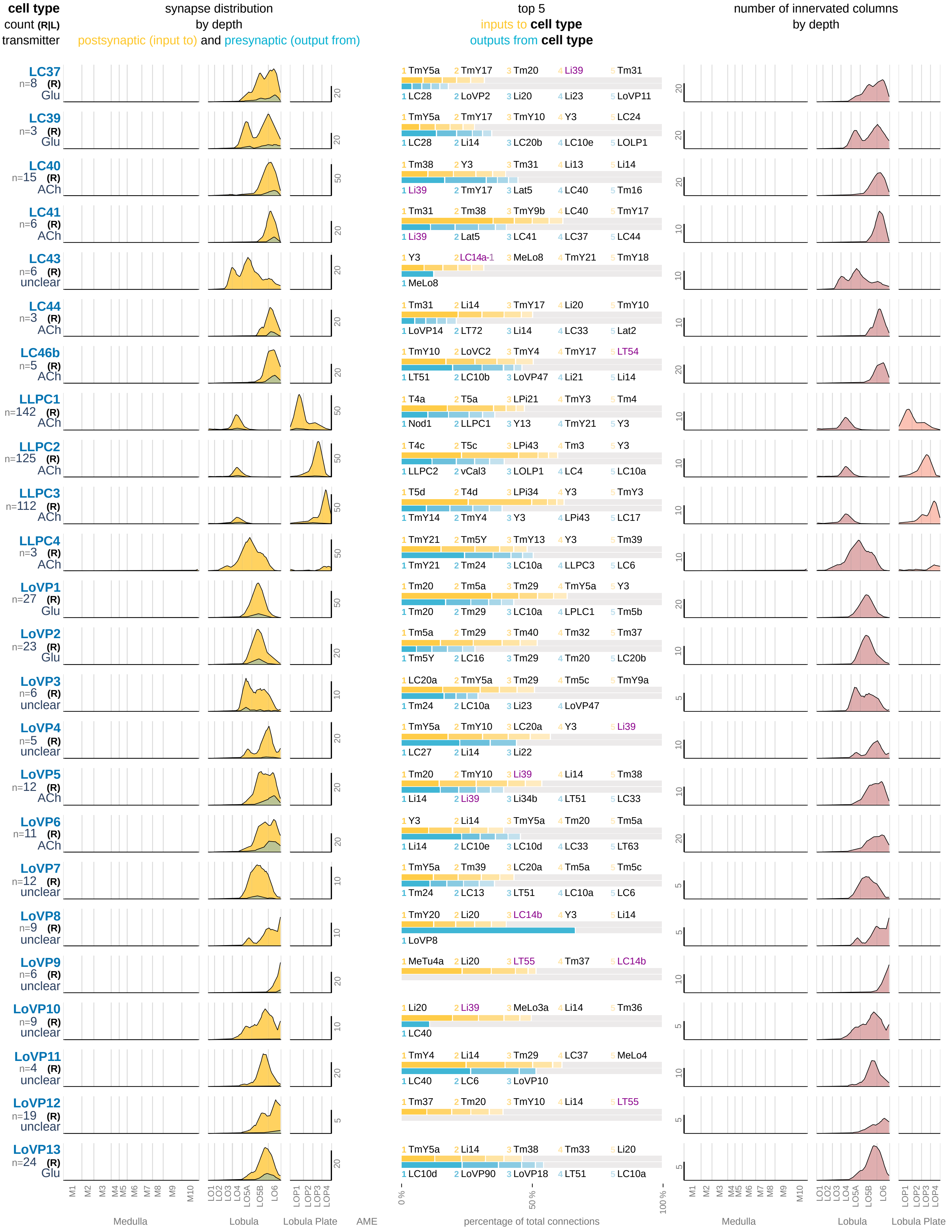

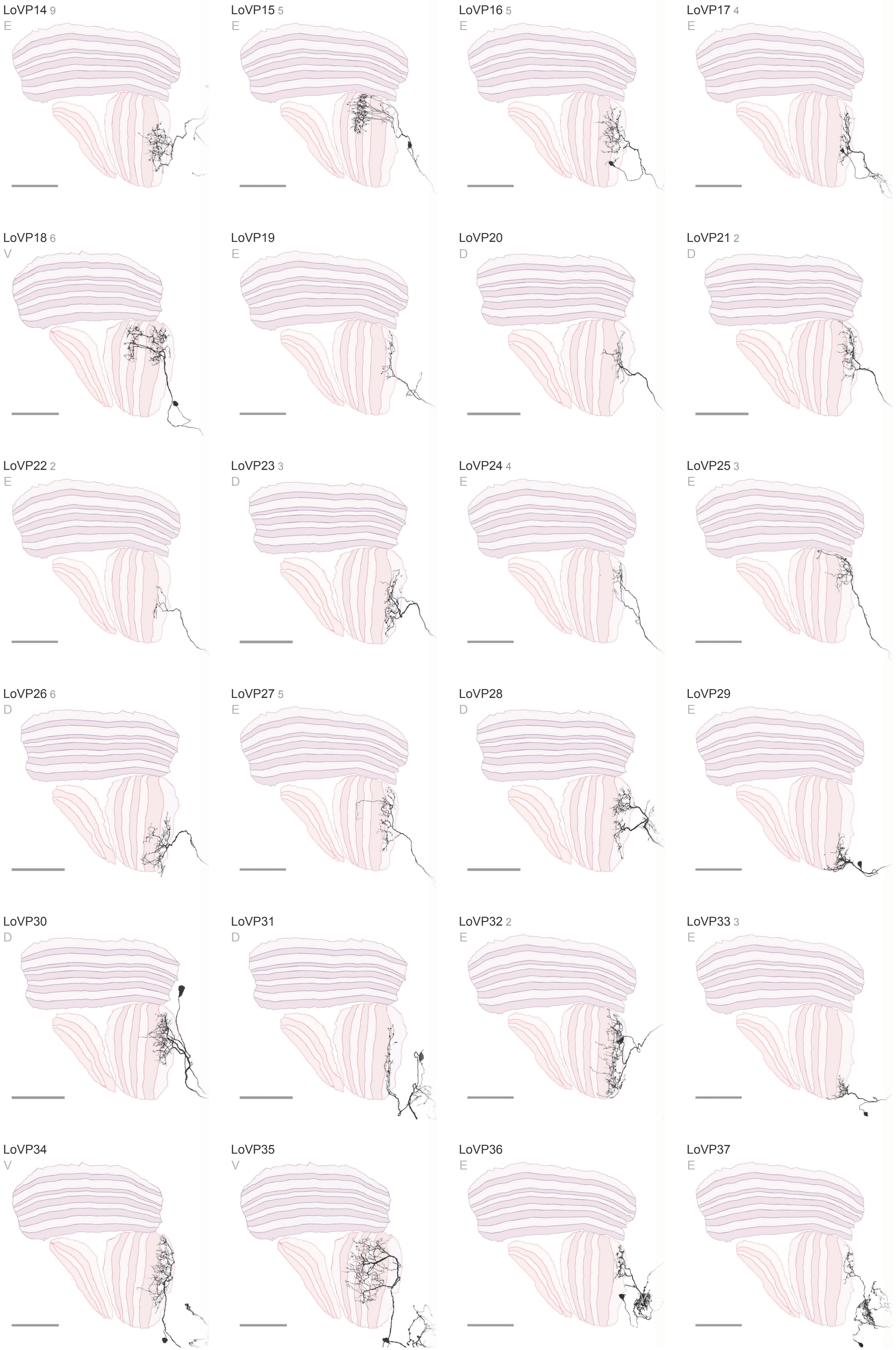

LoVP14 9
E
LoVP15 5
E
LoVP16 5
E
LoVP17 4
E
LoVP18 6
V
LoVP19
E
LoVP20
D
LoVP21 2
D
LoVP22 2
E
LoVP23 3
D
LoVP24 4
E
LoVP25 3
E
LoVP26 6
D
LoVP27 5
E
LoVP28
D
LoVP29
E
LoVP30
D
LoVP31
D
LoVP32 2
E
LoVP33 3
E
LoVP34
V
LoVP35
V
LoVP36
E
LoVP37
E

### Visual Projection Neurons 5 / 16

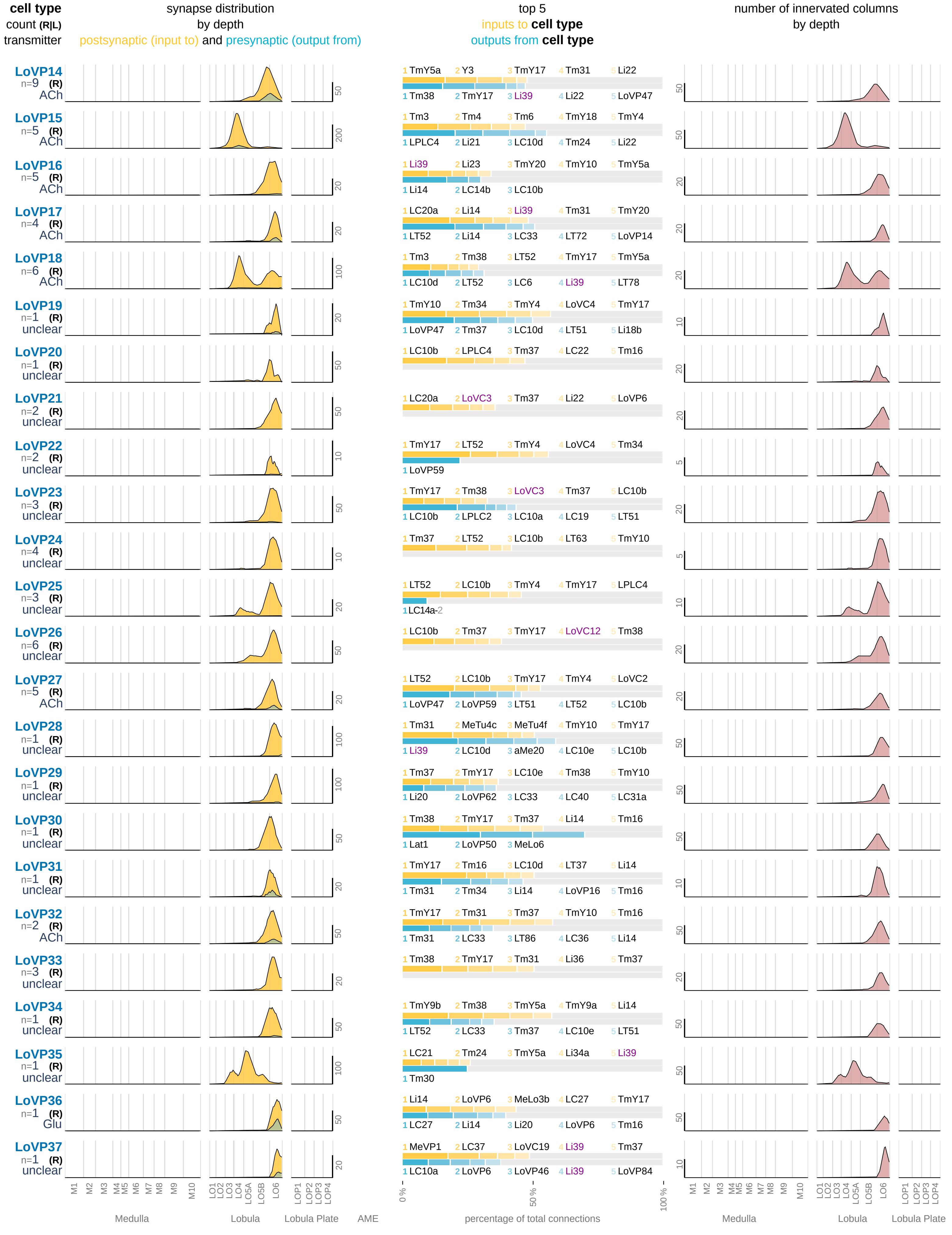

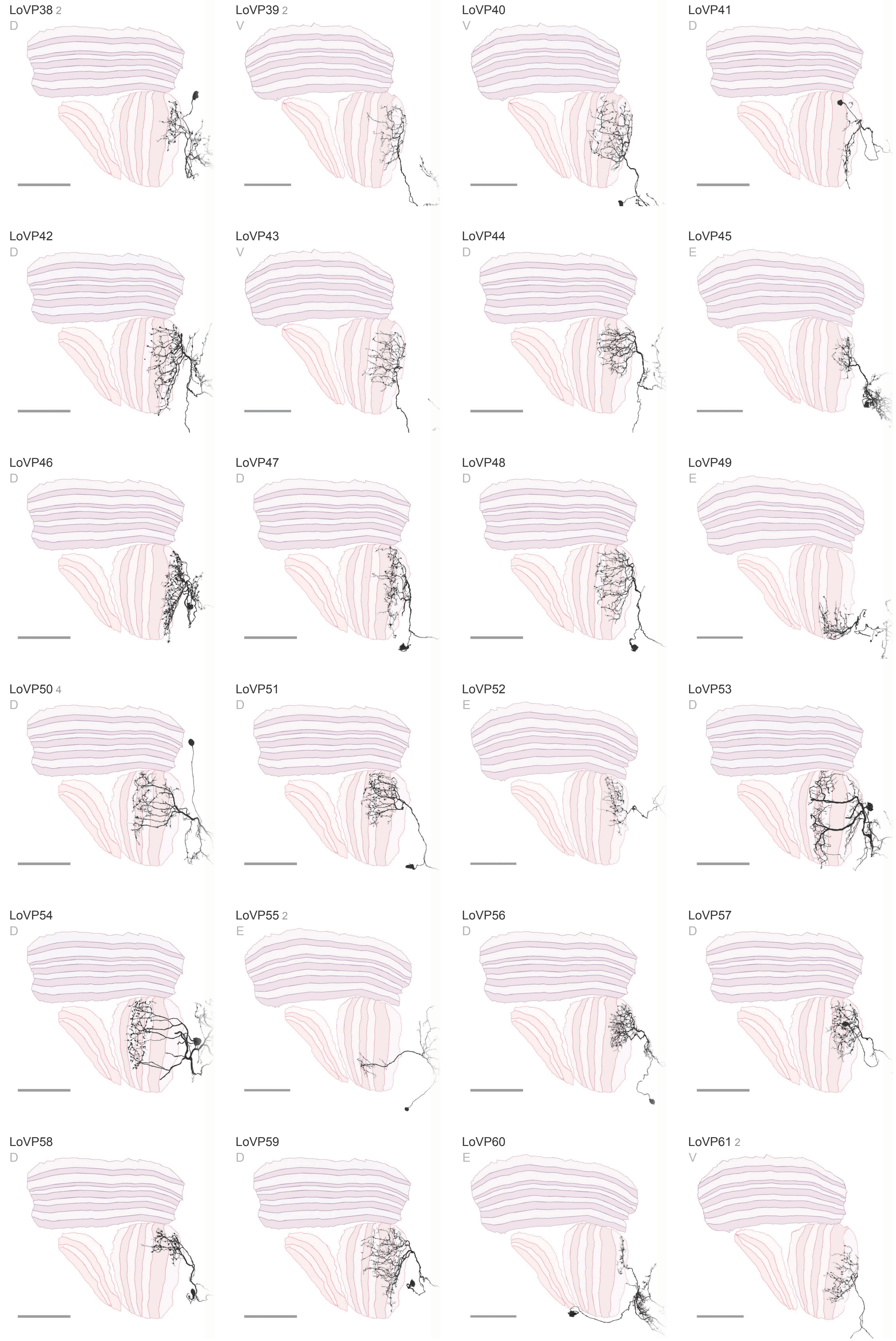

LoVP38 2
D
LoVP39 2
V
LoVP40
V
LoVP41
D
LoVP42
D
LoVP43
V
LoVP44
D
LoVP45
E
LoVP46
D
LoVP47
D
LoVP48
D
LoVP49
E
LoVP50 4
D
LoVP51
D
LoVP52
E
LoVP53
D
LoVP54
D
LoVP55 2
E
LoVP56
D
LoVP57
D
LoVP58
D
LoVP59
D
LoVP60
E
LoVP61 2
V

### Visual Projection Neurons 6 / 16

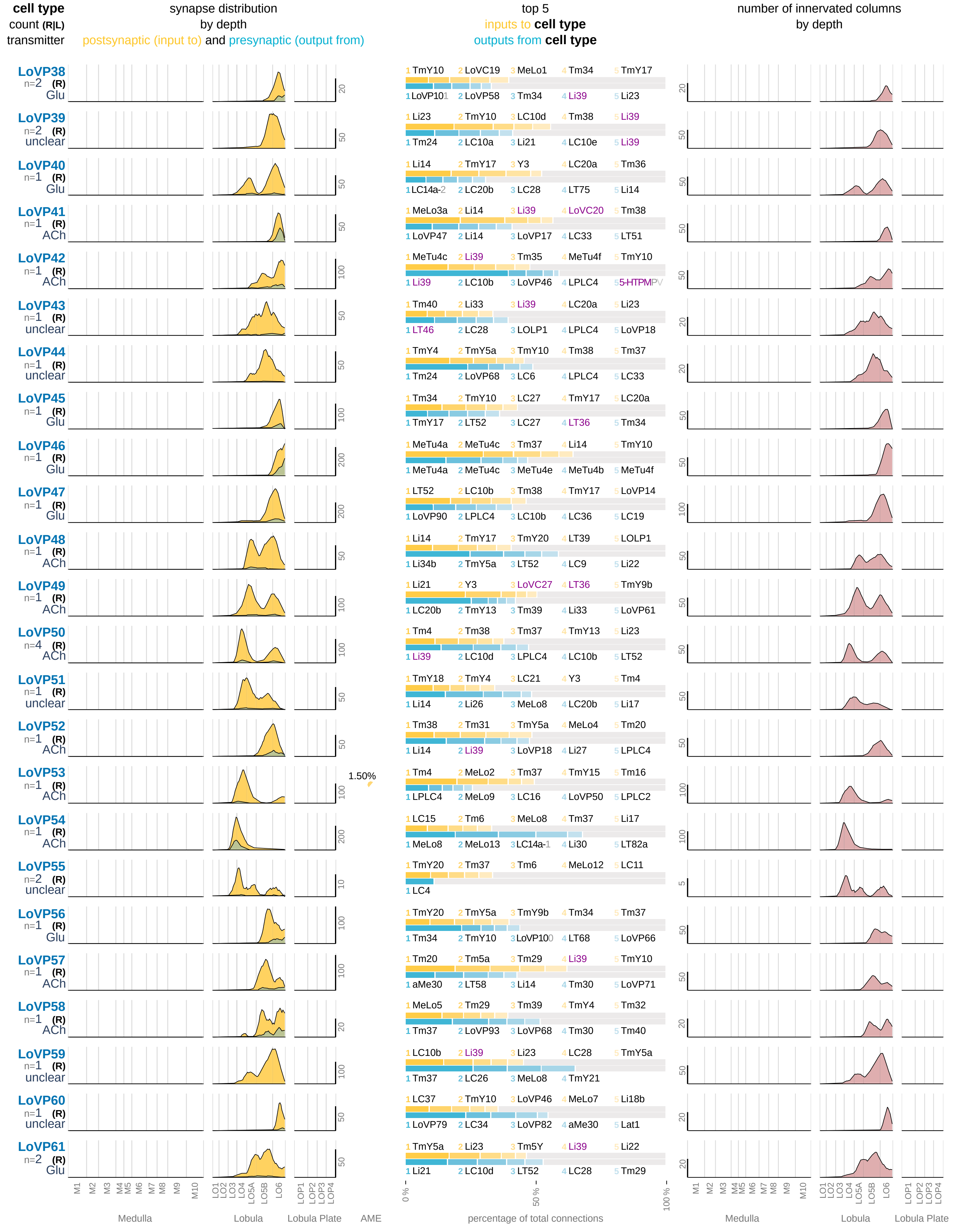

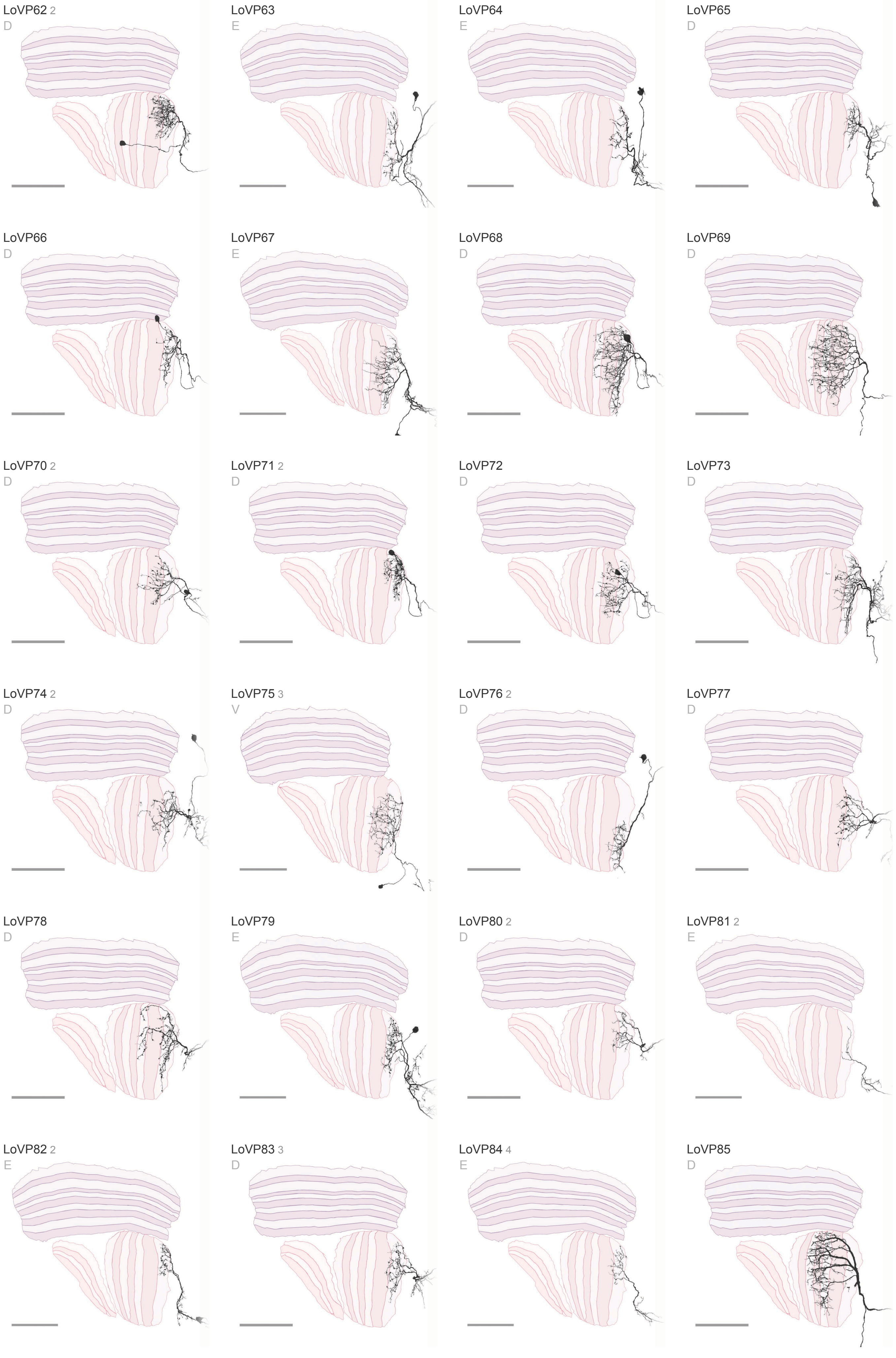
LoVP62 2
D
LoVP63
E
LoVP64
E
LoVP65
D
LoVP66
D
LoVP67
E
LoVP68
D
LoVP69
D
LoVP70 2
D
LoVP71 2
D
LoVP72
D
LoVP73
D
LoVP74 2
D
LoVP75 3
V
LoVP76 2
D
LoVP77
D
LoVP78
D
LoVP79
E
LoVP80 2
D
LoVP81 2
E
LoVP82 2
E
LoVP83 3
D
LoVP84 4
E
LoVP85
D

### Visual Projection Neurons 7 / 16

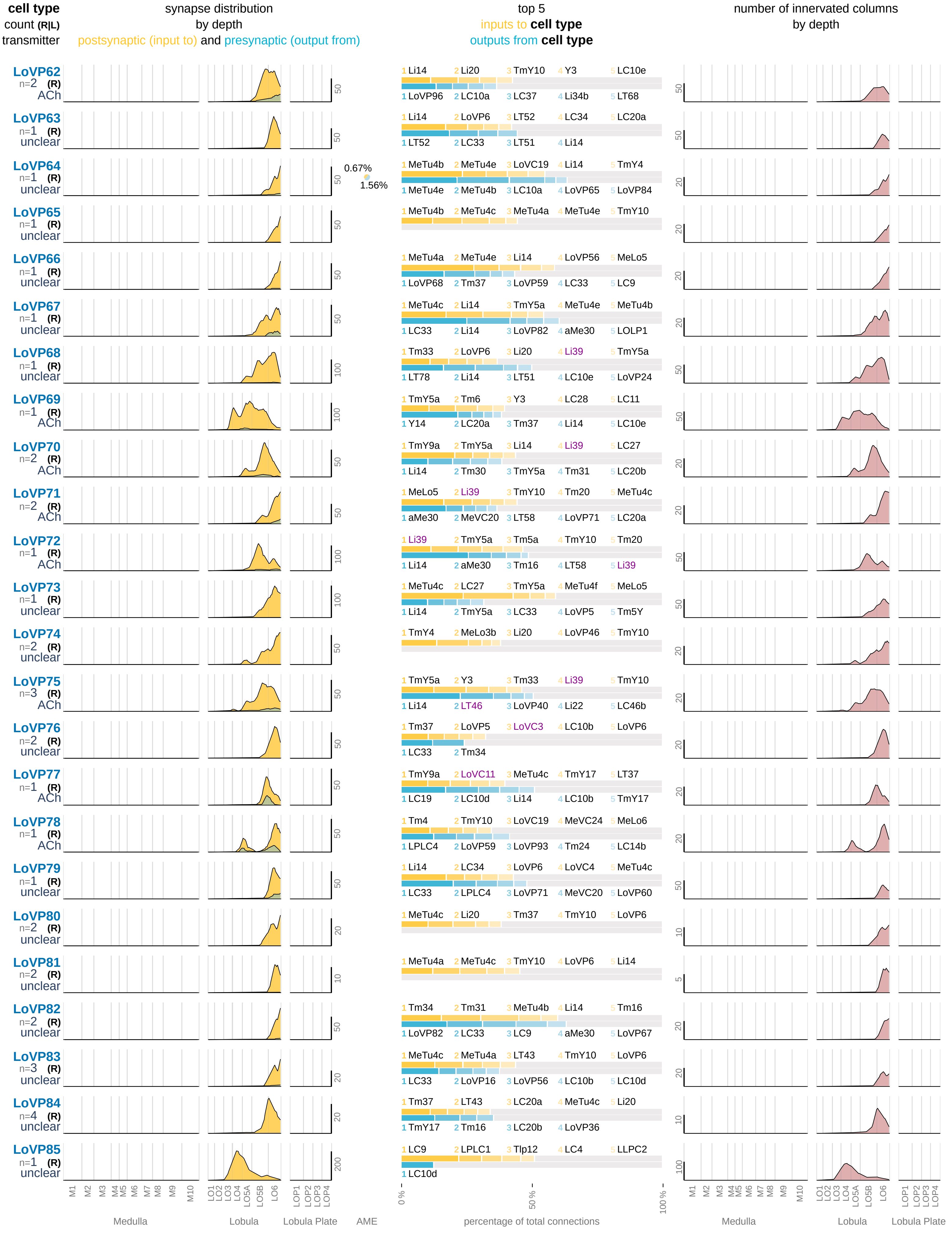

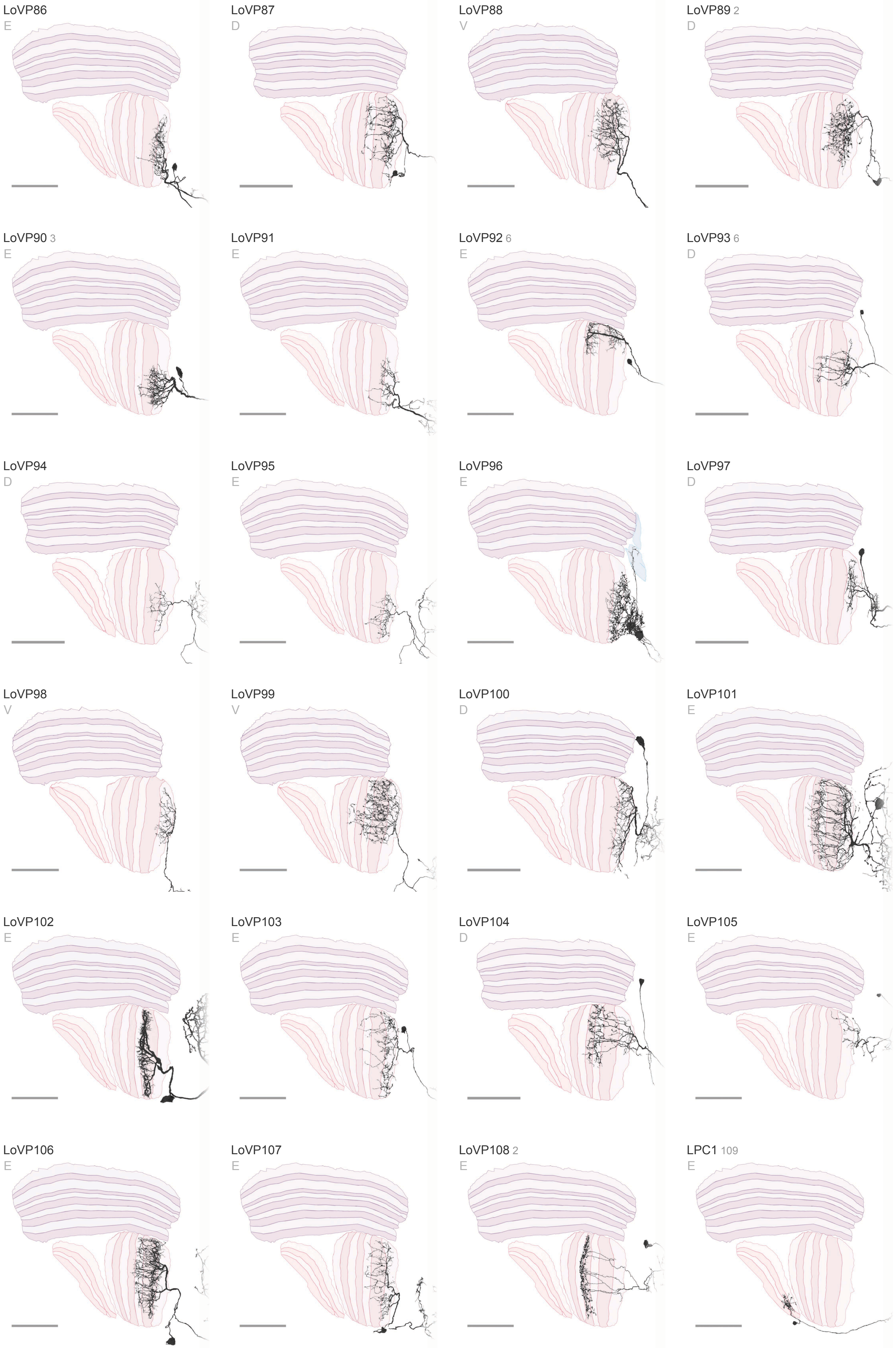

LoVP86
E
LoVP87
D
LoVP88
V
LoVP89 2
D
LoVP90 3
E
LoVP91
E
LoVP92 6
E
LoVP93 6
D
LoVP94
D
LoVP95
E
LoVP96
E
LoVP97
D
LoVP98
V
LoVP99
V
LoVP100
D
LoVP101
E
LoVP102
E
LoVP103
E
LoVP104
D
LoVP105
E
LoVP106
E
LoVP107
E
LoVP108 2
E
LPC1 109
E

### Visual Projection Neurons 8 / 16

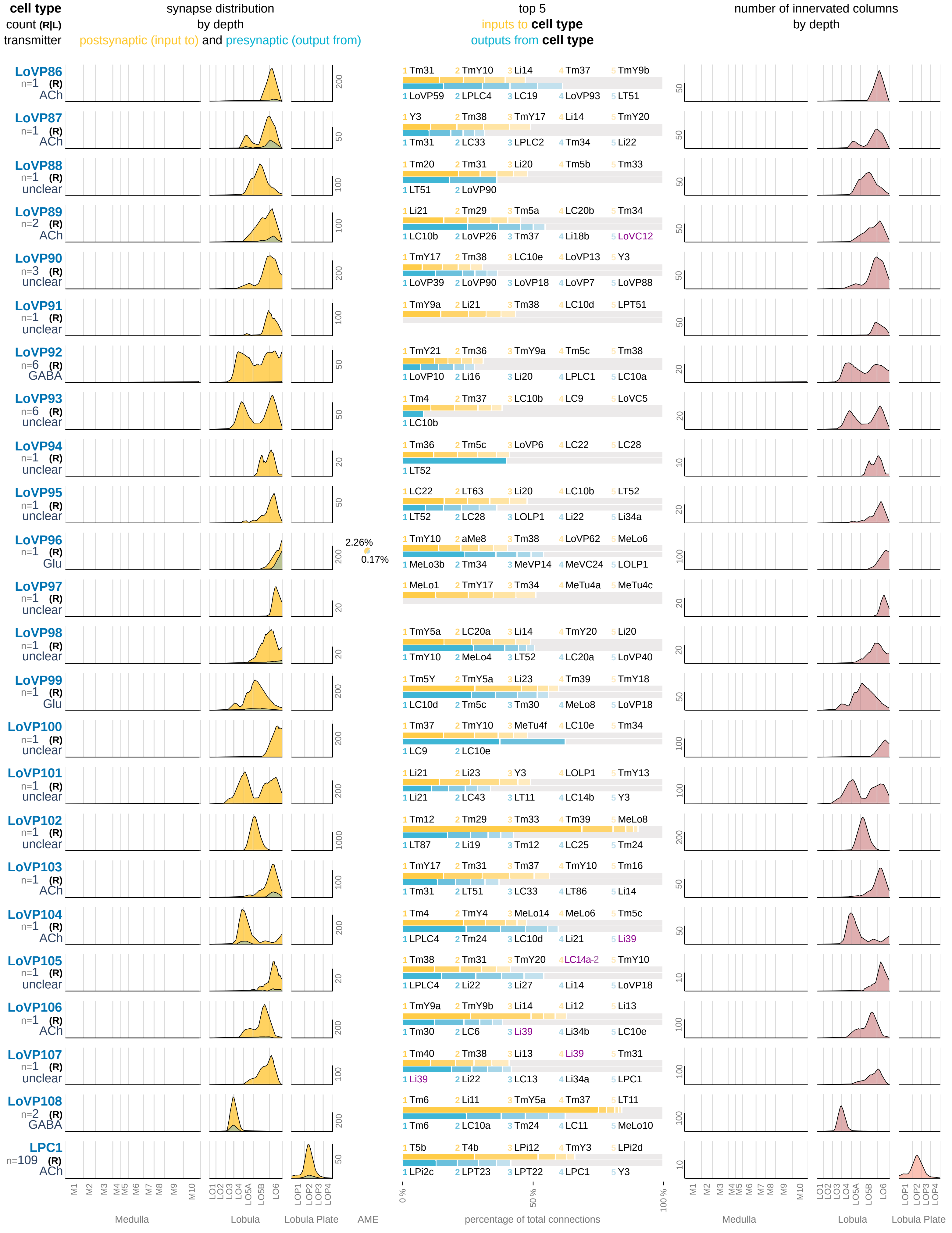

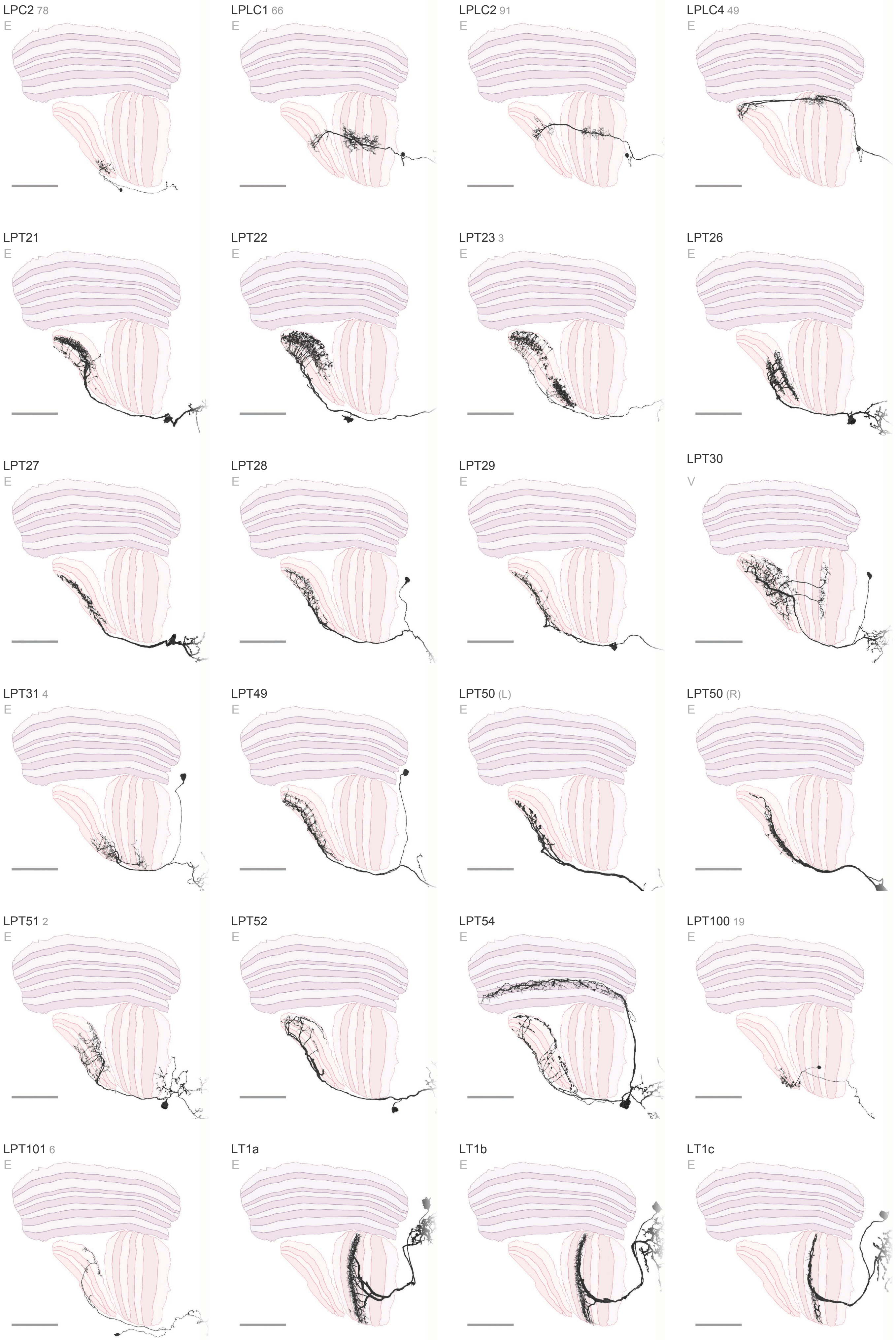

LPC2 78
E
LPLC1 66
E
LPLC2 91
E
LPLC4 49
E
LPT21
E
LPT22
E
LPT23 3
E
LPT26
E
LPT27
E
LPT28
E
LPT29
E
LPT30
V
LPT31 4
E
LPT49
E
LPT50 (L)
E
LPT50 (R)
E
LPT51 2
E
LPT52
E
LPT54
E
LPT100 19
E
LPT101 6
E
LT1a
E
LT1b
E
LT1c
E

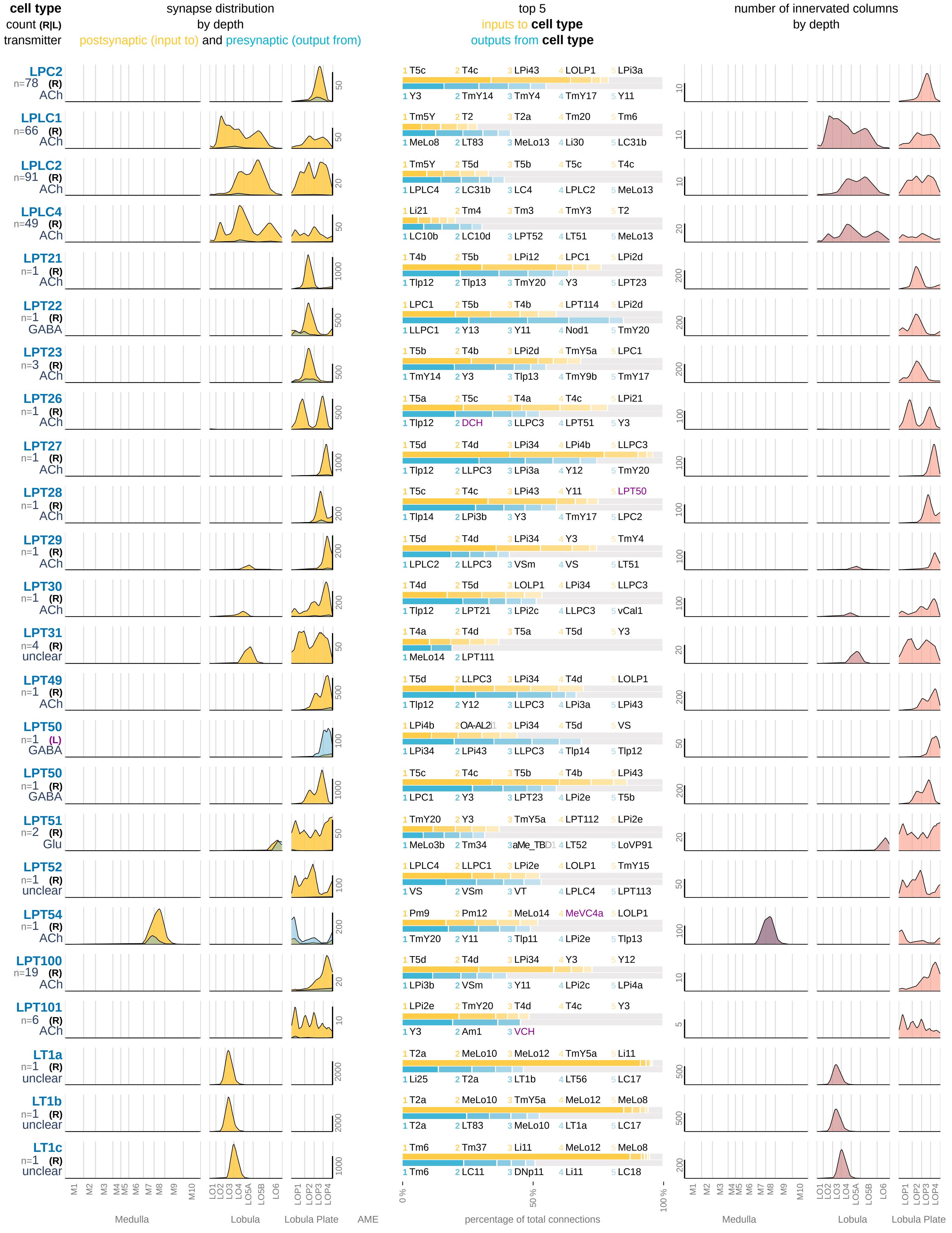
cell type
count (R|L)
transmitter
synapse distribution
by depth
postsynaptic (input to) and presynaptic (output from)
top 5
inputs to cell type
outputs from cell type
number of innervated columns
by depth
LPC2 n=78 (R) ACh
1 T5c 2 T4c 3 LPi43 4 LOLP1 5 LPi3a
1 Y3 2 TmY14 3 TmY4 4 TmY17 5 Y11
LPLC1 n=66 (R) ACh
1 Tm5Y 2 T2 3 T2a 4 Tm20 5 Tm6
1 MeLo8 2 LT83 3 MeLo13 4 Li30 5 LC31b
LPLC2 n=91 (R) ACh
1 Tm5Y 2 T5d 3 T5b 4 T5c 5 T4c
1 LPLC4 2 LC31b 3 LC4 4 LPLC2 5 MeLo13
LPLC4 n=49 (R) ACh
1 Li21 2 Tm4 3 Tm3 4 TmY3 5 T2
1 LC10b 2 LC10d 3 LPT52 4 LT51 5 MeLo13
LPT21 n=1 (R) ACh
1 T4b 2 T5b 3 LPi12 4 LPC1 5 LPi2d
1 Tlp12 2 Tlp13 3 TmY20 4 Y3 5 LPT23
LPT22 n=1 (R) GABA
1 LPC1 2 T5b 3 T4b 4 LPT114 5 LPi2d
1 LLPC1 2 Y13 3 Y11 4 Nod1 5 TmY20
LPT23 n=3 (R) ACh
1 T5b 2 T4b 3 LPi2d 4 TmY5a 5 LPC1
1 TmY14 2 Y3 3 Tlp13 4 TmY9b 5 TmY17
LPT26 n=1 (R) ACh
1 T5a 2 T5c 3 T4a 4 T4c 5 LPi21
1 Tlp12 2 DCH 3 LLPC3 4 LPT51 5 Y3
LPT27 n=1 (R) ACh
1 T5d 2 T4d 3 LPi34 4 LPi4b 5 LLPC3
1 Tlp12 2 LLPC3 3 LPi3a 4 Y12 5 TmY20
LPT28 n=1 (R) ACh
1 T5c 2 T4c 3 LPi43 4 Y11 5 LPT50
1 Tlp14 2 LPi3b 3 Y3 4 TmY17 5 LPC2
LPT29 n=1 (R) ACh
1 T5d 2 T4d 3 LPi34 4 Y3 5 TmY4
1 LPLC2 2 LLPC3 3 VSm 4 VS 5 LT51
LPT30 n=1 (R) ACh
1 T4d 2 T5d 3 LOLP1 4 LPi34 5 LLPC3
1 Tlp12 2 LPT21 3 LPi2c 4 LLPC3 5 vCal1
LPT31 n=4 (R) unclear
1 T4a 2 T4d 3 T5a 4 T5d 5 Y3
1 MeLo14 2 LPT111
LPT49 n=1 (R) ACh
1 T5d 2 LLPC3 3 LPi34 4 T4d 5 LOLP1
1 Tlp12 2 Y12 3 LLPC3 4 LPi3a 5 LPi43
LPT50 n=1 (L) GABA
1 LPi4b 2 OA-AL2i1 3 LPi34 4 T5d 5 VS
1 LPi34 2 LPi43 3 LLPC3 4 Tlp14 5 Tlp12
LPT50 n=1 (R) GABA
1 T5c 2 T4c 3 T5b 4 T4b 5 LPi43
1 LPC1 2 Y3 3 LPT23 4 LPi2e 5 T5b
LPT51 n=2 (R) Glu
1 TmY20 2 Y3 3 TmY5a 4 LPT112 5 LPi2e
1 MeLo3b 2 Tm34 3 aMe_TBD1 4 LT52 5 LoVP91
LPT52 n=1 (R) unclear
1 LPLC4 2 LPLC1 3 LPi2e 4 LOLP1 5 TmY15
1 VS 2 VSm 3 VT 4 LPLC4 5 LPT113
LPT54 n=1 (R) ACh
1 Pm9 2 Pm12 3 MeLo14 4 MeVC4a 5 LOLP1
1 TmY20 2 Y11 3 Tlp11 4 LPi2e 5 Tlp13
LPT100 n=19 (R) ACh
1 T5d 2 T4d 3 LPi34 4 Y3 5 Y12
1 LPi3b 2 VSm 3 Y11 4 LPi2c 5 LPi4a
LPT101 n=6 (R) ACh
1 LPi2e 2 TmY20 3 T4d 4 T4c 5 Y3
1 Y3 2 Am1 3 VCH
LT1a n=1 (R) unclear
1 T2a 2 MeLo10 3 MeLo12 4 TmY5a 5 Li11
1 Li25 2 T2a 3 LT1b 4 LT56 5 LC17
LT1b n=1 (R) unclear
1 T2a 2 MeLo10 3 TmY5a 4 MeLo12 5 MeLo8
1 T2a 2 LT83 3 MeLo10 4 LT1a 5 LC17
LT1c n=1 (R) unclear
1 Tm6 2 Tm37 3 Li11 4 MeLo12 5 MeLo8
1 Tm6 2 LC11 3 DNp11 4 Li11 5 LC18
M1 M2 M3 M4 M5 M6 M7 M8 M9 M10
LO1 LO2 LO3 LO4 LO5A LO5B LO6
LOP1 LOP2 LOP3 LOP4
Medulla
Lobula
Lobula Plate
AME
0 % 50 % 100 %
percentage of total connections

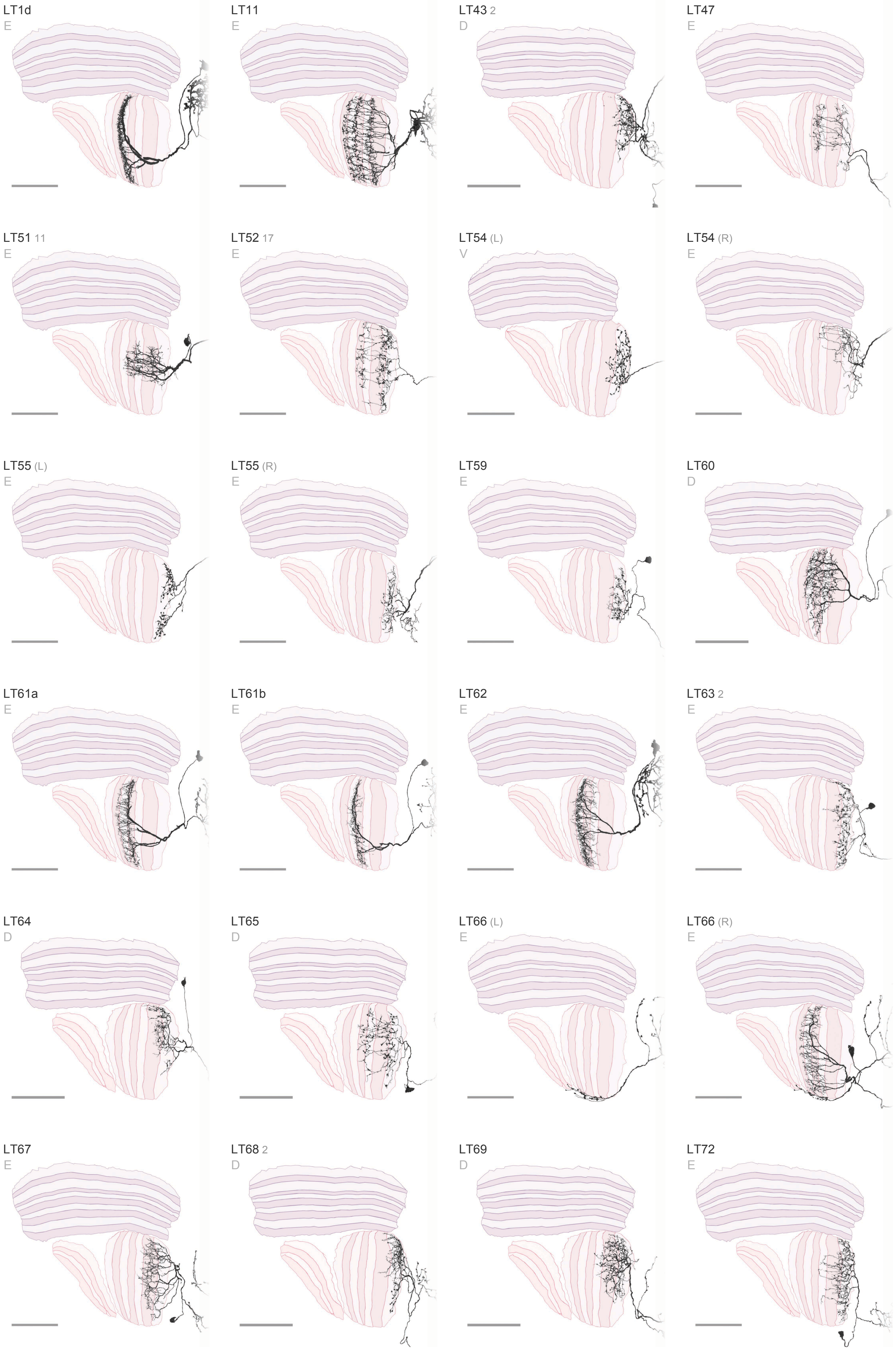

LT1d
E
LT11
E
LT43 2
D
LT47
E
LT51 11
E
LT52 17
E
LT54 (L)
V
LT54 (R)
E
LT55 (L)
E
LT55 (R)
E
LT59
E
LT60
D
LT61a
E
LT61b
E
LT62
E
LT63 2
E
LT64
D
LT65
D
LT66 (L)
E
LT66 (R)
E
LT67
E
LT68 2
D
LT69
D
LT72
E

### Visual Projection Neurons 10 / 16

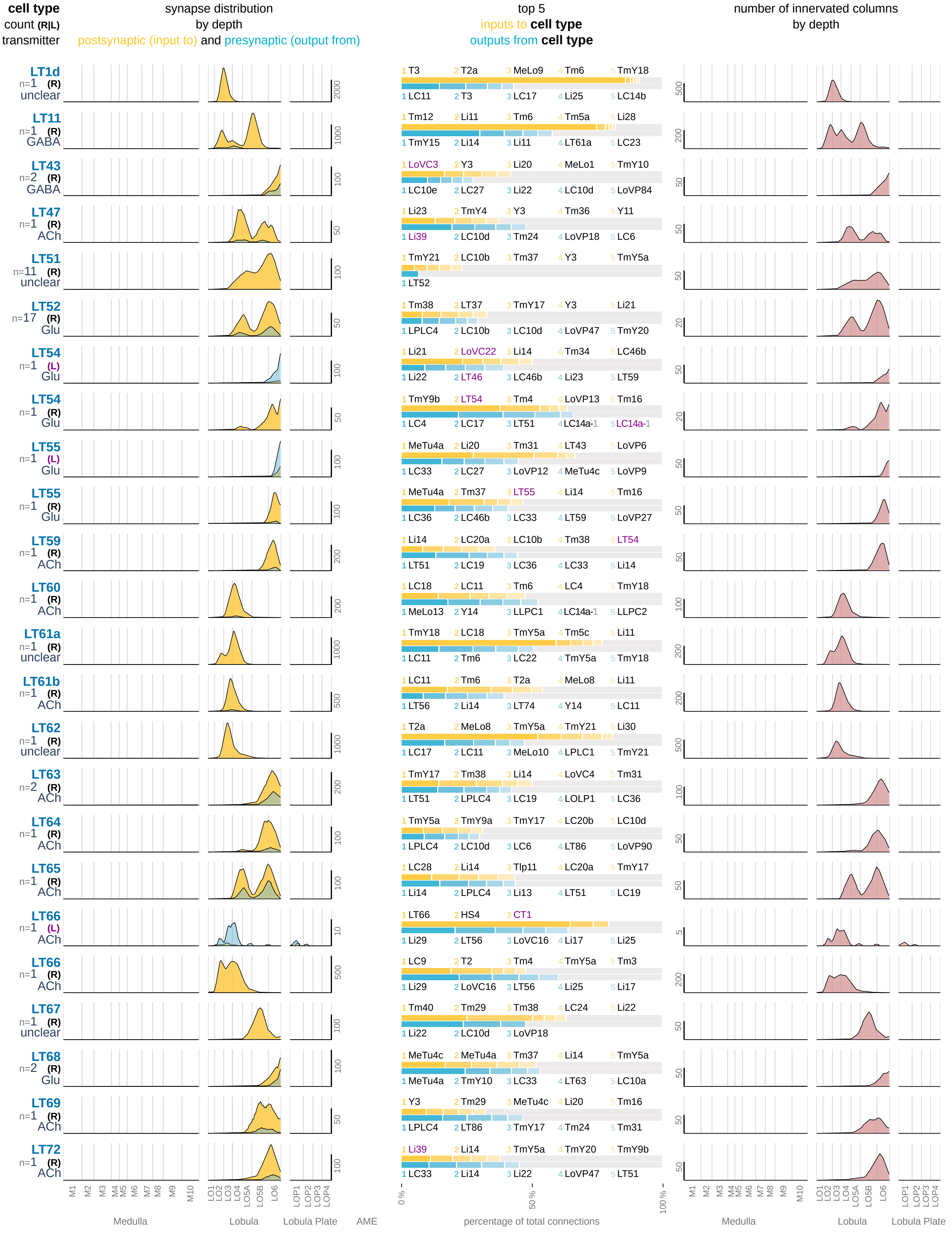

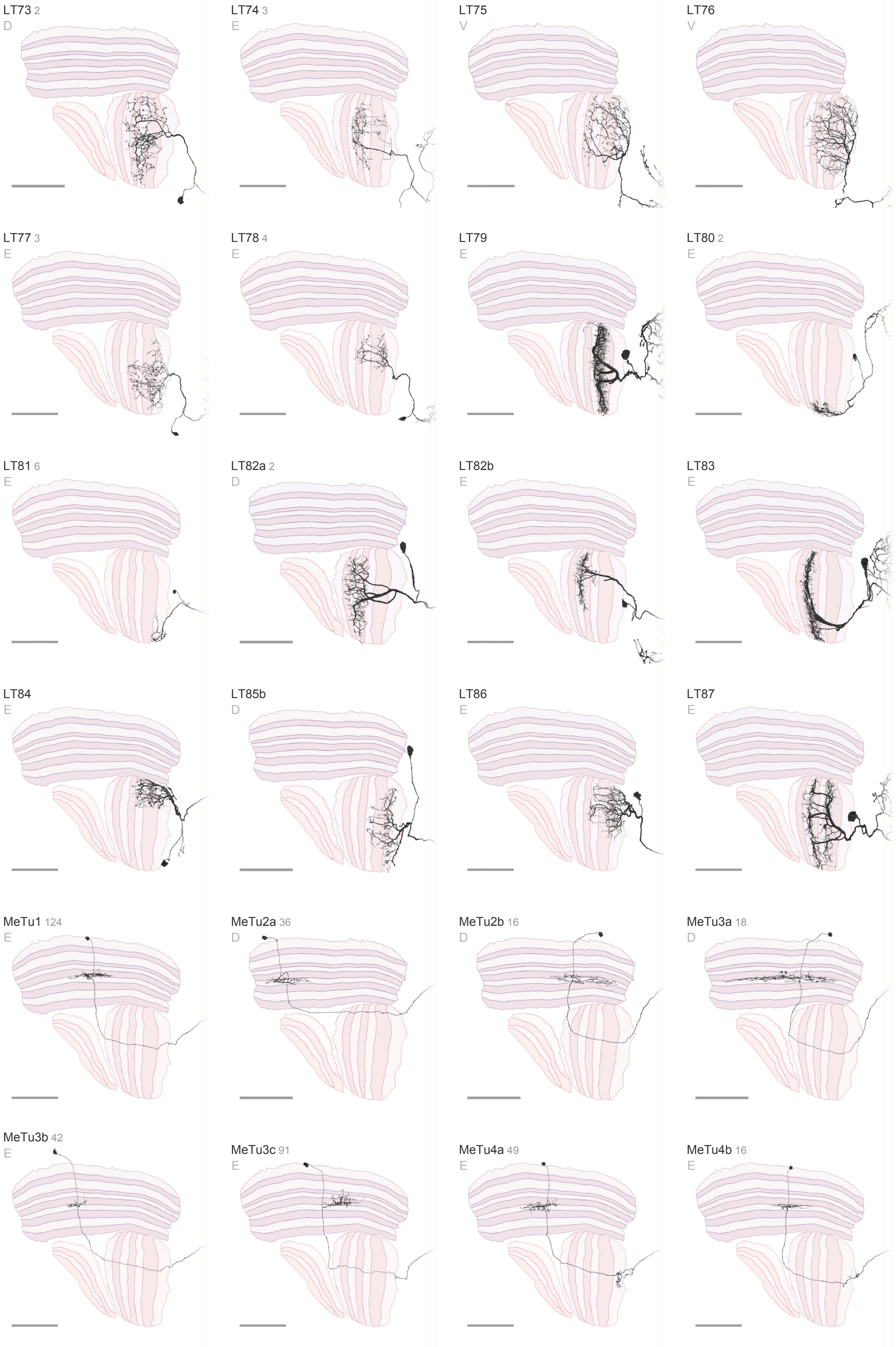
LT73 2
D
LT74 3
E
LT75
V
LT76
V
LT77 3
E
LT78 4
E
LT79
E
LT80 2
E
LT81 6
E
LT82a 2
D
LT82b
E
LT83
E
LT84
E
LT85b
D
LT86
E
LT87
E
MeTu1 124
E
MeTu2a 36
D
MeTu2b 16
D
MeTu3a 18
D
MeTu3b 42
E
MeTu3c 91
E
MeTu4a 49
E
MeTu4b 16
E

### Visual Projection Neurons 11 / 16

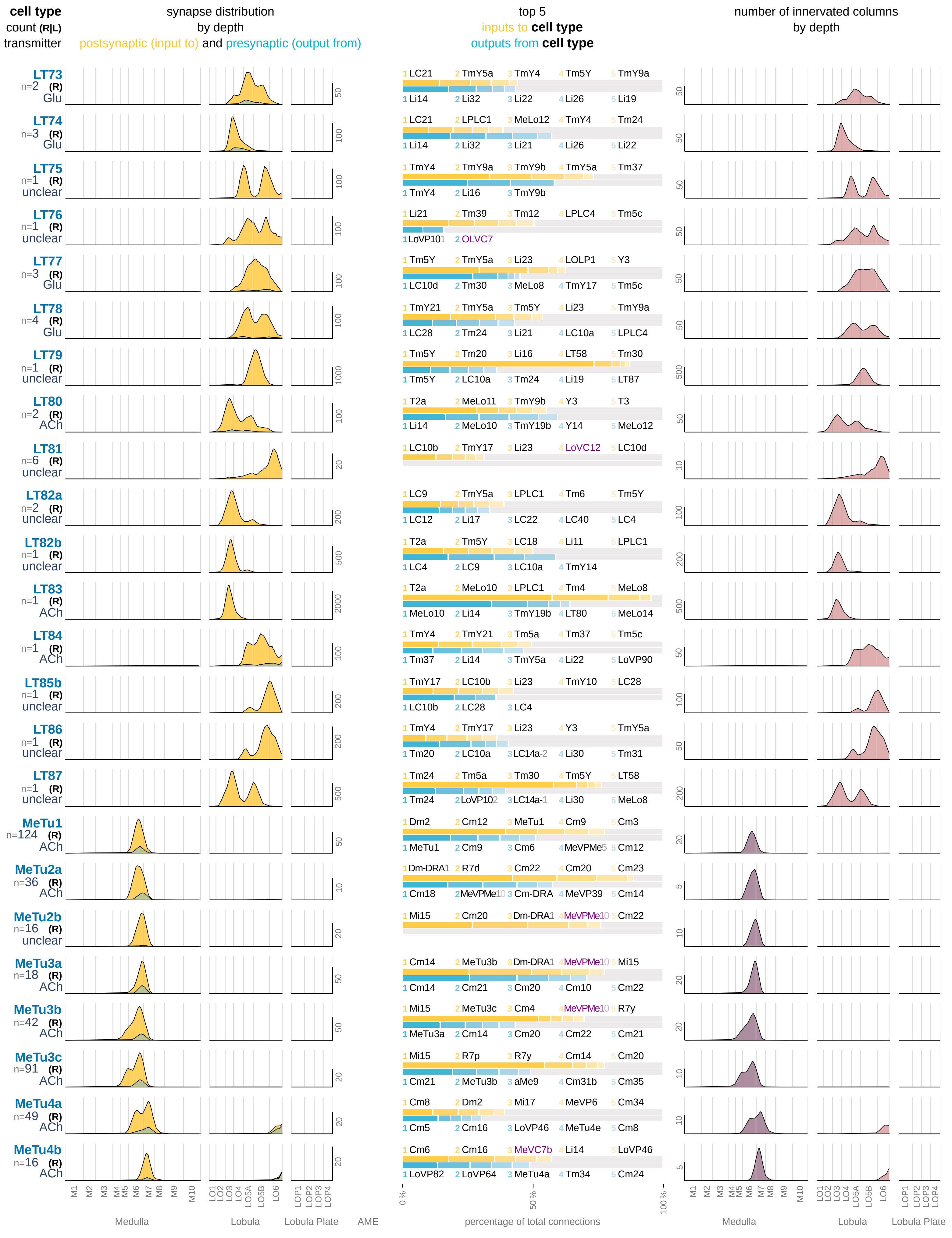

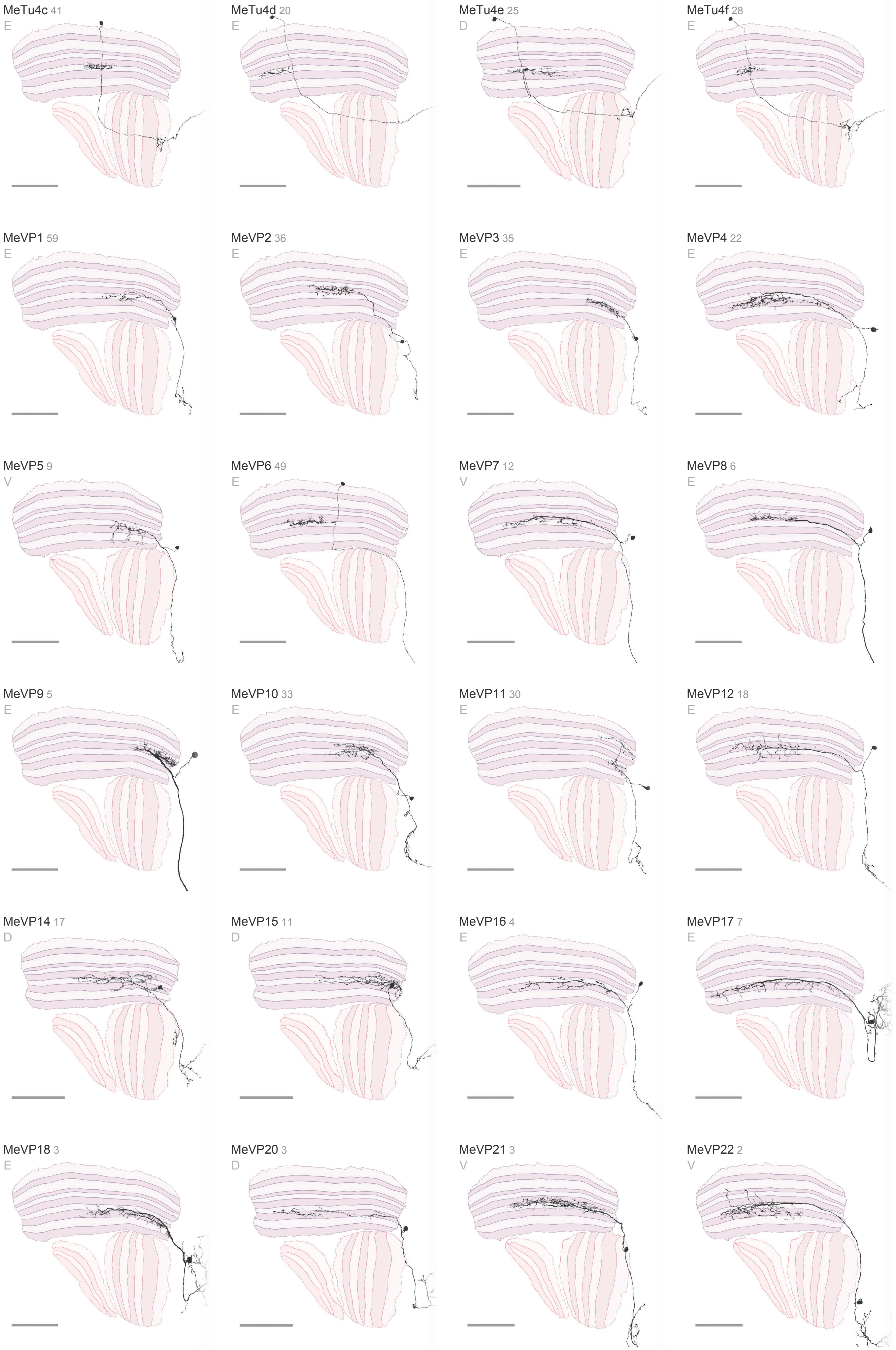

MeTu4c 41
E
MeTu4d 20
E
MeTu4e 25
D
MeTu4f 28
E
MeVP1 59
E
MeVP2 36
E
MeVP3 35
E
MeVP4 22
E
MeVP5 9
V
MeVP6 49
E
MeVP7 12
V
MeVP8 6
E
MeVP9 5
E
MeVP10 33
E
MeVP11 30
E
MeVP12 18
E
MeVP14 17
D
MeVP15 11
D
MeVP16 4
E
MeVP17 7
E
MeVP18 3
E
MeVP20 3
D
MeVP21 3
V
MeVP22 2
V

**Visual Projection Neurons** 12 / 16

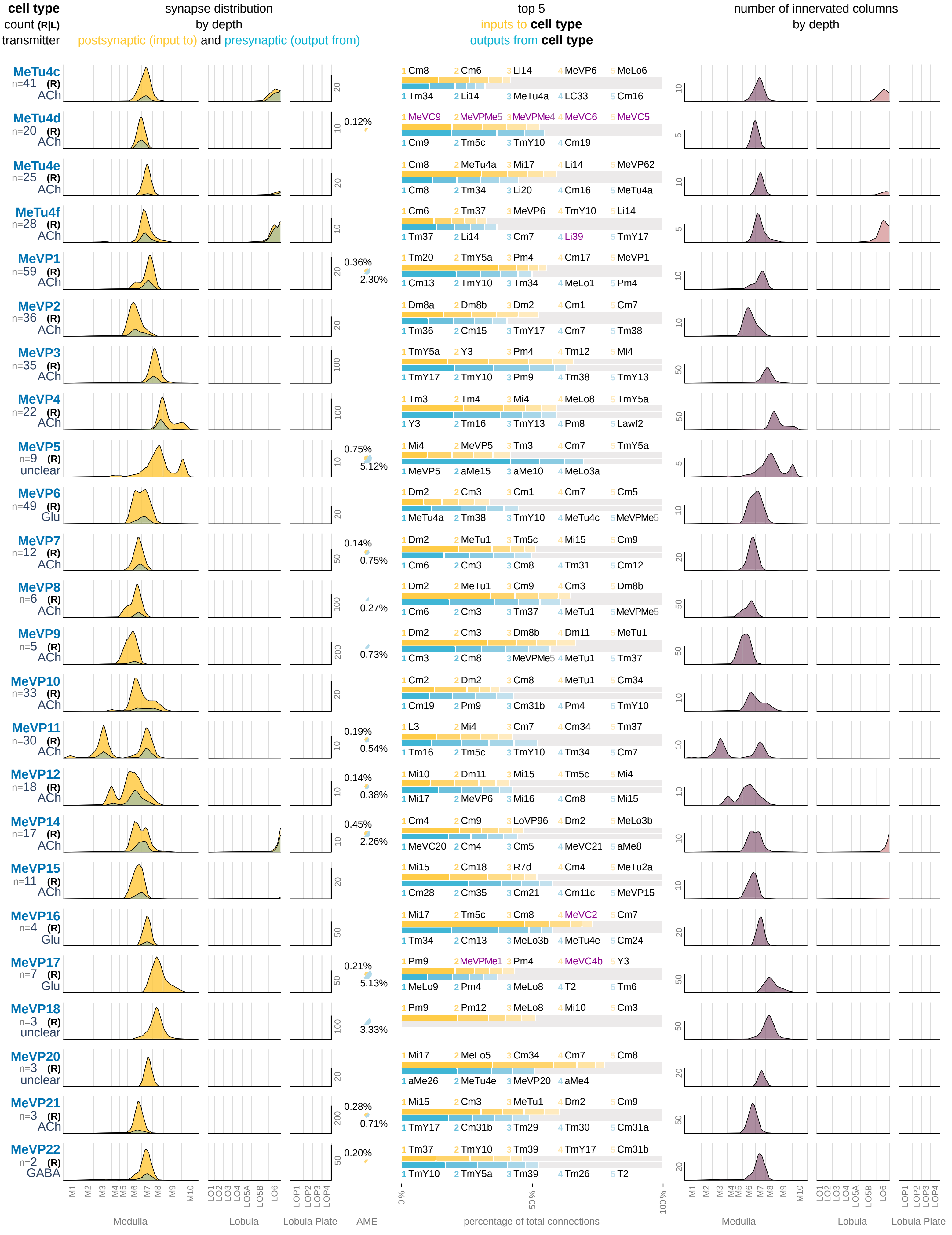

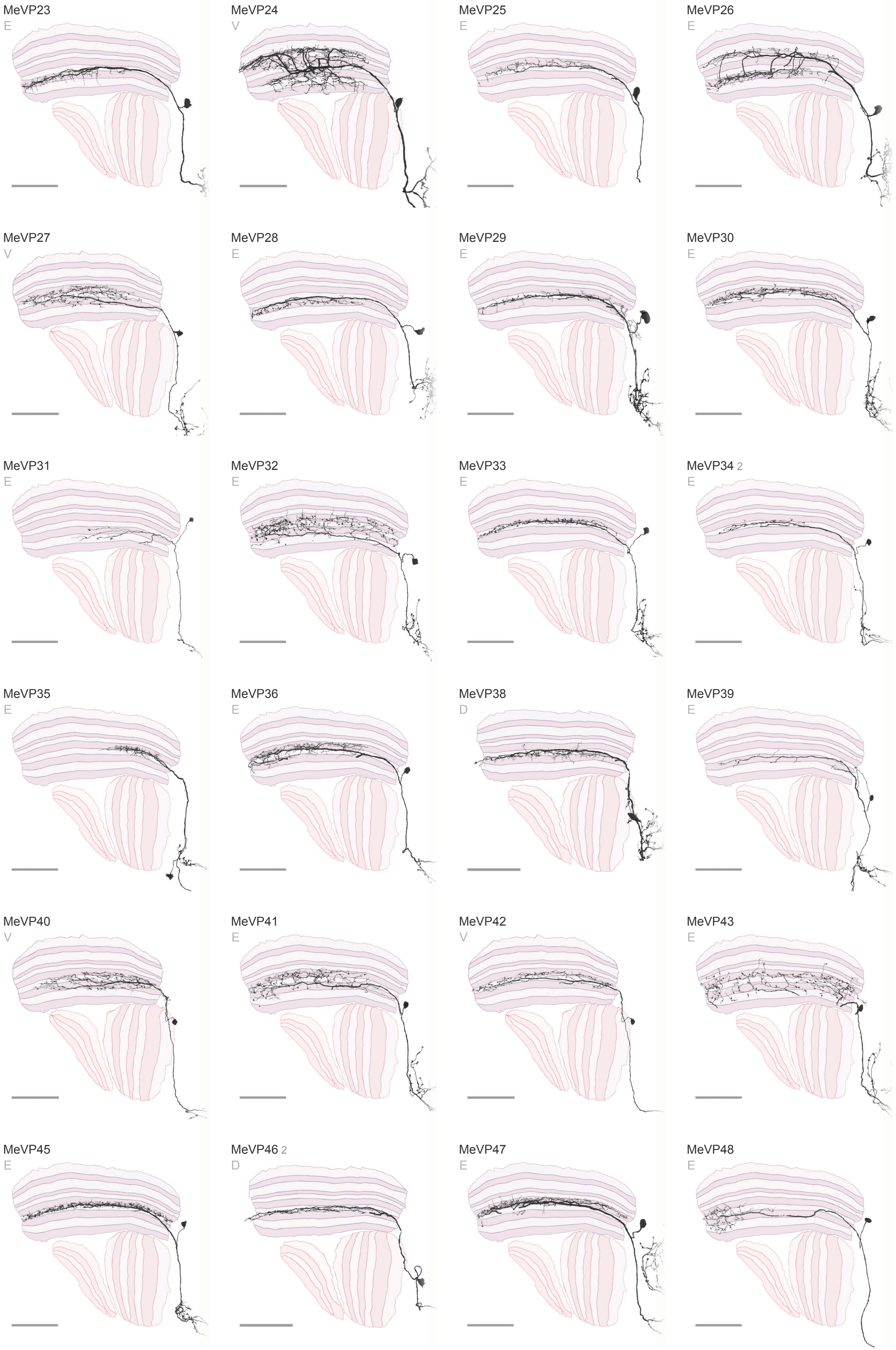
MeVP23
E
MeVP24
V
MeVP25
E
MeVP26
E
MeVP27
V
MeVP28
E
MeVP29
E
MeVP30
E
MeVP31
E
MeVP32
E
MeVP33
E
MeVP34 2
E
MeVP35
E
MeVP36
E
MeVP38
D
MeVP39
E
MeVP40
V
MeVP41
E
MeVP42
V
MeVP43
E
MeVP45
E
MeVP46 2
D
MeVP47
E
MeVP48
E

### Visual Projection Neurons 13 / 16

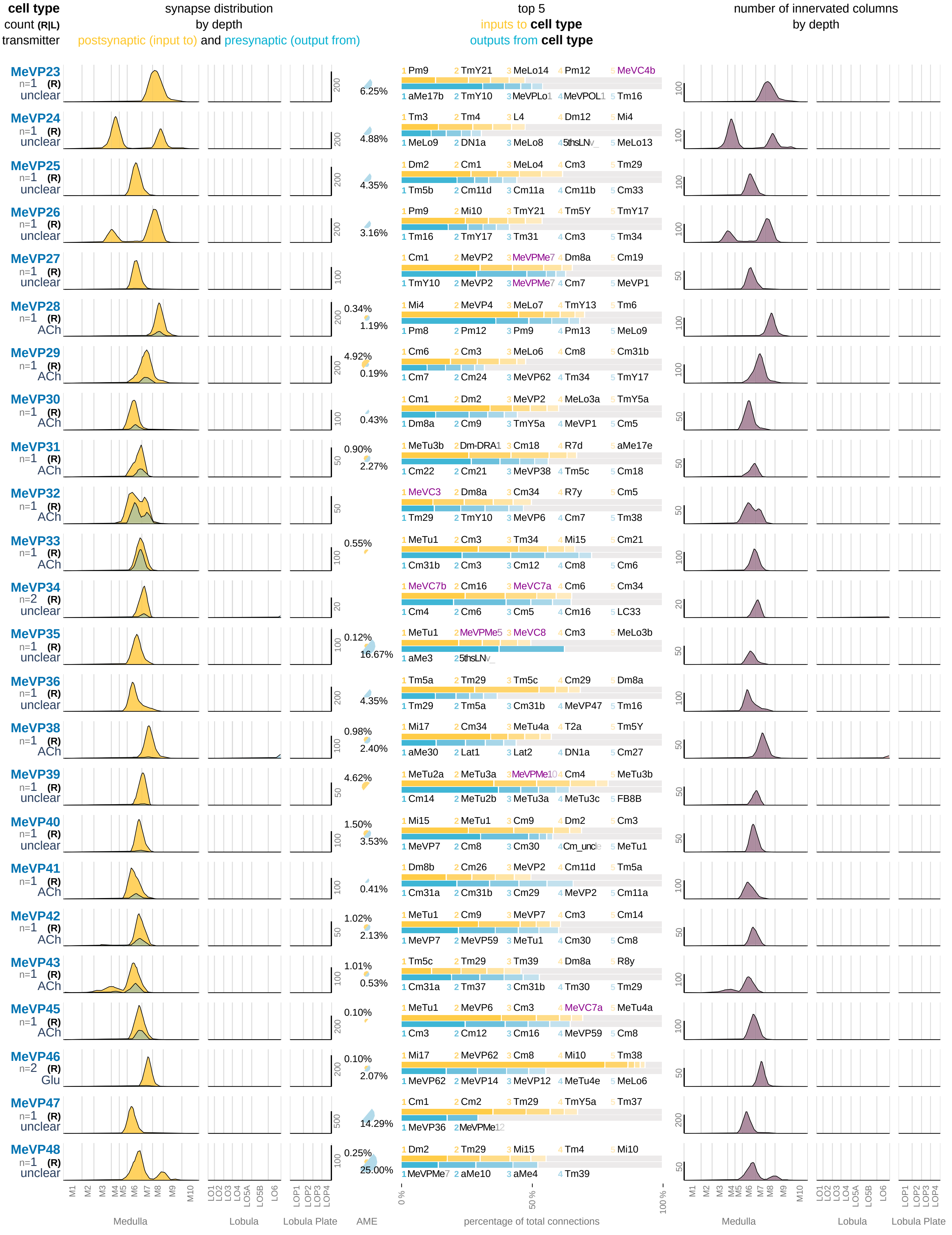

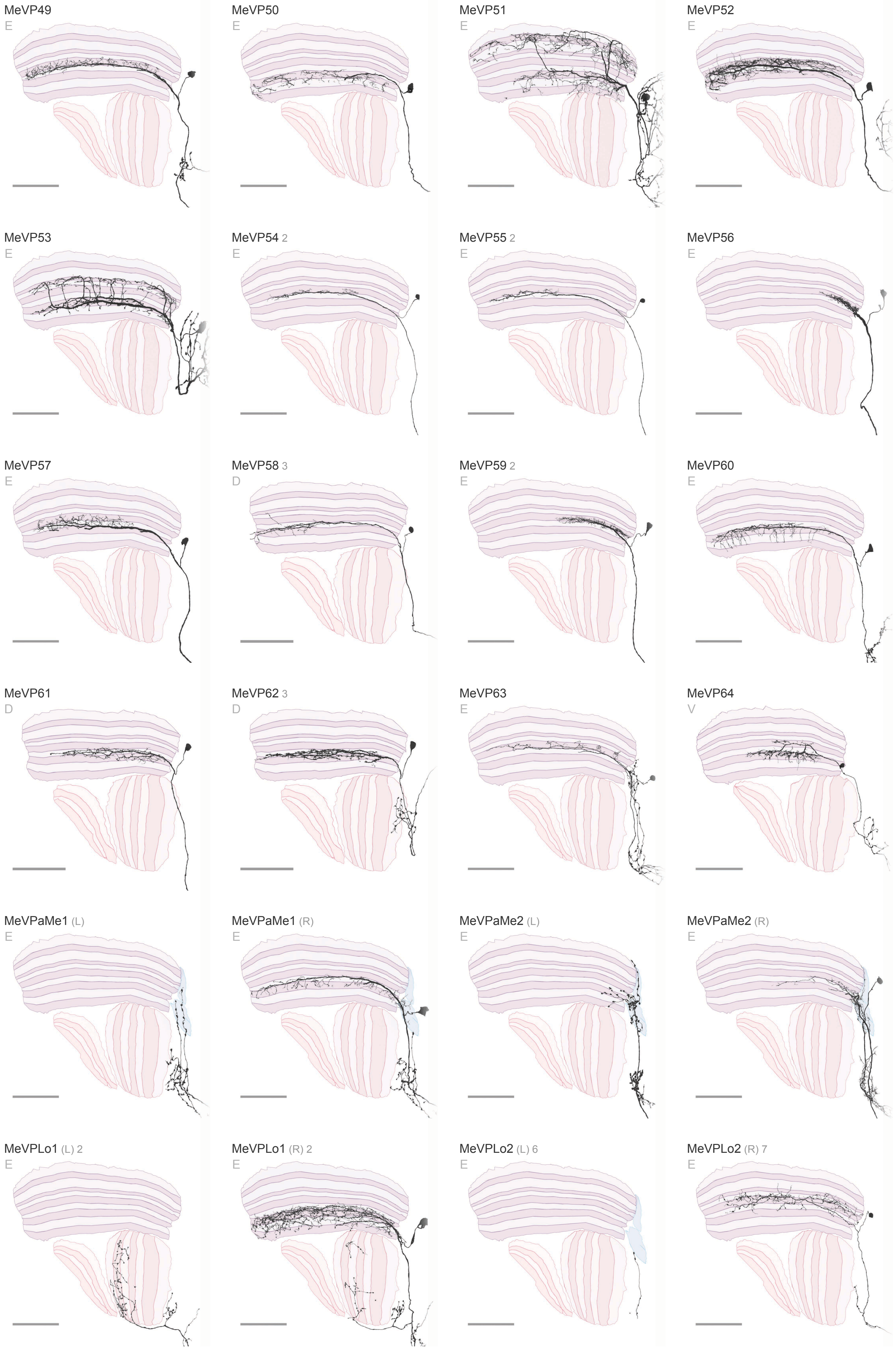

MeVP49
E
MeVP50
E
MeVP51
E
MeVP52
E
MeVP53
E
MeVP54 2
E
MeVP55 2
E
MeVP56
E
MeVP57
E
MeVP58 3
D
MeVP59 2
E
MeVP60
E
MeVP61
D
MeVP62 3
D
MeVP63
E
MeVP64
V
MeVPaMe1 (L)
E
MeVPaMe1 (R)
E
MeVPaMe2 (L)
E
MeVPaMe2 (R)
E
MeVPLo1 (L) 2
E
MeVPLo1 (R) 2
E
MeVPLo2 (L) 6
E
MeVPLo2 (R) 7
E

### Visual Projection Neurons 14 / 16

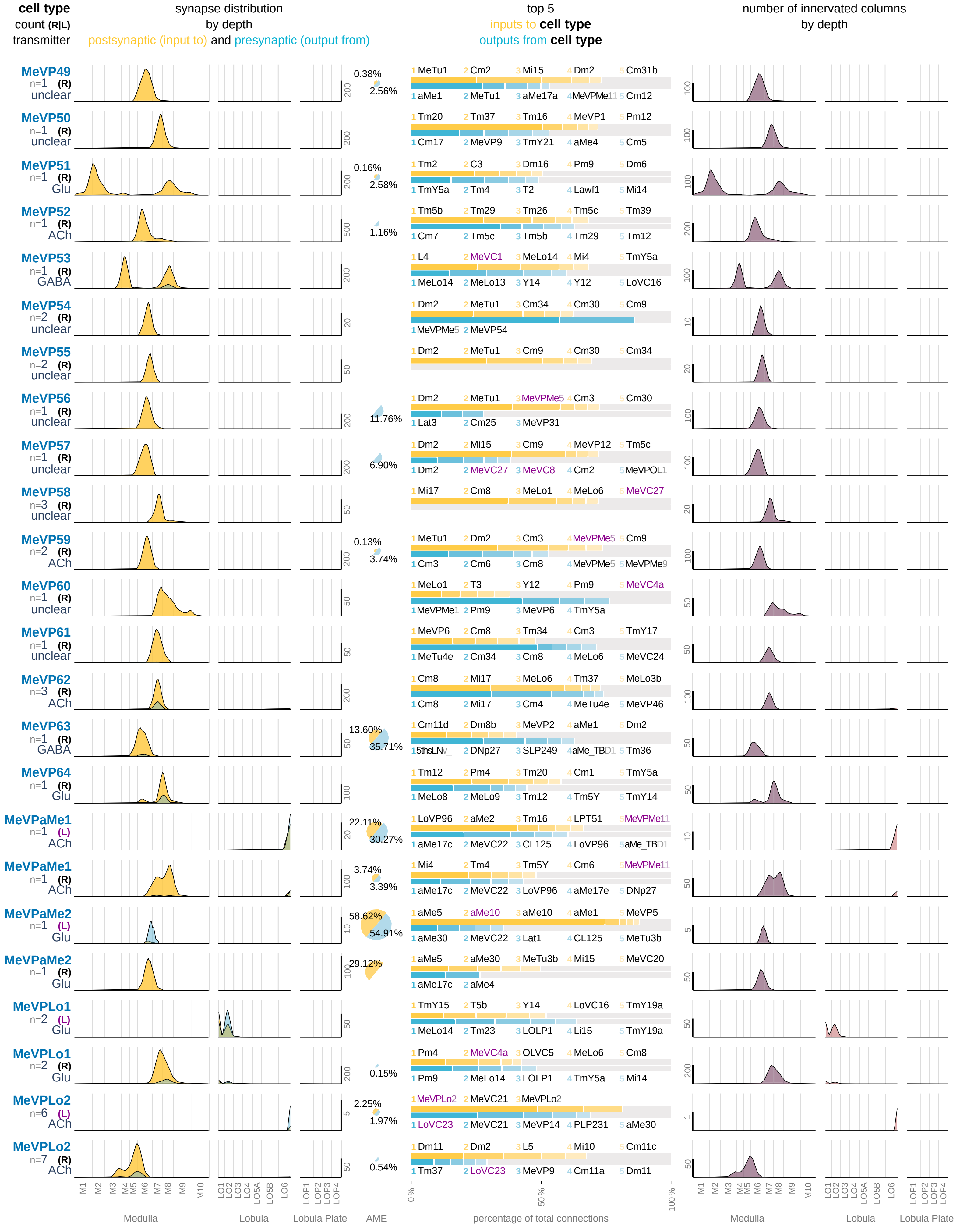

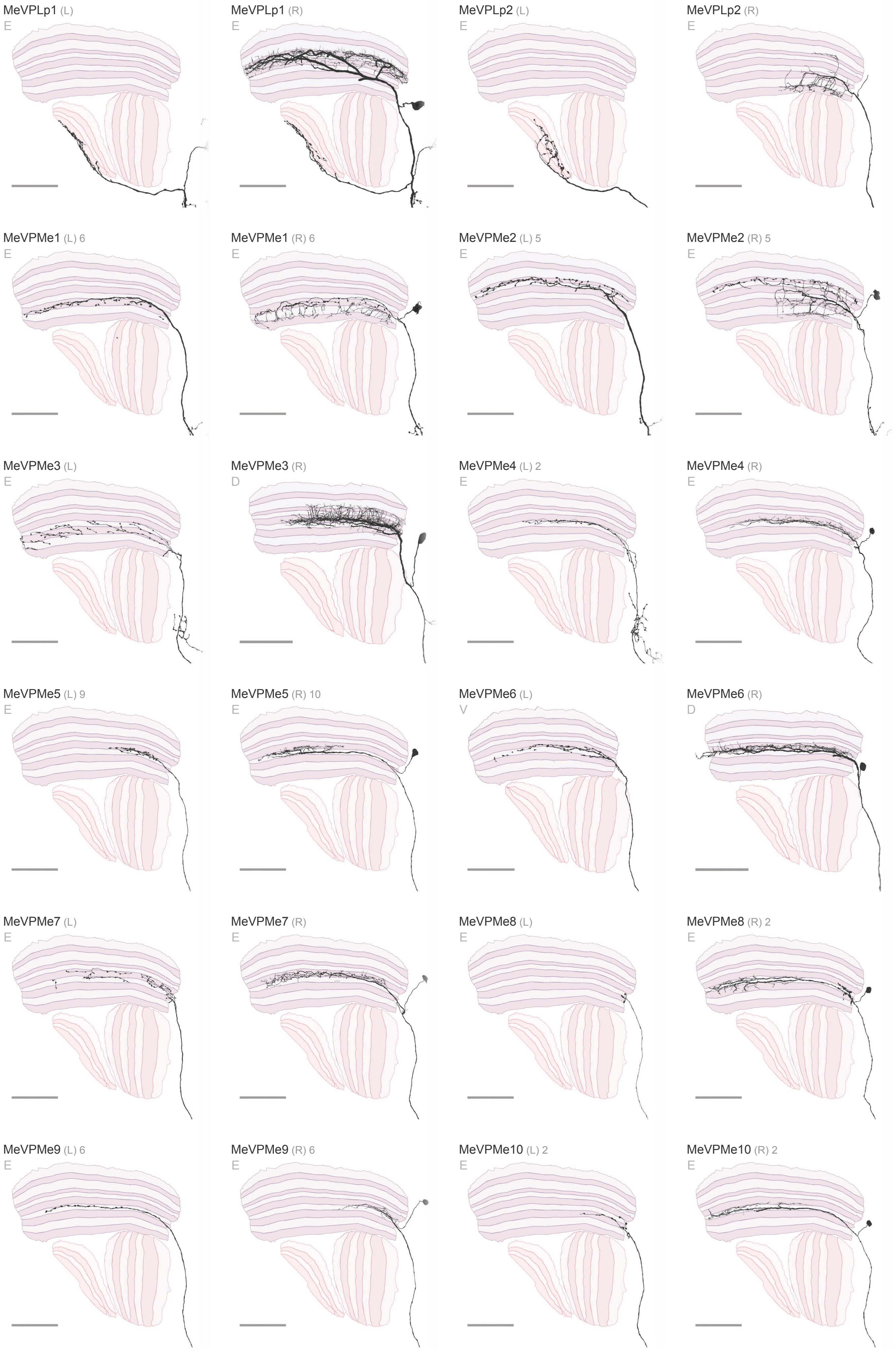
MeVPLp1 (L)
E
MeVPLp1 (R)
E
MeVPLp2 (L)
E
MeVPLp2 (R)
E
MeVPMe1 (L) 6
E
MeVPMe1 (R) 6
E
MeVPMe2 (L) 5
E
MeVPMe2 (R) 5
E
MeVPMe3 (L)
E
MeVPMe3 (R)
D
MeVPMe4 (L) 2
E
MeVPMe4 (R)
E
MeVPMe5 (L) 9
E
MeVPMe5 (R) 10
E
MeVPMe6 (L)
V
MeVPMe6 (R)
D
MeVPMe7 (L)
E
MeVPMe7 (R)
E
MeVPMe8 (L)
E
MeVPMe8 (R) 2
E
MeVPMe9 (L) 6
E
MeVPMe9 (R) 6
E
MeVPMe10 (L) 2
E
MeVPMe10 (R) 2
E

**Visual Projection Neurons** 15 / 16

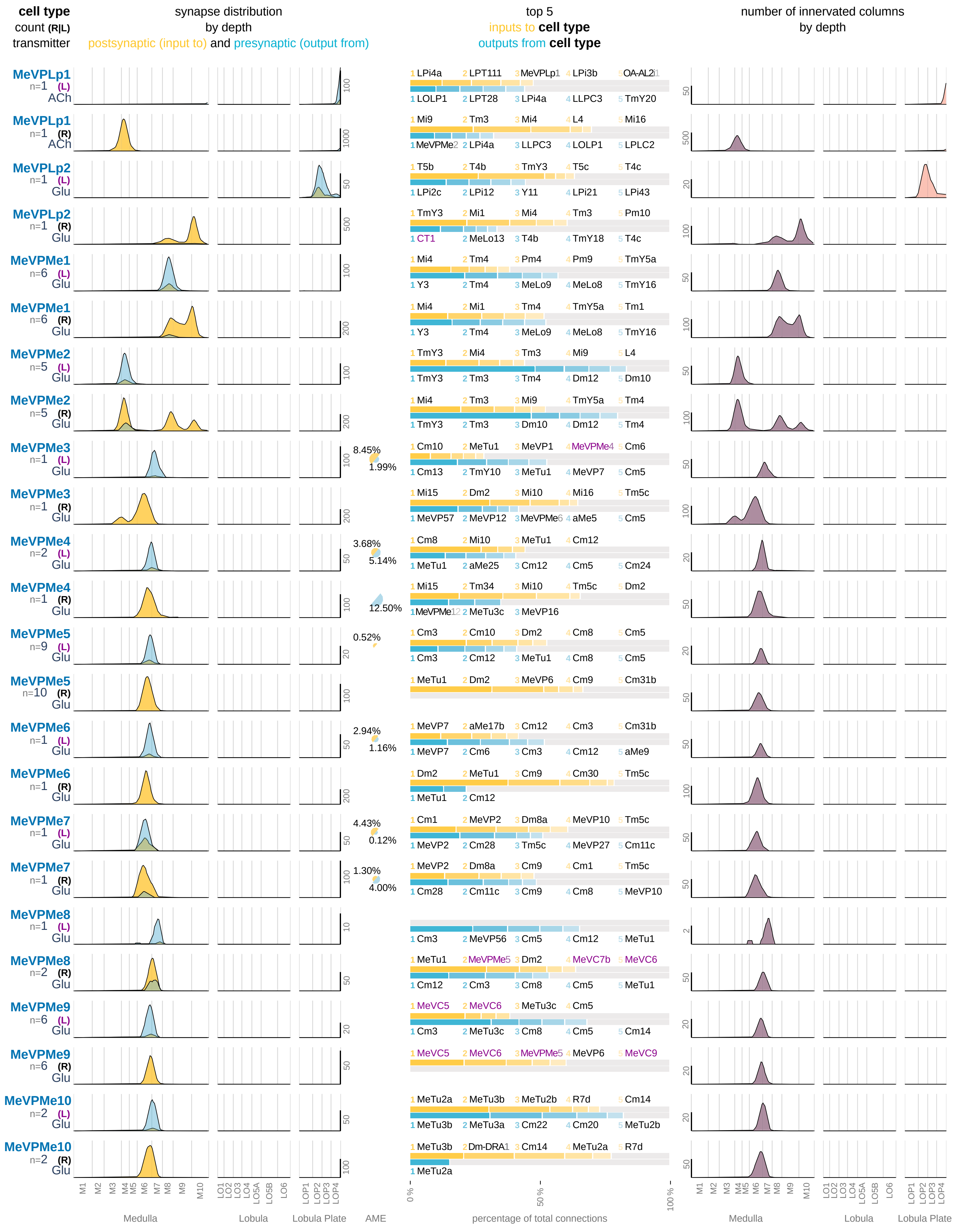

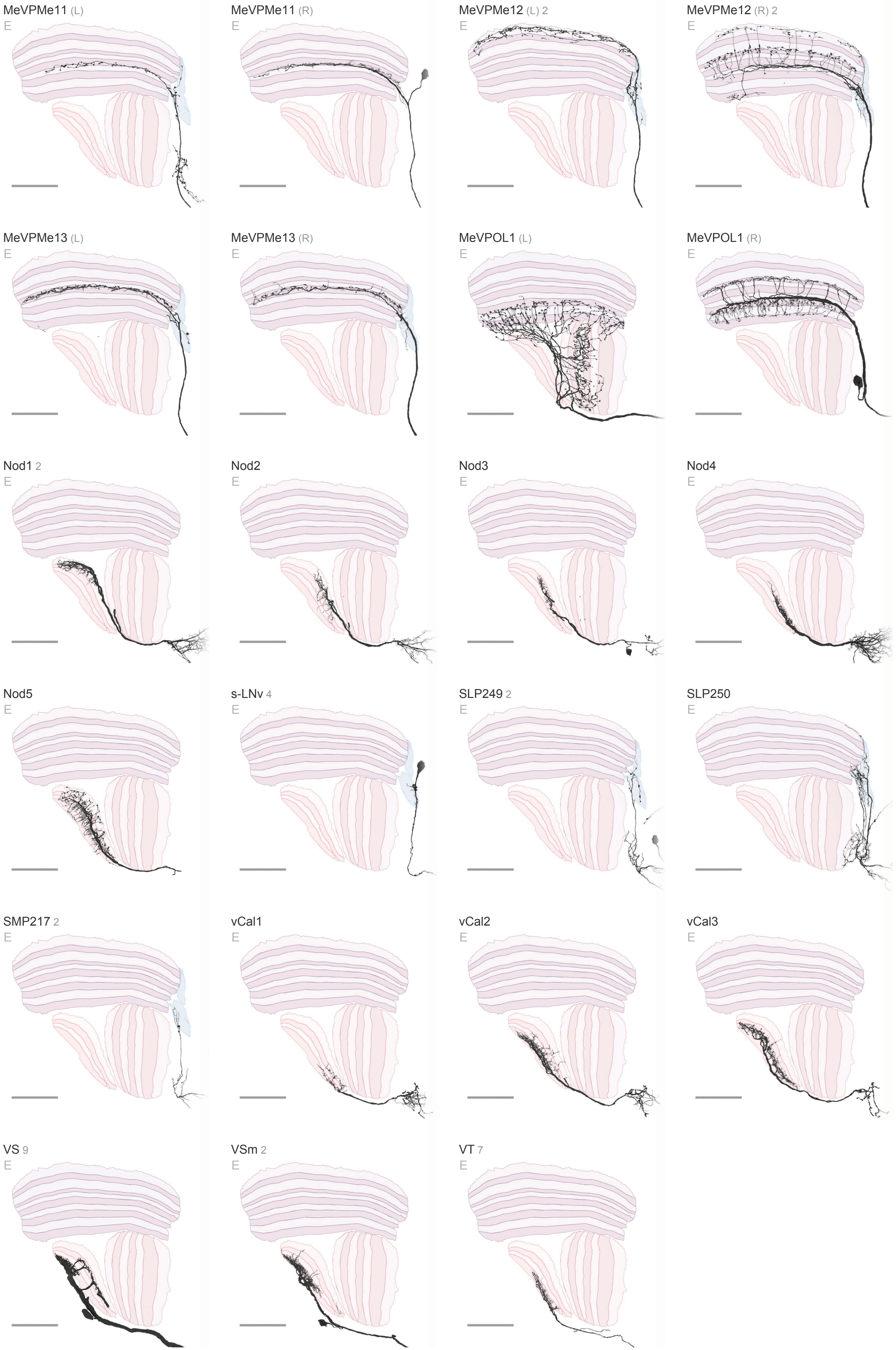
MeVPMe11 (L)
E
MeVPMe11 (R)
E
MeVPMe12 (L) 2
E
MeVPMe12 (R) 2
E
MeVPMe13 (L)
E
MeVPMe13 (R)
E
MeVPOL1 (L)
E
MeVPOL1 (R)
E
Nod1 2
E
Nod2
E
Nod3
E
Nod4
E
Nod5
E
s-LNv 4
E
SLP249 2
E
SLP250
E
SMP217 2
E
vCal1
E
vCal2
E
vCal3
E
VS 9
E
VSm 2
E
VT 7
E

**Visual Projection Neurons** 16 / 16

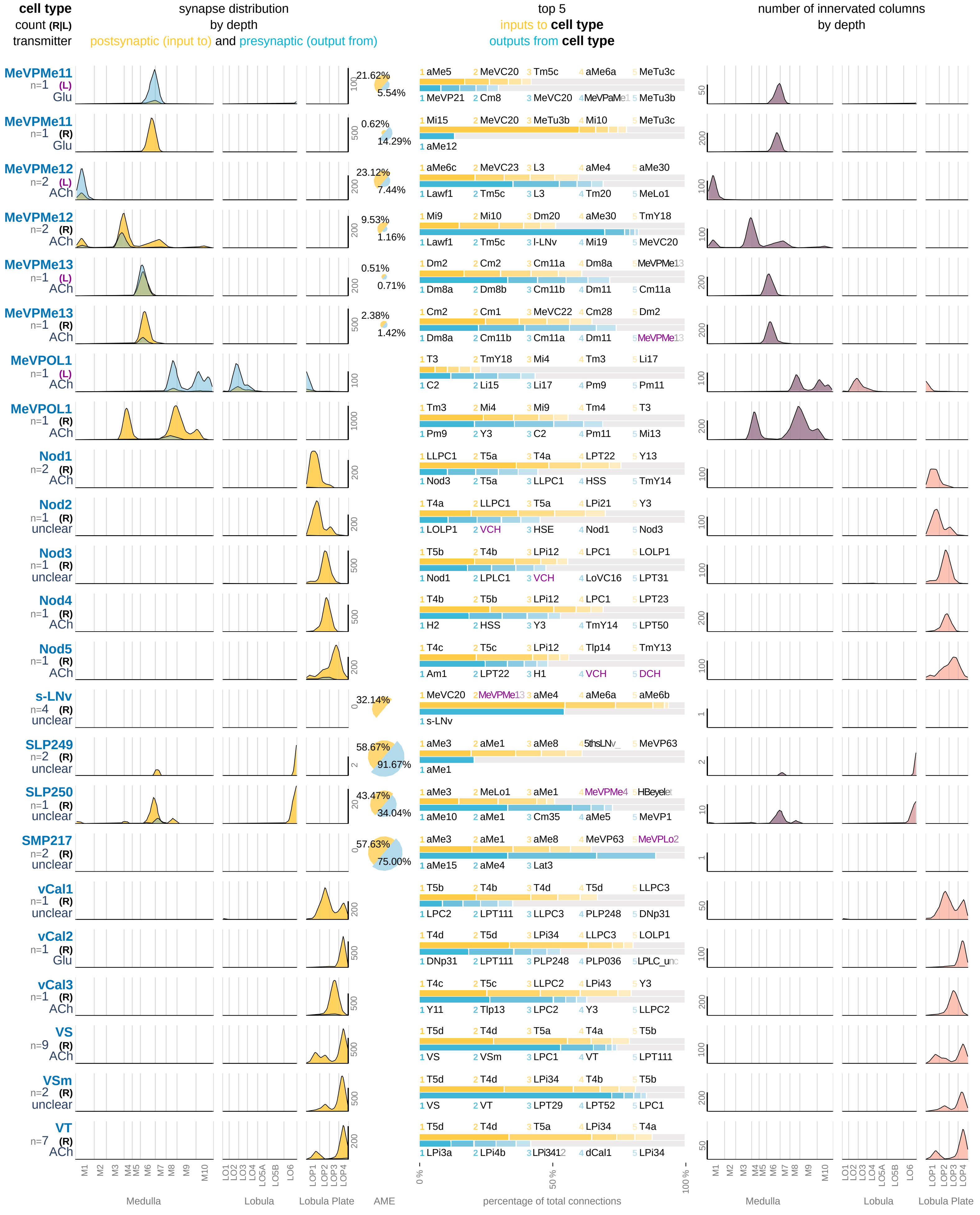

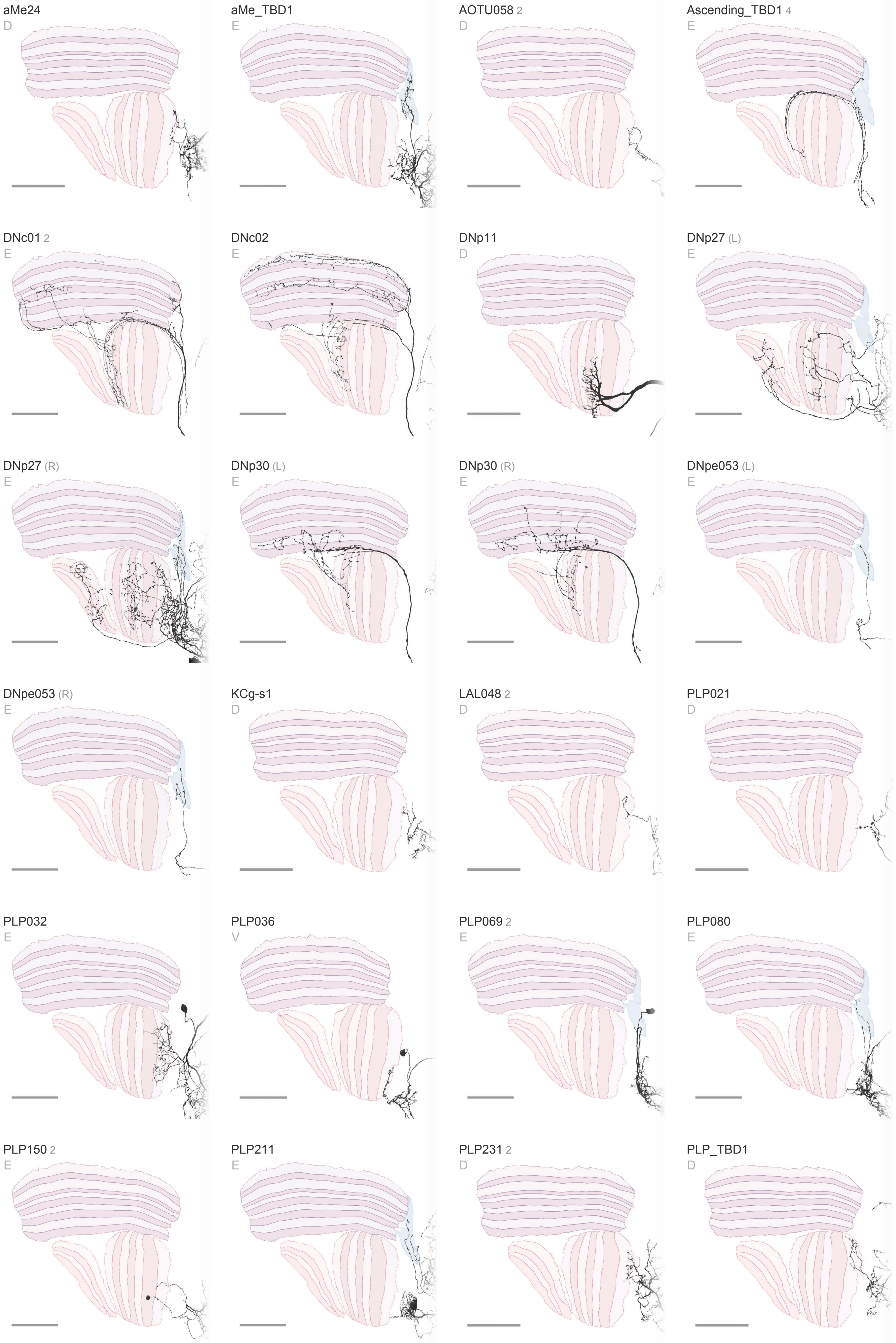
aMe24
D
aMe_TBD1
E
AOTU058 2
D
Ascending_TBD1 4
E
DNc01 2
E
DNc02
E
DNp11
D
DNp27 (L)
E
DNp27 (R)
E
DNp30 (L)
E
DNp30 (R)
E
DNpe053 (L)
E
DNpe053 (R)
E
KCg-s1
D
LAL048 2
D
PLP021
D
PLP032
E
PLP036
V
PLP069 2
E
PLP080
E
PLP150 2
E
PLP211
E
PLP231 2
D
PLP_TBD1
D

### Other Visual Neurons 1 / 2

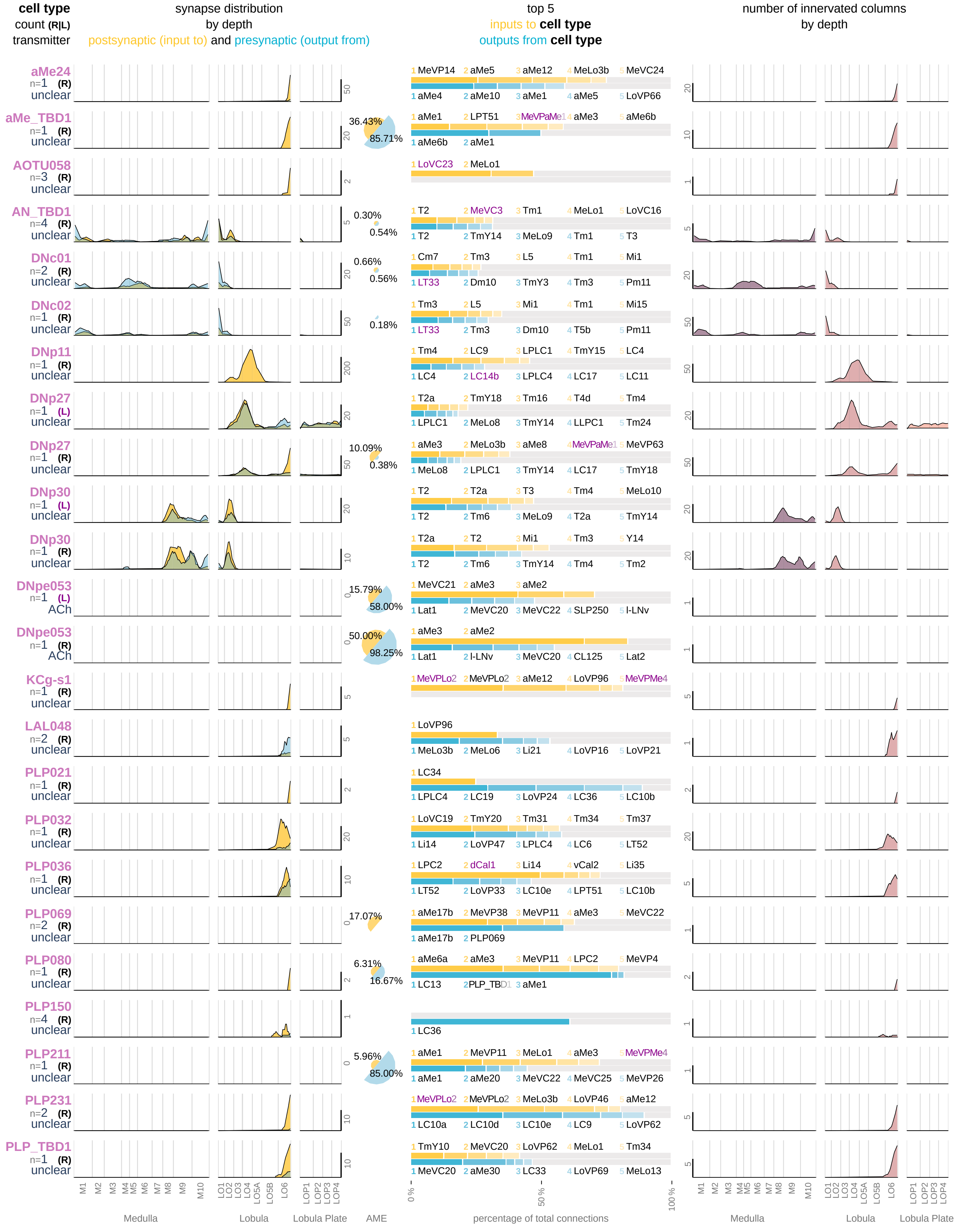

PS272 2
D
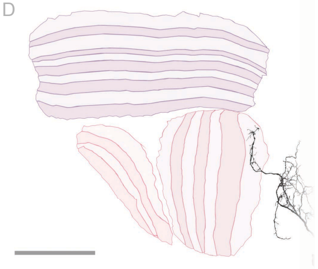

SLP359
D
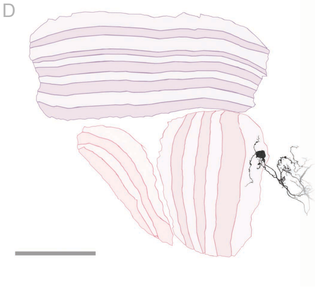

SMP200
E
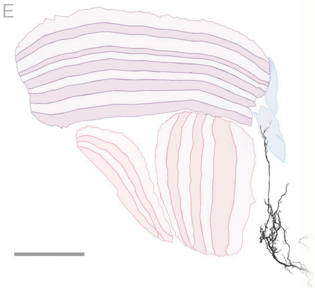

SMP528
D
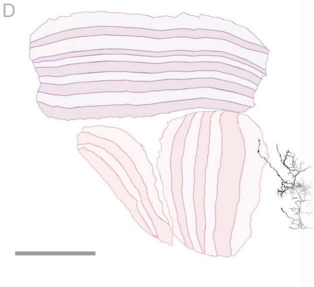

VLP_TBD1
V
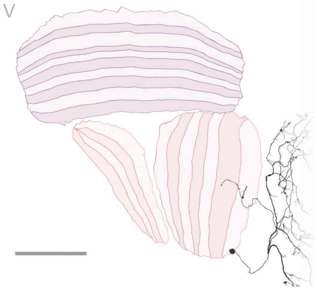

### Other Visual Neurons 2 / 2

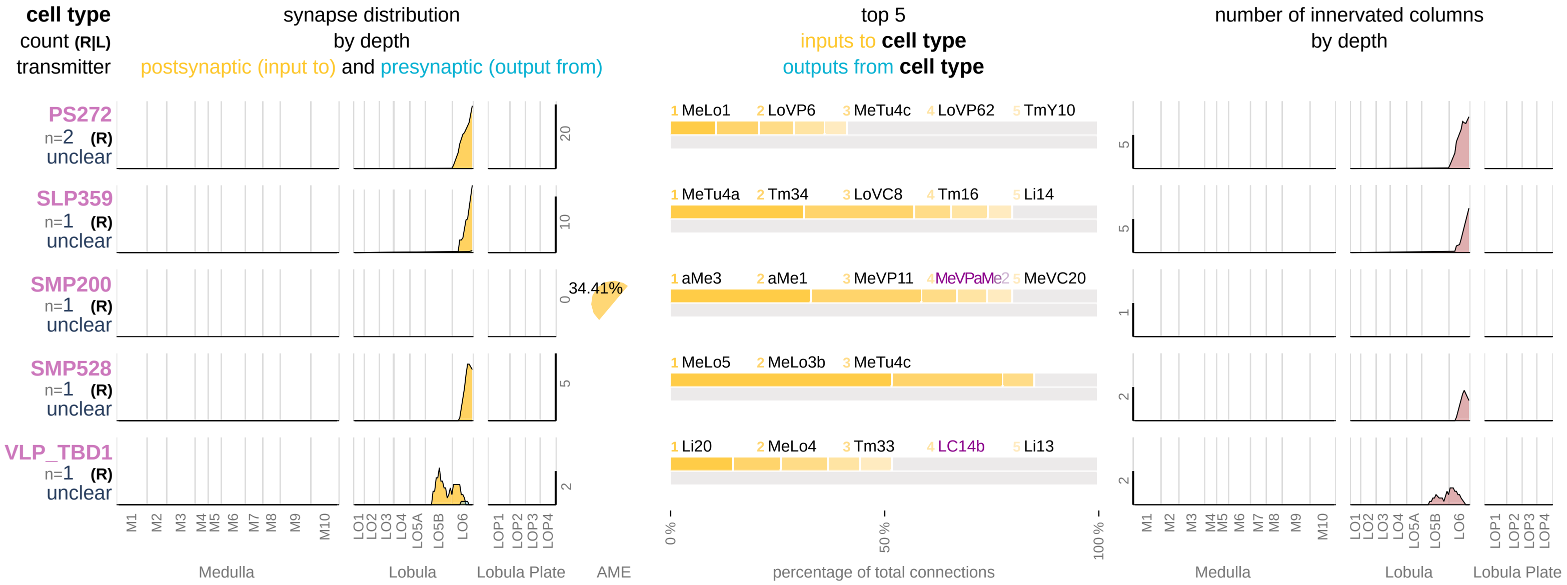
